# Tumour evolution as ground truth for cancer whole-genome sequencing

**DOI:** 10.64898/2026.06.09.731105

**Authors:** Lucrezia Valeriani, Giorgia Gandolfi, Elena Buscaroli, Katsiaryna Davydzenka, Giovanni Santacatterina, Alice Antonello, Azad Sadr, Virginia Anna Gazziero, Salvatore Milite, Elena Rivaroli, Anna Kabanova, Guido Sanguinetti, Alessio Ansuini, Leonardo Egidi, Stefano Cozzini, Alberto Cazzaniga, Giovanni Tonon, Trevor Graham, Andrea Sottoriva, Riccardo Bergamin, Nicola Calonaci, Alberto Casagrande, Giulio Caravagna

## Abstract

Cancer genomes are shaped by evolutionary processes that couple mutagenesis, clonal selection, chromosomal instability, spatial growth and treatment response into structured genomic patterns, yet current benchmarking strategies largely ignore this evolutionary dependency. Here, we present SCOUT, a large-scale synthetic whole-genome sequencing resource of over 200 samples, designed for systematic benchmarking of tumour genomic analysis and evolutionary inference under controlled evolutionary ground truth. Unlike conventional task-specific simulations, SCOUT models tumour evolution as a latent generative process that simultaneously shapes mutations, copy-number alterations, variant allele frequencies, mutational signatures and clonal architectures. SCOUT recapitulates key features of solid and haematological malignancies, including driver mutations, chromosomal instability, intratumour heterogeneity, spatial sampling and treatment-associated evolutionary dynamics in tumour and matched-normal longitudinal and multi-region sequencing designs. Using SCOUT, we benchmarked widely used methods for somatic variant detection, copy-number analysis, mutational signature inference and tumour evolutionary reconstruction. Across analytical tasks, performance deteriorated in low-purity, highly subclonal and structurally complex tumours, while spatial sampling bias and hypermutation generated spurious evolutionary signals that confounded tumour interpretation across multiple inference layers. Evolutionary simulations further distinguished lineage-restricted genetic bottlenecks from multi-lineage resistance dynamics associated with tumour plasticity. Tumour purity consistently exerted a stronger effect on inference accuracy than sequencing depth. Together, our results establish evolutionary ground truth as a prerequisite for reproducible benchmarking and biologically interpretable analysis of cancer whole-genome sequencing data.

## 1. Introduction

Tumour whole-genome sequencing (WGS) is being integrated into clinical and diagnostic workflows through national and institutional programmes [1–3]. Unlike targeted panels or whole-exome sequencing [4, 5], WGS captures the full spectrum of somatic and germline variation across coding and non-coding regions of the cancer genome [6], establishing tumour evolution as a key framework for interpreting intratumour heterogeneity (ITH), disease progression and therapeutic response [7, 8]. The adoption of WGS is shifting the main bottleneck from genome generation to genome interpretation [9], driving the development of computational methods for somatic variant detection, mutational signature analysis, clonal deconvolution and phylogenetic reconstruction [10–20]. However, unlike other areas of machine learning, cancer genomics lacks standardised datasets with known ground truth for reproducible benchmarking and systematic method evaluation [21]. Important community efforts have established reference datasets and benchmarking strategies for specific analytical tasks, including somatic mutation detection[22] and subclonal reconstruction [22]. Yet these approaches generally model isolated components of cancer genome analysis rather than the evolutionary processes that jointly shape mutagenesis, copy-number change, clonal dynamics and treatment response. In cancer genomes, these observables are not independent: they emerge from shared evolutionary histories and are therefore coupled across multiple layers of genomic data. As a result, simplified simulations and task-specific spike-in strategies cannot fully capture the nonlinear dependencies that confound tumour genome interpretation in real sequencing studies. Benchmarking therefore still frequently relies on heterogeneous cohorts, partial orthogonal validation or simplified simulations, limiting reproducibility and complicating the definition of performance standards for research and clinical applications.

The missing ground truth in cancer genomics is inherently evolutionary [23]. Somatic mutations, copy-number alterations and structural variants arise through stochastic processes of tumour growth, selection, clonal expansion and treatment response, and are subsequently observed through imperfect sampling and sequencing. These evolutionary processes generate correlated genomic observables whose interpretation depends on latent clonal structure, spatial growth dynamics and temporal ordering. Realistic benchmark datasets therefore require more than the insertion of predefined variants into a reference genome: they must jointly model tumour evolution, mutational processes and sequencing measurement. Population-genetic models provide a principled framework for simulating complex selective regimes [24], but are rarely integrated with realistic models of cancer mutagenesis, genome alteration and sequencing at the scale required for WGS benchmarking [25].

Here, we present a synthetic short-read WGS cohort designed as a reference resource for benchmarking tumour evolution and cancer genomics pipelines. The cohort spans multiple solid and haematological malignancies and captures clinically relevant evolutionary scenarios, including distinct sampling strategies as observed in real cohorts [26, 27], patterns of ITH and chromosomal instability [28, 29], endogenous and exogenous mutagenesis [30, 31], and acquired drug resistance [28, 32, 33]. To generate and analyse the cohort, we developed a programmable spatial tumour evolution framework that models tumour evolution as a latent generative process linking clonal dynamics, genome instability, mutagenesis and sequencing observables within an end-to-end bioinformatics workflow for the systematic evaluation of cancer genomics pipelines.

The resulting resource comprises more than 27 terabytes of synthetic WGS data across multiple tumour types, sequencing depths and sample purities, with complete access to the underlying evolutionary and genomic ground truth. We establish quantitative performance baselines for widely used analytical pipelines and identify major sources of variability across WGS analysis workflows. This benchmark provides a controlled framework for the development, comparison and validation of computational methods in cancer genomics, enabling more reproducible evaluation and clinical translation of WGS analyses.

## 2. Results

### 2.1 A simulated cohort of universal tumours (SCOUT) for benchmarking WGS analysis

We generated SCOUT, a synthetic WGS cohort comprising seven tumour models (five solid and two haematological) with matched tumour–normal sequencing across longitudinal, spatial and treatment-associated evolutionary scenarios. To generate the cohort, we developed ProCESS (Programmable Cancer Evolution Spatial Simulator; Methods), a new framework that integrates spatial population-genetic tumour evolution with synthetic short-read WGS generation (Fig. 1A, Extended Data Fig. 1, Supplementary Fig. S1 to S11, Supplementary Tables T1 to T6). Starting from 1000 Genome Project [34] germline genomes, ProCESS jointly models clonal expansion, cancer mutagenesis, treatment dynamics and sequencing noise, enabling complete evolutionary ground truth to be tracked from mutational origin to paired-end FASTQ generation (Supplementary Fig. S12).

**Figure 1.**
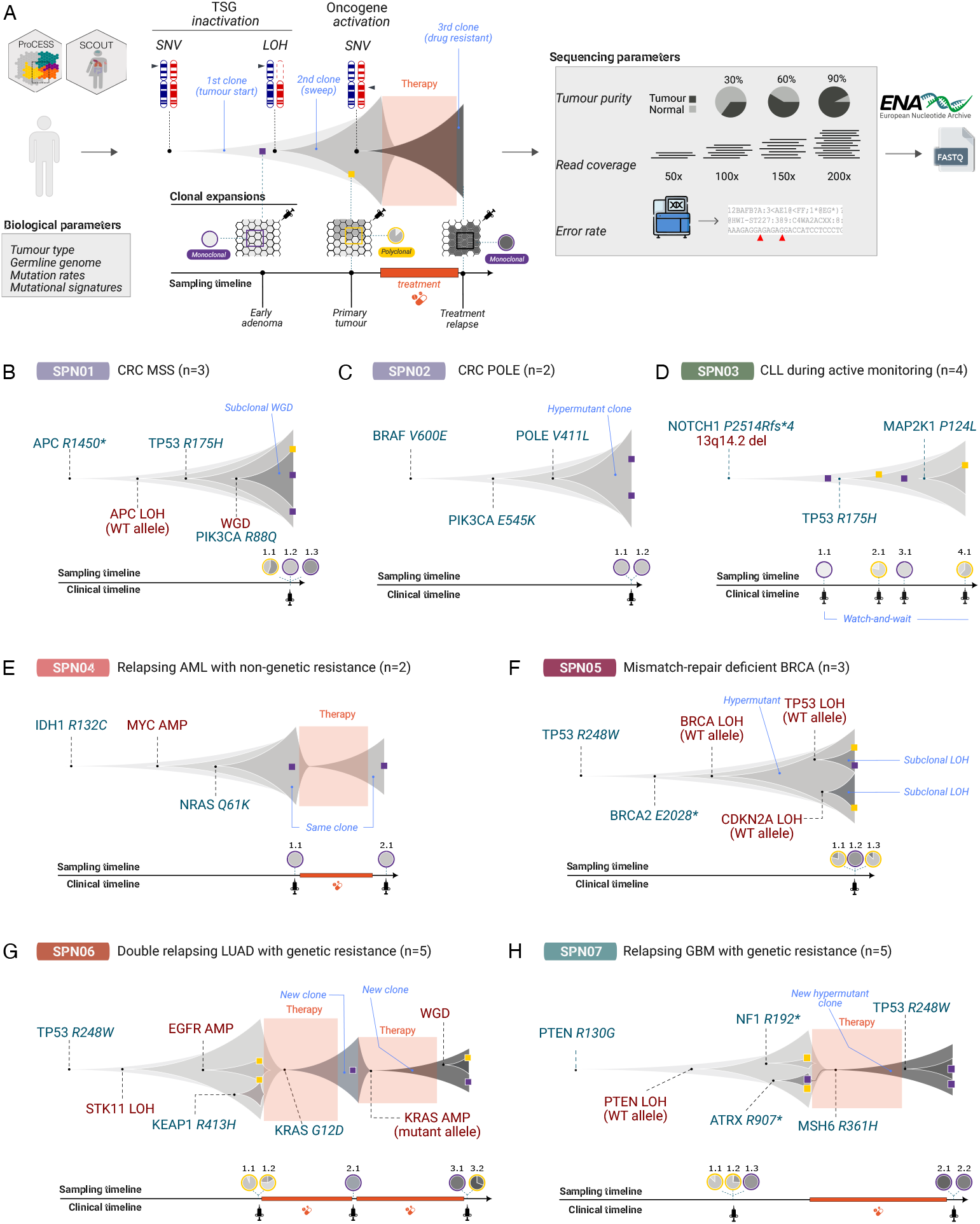
The Simulated Cohort of Universal Tumours (SCOUT). (A) Overview of the SCOUT framework. Starting from a real germline genome and tumour-specific biological parameters, clonal evolution is simulated using spatial population genetics models augmented with mutational processes and treatment dynamics. Synthetic tumour-normal pairs are sampled using multi-region and longitudinal designs, and Illumina short-read data are generated across combinations of sequencing coverage and tumour purity. FASTQ files are produced for benchmarking downstream analyses. (B–H) Clonal evolution models of the seven SCOUT tumours (SPN01–SPN07); *n* denotes the number of samples per tumour. SPN01 and SPN02 are colorectal cancers, respectively microsatellite-stable and POLE-mutant hypermutant cases, with multi-region sampling. SPN03 is an untreated chronic lymphocytic leukaemia followed longitudinally during active monitoring. SPN04 is an acute myeloid leukaemia developing non-genetic resistance to chemotherapy. SPN05 is a mismatch-repair-deficient breast cancer with multi-region sampling. SPN06 is a lung adenocarcinoma developing KRAS-associated resistance following two rounds of chemotherapy, with longitudinal and multi-region sampling. SPN07 is a glioblastoma developing MSH6-associated resistance to temozolomide. Driver events, sampling times and treatments are annotated along the evolutionary lineages; all relapsing tumours are sampled before and after treatment. Together, the tumours recapitulate diverse evolutionary scenarios, including monoclonal and polyclonal samples, aneuploidy, distinct mutational processes, treatment response and relapse dynamics.

Using ProCESS, we constructed tumour prototypes (Fig. 1B–H and Extended Data Fig. 2) of colorectal cancers (SPN01–02), chronic lymphocytic leukaemia (SPN03), acute myeloid leukaemia (SPN04), breast cancer (SPN05), lung adenocarcinoma (SPN06) and glioblastoma (SPN07), (Supplementary Fig. S13 to S19). These tumours recapitulate distinct evolutionary and clinical trajectories characteristic of the corresponding cancer types [35–51], including polyclonality, clonal sweeps and treatment resistance measured through distinct combinations of longitudinal and multi-region sampling (Supplementary Table T7 and T8A-I). Importantly, these evolutionary scenarios generate coordinated patterns across multiple genomic layers, linking mutational processes, copy-number dynamics, clonal composition and spatial structure within the same simulated tumours.

Across the cohort, we simulated 30 evolutionary clones sampled through 24 biopsies, with 7 matched normals. For each biopsy, matched tumour–normal WGS data were generated on an Illumina NovaSeq 6000 paired-end platform across three sequencing depths (50×, 100× and 150×) and three tumour purity levels (30%, 60% and 90%), generating nine sequencing configurations per sample (with reference genome GRCh38). In total, SCOUT comprises 223 FASTQ files, including 216 from tumour and 7 from matched-normal samples (total 223 samples), corresponding to approximately 27 terabytes of raw sequencing data (Extended Data Fig. 3). All data are publicly available through the European Nucleotide Archive, together with processed outputs from all analyses provided alongside this study (Data Availability). This design enables systematic benchmarking of analytical robustness across key sources of variability in clinical cancer genomics, including sequencing depth, tumour purity and sampling strategy.

The SCOUT genomes recapitulated major genomic features of human cancers. Tumour mutational burden (TMB; Fig.2A) distributions matched ranges observed in large cohorts [3, 52], including hypermutated profiles associated with homologous recombination deficiency (HRd) and mismatch repair deficiency (MMRd). Driver landscapes comprised single-nucleotide variants (SNVs), insertions and deletions (indels), and allele-specific copy-number alterations (CNAs) affecting canonical suppressor genes and oncogenes (Fig.2B), including APC, TP53, BRCA2, NOTCH1, KRAS, PIK3CA, and IDH1. The cohort also contained focal and arm-level CNAs, including EGFR and MYC amplifications and deletions of CDKN2A and 13q14.2.

Endogenous and exogenous mutational processes (Fig.2B) were modelled from reference signatures [53, 54], capturing ageing (SBS1), APOBEC activity (SBS2 and SBS13), tobacco exposure (SBS4), HRd (SBS3), MMRd (SBS26), as well as mutagenesis associated with exposure to platinum (SBS35) and alkylating agents (SBS11) [14, 55–61] (Supplementary Fig. S20 and S21). The simulated mutational processes generated realistic clonal and subclonal ITH, including, for example, POLE-associated hypermutators in colorectal cancers, and HRd in BRCA-deficient breast cancers.

SCOUT reproduced diverse configurations of chromosomal instability, including whole-genome doubling (WGD) and subclonal CNAs, at various levels of fractions of genome altered (FGA) [62–66]. Recurrent arm-level gains on chromosomes 7p, 8q and 12p, and deletions on 5q, 10q, 13q, 17p, 17q and 19p emerged from explicitly simulated evolutionary dynamics rather than from predefined mutational templates (Fig. 2C), enabling chromosomal instability, clonal selection and mutational processes to co-evolve within the same tumours.

**Fig. 2:**
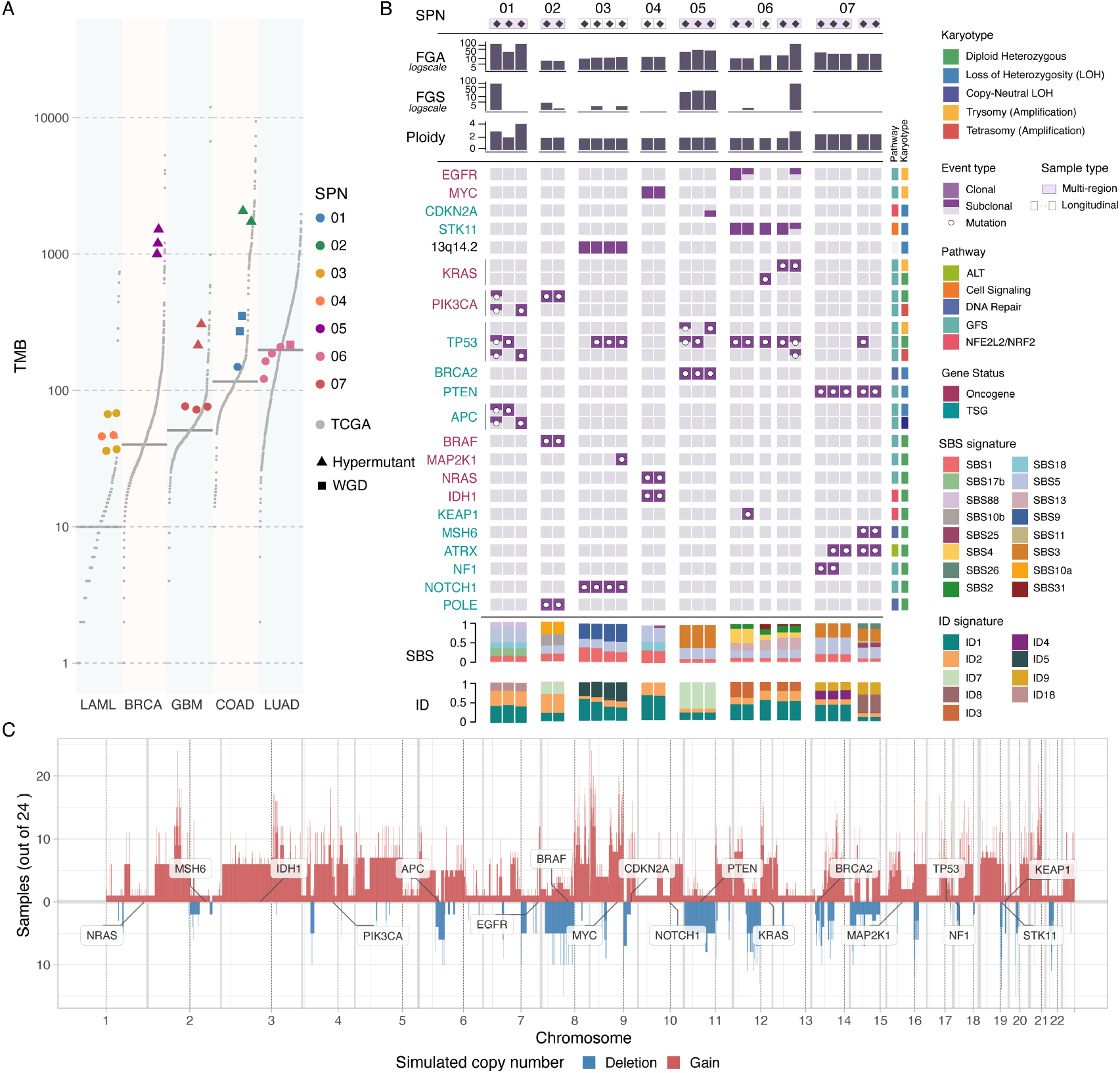
Genomic features of the SCOUT cohort. (A) Tumour mutational burden (TMB) across SCOUT samples compared with corresponding tumour types from The Cancer Genome Atlas (TCGA) [52]. SCOUT tumours span the range of mutational burdens observed in human cancers, including hypermutant profiles associated with complex mutagenesis (mismatch repair and homologous recombination deficiency). (B) Integrated genomic profiles of SCOUT tumours summarising key genomic features across samples, including whole-genome doubling (WGD), fraction of genome altered (FGA), fraction of genome with subclonal copy-number segments (FGS), tumour ploidy, mutational signatures, and driver alterations in canonical cancer genes. Driver SNVs and indels are annotated by gene class and pathway, illustrating the diversity of genomic architectures represented across the cohort. (C) Genome-wide allele-specific CNAs of SCOUT samples, showing gains and losses events across chromosomes. Recurrently altered cancer driver genes are annotated according to their genomic position.

These properties establish SCOUT as an evolutionarily coherent WGS benchmark in which evolutionary history, clonal composition, mutational processes and sampling structure are jointly specified rather than independently simulated. By preserving the coupling between tumour evolutionary dynamics and genomic observables, SCOUT enables rigorous evaluation of tumour somatic profiling, mutational signature inference, subclonal deconvolution and tumour evolution reconstruction from bulk WGS data under controlled evolutionary ground truth.

### 2.2. Molecular profiling with the nf-core/sarek bioinformatics pipeline

We used SCOUT to benchmark tumour–normal WGS workflows implemented in the nf-core/sarek pipeline (Methods), a widely adopted community-maintained framework for somatic variant detection [67, 68] built on the Genome Analysis Toolkit (GATK) [69, 70]. Using controlled combinations of sequencing depth, tumour purity and clonal architecture, we quantified the performance of somatic variant and CNA calling under realistic tumour evolutionary scenarios (Fig. 3A). Because tumour purity, subclonality and chromosomal instability emerge from the same underlying evolutionary dynamics, this framework enables systematic evaluation of how evolutionary complexity propagates across multiple layers of genome interpretation.

**Fig. 3:**
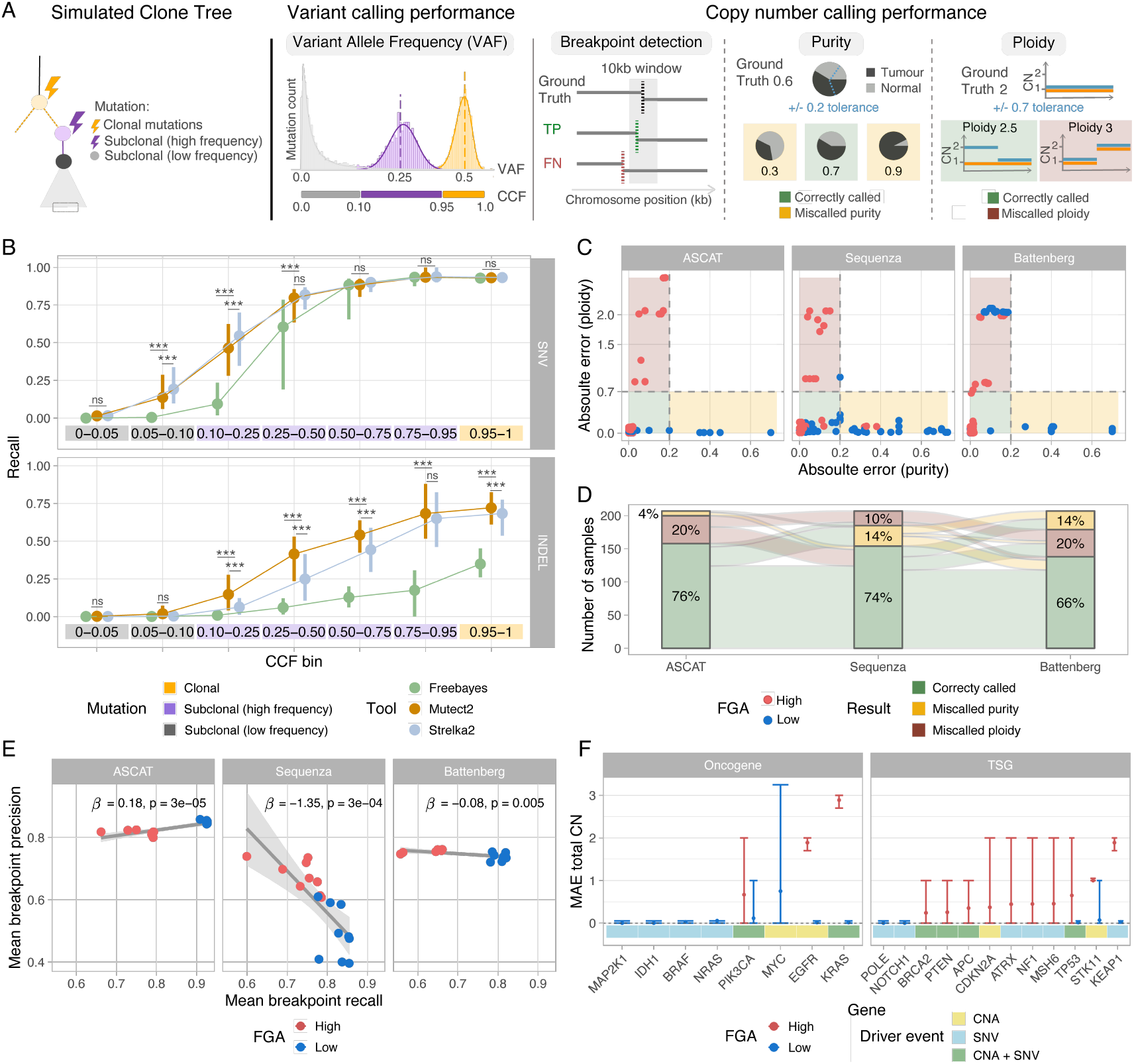
Benchmarking tumour molecular profiling with nf-core/sarek. (A) Given a simulated clone tree, we benchmark variant calling and copy number calling performance by comparing ProCESS simulated ground truth against nf-core/sarek results across four task: variant allele frequency (VAF) calling, breakpoint detection, and purity and ploidy inference. (B) Recall of somatic variant detection as a function of cancer cell fraction (CCF), stratified by mutation class (SNVs and indels) and variant caller (FreeBayes, Mutect2 and Strelka2). Variants are grouped into clonal and subclonal frequency bins, illustrating how detection recall depends on allelic prevalence. (C,D) Accuracy of CNA callers (ASCAT, Sequenza and Battenberg) in estimating tumour purity and ploidy (C). Absolute errors are shown for samples stratified by fraction of genome altered (FGA), highlighting differences in performance between low- and high-FGA tumours. Summary comparative plots are reported in panel (D). (E) Breakpoint detection performance of CNA callers, showing mean recall and precision across samples. Results are stratified by FGA. (F) Accuracy of CNA detection for SCOUT driver mutations, measured as mean absolute error (MAE) of total copy number (CN) estimate.

We first evaluated somatic variant detection recall as a function of cancer cell fraction (CCF), defined as the allelic prevalence of mutations after accounting for tumour purity and copy number (Supplementary Fig. S22, Supplementary Table T9A). Clonal mutations were consistently detected with high recall (0.87 ± 0.33), whereas subclonal variants markedly reduced sensitivity. With a practical detection threshold of recall < 0.5, all callers failed to detect variants below approximately 25% CCF. Above this threshold, subclonal recall was 0.67 ± 0.60, with Mutect2 [71] (0.75 ± 0.49) and Strelka2 [72] (0.73 ± 0.39) performing comparably and consistently outperforming FreeBayes [73] (0.33 ± 0.76) (Fig. 3B).

Tumour purity was the dominant determinant of recall variation, substantially outweighing the effect of sequencing depth. At 90% purity, SNVs remained detectable down to ~ 10% CCF, whereas indel detection was substantially less sensitive, especially below 50% CCF. Reducing purity to 60% and 30% progressively shifted these limits upward, and indel detection failed to achieve reliable recall even for clonal variants at the lowest purity. Increasing tumour purity improved recall by approximately 15%, compared with only ~ 7% gained from increasing sequencing depth (Extended Data Fig. 4). Across all callers, indels remained consistently more difficult to detect than SNVs at matched CCFs, highlighting a persistent limitation of current short-read approaches. We benchmarked the performance of three germline variant callers, FreeBayes [74], Strelka2 [72], and HaplotypeCaller [12], for the detection of single nucleotide polymorphisms (SNPs) in the simulated normal sample. All methods achieved high F1-scores (0.988–0.998), although FreeBayes showed comparatively lower performance (Extended Data Fig. 4, Supplementary Fig. S23, Supplementary Table T9B).

We next assessed CNA calling using ASCAT [75], Sequenza [76] and Battenberg [20] (Supplementary Fig. S24, Supplementary Table T9C). Accurate CNA inference requires joint estimation of tumour purity, ploidy and segment breakpoints, making copy-number reconstruction intrinsically underconstrained.

CNA calling performance depended strongly on tumour genomic architecture (Extended Data Fig. 5). Tumours with low FGA (≤ 15%) showed reduced accuracy in purity estimation, consistent with insufficient copy-number signal. Conversely, tumours with high FGA enabled more accurate purity estimation but introduced ambiguity in ploidy inference owing to increased genomic complexity (Fig. 3C,D). These results reveal an inherent trade-off between signal availability and model identifiability: genomes with limited CNAs lack sufficient information for robust inference, whereas highly rearranged genomes challenge accurate reconstruction. Importantly, these failure modes arise from distinct evolutionary regimes, indicating that tumour evolutionary dynamics directly constrain the interpretability of bulk WGS data. Consistent with this interpretation, breakpoint recall decreased in high-FGA tumours (mean ~ 0.71) compared with low-FGA tumours (~ 0.84), while precision remained relatively stable across methods, except for Sequenza, which showed reduced precision in low-FGA genomes (Fig. 3E). Subclonal copy-number callers were similarly affected: Battenberg performed worst in highly subclonal genomes (high-FGS, mean ~ 0.324) relative to those with fewer subclonal CNAs (low-FGS, ~ 0.537, Extended Data Fig. 4). Overall ASCAT achieved the best balance between precision and recall in breakpoint detection, followed by Sequenza and Battenberg, with purity and ploidy jointly inferred within acceptable error bounds in approximately 76% of samples. Despite these genome-wide limitations, copy-number states at simulated driver loci were inferred with high accuracy (0.83, Fig. 3F). Copy-number states were inferred accurately in low-FGA tumours (mean error 0.08), with most errors confined to genomes with high FGA and WGD (mean error 0.87, Extended Data Fig. 5).

Together, these results show that current state-of-the-art WGS workflows for molecular profiling remain fundamentally constrained by tumour purity, subclonal architecture and genome complexity. Although they achieve high accuracy for clonal mutations and major CNAs, performance deteriorates systematically in low-purity, highly subclonal and structurally complex tumours. Importantly, these sources of variability are coupled consequences of tumour evolutionary dynamics that propagate molecular layers. By providing a controlled setting with known evolutionary ground truth, SCOUT enables quantitative delineation of these failure regimes and their biological determinants.

### 2.3. Tumour evolutionary inference with the nf-core/tumourevo machine learning pipeline

We developed nf-core/tumourevo, the first nf-core pipeline dedicated to tumour evolution inference from bulk DNA sequencing data (Methods and Extended Data Fig. 6). Built to operate downstream of variant-calling workflows such as nf-core/sarek, nf-core/tumourevo unifies quality control, driver annotation, mutational signature analysis and subclonal reconstruction within a standardised and reproducible framework (Supplementary Fig. S25). Because these analytical layers reflect coupled consequences of tumour evolution, nf-core/tumourevo enables joint interpretation of mutagenesis, clonal dynamics and treatment-associated evolutionary processes from bulk WGS data.

Prior to evolutionary analysis, quality control of identified somatic mutations and relative inferred copy number profiles was performed using CNAqc [77], while normal contamination was assessed using TINC [78]. Across all combinations of tumour purity and sequencing coverage, CNAqc assigned 84.7% of samples as QC PASS, with the remainder classified as QC FAIL. Among QC FAIL samples, 44.4% and 21.4% were associated with incorrect purity and ploidy estimates, respectively. However, some inaccurate CNA calls were still classified as QC PASS, typically in the presence of sampling bias or undetectable subclones that generate confounding VAF peaks and obscure underlying CNA inference errors (Supplementary Fig. S26, Supplementary Table T9E). In addition, Tumour-In-Normal (TIN) contamination by TINC was correctly inferred to be negligible, with a cohort-wide mean estimate of 0.00156 ± 0.0129 (Supplementary Fig. S27, Supplementary Table T9F).

Driver somatic mutations were identified by integrating functional impact predictions [79] with tumour-type-specific annotations from the Cancer Gene Census [80]. The pipeline achieved consistently high accuracy across non-hypermutant tumours (SPN01, 04, 06 and 07; median F1 = 0.8, range 0.65 − 1; Fig. 4A, Supplementary Fig. S28, Supplementary Table T9D). In contrast, performance deteriorated substantially in hypermutant samples (SPN02, 05 and 07; median F1 = 0.3), primarily owing to false positive driver calls. This reduction was driven by elevated TMB (median log10-TMB 3.04 versus 2.2), which increased the probability of passenger mutations occurring within cancer genes and consequently reduced the precision of annotation-based approaches (median precision 0.18 versus 0.75).

**Fig. 4:**
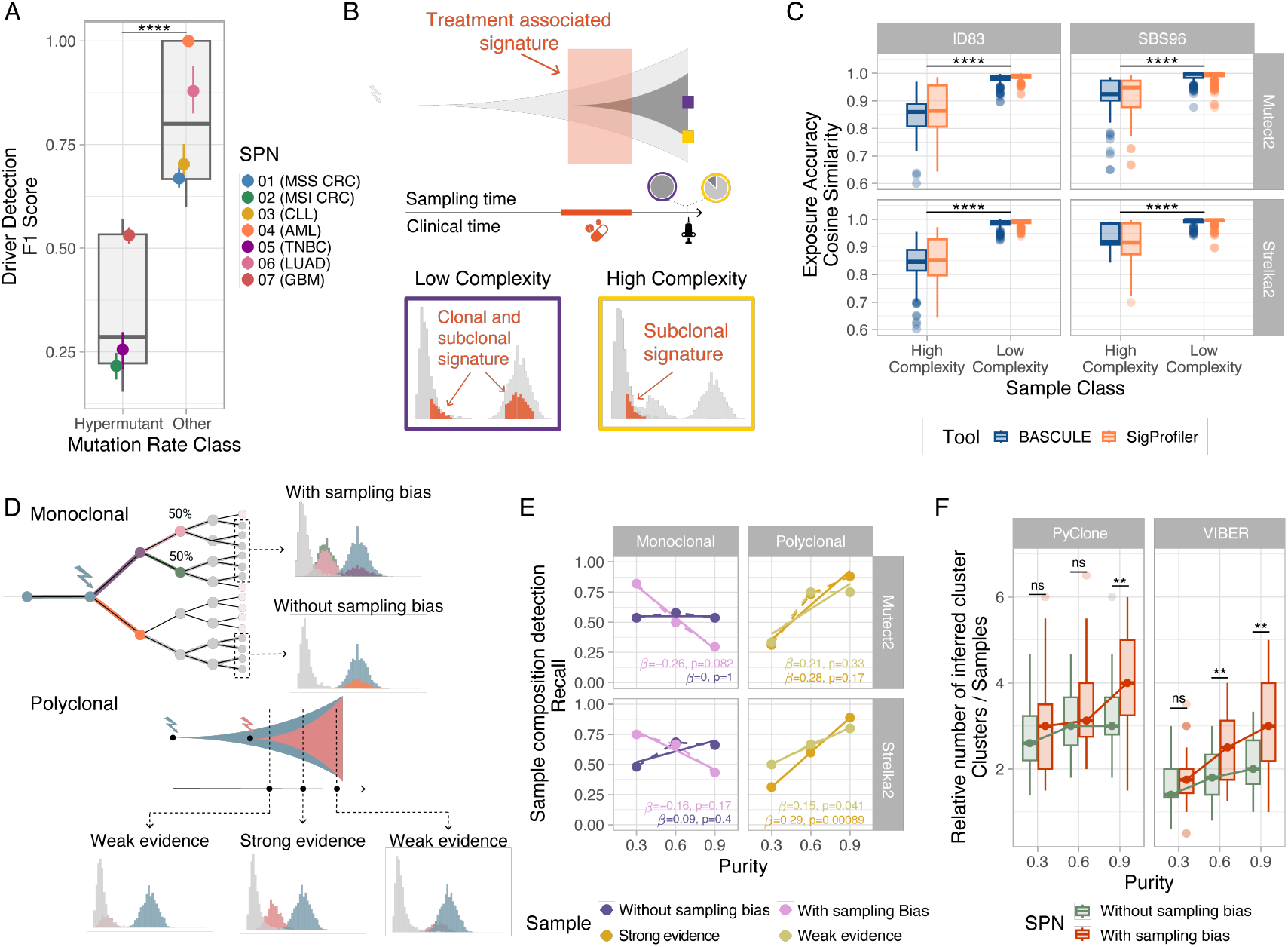
Benchmarking the nf-core/tumourevo pipeline with SCOUT. (A) F1 score for driver mutation detection across SCOUT tumours, stratified by TMB (hypermutant versus non-hypermutant tumours). Each point represents a single tumour. (B) Schematic of clone-specific mutational processes and their effects on mutational signature detection. Signatures associated with low-frequency subclones can be obscured by dominant clonal processes. (C) Cosine similarity of mutational signature deconvolution obtained with BASCULE and SigProfiler, stratified by signature class (SBS96 and ID83), sample complexity (high and low), and somatic variant caller (Mutect2 and Strelka2). (D) Schematic representation of monoclonal and polyclonal samples in the presence of spatial sampling bias or subclonal populations. Spatial sampling bias can generate multimodal VAF distributions in the absence of ongoing subclonal expansions. (E) Recall of monoclonal and polyclonal composition detection with MOBSTER following variant calling with Mutect2 or Strelka2 across tumour purity levels (0.3, 0.6 and 0.9), stratified by sample clonality and confounding factor (sampling bias or weak subclonal evidence). (F) Number of clusters inferred by PyClone and VIBER, normalised by the number of samples, across tumour purity levels (0.3, 0.6 and 0.9), stratified by spatial sampling bias.

**Fig. 5:**
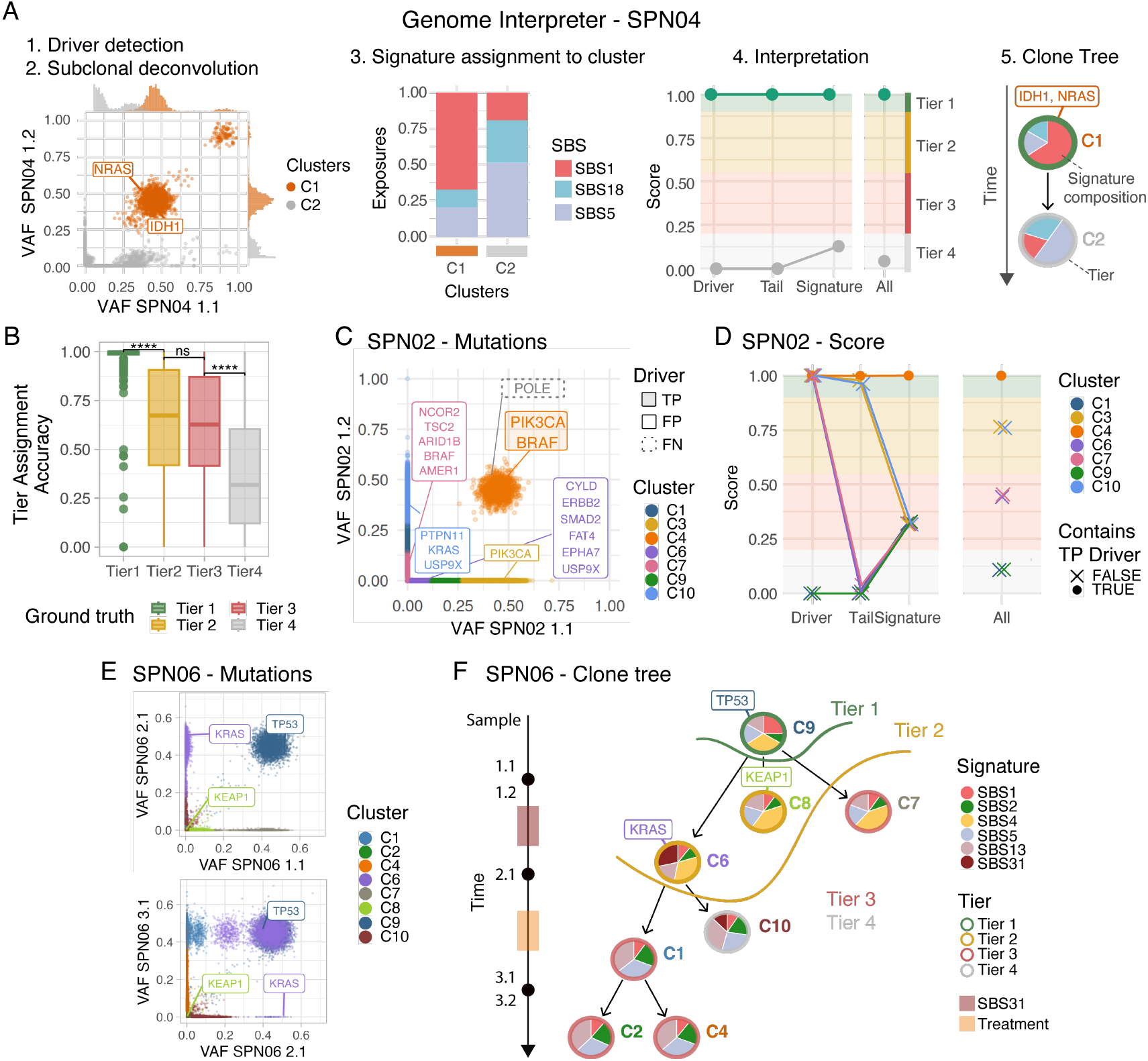
Interpretation of tumour evolutionary trajectories with nf-core/tumourevo’s genome interpreter. (A) The genome interpretation scoring implemented in nf-core/tumourevo integrates driver mutations, mutational signatures, subclonal deconvolution and clone tree reconstruction to rank mutation clusters (tiers 1 – 4) likely linked with subclonal expansions. The final output is a clone tree of evolutionary relations among clones, together with a pie chart reporting mutational signature activity per clone and the tier score estimated by the interpreter. (B) Accuracy of genome interpreter evidence-tier assignments evaluated against ProCESS simulation ground truth. (C,D) Subclonal mutations of the hypermutant colorectal cancer SPN02. The high TMB inflates spurious driver mutations across multiple low-frequency subclonal clusters, potentially confounding the evolutionary interpretation of this tumour (C). The Genome interpreter identifies which mutation clusters are supported by independent evolutionary evidence and flags as Tier 3 spurious clusters, reducing the confounding effects of hypermutation (D). (E) Multi-sample subclonal deconvolution with VIBER of SPN06, with annotated driver mutations. (F) Clone tree reconstruction of SPN06, obtained by nf-core/tumourevo using the genome interpreter. Chemotherapy mutagenesis (SBS31) is detected in clusters C6 and C10. The interpreter prioritises C6 (Tier 2 vs Tier 4) due to the presence of a KRAS driver mutation explaining disease resistance.

We next inferred single-base substitution (SBS) and insertion–deletion (ID) mutational signatures using SigProfiler [81] and BASCULE [82] (Methods). Across the cohort, inferred signature exposures closely matched the ground truth (median cosine similarity 0.988 for SBS and 0.981 for ID signatures). However, signature extraction was strongly influenced by clonal architecture, and subclonal mutational processes were generally harder to detect, especially when low-frequency signatures were masked by dominant clonal signals in SPN03, 04 and 07. This complexity impacted performance, with cosine similarity dropping from 0.991 to 0.922 for SBS signatures, and from 0.984 to 0.831 for ID signatures (Fig. 4B,C, Supplementary Fig. S29, Supplementary Table T9G). For example, this effect was particularly evident in SPN03, a longitudinal chronic lymphocytic leukaemia model under active surveillance [37–40]. In the earliest SPN03 sample, the mutational profile was dominated by clonal SBS1 mutations accumulated during the pre-neoplastic phase (Extended Data Fig. 7). In this context, detection of a low-frequency SBS5 signal (250 mutations with cancer cell fraction (CCF) <10%) required the use of an informative prior in BASCULE and was not recovered by SigProfiler. In contrast, in a later sample, after the SBS5-associated mutations had expanded to higher frequencies, both methods accurately recovered the underlying exposure profile. A similar effect was observed in the post-treatment sample of SPN07, a glioblastoma that acquired a hypermutant phenotype after developing resistance to temozolomide. Although both BASCULE and SigProfiler identified clonal SBS11 mutations, neither detected signatures of treatment (SBS25) and defective DNA mismatch repair (SBS26), which remained confined to a low-frequency hypermutant subclone (Extended Data Fig. 7). These results indicate that mutational signatures cannot be interpreted independently of tumour evolutionary structure, as temporal ordering and clonal expansion directly shape signature detectability.

We next performed subclonal deconvolution [83] using MOBSTER, VIBER [84] and PyClone-VI [18], with the aim to classify tumours as monoclonal or polyclonal under varying conditions of tumour purity and spatial sampling bias (Fig. 4D and Extended Data Fig. 8). Spatial sampling bias typically arises as cells collected from distinct tumour regions belong to different branches of the tumour phylogeny [23], leading to region-specific genomic profiles and potentially divergent interpretations of tumour clonality. Univariate analysis of individual tumour biopsies revealed that, in monoclonal tumours without spatial confounders, recall improved moderately with increasing purity (range 0.48-0.68; Fig. 4E, Supplementary Fig. S30, Supplementary Table T9H). In contrast, spatial sampling bias introduced a pronounced purity-dependent reduction in performance, with recall decreasing from 0.79 at purity 0.3 to 0.36 at purity 0.9. In particular, the overall amplification of the mutation frequency spectrum at higher tumour purity also enhanced spurious subclonal signals arising from spatial heterogeneity, rather than from genuine positive selection. Conversely, in polyclonal tumours, recall increased substantially with purity (0.31 to 0.89), consistent with improved detectability of positively selected subclones. This effect was particularly evident in SPN06 and SPN07, which contained expanding subclonal populations with larger CCF (> 0.4; Extended Data Fig. 8). These results show that tumour purity does not simply improve signal detection, but can amplify lineage-specific sampling biases embedded within the tumour phylogeny.

We next evaluated multivariate clustering, enabled by PyClone-VI [18] and VIBER [84], for reconstruction of clonal architectures across multiple samples (Supplementary Fig. S31, Supplementary Table T9I). To allow comparison across SPNs, we quantified the number of inferred clusters normalised by the number of samples (Fig. 4F). PyClone-VI systematically inferred more clusters than VIBER across tumour types and purity levels (median 3.12 versus 2.09, respectively). Consistent with the single-sample analyses, the effect of tumour purity on subclone quantification was largely restricted to tumours affected by spatial sampling bias (SPN02, 03 and 04). In these tumours, increasing purity systematically inflated the number of inferred clusters, indicating amplification of spurious subclonal structure. For example, in VIBER the median number of clusters increased from 1.75 at purity 0.3 to 3 at purity 0.9.

Together, these analyses establish nf-core/tumourevo as a unified framework for tumour evolutionary inference from bulk sequencing data, while revealing that multiple analytical tasks are jointly constrained by the same underlying evolutionary processes. Hypermutation, spatial sampling and treatment-associated clonal dynamics simultaneously distort mutational signatures, subclonal reconstruction and phylogenetic interpretation, generating coupled inference failures across analytical layers. These results indicate that tumour genome interpretation depends not only on sequencing quality, but also on the evolutionary regime from which the data emerge.

#### 2.3.1. Interpretation of tumour evolutionary dynamics with the genome interpreter

Tumour evolutionary analyses often generate fragmented and partially conflicting signals, making it difficult to distinguish genuine clonal expansions from passenger-dominated or artefactual processes. To address this, nf-core/tumourevo provides a genome interpreter framework that integrates multiple layers of evolutionary information to prioritise mutation clusters showing evidence of positive selection and biologically meaningful expansion. This multi-tier scoring system computes a ranking based on driver mutations, mutational signatures and subclonal structure (Fig. 5A), enabling joint interpretation of tumour evolutionary dynamics across otherwise disconnected analytical layers.

Across SCOUT, tier (1 to 4) assignments strongly stratified the probability that a mutation cluster represented a genuine clonal expansion. Concordance with the ProCESS ground truth progressively decreased from 0.99 (Tier-1) to 0.32 (Tier-4), indicating that higher-confidence tiers were enriched for robust evolutionary signals, whereas lower tiers captured increasingly ambiguous or passenger-dominated patterns (Fig. 5B, Supplementary Fig. S32, Supplementary Table T9J-K).

Importantly, the interpreter adapted flexibly to each SPN and successfully mitigated biases arising from individual analysis layers. For example, in the hypermutant colorectal cancer SPN02, which was overloaded by miscalled driver mutations (Fig. 5C), the interpreter classified all spurious mutation clusters (C6, C7, C9 and C10) as low-confidence Tier-3 (Fig. 5D). In SPN06, instead, the interpreter correctly distinguished biologically relevant expansion from passenger-dominated clusters, despite similar mutational signature distributions (Fig. 5E). Although two clusters (C10 and C6) were both associated with chemotherapy-related mutagenesis (SBS31 signature), one was assigned to a lower tier owing to enrichment for low-frequency passenger mutations and lack of evidence for sustained clonal expansion (Fig. 5F and Extended Data Fig. 9).

We next asked whether SCOUT could support analysis of treatment relapse dynamics, where treatment-induced mutagenesis, clonal selection, phylogenetic history and sampling jointly shape WGS patterns. SCOUT models these processes in SPN04 (chemotherapy resistance), SPN06 (platinum resistance) and SPN07 (temozolomide resistance). Although all three tumours converged on relapse, they did so through distinct evolutionary routes (Fig. 6A).

**Fig. 6:**
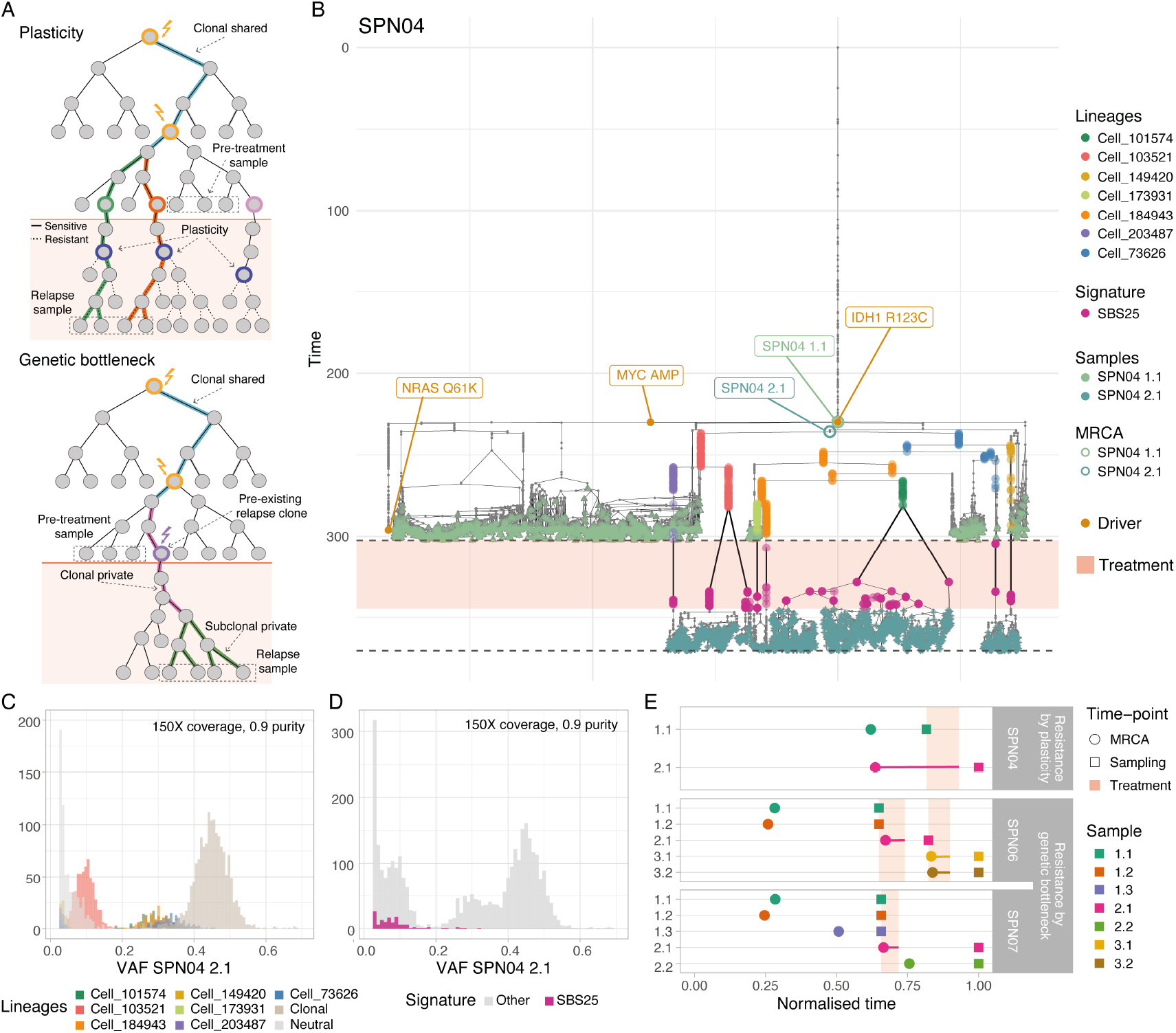
Simulated evolutionary dynamics of treatment resistance in SCOUT tumours. (A) Cell phylogenies comparing two resistance mechanisms: phenotypic plasticity (top) and selection of a pre-existing resistant clone (bottom). Under plasticity, resistance arises from recurrent, independent phenotype switches across multiple lineages; under clonal selection, treatment drives a strong bottleneck favouring a clone carrying a genetic or epigenetic driver that escapes negative selection. (B) Phylogeny of SPN04 with plasticity-associated resistance. Annotations indicate the MRCAs of the two samples, cells sampled at each time-point and treatment-associated SBS25 signature. Cell lineages resistant to treatment are also highlighted. (C) VAF distribution resolving mutations by class: clonal-cluster, neutral and resistance-clade-associated. The resistance-clade mutations form distinct VAF peaks whose frequencies reflect the size of the corresponding clade. (D) VAF distribution highlighting SBS25 treatment-associated mutations, showing their low frequency and abundance, which renders them largely undetectable in sequencing data. (E) Simulation timelines for SPN04, SPN06, and SPN07, displaying sampling timepoints, corresponding MRCAs, and the treatment window. SPN04 displays resistance through phenotypic plasticity, with the post-treatment MRCA located outside the treatment window, while SPN06 and SPN07 exhibit resistance driven by genetic bottlenecks, with post-treatment MRCAs coinciding with the treatment window.

In SPN04, resistance emerged through multi-lineage adaptation, consistent with tumour plasticity and the absence of relapse-associated driver alterations (Fig. 6B). Resistant populations remained phylogenetically close to pre-treatment cells. This multi-lineage adaptation produced multiple VAF peaks corresponding to independent resistant lineages (Fig. 6C), with SBS25 mutations diluted at low VAF because of the polyphyletic structure of the relapse population (Fig. 6D). By contrast, SPN06 and SPN07 followed a genetic bottleneck model, in which a pre-existing resistant clone carrying novel driver alterations expanded after treatment (Extended Data Fig. 10). The bottleneck produced a dominant clonal VAF peak, while treatment-associated mutations (SBS11/SBS31) spanned the clonal and subclonal VAFs (Extended Data Fig. 10). These differences arose mechanistically from the timing of the most recent common ancestor, which was earlier than the emergence of multi-lineage resistance (Fig. 6E), as predicted by evolutionary theory [23].

We used this simulation-based knowledge to establish an automated decision framework to interpret relapse dynamics in SCOUT, integrating evidence from the Genome Interpreter and clinical treatment annotations (Methods; Supplementary Table T9L). In both SPN06 and SPN07, we flagged the clone driving resistance supported by Tier 2 class evidence, exposure to treatment-associated mutational signatures and increased frequency following treatment. In both cases, the presence of relapse driver mutations enabled classification of this resistance as genetic, even though in one patient (SPN07) the causative driver event was misclassified. In SPN04, the resistant population was likewise detected as a population shared by both pre- and post-treatment samples. In this case, non-genetic resistance was correctly identified by combining the lack of relapse-associated driver events and diluted treatment-associated mutational signatures.

These results show that integrated interpretation of driver mutations, mutational signatures and subclonal structure resolves ambiguities that remain inaccessible to isolated analytical approaches, enabling coherent reconstruction of tumour evolutionary dynamics from bulk sequencing data. More broadly, SCOUT enables inference not only of clonal composition, but also of distinct modes of tumour adaptation and relapse. This type of mechanistic evolutionary inference is difficult to reproduce with spike-in simulations, which can emulate mutational mixtures but do not preserve lineage continuity, treatment-driven evolutionary trajectories and topology-dependent sampling effects.

## 3. Discussion

Advances in whole-genome sequencing (WGS) have shifted the central challenge of cancer genomics from data generation to data interpretation [6], with tumour evolution emerging as a unifying framework for understanding cancer genomes [23]. While the foundational bioinformatics methods developed over the past decades are now largely consolidated, modern cancer genome analysis increasingly relies on statistical and machine learning approaches that infer tumour composition, mutational processes and evolutionary history from noisy and incomplete observations. This evolutionary perspective is essential because somatic mutations, chromosomal instability, mutagenesis and treatment resistance are not independent phenomena, but manifestations of evolutionary dynamics unfolding across space and time. As these inference frameworks become increasingly central to biological discovery and clinical decision-making, systematic and reproducible benchmarking becomes essential. Our results argue that the missing ground truth of cancer genomics benchmarking is fundamentally evolutionary.

In this work, we developed ProCESS and nf-core/tumourevo to generate SCOUT, a synthetic tumour WGS benchmark with complete evolutionary and genomic ground truth. Across multiple tumour types and clinically relevant scenarios, SCOUT jointly models tumour growth, mutagenesis, chromosomal instability, treatment dynamics and sequencing generation from FASTQ level onward. Unlike conventional spike-in approaches that independently perturb isolated analytical layers, SCOUT preserves the evolutionary dependencies linking clonal architecture, mutational processes, copy-number dynamics and spatial sampling structure. Although no finite cohort can capture the full diversity of human cancers, the simulated tumours span a broad range of evolutionary behaviours, including hypermutation, whole-genome doubling, subclonal selection, spatial heterogeneity and therapy-associated resistance. Together, these scenarios provide a robust framework for benchmarking methods that profile tumour genomes and reconstruct their evolutionary trajectories.

Using SCOUT, we systematically evaluated state-of-the-art workflows for mutation and copy-number calling, mutational signature inference and tumour evolutionary reconstruction. Across methods, performance deteriorated reproducibly in low-purity tumours, highly subclonal samples, hypermutated genomes and structurally complex cancers. Notably, tumour purity consistently exerted a stronger effect on inference accuracy than sequencing depth, indicating that biological composition rather than sequencing volume defines a major operational limit of current WGS analyses. Importantly, these sources of variability are not independent technical confounders, but emergent consequences of tumour evolutionary dynamics that propagate across multiple inference layers.

Our analyses further revealed that spatial tumour structure remains a major and underappreciated confounder for evolutionary inference. Increasing tumour purity amplified not only genuine subclonal signals, but also artefactual ITH arising from spatial sampling bias, leading to systematic overestimation of subclonal complexity. Similarly, low-frequency mutational processes frequently escaped accurate reconstruction despite deep sequencing and advanced inference methods. These observations suggest that many analytical limitations arise not only from technical noise but also from the intrinsic evolutionary complexity of tumours.

More broadly, this work argues that realistic benchmarking in cancer genomics cannot rely on isolated variant spike-ins or simplified simulations. Because cancer genomes are shaped by dynamic processes of mutation, selection, genome instability, treatment response and spatial growth, benchmarking frameworks must model these processes jointly to reproduce the coupled signals observed by sequencing technologies. In this respect, SCOUT establishes a new class of evolutionary-aware benchmark resource for cancer genomics.

Importantly, integrated evolutionary interpretation enabled reconstruction not only of clonal composition, but also of distinct modes of tumour adaptation to treatment. By jointly analysing mutational signatures, phylogenetic topology, temporal ordering and clonal continuity, we distinguished treatment resistance driven by adaptive multi-lineage evolution from resistance emerging through selection of pre-existing resistant clones. These mechanistically distinct trajectories generated different patterns across multiple analytical layers, including VAF distributions, mutational signatures and ancestor timings. Such evolutionary structures are difficult to reproduce using spike-in simulations, which can emulate mutational mixtures but do not preserve lineage continuity, treatment-driven evolutionary trajectories and topology-dependent sampling effects. More generally, these results suggest that tumour adaptation may become inferable from integrated evolutionary signals embedded within bulk sequencing data.

Finally, we view this study as a starting point rather than a complete solution. We intentionally focused on bulk short-read WGS because it remains the most widely adopted platform for large-scale cancer genomics and a promising direction for clinical sequencing. Similarly, we benchmarked a selected set of widely used analytical methods, recognising that many additional tools and workflows could not be systematically evaluated within the scope of this study. Rather than providing an exhaustive comparison, we aimed to establish a reproducible evolutionary benchmark framework that can enable broader community-driven evaluation efforts. The underlying framework is inherently extensible and can naturally support additional tumour types, evolutionary scenarios and orthogonal assays, including long-read sequencing, single-cell genomics, transcriptomics and epigenomic profiling.

Extending these frameworks towards phenotypic tumour evolution will require solving major mathematical and computational challenges that connect genetic evolution with partially heritable epigenetic and cellular states, ultimately enabling integrated modelling of genotype–phenotype dynamics during tumour progression and therapeutic adaptation. We anticipate that SCOUT will provide a reproducible foundation for the development, benchmarking and clinical validation of future cancer genomics and machine learning methods, while supporting a broader transition towards benchmarking strategies in which tumour evolution itself becomes the central organising principle for interpreting cancer genomes.

### Data, software, and reproducibility ecosystem

To maximise transparency, reuse, and long-term reproducibility, we release the complete SCOUT/ProCESS/nf-core/tumourevo ecosystem, including raw sequencing data, processed benchmark outputs, fully reproducible computational environments, workflow pipelines, and open-source software packages.

A central project hub aggregating all resources, documentation, and release information is available at:

https://caravagnalab.github.io/SCOUT-website/

### Raw sequencing and benchmark data

The complete collection of FASTQ files generated for the SCOUT benchmark cohort, including all simulated sequencing experiments across 7 SPNs, multiple tumour purities, and sequencing depths, has been deposited at the European Nucleotide Archive (ENA) under accession:

PRJEB97253T

The data will be released the 30th June 2026.

We release the full set of processed outputs generated across all executions of nf-core/sarek and nf-core/tumourevo, enabling direct inspection, downstream benchmarking, and independent validation of all analyses presented in this study:

data are stored in the Zenodo DOI listed in Supplementary Table T10

### Open-source software

This work is accompanied by the release of three complementary open-source resources:

- SCOUT – an R package providing ground-truth annotations, reference tables and parser for raw and processed data for all SPNs in the SCOUT cohort [85]: https://caravagnalab.github.io/SCOUT All results presented in this work were generated using version 1.1.0.
- ProCESS – an open-source R package for tumour evolutionary reconstruction, benchmarking, and simulation-aware analysis [86]: https://caravagnalab.github.io/ProCESS All analyses presented in this manuscript were generated using version 1.1.0 (Release 1.1.0 “Bastille”).
- nf-core/tumourevo – a fully reproducible nf-core workflow for scalable and portable execution across hetero-geneous computational infrastructures: https://nf-co.re/tumourevo All workflow executions reported in this study were performed using version dev0.0.1 that can be found here: https://github.com/caravagnalab/tumourevo/releases/tag/dev0.0.1 In addition, all scripts, workflow configurations, and figure-generation code required to reproduce every analysis and benchmark presented in this manuscript are publicly available at: https://github.com/caravagnalab/process_reproducibility

### Portable reproducibility

To ensure exact computational reproducibility across infrastructures, we release fully containerised execution environments.

- Docker image for ProCESS v1.1.0 is publicly available at: https://hub.docker.com/repository/docker/lvaleriani/process/tags/1.1.0/

## Authors contribution

ACas developed ProCESS, with support from LV, GG, RB, GiS, AA, EB, NC, AS under the supervision of GC. LV, GG, EB, KD, VAG developed nf-core/tumourevo, under the supervision of NC and GC. LV, GG, GiS, AA, AS, VAG, SM developed SCOUT, under the supervision of RB and GC. LV and GG run data generation and analysis with both nf-core/sarek and nf-core/tumourevo, supported by EB, GiS, KD, AA, AS, VAG, SM and ER. All authors examined data and interpreted results. LV, GG, NC, RB, ACas and GC drafted the manuscript, which all authors approved. GC conceived, designed, and coordinated the study.

## Acknowledgments

The authors acknowledge the Area Science Park supercomputing platform ORFEO made available for conducting the research reported in this paper. We thank Konstantin Winter and Anastasiia Romanova for the invaluable support to the Computational Biology Research Centre at Human Technopole.

## Ethical approvals

Not required.

## 4. Methods

### The ProCESS simulation framework

ProCESS (*Pro*grammable *C*ancer *E*volution *S*patial *S*imulator)[86] is an R interface to CLONES (*C*++ *L*ibrary f*O*r *N*eoplastic *E*volution *S*imulations), a C++ library for simulating tumour evolution. These tools were developed to generate the SCOUT cohort and enable controlled simulation of tumour growth, mutational processes and sequencing data. A detailed description of the algorithms and implementation is provided in Supplementary Note A.

ProCESS organises simulations across three hierarchical levels:

#### Tissue level

Tumour growth is modelled on a two-dimensional lattice, initialised from a single founding cell. Each clone is defined by user-specified growth and death rates, governing a stochastic birth–death process that drives spatial expansion[87]. New clones arise through evolutionary transitions from pre-existing states. Cells can be sampled from the lattice under multi-region or longitudinal designs, generating bulk samples with controlled clonal composition. A ground-truth phylogeny describing the evolutionary relationships between sampled cells is recorded.

#### Mutation level

Somatic mutational processes are specified through clone-specific or global parameters that define mutation rates and spectra. Each clone is associated with a molecular profile comprising driver events and passenger mutations across SNVs, indels and copy-number alterations. Passenger mutation burden is determined by user-defined mutation rates. Germline variation is introduced by sampling haplotypes from the 1000 Genome Project [34]. Time-varying mutational processes can be modelled using COSMIC (v3) SBS and indel signatures with user-defined exposure trajectories. Mutations are stochastically assigned along the simulated phylogeny.

#### Sequencing level

Matched tumour–normal bulk WGS data are generated under user-defined sequencing conditions. Illumina sequencing is modelled by specifying read length, error rates and library configuration (paired-end or single-end). Allele-specific read counts for somatic variants are simulated using a Beta–Binomial model to capture overdispersion. Sequencing data are generated at user-specified tumour purity and coverage, producing synthetic datasets suitable for downstream analysis.

ProCESS simulations provide fully specified ground truth for tumour evolution, enabling systematic benchmarking of bioinformatics and machine-learning methods, including somatic variant detection, mutational signature analysis, subclonal reconstruction, copy-number inference and phylogenetic modelling.

### The SCOUT cohort

The SCOUT cohort comprises seven tumour profiles (SPN01–07) selected to represent well-characterised disease trajectories across solid and haematological malignancies, as described in the literature. This section summarizes the key evolutionary features of each sample in the cohort, highlighting patient-specific genomic configurations and the main analytical challenges encountered in both molecular profiling and subclonal evolutionary reconstruction. In Supplementary Note B we provide a detailed description of each SPN.

SPN01 and SPN02 simulate colorectal cancer [35, 36] and follow canonical trajectories determined by early tumour-suppressor inactivation. SPN01 is characterized by two canonical driver SNVs in *APC* and *TP53*, followed by a copy number deletion of the wild-type allele of *APC*. SPN01 is characterized by the expansion of a subclone harbouring whole-genome duplication (WGD), alongside one sample exhibiting a fully clonal WGD event. The primary challenge in this case arises from limitations in copy number calling algorithms, which often fail to robustly detect WGD and, more critically, to distinguish between clonal and subclonal WGD events.

SPN02 is a microsatellite instable colorectal cancer and is characterized by *KRAS, BRAF, PIK3CA* and *POLE* driver mutations, which are normally found in hypermutant cases [36, 88]. The main challenge is accurate driver mutation identification. Despite restricting the analysis to known colorectal cancer drivers, numerous false positives are detected, particularly at low variant allele frequencies. These spurious calls can be mitigated through cluster-based interpretation, as they are largely assigned to tail mutation clusters by univariate subclonal deconvolution methods. SPN03 represents chronic lymphocytic leukaemia under active surveillance [37–40], characterised by slow clonal diversification driven by alterations in genes such as *NOTCH1, MAP2K* and *TP53*, together with deletions at 13q14. The tumour is sampled longitudinally to capture continuous evolutionary dynamics in the absence of treatment and it is characterized by a low mutation rate. The main challenge is mutational signature deconvolution, which is hindered by the limited number of mutations available, particularly at low variant allele frequencies. Additionally, the final clonal expansion is driven by a mutation absent from common driver annotation databases, resulting in its non-detection by nf-core/tumourevo driver annotation step.

SPN04 models acute myeloid leukaemia [41–43] undergoing relapse driven by non-genetic resistance. The tumour includes canonical AML drivers such as *NRAS* and *IDH1*, together with a *MYC* amplification. The tumour is treated simulating chemotherapy, and the evolutionary trajectory is characterised by limited genetic divergence between diagnosis and relapse, resulting in genetically similar MRCAs. The tumour is sampled longitudinally across treatment to capture this behaviour. The main challenge is the detection of the treatment-induced mutational signature, characterized by low frequencies mutations.

SPN05 corresponds to breast cancer [44–46] and follows a trajectory combining early driver mutations, including *PIK3CA* and *TP53*, with later genome instability associated with *BRCA2* alterations, copy-number changes and whole-genome doubling. The tumour is sampled using a multi-region design to capture spatial heterogeneity, and combines a particularly complex pattern of homologous recombination deficiency typical of BRCA-loss breast cancers. The hypermutant phenotype introduces a substantial number of false positives in driver annotation, often resulting in the identification of spurious mutation clusters. Concurrently, copy number–altered drivers pose a challenge for multivariate subclonal inference methods that do not explicitly model copy number (e.g., VIBER), as these approaches may fail to associate such drivers with the correct clusters, leading to apparent driverless subclones despite the presence of biologically relevant events.

SPN06 represents lung adenocarcinoma [47] with treatment-driven evolution, characterised by alterations in *KRAS, EGFR, KEAP1* and *STK11*, and by the emergence of resistant clones following two rounds of chemotherapy. The tumour integrates both longitudinal and multi-region sampling to capture clonal replacement and spatial heterogeneity at relapse. In this case, resistance is linked with the emergence of de novo *KRAS* mutations and their subsequent amplifications. SPN06 exhibits genetic resistance to therapy, resulting in a highly subclonal tumour structure. The MRCAs of the pre- and post-treatment samples are genetically distant, leading to a greater number of clonal mutations in the later sample and a more detectable treatment-induced mutational signature. However, the highly subclonal architecture introduces substantial complexity in subclonal reconstruction, as mutations at elevated frequencies may be mistakenly attributed to clonal expansions, complicating the distinction between true subclonal dynamics and artefactual signals.

SPN07 corresponds to glioblastoma [48–51] with therapy-associated hypermutation and rapid evolutionary dynamics, including alterations in *TP53, ATRX* and mismatch-repair genes such as *MSH6*. The tumour combines longitudinal and multi-region sampling to model treatment-induced mutational processes, linked to exposure to temozolomide. A key challenge arises from the presence of driver mutations affected by copy number alterations, such as PTEN loss of heterozygosity, which complicates their detection by multivariate methods that do not account for copy number. As a result, correctly identifying the clonal drivers and reconstructing the evolutionary trajectory of the tumour becomes more difficult, particularly in the context of treatment-induced genomic changes. An additional challenge is the identification of subclonal processes linked to therapy, as these are often represented by mutational signatures with low exposure and low variant allele frequency, further limiting their detectability.

### Computational resources (time, CPU, memory, RAM)

The computational workflow comprised four main stages. SPNs simulation and in silico sequencing were performed using ProCESS within a Singularity container on an AMD EPYC partition (dual EPYC 7H12 processors), optimized for highly parallel CPU workloads. Downstream analyses, including alignment, variant calling and tumour evolution inference, were carried out on AMD EPYC 9374F (Zen 4) processors (64 cores, 512 GB RAM per node) using nextflow pipelines, nf-core/sarek and nf-core/tumourevo. A custom nextflow pipeline was develop to allow systematic benchmarking of molecular profiling and subclonal and signature deconvolution methods. All workflows were parallelized across available compute cores and managed via SLURM job scheduler. Typical runtimes were ~220 min per sample for alignment and variant calling and ~50 min for downstream copy-number analyses, corresponding to a total computational cost of ~1,500 CPU hours per sample, with peak memory usage of up to ~166 GB per task.

### Standard metrics of benchmarking

For benchmarking molecular profiling and tumour evolutionary inference tools, we use standard machine learning metrics that compare the ground truth signal with the inferred one. These metrics are defined based on the number of true positives (TP), false negatives (FN), false positives (FP), and true negatives (TN), which may vary depending on the specific context (e.g. somatic mutation calling or driver detection algorithms). From these quantities, Precision, Recall, F1 score, and Accuracy are calculated as follows:

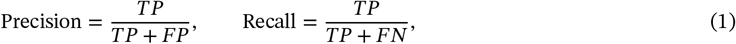

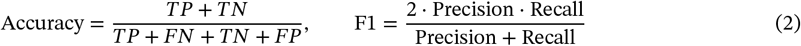

For tasks involving the inference of continuous quantities, such as VAFs (somatic and germline) and mutational signature exposures, we quantified the discrepancy between inferred and simulated values using the Mean Squared Error (MSE) and Root Mean Squared Error (RMSE) computed as follows:

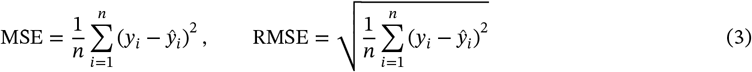

### Benchmarking with nf-core/sarek

We applied the nf-core/sarek pipeline (v3.5.1) to perform standardized alignment and variant calling on matched tumor-normal whole-genome sequencing data in all coverage-purity combinations. The pipeline includes read alignment with BWA-MEM2 (v2.2.1) [89] and preprocessing steps such as duplicate marking and base quality score recalibration using GATK tools [90]. Somatic variant calling was performed with Strelka2 [72], FreeBayes [73], and Mutect2 [71], the latter run in joint mode to identify both shared and private mutations across all samples from a single SPN. For copy number analysis, we used ASCAT [75] to infer allele-specific CNAs and estimate tumour purity and ploidy. To expand the set of allele-specific CNA callers, we developed a custom nextflow pipeline to integrate Sequenza [76] and Battenberg [20]. Additionally, germline variant calling was conducted on the normal samples using HaplotypeCaller [12], FreeBayes [73], and Strelka2 [72]. In Supplementary Note C we provide a detailed analysis of the results for full cohort and each SPN.

#### Performance metrics of variant callers

For each somatic variant caller (Mutect2, Strelka2, FreeBayes), we evaluated its ability to detect SNV and INDEL mutations and to estimate their variant allele frequencies (VAFs) against the known ground truth across varying combinations of tumour purity and coverage. A variant was considered truly somatic in the simulated BAMs when its tumour VAF was ≥ 0.02. For Strelka2 and Mutect2, somatic calls corresponded to variants marked PASS in their somatic VCFs. For FreeBayes, which lacks explicit somatic calling, we classified variants as somatic if they met the following empirical thresholds: quality ≥ 20, tumour VAF ≥ 0.01, normal VAF ≤ 0.02, and depth ≥ 10. Based on these definitions, we derived precision, recall, and accuracy according to Equations 1,2. To assess the impact of tumour subclonality, we computed these metrics across cancer cell fraction (CCF) bins spanning 0–5%, 5–10%, 10–25%, 25–50%, 50–75%,75–95%, and clonal (CCF ≥ 95%), enabling detailed comparison of caller performance across the full spectrum of allelic representation.

We applied a similar benchmarking strategy for germline variant callers, comparing their calls against the known germline mutations included as ground truth in the simulated data. For germline analysis, CCF binning was not used; instead, we directly calculated metrics in Equations 1,2, together with B-allele frequency (BAF) Pearson correlation and root mean square error (RMSE) (Equation 3) with respect to the simulated BAF on high quality variants.

#### Performance metrics of CNA callers

To benchmark copy number alteration (CNA) callers, we stratified samples according to the fraction of genome altered (FGA), defined as the proportion of the autosomal genome deviating from a diploid copy number state:

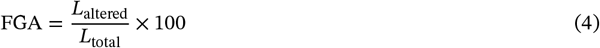

where *L*_altered_ is the total length of segments with copy number changes (CN ≠ 1:1 or ratio < 1), and *L*_total_ is the total length of autosomal segments. Samples with FGA > 15% were classified as High FGA; all remaining samples were designated Low FGA. CNA caller performance was then evaluated separately within each category.

We first compared purity and ploidy estimates produced by ASCAT, Sequenza, and Battenberg against the ground truth generated by ProCESS. For each sample, we computed the absolute error between predicted and true values, (*δ*_*π*_ for purity and *δ*_*σ*_ for ploidy) allowing systematic biases to be quantified across FGA classes. We classify then samples with *δ*_*π*_ > 0 as Miscalled purity and *δ*_*σ*_ > 0.7 as Miscalled ploidy.

We next assessed breakpoint detection accuracy and copy number state inference. Because ProCESS simulations contain both clonal and subclonal CN segments, the latter defined by a cancer cell fraction CCF < 1, segments were categorised accordingly prior to evaluation.

Breakpoint detection was evaluated independently for each caller using a tolerance-based matching procedure. True breakpoints were derived from the simulated segments and predicted breakpoints from each tool; the two sets were then compared within a predefined window (*δ* = 10 kb). For each predicted breakpoint, the nearest unmatched true breakpoint falling within *δ* was identified and recorded as a true positive (TP). Predicted breakpoints with no matching true breakpoint were counted as false positives (FP), and unmatched true breakpoints as false negatives (FN). The resulting confusion matrix quantifies breakpoint detection accuracy for each caller.

For each segments *s*, classified as clonal, we additionally assessed whether the predicted copy number state matched the simulated state. The proportion of the genome correctly inferred for clonal segments (*ρ*_clonal_) was calculated as:

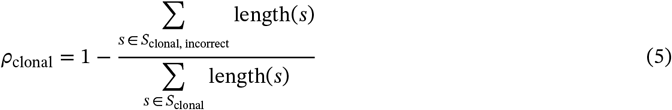

An analogous evaluation was performed for subclonal segments. Samples were further stratified by the fraction of genome subclonal (FGS), defined as the proportion of the autosomal genome harbouring subclonal copy number states:

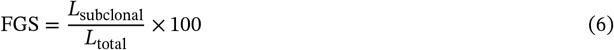

where *L*_subclonal_ is the total length of subclonal segments and *L*_>total_ is the total genome length. The proportion of correctly identified subclonal copy number states (*ρ*_subclonal_) was then calculated using the same approach as for clonal segments.

### Benchmarking with nf-core/tumourevo

We developed nf-core/tumourevo, the first community-based pipeline that uses tumour molecular profiles to perform automatic and reproducible tumour evolution analysis from bulk DNA sequencing data, and applied it to SCOUT. The pipeline naturally integrates with nf-core/sarek and provides access to state-of-the-art machine-learning tools for the interpretation of tumour genomes, including quality control of variant calls and detection of subclonal expansions that integrate driver-mutation data and mutational signatures. The pipeline requires somatic variant calls and allele-specific copy-number profiles, along with inferred ploidy and purity estimates. The nf-core/tumourevo workflow includes the following main steps: (i) variant annotation with Ensembl VEP [79] and driver annotation using the COSMIC database [91]; (ii) quality control of inferred VAF and CNA profiles using CNAqc [92] and TINC [78]; (iii) subclonal deconvolution of high-quality mutations with MOBSTER, VIBER [84], and PyClone-VI [18]; (iv) mutational-signature deconvolution with SigProfiler [81] and SparseSignatures [93, 94]; (v) signature deconvolution analysis in multivariate subclonal inferred cluster and (vi) downstream genome interpretation modules. To broaden the set of signature-deconvolution methods, we additionally developed a custom pipeline integrating BASCULE [82]. SparseSignatures, even if present in nf-core/tumourevo pipeline, was not applied, as robust inference with this method typically requires at least 100 samples to ensure accurate signature extraction, which exceeds the size of the dataset analysed here. To assess how caller accuracy affects downstream evolutionary analyses (e.g., subclonal and mutational-signature deconvolution), we executed the pipeline across multiple combinations of somatic variant and CNA callers. In total, we evaluated four combinations of somatic callers (Mutect2, Strelka) and CNA callers (Sequenza, ASCAT), together with nine purity–coverage settings. In Supplementary Note D we provide a detailed analysis of the results for the full cohort and each SPN.

#### Performance metrics for driver detection

Driver identification performance was evaluated using standard binary classification metrics according to Equations 1,2. All metrics were computed in the multisample setting, yielding a single performance value per SPN rather than per individual sample. To assess whether performance varies across mutation rate regimes, SPNs were stratified by their simulated mutation rate into hypermutant (*μ* = 1 × 10^−7^) and other (*μ* < 1 × 10^−7^) categories.

#### Performance metrics for subclonal deconvolution

Univariate subclonal deconvolution methods were evaluated on their ability to detect subclonal structure at the single-sample level. Each sample was annotated according to the presence or absence of at least one subclone in the ProCESS ground truth, and accordingly classified as *monoclonal* (no subclonal expansion present) or *polyclonal* (at least one subclonal expansion present). The evaluation was deliberately restricted to this binary detection task, without assessing the accuracy of the inferred number of subclonal clusters. For each category, we defined TP and FN separately: for monoclonal samples, TP are samples correctly classified as monoclonal, and FN are monoclonal samples incorrectly classified as polyclonal; conversely, for polyclonal samples, TP are samples correctly classified as polyclonal, and FN are polyclonal samples incorrectly classified as monoclonal. We then compared the recall (Equation 1) between the two categories as a function of increasing tumour purity.

We further evaluate the multivariate subclonal deconvolution methods VIBER and PyClone-VI by comparing the number of inferred clusters relative to the number of samples per SPN, reported as the ratio of number of detected clusters over the number of samples. This ratio is compared across SPNs with and without sampling bias, in order to assess the effect of spatial sampling on the inference of clonal architecture.

#### Performance Metrics for Signature Reconstruction methods

For each signature deconvolution tool, we benchmarked its ability to accurately identify the simulated mutational signatures, both in the SBS96 and ID83 contexts, and their corresponding exposures. For each sample, we first computed TP, FP, and FN by comparing the known ground-truth signatures with the tool-inferred signatures, and then used to compute the metrics described above (Equations 1 and 2).

To evaluate performance in estimating signature exposures, we calculated the cosine similarity and mean squared error (MSE) between the ground-truth exposure vector *x* and the inferred exposure vector *y*. Cosine similarity was computed as:

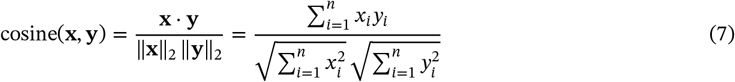

The mean squared error was calculated for each inferred exposure and summarized as the average MSE per sample (Equation 3). To evaluate the potential confounding effect of subclonal mutational signatures, samples were stratified into high- and low-complexity groups based on differences in signature composition between clonal and subclonal mutations. Specifically, we compared the sets of mutational signatures assigned to clonal and subclonal clusters using the Jaccard similarity index computed as:

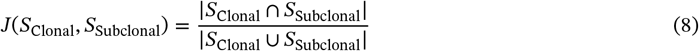

Values of *J* approaching 1 indicate highly similar signature compositions between clonal and subclonal mutations and were therefore classified as low-complexity samples. In contrast, samples with *J* ≤ 0.9 were considered high complexity, reflecting the emergence of distinct mutational processes associated with subclonal expansion. The Jaccard index was averaged across all coverage–purity combinations and computed separately for single base substitutions (SBS) and insertions/deletions (ID) using the simulated ground truth data.

#### Performance Metrics for Genome Interpretation

To evaluate the performance of the Genome Interpreter scoring system, each mutation was annotated with two independent subclonal expansion evidence labels: (i) the evidence Tier assigned to the mutation cluster inferred by the nf-core/tumourevo pipeline, and (ii) the corresponding Tier derived from the ProCESS ground-truth simulation. Classification performance was then assessed separately for each Tier by treating Tier membership as a binary prediction problem at the level of individual mutations. Specifically, for a given Tier, mutations assigned to that Tier in the ground truth were considered positive instances, whereas all remaining mutations were considered negative instances. Standard binary classification metrics were subsequently computed according to Equations 1 and 2, enabling quantitative evaluation of the concordance between inferred and simulated expansion evidence across individual mutations.

#### Framework for automated classification of resistance mechanisms

To identify mechanisms of treatment resistance, we developed an algorithm that integrates clinical treatment annotations with evidence of subclonal expansion under therapeutic selection. We assumed that simulated sequencing samples were annotated as pre- or post-treatment, whereas evidence of subclonal expansion was obtained from the Genome Interpreter through subclonal deconvolution, driver alteration assignment, and mutational signature analysis performed at the level of individual genetic clones. This information was used to classify tumour clones according to their putative role in treatment resistance. The procedure consists of three steps:

##### Step 1

Clones assigned to Tier 3 or lower were excluded owing to insufficient evidence of subclonal expansion.

Only Tier 1 and Tier 2 clones were retained as candidate therapy-resistant populations.

##### Step 2

For each clone passing Step 1, we computed a continuous resistance score ℛ defined as:

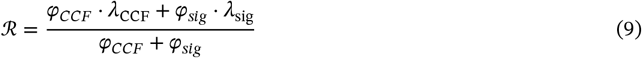

where *λ*_CCF_ ∈ {0, 1}and *λ*_sig_ ∈ {0, 1}are indicator variables equal to 1 if (i) the mean cancer cell fraction (CCF) across post-treatment samples exceeded the mean CCF across pre-treatment samples, and (ii) at least one mutational signature active in the clone was treatment-associated (for example, belonging to the Chemotherapy, Immunosuppressants, or Treatment categories of the COSMIC v3 catalogue [31]), respectively. For the CCF increase component, we additionally required the mean pre-treatment CCF to exceed 10^−3^ in order to exclude clones detected at negligible abundance before treatment, which may arise from sequencing noise. The parameters *φ*_*CCF*_ and *φs*_*ig*_ determine the relative contribution of the two components. By default, the CCF increase term was assigned twice the weight of the mutational signature term (*φ*_*CCF*_ = 2/3 and *φ*_*sig*_ = 1/3), reflecting the fact that expansion despite therapeutic pressure constitutes direct evidence of positive selection and resistance. In contrast, treatment-associated mutational signatures do not provide evidence of resistance per se, but rather indicate that the clone was exposed to and continued to accumulate treatment-induced mutations. Clones with ℛ > 0.6 were classified as candidate treatment-resistant populations.

##### Step 3

For clones classified as treatment-resistant, we assessed whether resistance was likely driven by genetic or non-genetic mechanisms. Clones harbouring at least one driver alteration not shared with any other clone were classified as *genetic resistance candidates*. Candidate resistant clones lacking identifiable private driver alterations, as well as resistant clonal populations that by definition do not possess newly acquired clone-specific genetic events, were instead classified as *non-genetic resistance candidates*.

## 5. Figures

**Extended Data Fig. 1:**
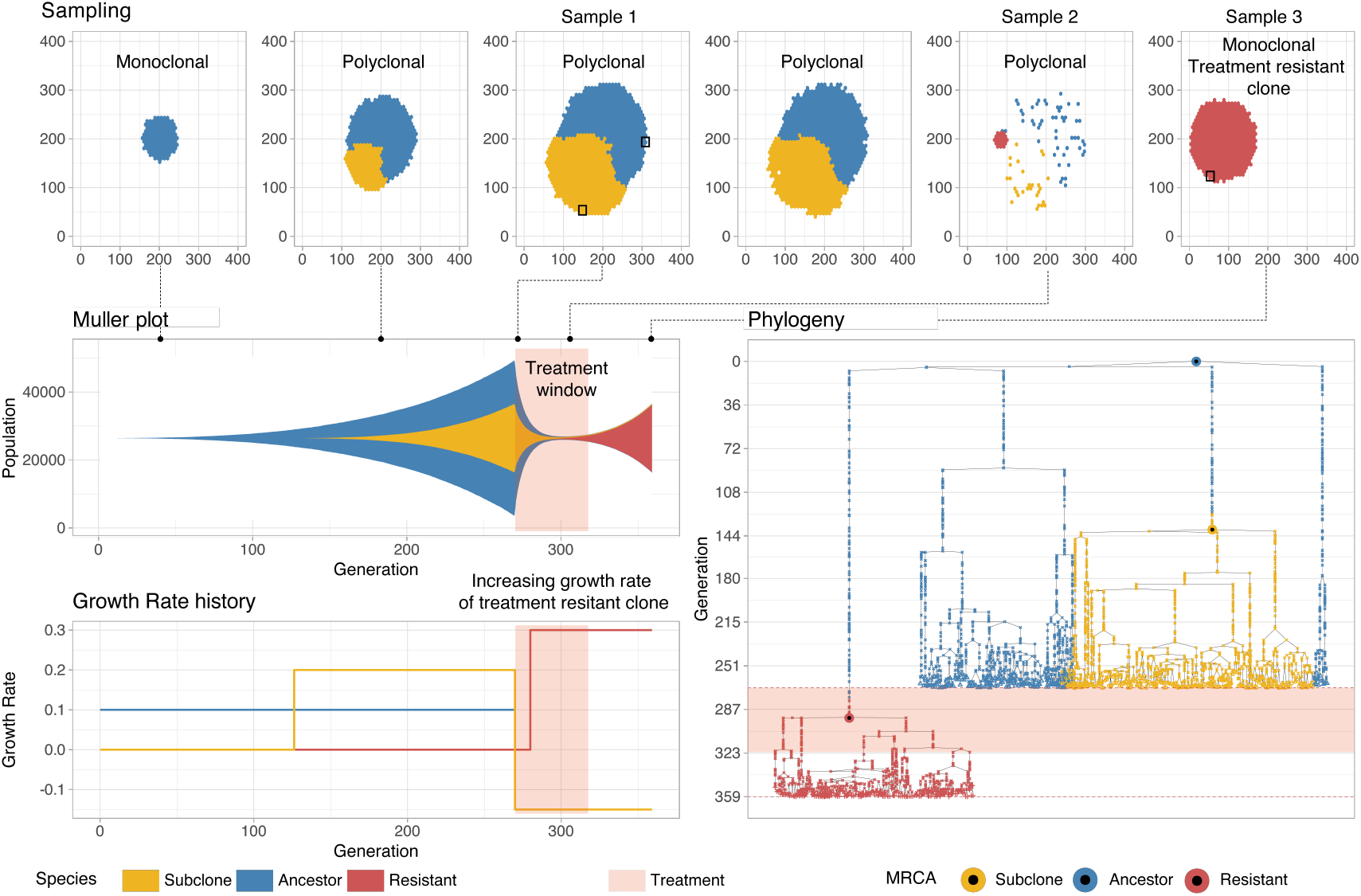
Main features of the ProCESS simulation framework. ProCESS simulates tumour growth as a spatial stochastic branching process on a two-dimensional lattice, modelling the expansion of multiple cell populations with distinct evolutionary parameters, including growth and death rates. Mutational processes are introduced over time and can be modulated to reflect endogenous and treatment-associated dynamics, enabling the simulation of clonal sweeps and therapeutic responses. Tumours can be sampled through multi-region or longitudinal designs, and the full genealogy of sampled cells is recorded at single-cell resolution. These features enable the generation of synthetic tumours with controlled evolutionary histories and known ground truth for benchmarking downstream analyses.

**Extended Data Fig. 2:**
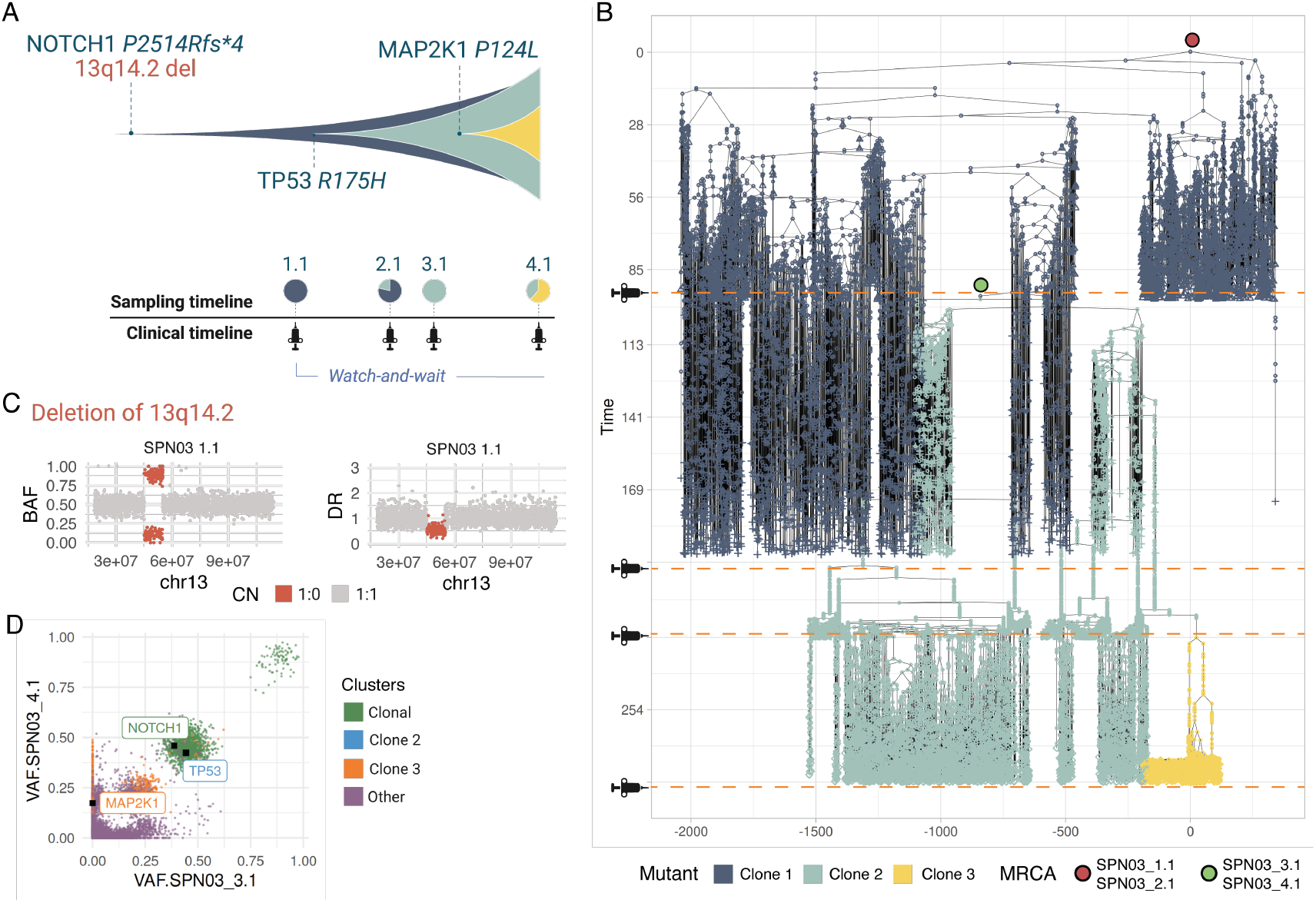
SPN03 main evolutionary features. (A) Muller plot depicting the clonal dynamics of the SPN03 simulation prototype. (B) Simulated phylogenetic forest with the most recent common ancestor (MRCA) of each sample and corresponding sampling times annotated. (C) B-allele frequency (BAF) and depth ratio (DR) of the 13q14.2 driver deletion in sample 1.1. (D) Multivariate variant allele frequency (VAF) distribution across the last two samples (3.1 and 4.1), illustrating simulated driver SNV and indel mutations and their respective frequencies.

**Extended Data Fig. 3:**
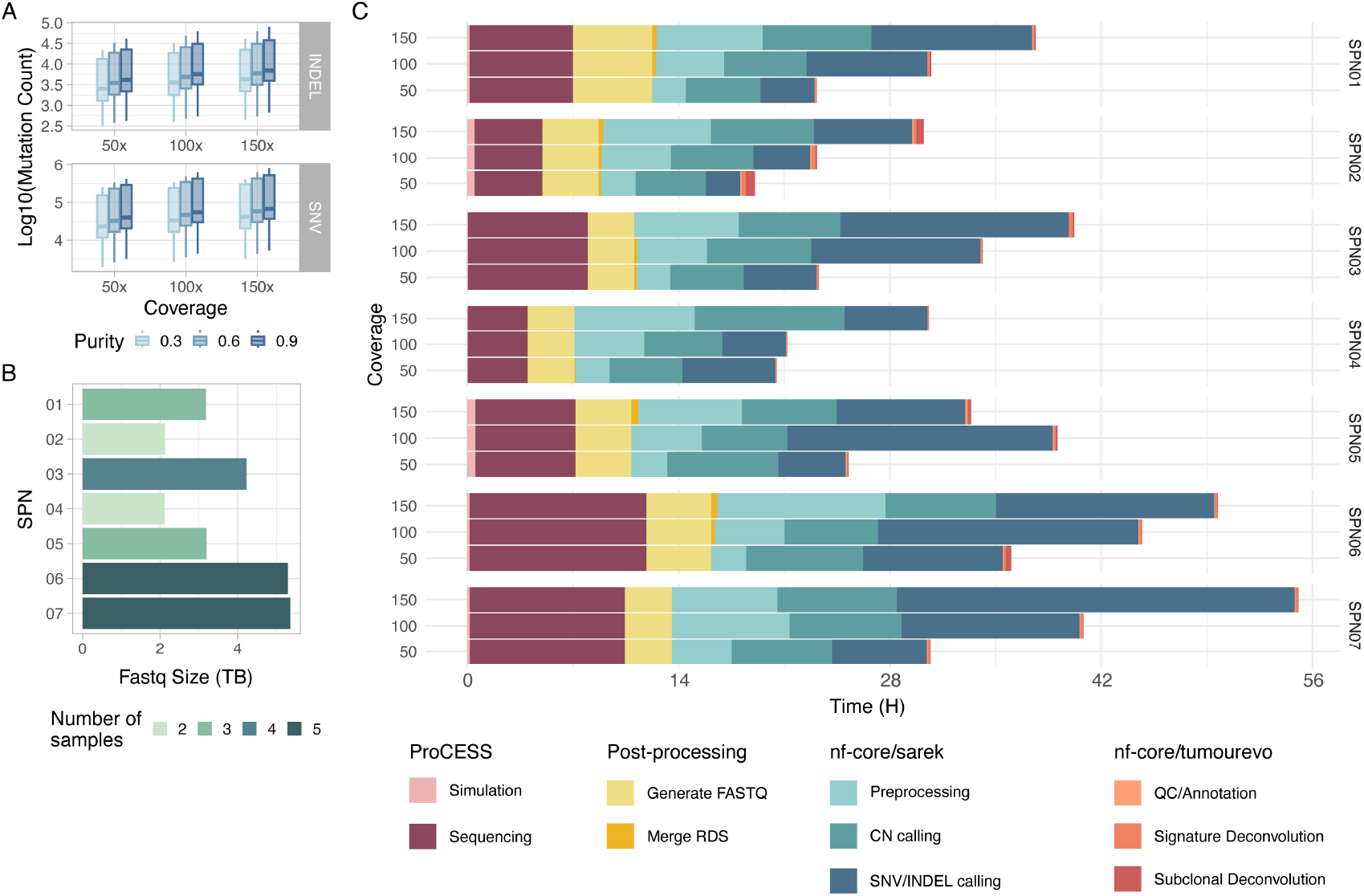
Statistics of the SCOUT dataset. (A) Distribution of simulated mutation counts across sequencing depths (50×, 100× and 150×) and tumour purity levels (0.3, 0.6 and 0.9), shown separately for SNVs and indels, illustrating the effect of sequencing parameters on observable mutation burden. (B) Size of generated FASTQ files for each SCOUT tumour at all simulated sequencing depth and purities, highlighting variability in data volume across tumour types and number of collected samples. (C) Timeline for the SCOUT cohort, including ProCESS simulation and sequencing, preprocessing and FASTQ generation, nf-core/sarek copy number and SNV/indel calling, and downstream nf-core/tumourevo analysis, showing the computational workflow and runtime associated with the dataset.

**Extended Data Fig. 4:**
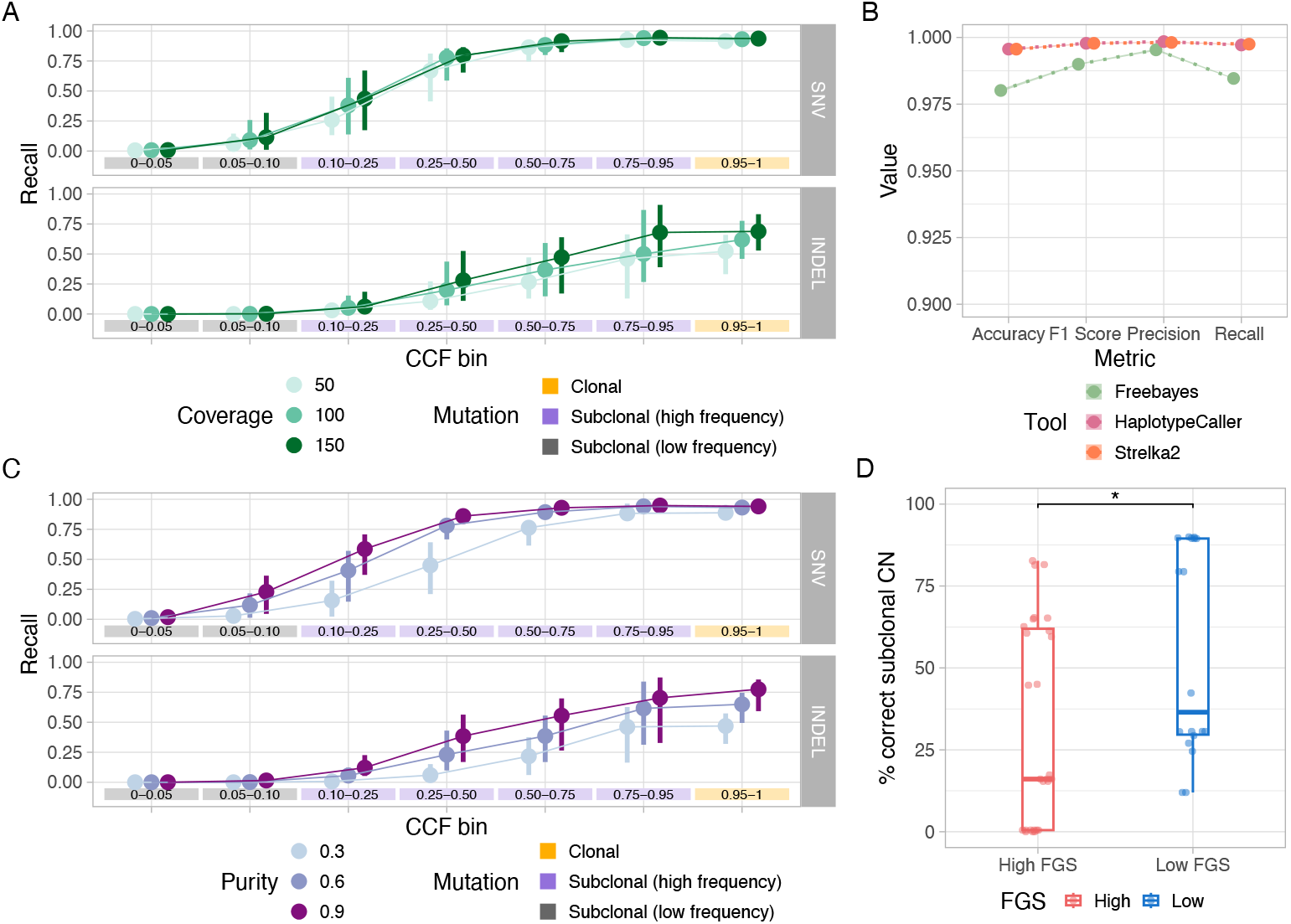
Variant calling performance across sequencing conditions. (A) Recall of SNV and indel detection as a function of cancer cell fraction (CCF), stratified by sequencing coverage (50×, 100× and 150×), for each variant caller (FreeBayes, Mutect2 and Strelka2). (B) Aggregate scores for the germline variant callers (FreeBayes, HaplotypeCaller, Strelka2). (C) Recall of SNV and indel detection as a function of cancer cell fraction (CCF), stratified by tumour purity (0.3, 0.6 and 0.9) and sequencing coverage (50×, 100× and 150×), for each variant caller (FreeBayes, Mutect2 and Strelka2). (D) Boxplot for the percentage of correctly detected subclonal CNAs by Battenberg as a function of fraction of genome subclonal (FGS).

**Extended Data Fig. 5:**
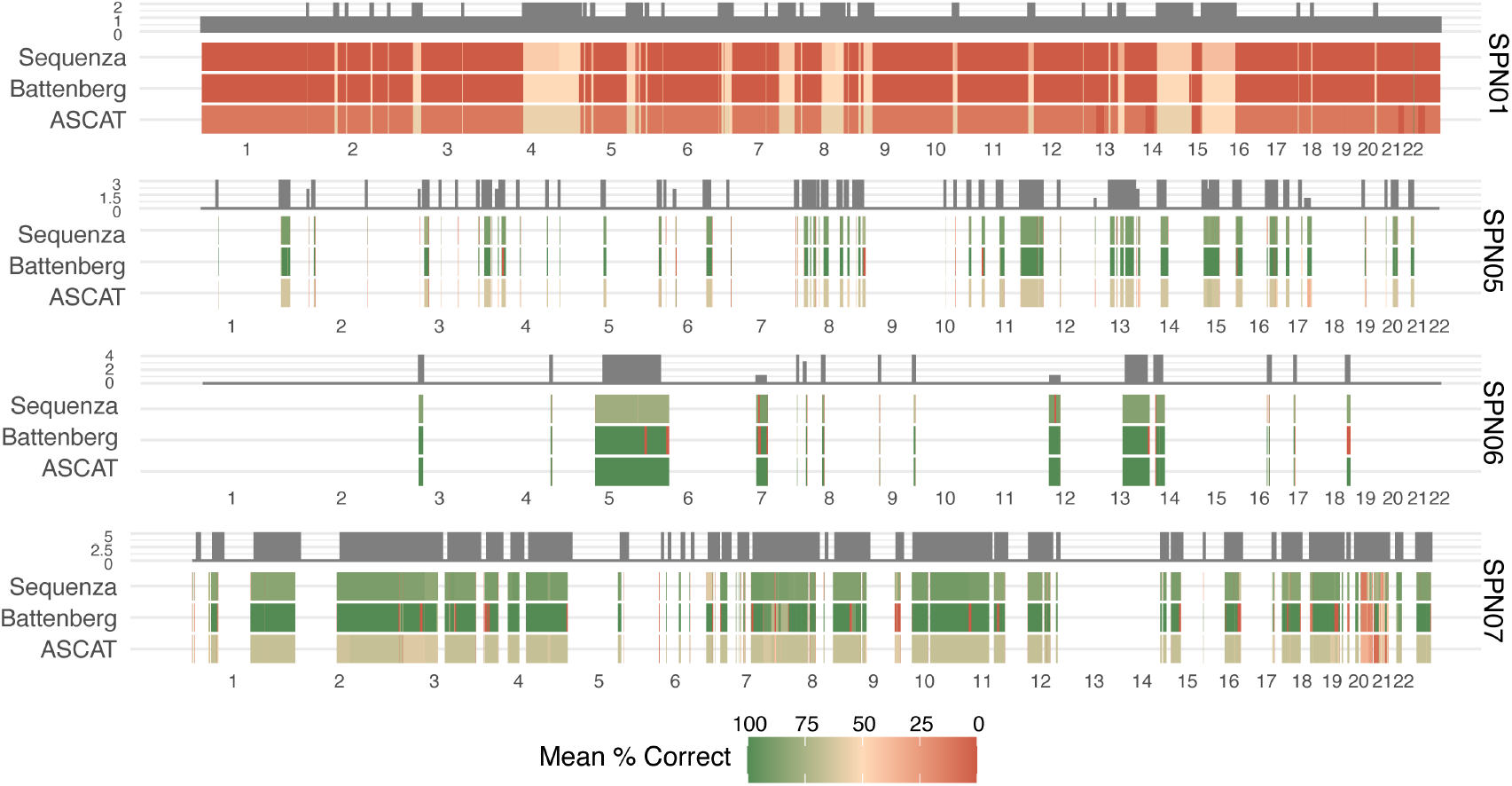
Reconstructed tumour genome profiles of high FGA patient. For high-FGA SPNs (01, 05, 06 and 07), the plot reports the mean percentage of correctly inferred clonal copy number events across the three copy number callers, ASCAT, Sequenza, and Battenberg. These values are aggregated over purity, sequencing coverage, and samples. The frequency of events within the SPNs is shown in the upper part of the plot.

**Extended Data Fig. 6:**
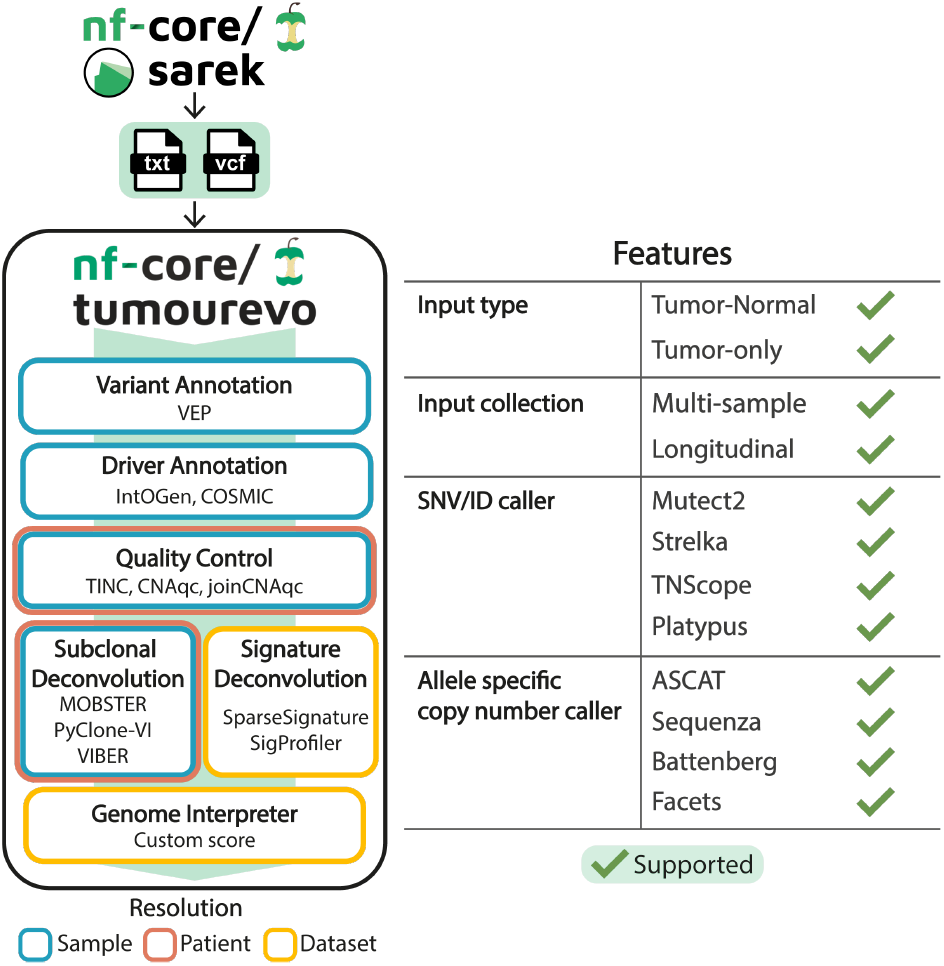
Main features of the nf-core/tumourevo pipeline. nf-core/tumourevo accepts as input VCF files and copy number call files generated by supported tools, either produced by nf-core/sarek or independent callers. The main workflows, implemented tools, and the resolution at which analyses are performed are shown.

**Extended Data Fig. 7:**
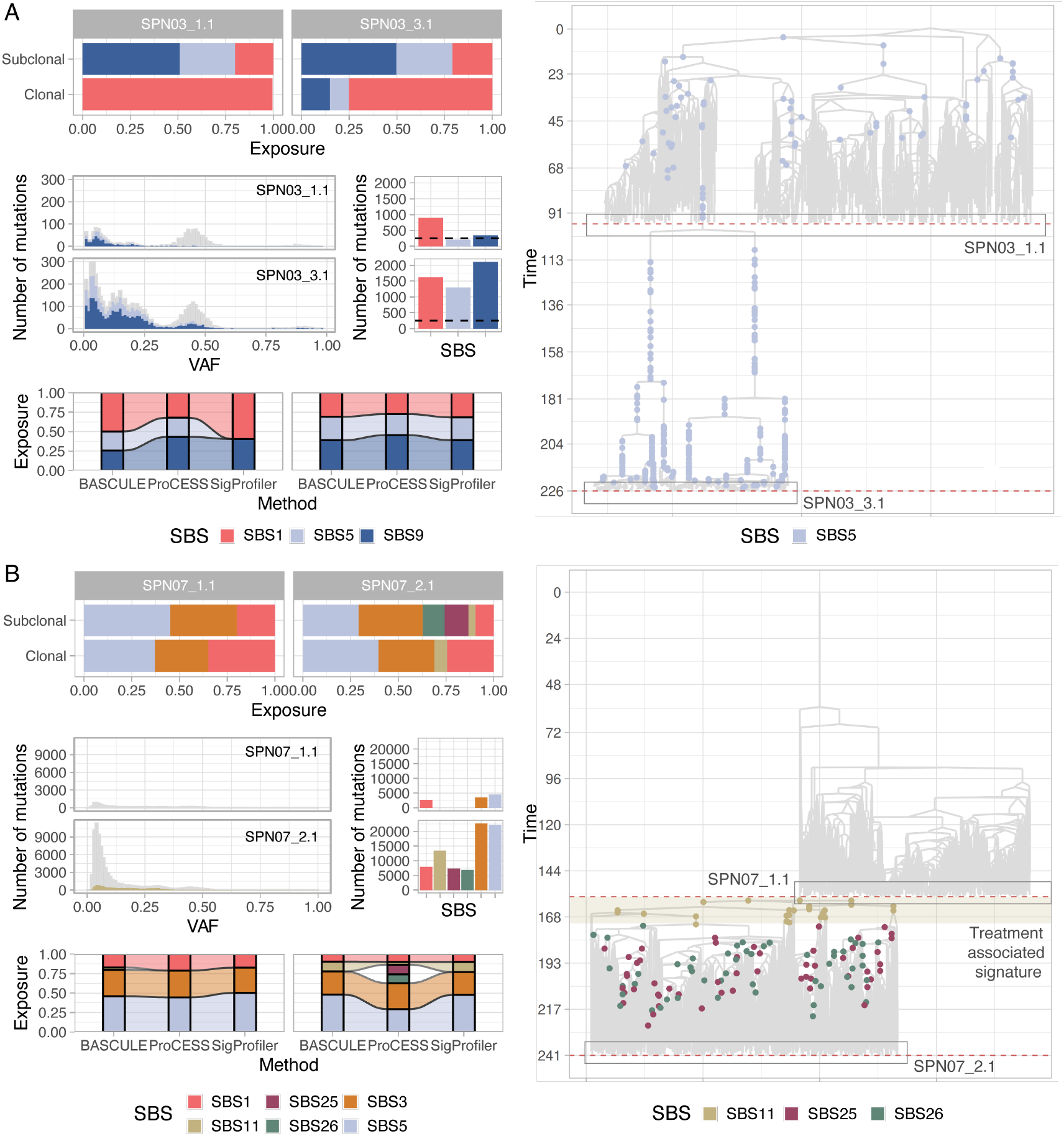
Performance of signature detection tools for subclonal mutational processes. (A) For SPN03 samples 1.1 and 3.1: simulated ground truth signature exposures in clonal and subclonal mutations; VAF spectra of mutations detected by Mutect2, with SBS5 and SBS9 highlighted; absolute mutation counts per signature, with a dashed line indicating the limit of detectability; ground truth (ProCESS) and inferred signatures and exposures by BASCULE and SigProfiler. The phylogeny annotates the samples and cells carrying the highlighted signature. (B) As in (A), for SPN07 samples 1.1 and 2.1.

**Extended Data Fig. 8:**
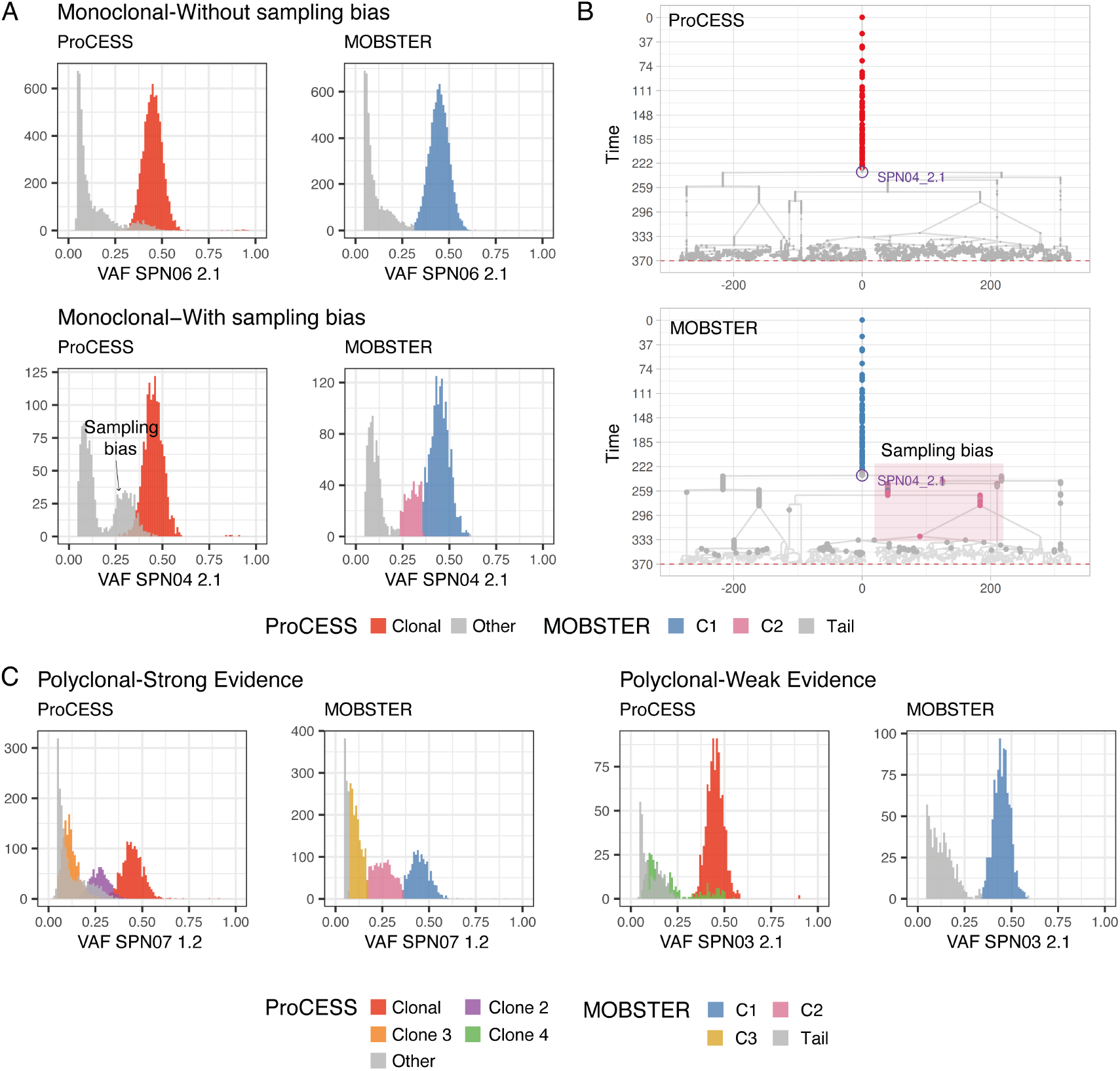
Sources of bias in univariate subclonal deconvolution. (A) In univariate subclonal deconvolution, monoclonal samples may (SPN04 2.1) or may not (SPN06 2.1) show evidence of sampling bias, confounding the inference of clonal composition by MOBSTER. (B) For SPN04 sample 2.1, affected by sampling bias, the ground truth phylogeny is shown with ground truth labels (top) and compared against the phylogeny inferred by MOBSTER (bottom), in which the sampling bias signal is erroneously fitted as a subclone. (C) In univariate subclonal deconvolution, weak evidence of subclonal signals in polyclonal samples (SPN03 2.1) confound the accurate inference of clonal composition by MOBSTER.

**Extended Data Fig. 9:**
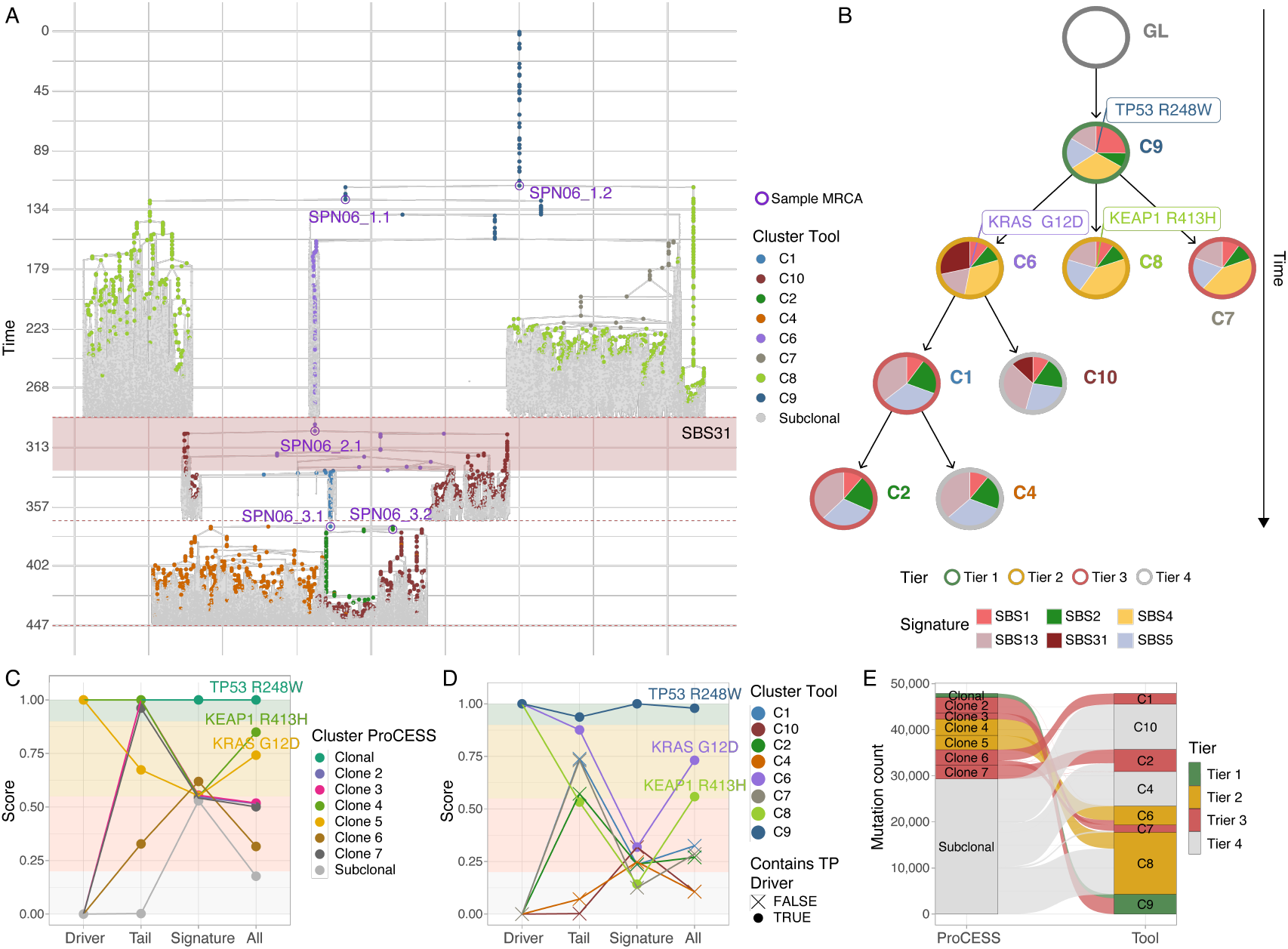
SPN06 genome interpreter. (A) Phylogeny from ProCESS simulation with annotated samples and their most recent common ancestor (MRCA). Nodes are coloured by the cluster inferred by nf-core/tumourevo subclonal deconvolution, with SBS31 treatment windows annotated. (B) Clone tree produced by nf-core/tumourevo subclonal deconvolution and interpretation, with signature proportions reported for each cluster. Nodes are also annotated by the Tier assigned by the genome interpreter score. (C) Ground truth ProCESS clusters with their corresponding scores and simulated drivers, used as reference for evaluating the genome interpreter score. (D) Genome interpreter score for clusters inferred by nf-core/tumourevo, annotated with the presence or absence of a true positve ProCESS simulated driver in each inferred cluster. (E) Sankey plot illustrating how simulated ProCESS mutations and their corresponding Tier assignments are distributed across clusters in the nf-core/tumourevo inference.

**Extended Data Fig. 10:**
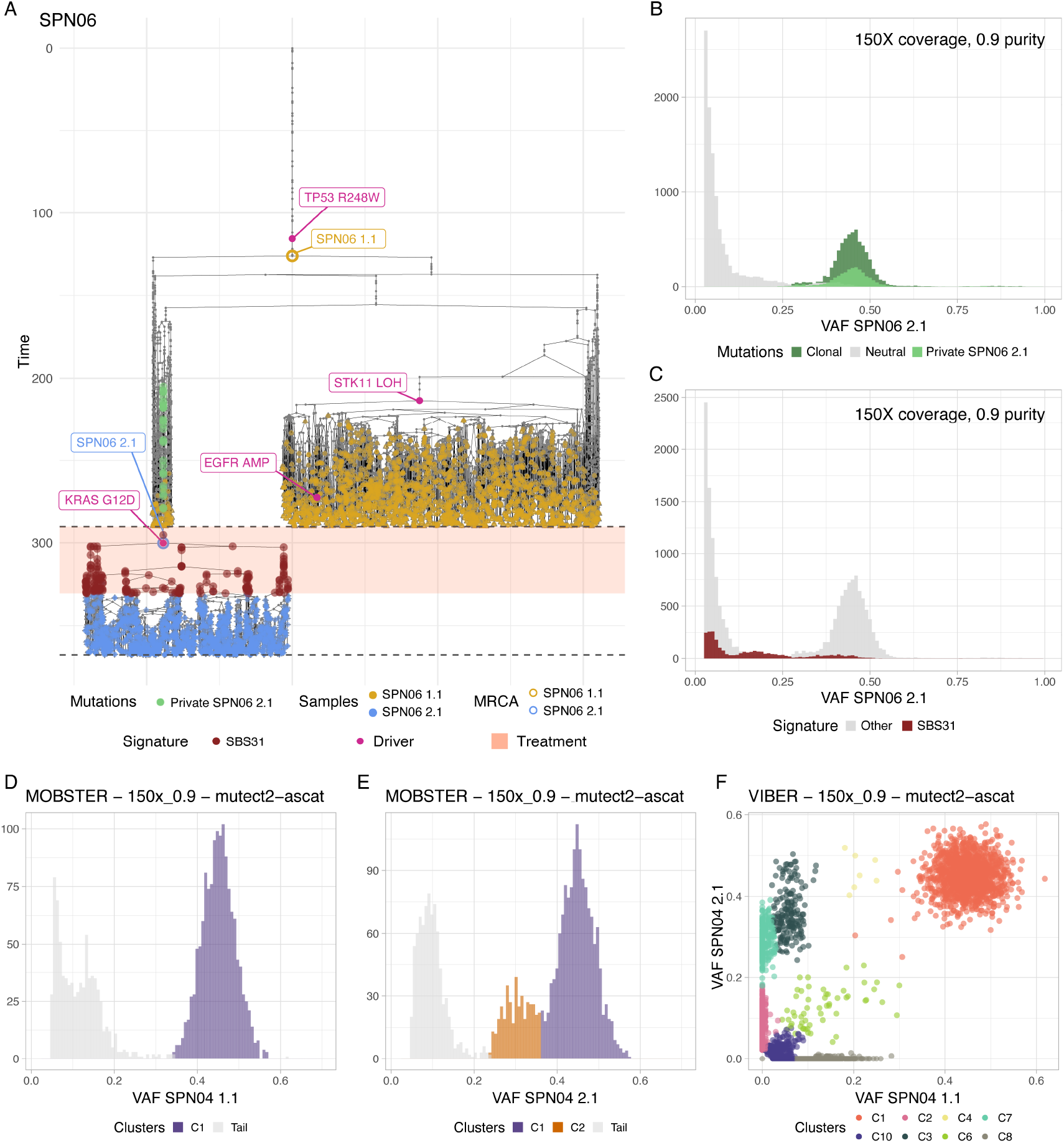
Mechanisms of therapy response. (A) Phylogeny of SPN06 with a pre-existing resistance mechanism, annotated with the most recent common ancestors (MRCAs) of both samples, per-timepoint sampled cells, mutations private to the post-treatment sample, and the treatment-associated SBS31 mutational signature. (B) VAF spectra resolving mutations by class: clonal-cluster, neutral and private to the relapsed sample. (C) VAF distribution of treatment-associated mutations, spanning both high and low frequencies, confirming their detectability in the relapse sample. (D,E) VAF deconvolution by MOBSTER for samples 1.1 (D) and 2.1 (E) of SPN04 with inferred clusters and neutral tail highlighted. (F) Multi-sample subclonal deconvolution by VIBER for SPN04, with inferred mutation clusters shown in distinct colours.

## A. Supplementary Notes

Detailed description of the ProCESS simulation framework, of the nf-core/tumourevo pipeline for tumour evolutionary analysis and of the analysis results of each SPN of the SCOUT cohort are available as Supplementary Notes.

## B. Supplementary Tables

**Supplementary Tables T8A–I** | **Simulation parameters and ground truth data for the** SCOUT **cohort**. Parameters governing ProCESS simulations and associated ground truth annotations for copy number alterations, mutational exposures, and driver events across all SPNs. (A) Kinetics parameters, foundation time, growth rate, and death rate, for each simulated clone per SPN. (B) Cell counts, sampling timepoints, and clonal proportions for each sample per SPN. (C) Simulated driver mutation rates (SNV, INDEL, and CNV) per SPN. (D) Simulated mutational signatures and their temporal exposures per SPN. (E) Sample-level metadata per SPN. (F) Ground truth allele-specific copy number profiles with corresponding cancer cell fractions (CCFs) per sample and SPN. (G) Ground truth driver mutations per SPN. (H) Ground truth driver mutations annotated with copy number status and CCF per SPN. (I) Ground truth mutational signature exposures stratified by sequencing coverage and tumour purity per sample and SPN.

**Supplementary Tables T9A–K** | **Benchmarking results of** nf-core/sarek **and** nf-core/tumourevo **for the** SCOUT **cohort**. Performance metrics evaluated per SPN, sample, purity, coverage, and tool across all benchmarked analyses. (A) Benchmarking results for somatic SNV and INDEL variant calling. (B) Benchmarking results for germline variant calling. (C) Benchmarking results for somatic clonal copy number calling. (D) Benchmarking results for driver annotation. (E) CNAqc quality control results for each analytical combination. (F) TINC estimates for each analytical combination. (G) Benchmarking results for mutational signature deconvolution. (H) Benchmarking results for univariate subclonal deconvolution. (I) Benchmarking results for multivariate subclonal deconvolution. (J) Bench-marking results for Genome Interpreter scores at the clone level. (K) Benchmarking results for Genome Interpreter scores at the mutation level. (L) Results of algorithm for automated classification of resistance mechanisms.

**Supplementary Tables T10** | **Zenodo DOI for ground-truth** ProCESS **data**, nf-core/sarek **and** nf-core/tumourevo **outputs for the** SCOUT **cohort**.

