## Supplementary Notes for "Tumour evolution as ground truth for cancer whole-genome sequencing"

SPN01 • SPN02 • SPN03 • SPN04 • SPN05 • SPN06 • SPN07

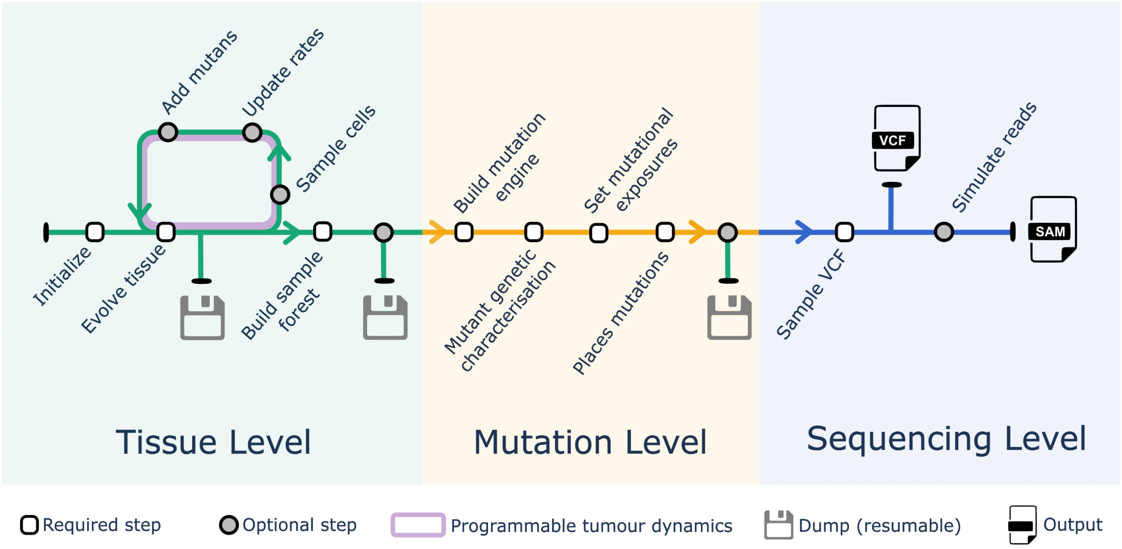

**Supplementary Fig. S1.** A schematic flow diagram of ProCESS/CLONES. The overall pipeline is split into three different levels. The tissue level deals with the simulation of the tissue and cell sampling. It abstracts from the cell genomic and takes care of the spatial and epidemiological aspects of the tumour growth, mimicking the interactions between different cell families. Furthermore, it simulates sampling and reconstructs sampled cell genealogy. The mutation level assigns each sampled cell a genealogy-consistent genome using a realistic, environment-aware mutation process. Finally, the sequencing level emulates sequencing and produces simulated reads and their alignments in standard SAM files.

#### A. The ProCESS spatial population genetics simulator

ProCESS is a tool that relies on CLONES (*C*++ *L*ibrary f*O*r *N*eoplastic *E*volution *S*imulations), a tool that simulates tumours and their analysis on different levels:

1. the *tissue level* reproduces the cancer growth on a simulated tissue, deals with cell sampling, and reconstructs sample cell families;
2. the *mutation level* simulates somatic alterations that accrue onto the tumour DNA during cell division, augmenting the intra-tumour heterogeneity (ITH). This level also labels the sampled cell with simulated genomes consistent with their spatial evolution history;
3. the *sequencing level* produces realistic artificial sequencing data for the simulated tumours.

ProCESS (*Pro*grammable *C*ancer *E*volution *S*patial *S*imulator) is an R package that wraps CLONES functions and data structures using RCpp [1]. ProCESS automates the simulation setup task and makes it easy to retrieve the required data and files. It also features many useful presentation functions to analyse simulation outcomes. The overall, step-by-step, functioning of ProCESS/CLONES is summarised in Supplementary Figure S1.

##### A.1. State-of-the-art in Cancer Simulation Tools

Genomic data simulators are essential for validating computational methods to both investigate tumour evolution and design effective personalised cancer treatments. Many different simulators have been proposed so far in the literature. Each of them is characterised by a specific approach and the corresponding limitations. *RSVSim* is an R package to simulate structural variations from a template sequence [2]. *SCNVSim* simulates somatic CNA and structural variations [3]. *VarSim* integrates read simulators, such as *ART*, and simulates high-throughput sequencing [4]. *BAMSurgeon* adds synthetic SBSs, indels, and structural variations on reads contained in a BAM file [5]. *tHapMix* combines different haplotypes to generate synthetic tumour samples that meet the specified polyclonality, purity, and ploidy [6]. *Pysim-sv* is a package for simulating high-throughput sequencing data that can introduce a wide range of germline and somatic genomic variations, accounting for GC biases [7]. *Xome-Blender* generates synthetic cancer genomes with user-defined features, such as diploidy, somatic variants, and CNVs [8]. *SVEngine* simulates next-generation sequencing data embedded with structural variations, allowing specification of locus-specific variant fractions and allelic haplotypes [9]. *J-SPACE* is a Julia package for simulating the spatial growth and genomic evolution of a cell population, as well as the sequencing experiment for the sampled cells [10]. It decouples tissue and genomic simulations, building only the phylogenetic tree for a simulated tissue sample. *cancerSimCraft* is an R package for simulating cancer genomes at both clonal and single-cell resolution, combining deterministic rules and stochastic processes while tracing ground truth [11]. *SISTEM* ses an agent-based approach for modelling metastasis at single-cell resolution [12]. Finally, *MOV&RSim* uses COSMIC and TCGA datasets for simulating 21 types of cancer [13].

Among the above tools, only J-SPACE simulates intra-tumour spatial dynamics and spatial-constrained biopsy mutation heterogeneity by representing cells on either a 2D or a 3D grid. SISTEM models tumour metastatic migrations, but it ignores the exact position of each cancer cell. cancerSimCraft, SVEngine, and MOV&RSim mimic clonal expansion according to a user-specified phylogenetic tree. The phylogenetic trees are instead built from stochastic simulations in SISTEM and J-SPACE.

cancerSimCraft uses a continuous-time Markov model to represent discrete cell events, such as birth and death, and allows the specification of distinct birth and death rates. SISTEM users specify genotype-related cellular lifespan and turnover rates. Instead, J-SPACE accepts stochastic parameters, such as birth and death rates, to govern the epidemiologic evolution of the cancer cohort. MOV&RSim models 21 different types of tumour and integrates COSMIC and TCGA data to simulate somatic SBSs and indels. J-SPACE supports COSMIC SBS mutational signatures [14]; users can specify the list of active signatures to simulate mutation accumulation consistently with the modelled biological processes. Passenger indels are instead simulated by using the Lavalette law.

SISTEM can simulate 2.5 million cells distributed over 5 metastatic sites in about 25 minutes on an Apple M3 MacBook Pro with 16GB of RAM by modelling each clone as a single agent. [12]. This approach scales up very well in terms of cell number, but abstracts from passenger mutations and cell locality. J-SPACE can instead simulate 200 000 cells in about 1.5 hours on an Intel® Xeon® Gold 6240@2.60GHz, and the time "*increases exponentially with respect to the number of simulated cells*" [10]. J-SPACE decouples tissue and genomic simulations. According to the authors, it builds the phylogenetic tree of a 1000-cell sample in about 6 minutes on the same hardware used to simulate the tissue. However, neither the setup passenger mutation rate nor the placed passenger mutations are mentioned in [10].

ProCESS/CLONES is a fully programmable R package for tumour simulation. It supports tissue, sampling and sequencing simulations. Tumour evolution in the tissue is governed by first-order formulas whose variables are mutant sizes, simulated time, and the number of duplications and deaths per mutant. To ensure scalability, tissue and genomic simulations are decoupled, as in J-SPACE, and simulated cells are not associated with genomic data at the tissue level. The cells are labelled by their genome only after the sampling, when the phylogenetic forest is built.

Supplementary Table T1 summarises the main differences between some of the most used genomic data simulators and ProCESS/CLONES.

**Supplementary Table T1.** Comparative Table of Cancer Genome Simulators

| Simulator | SBS Signatures | Indel Mutations | Germline Mutations | Programmability | Max Cell Scale | Spatial Representation |
|---|---|---|---|---|---|---|
| **cancerSimCraft** | Real data | Supported | Yes (SNP table) | Det. & stochastic rules | Thousands (scDNA-seq) | None |
| **SISTEM** | MSK-MET patterns | No (SBS & CNA only) | N.A. | Agent-based models | Millions (clonal eff.) | Discrete sites & migration graphs |
| **J-SPACE** | Temporal SBS | Yes (Lavalette law) | N.A. (spatial focus) | Julia (graph interaction) | $> 10^5$ cells | Arbitrary graphs or 2D-3D lattices |
| **MOV&RSim** | 21 cancer presets | Yes (signatures/presets) | Yes (manual in VAR file) | Guided pipelines / Docker | N.A. (genomic realism) | None |
| **SVEngine** | Meta-distributions | Yes (Structural INS/DEL) | Yes (complex germline) | Parallel divide-and-conquer | Thousands of variants | None |
| **Xome-Blender** | 1KGP data | Yes (heterozygous loci) | From 1KGP | Modular (Bash/R/C++) | N.A. (bulk/WES focus) | None |
| **ProCESS/CLONES** | Temporal linear comb. of COSMIC SBSs | Temporal linear comb. of COSMIC IDs | 1KGP SBS & indels | R/First-order logic | Millions of cells & Hundreds of mutants | 2D lattices |

#### A.2. Tissue level

**Spatial organisation of cells in a tissue.** CLONES models tissues as 2D squared lattices, i.e., grids. Every element of a lattice represents a cell in the tissue that is either a wild-type or a tumour cell. A coordinate system based on the grid's columns and rows identifies any cell in the lattice. A cell is *internal* if it does not lie on the first or last column/row of the grid. The grid is initially empty, and new tumour cells can be individually placed in it by specifying their positions.

We write $p_c$ to indicate the position of the cell $c$ in the lattice. Any internal cell $c$ with $p_c = (x, y)$ has 8 *neighbour cells* $c_{\uparrow}$, $c_{\nwarrow}$, $c_{\leftarrow}$, $c_{\swarrow}$, $c_{\downarrow}$, $c_{\searrow}$, $c_{\rightarrow}$, and $c_{\nearrow}$ whose positions are $(x, y+1)$, $(x-1, y+1)$, $(x-1, y)$, $(x-1, y-1)$, $(x, y-1)$, $(x+1, y-1)$, $(x+1, y)$, and $(x+1, y+1)$, respectively. By extension, we use the notation $c_*$ with $* \in \{\uparrow, \nwarrow, \leftarrow, \swarrow, \downarrow, \searrow, \rightarrow, \nearrow\}$ also when $p_{c_*}$ is not in the grid.

The 8 neighbours induces the *set of admissible directions* $\mathcal{D} \stackrel{\text{def}}{=} \{\uparrow, \nwarrow, \leftarrow, \swarrow, \downarrow, \searrow, \rightarrow, \nearrow\}$. The $*$-*directional distance of a cell c from the tumour border*, with $* \in \mathcal{D}$, is defined as:

$$d_{b,*}(c) \stackrel{\text{def}}{=} \begin{cases} 0 & \text{if } c \text{ is a wild-type cell} \\ 1 & \text{if } c \text{ is a tumour cell and } p_{c_*} \text{ is not in the grid} \\ 1 + d_{b,*}(c_*) & \text{if } c \text{ is a tumour cell and } p_{c_*} \text{ is in the grid} \end{cases} \quad (10)$$

The *distance of a cell c from the tumour border* is the minimum among the directional border distances, i.e.,

$$d_b(c) \stackrel{\text{def}}{=} \min_{* \in \mathcal{D}} d_{b,*}(c) \quad (11)$$

Even though the cells are not genomically characterised at the spatial evolution level, tumour cells are grouped according to their potentially unknown functionally relevant mutations. A *mutant* is a set of cells that have the same driver mutations. All the cells of a mutant have the same liveness rates, i.e., rates for cell division and death.

**Clonal organisation of cells.** Every tumour cell is labelled by its mutant, an identifier, unique among all simulation cells, and the cell birth time, which is the simulated time at which the cell was created (the initial cell is assigned time 0). Whenever a new tumour cell is created, its data are stored in an indexed binary log file named *cell information file*. This file is eventually queried to retrieve cell descendant information and build the sample forest of sampled cells.

CLONES reproduces tissue evolution by successively firing one of the three possible cell event types: death, duplication, and mutant evolution. The death event removes the original cell from the tissue and replaces it with a wild-type cell. Both duplication and mutant evolution events replace the original cell with two new cells, called child cells, but they differ in the mutant of the newly created cells. The cells created by duplications belong to the same mutant as their parent. On the contrary, one of the cells created by a mutant evolution belongs to its parent mutant, while the other one to a different mutant. The mutant evolution event is non-reversible, following the no-back mutations model: if one of the ancestors of a cell belongs to the cell mutant, then all descendants of that ancestor down to the cell belong to the same mutant.

Duplications and mutant evolution replace a single tumour cell with two cells. One of the two children replaces the original cell $c$ in the lattice, while the other is placed in one of the neighbouring positions. The position of the second child is randomly chosen according to the discrete probability distribution

$$\mathcal{P}\left(p_{c_*}\right) \stackrel{\text{def}}{=} \frac{1}{d_{b,*}(c)} \frac{1}{\sum_{x\in\mathcal{D}}(1/d_{b,x}(c))}. \tag{12}$$

Hence, the probability $\mathcal{P}(p_{c_*})$ for $p_{c_*}$ to be the second child position is inversely proportional to the distance between $c$ and the tumour border along the direction $*$.

If $p_{c_*}$ is not in the grid, the child is tagged as *lost* and removed from the simulation. If it is in the lattice, then either $c_*$ is a wild-type or a tumour cell. In the former case, the child cell replaces the wild-type cell in $p_{c_*}$. When $c_*$ is instead a tumour cell, CLONES makes room for the second child by pushing all the tumour cells along the direction $*$ up to the tumour border or the lattice border.

Mutant evolution events are triggered by users during the simulation. Instead, death and duplication are stochastic events whose rates depend on the cell mutant.

**Growth models.** Each tumour cell is either enabled or not enabled for a cell event. A *growth model* is the set of rules establishing whether a cell is enabled. CLONES implements the *homogeneous growth model* and the *border-driven growth model*. The homogeneous growth model enables all cells for all events (Supplementary Fig. S2a). In contrast, the border-driven growth model admits duplications and mutant evolutions only on the cells whose distance from the tumour border is 1, i.e., $d_b(c) = 1$, (Supplementary Fig. S2b). Both models enable all cells for the death event. *ProCESS* adopts the border-driven growth model by default. However, it can switch to the homogeneous model on request. See the growth model page on the ProCESS site for more details.

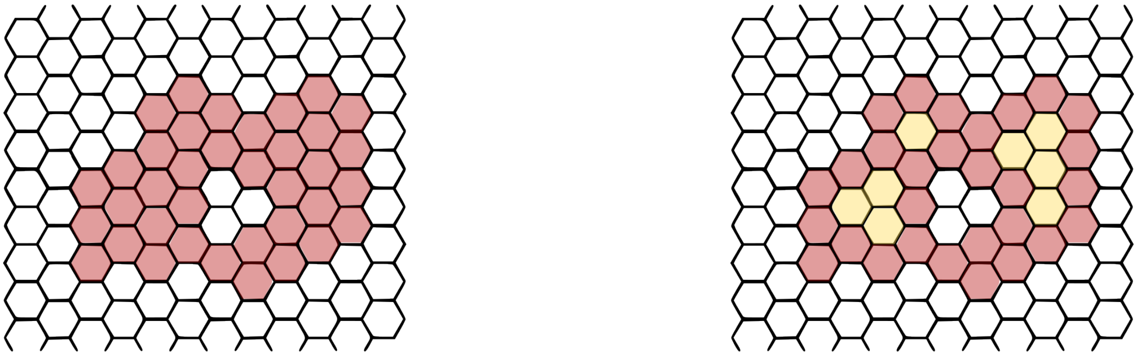

**(a)** The homogeneous growth model allows the duplication and the mutant evolution of any non-wild-type cell – the red cells in the figure – in the simulated tumour. In the figure, the white cells are wild-type cells.

**(b)** The border-driven growth model, instead, admits duplications and mutant evolutions exclusively on the tumour border – the red cells in the figure –. In the figure, white cells are wild-type cells. The orange and red cells are internal and border tumour cells, respectively.

**Supplementary Fig. S2.** The two growth models implemented in CLONES: the homogeneous growth model (Fig. S2a) and the border-driven growth model (Fig. S2b).

**Tissue temporal evolution.** The next event to be fired, its timing, and the affected mutant are selected based on the event rates and the number of cells enabled for the event by using the first-reaction method [15]. The duplication and death events are denoted by $\lambda$ and $\delta$, respectively, and $\lambda_j$ and $\delta_j$ indicate their rates for the mutant $M_j$. Moreover, let $n_{j,\lambda}$ and $n_{j,\delta}$ be the number of cells in $M_j$ active for duplication and death, respectively, and let $k$ be the number of mutants. Then, the timing $\tau$ of the next event is the minimum among

$$\tau_{j,\sigma} = -\frac{\ln j}{n_{j,\sigma} * \sigma_j} \tag{13}$$

where $j \in [1,k]$, $\sigma \in \{\lambda, \delta\}$, and $j$ is a uniformly random value in $[0,1]$. The event type $\sigma$ and mutant $M_j$ that will occurs after time $\tau$ are those that minimise $\tau_{j,\sigma}$. Once $j$, and $\sigma$ establied, CLONES randomly selects, with uniform distribution, one cell among the $n_{j,\sigma}$ cells enabled for the event $\sigma$, and it applies $\sigma$ to it.

**Constraining tissue simulation with logic.** The tissue simulation proceeds until a user-defined condition is met. ProCESS allows to specify the condition by a first-order unquantified logic having variables representing the cardinality of the mutants, the number of events fired in a mutant (being duplications or deaths), and the simulation time. These variables and real values form expressions by being summed (+), subtracted (−), and multiplied (∗). The expressions are then compared by using the relational symbols $>$, $>=$, $==$, $!=$, $<=$, and $<$ to form relational expressions. A formula in this language is either a relation, the conjunction of two formulas (&), or the non-exclusive disjunction of two formulas (|). Eq. 14 and 15 report the formal BNF syntax of the tissue simulation logic. Eq. 14 specifies the valid variable names for this logic, while Eq. 15 declares the syntax of the numeric and logic expressions.

$$\begin{aligned}
\texttt{<var_name>} &::= \texttt{<mutant_var_name>} \mid \textit{"Time"} \\
\texttt{<mutant_var_name>} &::= \texttt{<mutant_name>} \mid \texttt{<mutant_name>}\ \textit{".duplications"} \\
&\quad\mid \texttt{<mutant_name>}\ \textit{".deaths"} \\
\texttt{<mutant_name>} &::= [_a-zA-Z][_a-zA-Z0-9]+
\end{aligned} \tag{14}$$

All symbols in Eq. 15 are actual R objects and, in particular, the symbol `sim_var` in Eq. 15 represents the name of an R variable referring to a tissue simulation.

$$\begin{aligned}
\texttt{<logic_formula>} &::= \texttt{<logic_term>} \mid \texttt{<logic_formula>}\ \textit{"|"}\ \texttt{<logic_term>} \\
\texttt{<logic_term>} &::= \texttt{<logic_factor>} \mid \texttt{<logic_term>}\ \textit{"\&"}\ \texttt{<logic_factor>} \\
\texttt{<logic_factor>} &::= \texttt{<rel_expr>} \mid \textit{"("}\ \texttt{<logic_formula>}\ \textit{")"} \\
\texttt{<rel_expr>} &::= \texttt{<expr>}\ (\textit{"<"} \mid \textit{">"} \mid \textit{"=="} \mid \textit{"<="} \mid \textit{">="} \mid \textit{"!="})\ \texttt{<expr>} \\
\texttt{<expr>} &::= \texttt{<term>} \mid \texttt{<expr>}\ (\textit{"+"} \mid \textit{"-"})\ \texttt{<term>} \\
\texttt{<term>} &::= \texttt{<factor>} \mid \texttt{<term>}\ \textit{"*"}\ \texttt{<factor>} \\
\texttt{<factor>} &::= \texttt{<sim_var>}\ \textit{"\$var("}\ [\textit{"}]\ \texttt{<var_name>}\ [\textit{"}]\ \textit{")"} \mid \texttt{<number>} \mid \textit{"("}\ \texttt{<expr>}\ \textit{")"} \\
\texttt{<sim_var>} &::= [_a-zA-Z][_a-zA-Z0-9]+ \\
\texttt{<number>} &::= [0-9]+(.[0-9]+)?
\end{aligned} \tag{15}$$

The semantics of the tissue simulation logic is evaluated on the current state of the tissue simulation. The variable "*Time*" models the elapsed time, while the variables `<mutant_name>`, `<mutant_name>` "*.duplications*", and `<mutant_name>` "*.deaths*" represent the number of cells, and the number of duplications and deaths that occurred along the simulation, respectively, of the mutant having the name `mutant_name`. The numeric expressions and logic formulas have the usual semantics.

Once a formula is satisfied, ProCESS suspends the simulation. Then, users can update some simulation parameters, such as the duplication rate of a mutant, let a mutant evolve, or sample the simulated tissue, and proceed with the simulation or end it.

For instance, users can let the simulation modelled in the variable `sim` proceed until the mutant B consists of 1000 cells at least by using the formula "`sim$var("B") >= 1000`". The formula "`3*sim$var("A.deaths") >= 4*sim$var("C.deaths")`" holds when the number of deaths for mutant A is at least three-fourths of those for mutant C. Users can also combine different relation expressions by using the logic operator "`&`" (the conjunction "and") and "`|`" (the non-exclusive disjunction "or"). For instance, they can let the simulation run for at least 100 simulated time units and until both mutants A and B individually consist of at least 10 000 cells, or they cumulatively account for 40 000 cells by using the formula

```
sim$var("Time")>=100 & ((sim$var("A") >= 10000 & sim$var("B") >= 10000)
                         | sim$var("A") + sim$var("B") >= 40000)
```

See the formula-based simulation constraints page on the ProCESS site for more details.

**Sampling simulated cells.** CLONES can sample tumour cells at any time during the simulation. It stores the sampled cells in an internal data structure and removes them from the tissue. Then, the tumour evolution can proceed. Samples are sets of tumour cells. ProCESS allows users to either collect all the tumour cells in a rectangle of tissue, which simulates a solid biopsy, or uniformly sample a subset of cells within the rectangle, mimicking a liquid biopsy.

ProCESS allows users to search tissue rectangles containing at least a specified number of cells per mutant. For instance, they can search in the simulated tissue a 70 × 70-rectangle of cells containing at least 100 cells of mutant D and 120 cells of mutant B by issuing the R line

```
sim$search_sample(c("D" = 100, "B" = 120), 70, 70)
```

where `sim` is an R variable referring to the tissue simulation (e.g., see Fig. S3c).

**Sample forest.** The sampled cells and their ancestors, which were previously stored in the cell information file, can be used to build the *sample forest*. Each node of the forest represents a cell of the simulation, and its parent corresponds to the parent cell. The cells initially placed in the simulated tissue by the user are embodied by the forest roots, while the forest leaves model the sampled cells (e.g., see Figure S3f).

Supplementary Fig. S3a, S3b, and S3c depict three instants of a tumour evolution involving four mutants and three samples simulated by ProCESS. The space left after collecting the samples is visible in Supplementary Fig. S3b and S3c). Supplementary Fig. S3f represents the sample forest.

**ProCESS's getters and plots** ProCESS provides getter functions to collect simulation data. It also offers plotting functions to represent the simulated tissue (e.g., Supplementary Fig. S3a), the composition of the tumour mass (e.g., Supplementary Fig. S3d), and the cardinalities of the mutants over time (e.g., Supplementary Fig. S3e).

##### A.3. Mutation level

CLONES simulates the genomic mutations of the sampled cells according to the sample forest: a mutation occurring in a cell also appears in its progeny. The *mutation engine* is the CLONES component devoted to labelling each node in the sample forest. It recursively visits the sample forest from its roots to the leaves, places mutations along the forest branches, and builds the *phylogenetic forest*. The phylogenetic forest also contains a list of sampled cell mutations, and a map that associates any mutation with the cell in which it occurs for the first time: the *first-occurrence map*.

**Mutation taxonomy.** A mutation is *germinal* when it is present in the germline, and therefore appears in all host cells that did not lose it because of a deletion. A mutation is *somatic* if it is not germinal. They affect all cells in the phylogenetic forest. Somatic mutations are partitioned into *driver mutations*, which genomically characterise the simulated mutants and rule their liveness rates, and *passenger mutations*, which occur at a mutant-specific rate and do not significantly affect the cell phenotype. The *pre-neoplastic* mutations are the passenger mutations that arise before the first tumour cell.

**Mutation types.** CLONES implements three kinds of mutations: single base substitution (SBS), indel (ID), and copy number alteration (CNA). Furthermore, it supports, as a driver mutation event, whole-genome doubling (WGD), which is a complex mutational event consisting of a complete doubling of all the alleles in the genome chromosomes. In the following part of this section, we may write SID, meaning a mutation that is either an SBS or an ID.

**Infinite site model.** A *locus* is a position in the genome. One locus may correspond to distinct bases in different alleles. A *region* is a sequence of contiguous loci. A *mutation specification* is a mutation labelled with the locus and allele in which it occurs. The *context of a mutation* is the minimal region containing the loci just before and just after the mutation reference sequence. The *infinite site model* assumes that only one SID can independently occur in a locus, even if the locus corresponds to different alleles. CLONES adopts *infinite site model* by default, and it also supports the non-infinite site models. ProCESS implements an easy-to-use interface to switch between these two models.

A locus is *available for mutation* if either no mutation occurs in the context of the locus in any allele or in some allele when the infinite site model is adopted or not adopted, respectively. A locus is considered *available for mutation* under two conditions: when no mutation is present in the context of the locus across any allele, if the infinite site model is adopted, or when there is at least one allele in which its context does not contain any mutation, if the infinite site model is not adopted.

**Representing cell genome.** CLONES represents the DNA of each sampled cell. However, it does not maintain the complete DNA sequences. Instead, it stores, for each cell, the mutations from the reference genome. Cellular mutations are retained hierarchically to mimic DNA organisation. At the highest level, we have the genome, which is partitioned into chromosomes. The chromosomes are represented by a potentially empty set of alleles. Any allele is a set of non-overlapping allele fragments indexed by their positions and storing their SIDs. This representation makes CNA simulations easy to implement. Duplications are obtained by copying the duplicated allele fragments into new alleles, while deletions are modelled by removing the target fragments from alleles.

At each level, the data structures maintain references to lower-level objects rather than storing full instances. This approach speeds up simulations by using lazy copying: initially, lower-level object references are copied rather than the entire object, and a full duplication occurs only upon request. For instance, when a cell genome is lazy-copied, the new instance refers to the same chromosome as the original object, and no new chromosome object is created.

**Placing mutations in cell DNA.** CLONES places mutations in cell DNA according to their specifications. Thus, it not only requires the mutation type and locus, but also the aimed allele identifier. This choice is consistent with germinal mutation descriptions, which are usually allele-specific, and supports relative driver placement, e.g., mutations *a* and *b* lie on the same allele. The special allele identifier `RANDOM_ALLELE` is used by default to provide a valid specification for the mutations whose allele identifiers are not relevant or unknown, e.g, passenger mutations. In these cases, CLONES records the mutation in an allele that is uniformly sampled among those available for the mutation and updates the allele identifier in the mutation specification.

**Mutational signatures.** *Mutational signatures* are discrete probability distributions of the somatic mutation types [16]. There are 6 different classes of mutational signatures affecting different kinds of mutations: the Single Base Substitution (SBS)

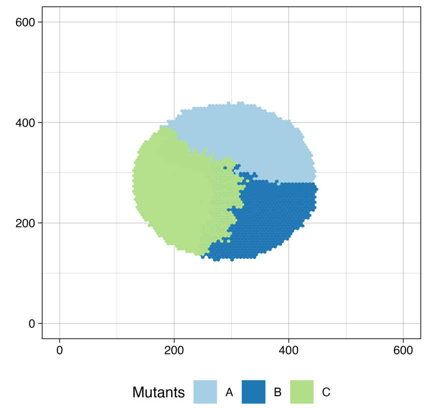

**(a)** Initially, the tissue contained only one cell of mutant A. The mutant grew up to 10 cells, and a new mutant B arose. When B accounted for 30 cells, a more aggressive mutant C emerged from A and surged up to 25 000 cells.

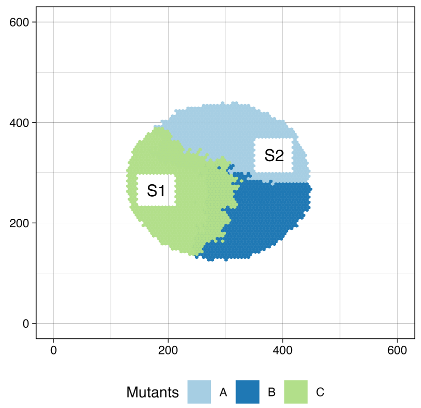

**(b)** Two 71 × 71 samples, S1 and S2, were collected from the tissue. S1 mainly contains C cells, whereas S2 consists of A cells. The sample labels are not automatically generated by ProCESS and were added for clarity.

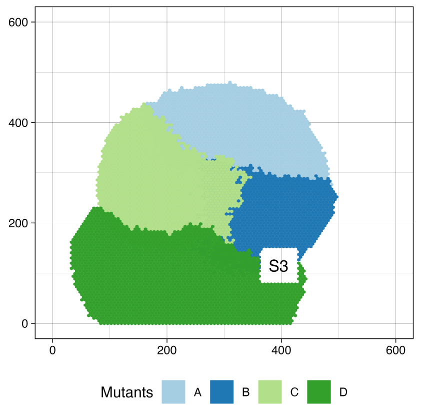

**(c)** After the sampling of the tissue, an even more aggressive mutant D arose as a sub-clone of B, and the simulation evolved until C and D globally accounted for 100 000 cells. A new sample S3 containing both D and B was collected.

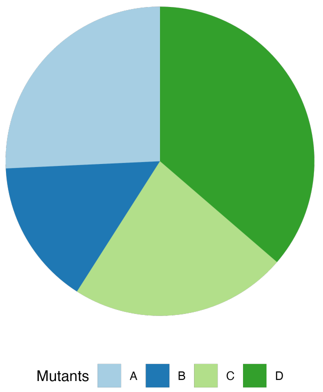

**(d)** The ratio among mutant cardinalities at the end of the tissue simulation.

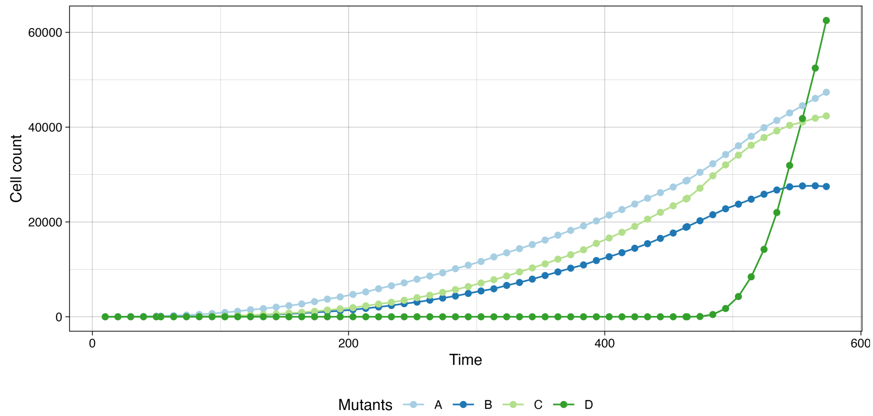

**(e)** The evolution of the mutant cardinalities along the simulation. We can observe a decline in the growth rates of mutants A, B, and C as D surges. This is due to the aggressiveness of D, which tends to fill all the non-tumour spaces on the lower border of the mass.

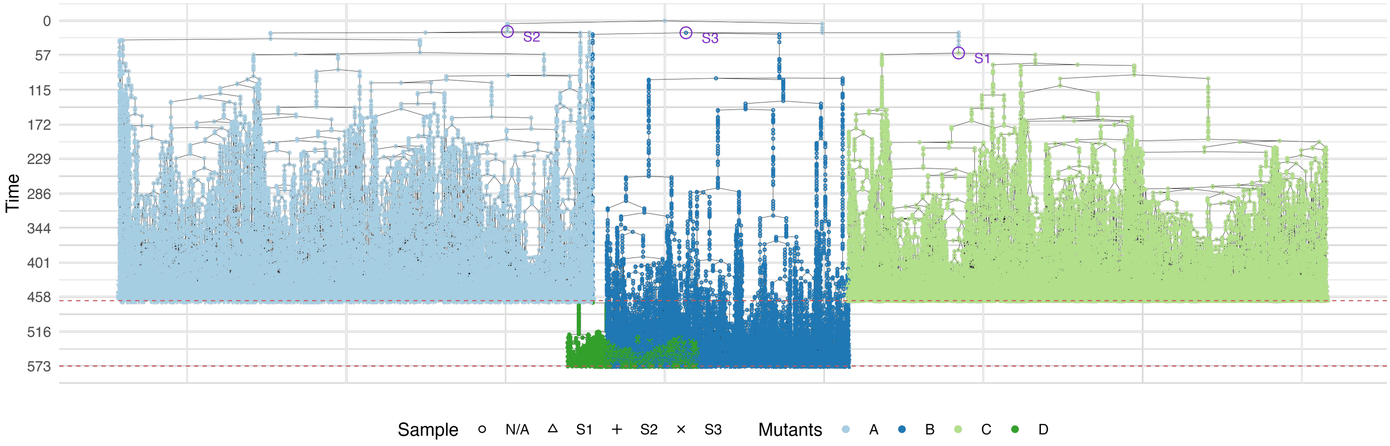

**(f)** The sample forest associated to the collected samples S1, S2, and S3. The nodes represent the samples' cells and their ancestors. The node colour corresponds to the cell mutant. The most recent common ancestor (MRCA) of each sample is labelled by the sample name. The $y$-axis represents the simulated time. Samples S1 and S2 were collected at about 458 simulated time units. Instead, S3 was collected at about 573, that is why the sub-tree of S3 is deeper than those of S1 and S2.

**Supplementary Fig. S3.** A ProCESS tumour model involving four mutants without epigenetic state.

signatures, the indel (ID) signatures, the Doublet Base Substitution (DBS) signatures, which rule the concurrent changes of two consecutive bases in the DNA, the Copy Number Variation (CN) signatures, related to CNAs, Structural Variation (SV) signatures, affecting large genomic changes involving the arrangement of the genome, and the RNA Single Base Substitution (RNA-SBS) signatures, that deal with RNA changes. Currently, CLONES supports SBS and ID signatures, and we plan to take care of DBS and CN signatures in the future.

The SBS mutational signatures establish the probability of an SBS and depend on both the SBS and its context. Each mutation in the SBS mutational signature is a class of mutations grouping a mutation and its inverse complement, e.g., the class `A[C>T]G` includes both `A[C>T]G` and `C[A>G]T`.

The ID mutational signatures are discrete probability distributions on the ID and the features of the repeated region in which the ID occurs. The signatures distinguish between IDs on homopolymers, repeated heteropolymers, and microhomologies. As far as the homopolymers are concerned, the ID probability depends on the base and on whether the ID is an insertion or a deletion. The probability of an ID on the heteropolymers, instead, depends on the number of repeated units, the size of the repeated unit, and the type of ID. Finally, only deletions are considered on microhomologies, and their probabilities depend on the length of the deletion.

A proposed aetiology characterises each mutational signature. For instance, the SBS signature SBS1 depicts an endogenous mutational process due to the age of the cell, and SBS4 is related to tobacco smoking [14].

**Mutational exposures.** *Mutational exposures* refers to the extent of exposure to multiple factors that can cause specific mutations. Each factor corresponds to a mutational signature. CLONES supports mutational exposures and models them as a probability distribution over a set of mutational signatures. The higher the probability of a signature, the more exposed the cell is to the factor associated with it. If $\mathcal{S}_1, \dots, \mathcal{S}_e$ are the signatures available for a mutation type $t$, a mutation exposure $\mathcal{M}_t$ for that type is a function $\mathcal{M}_t : \{\mathcal{S}_1, \dots, \mathcal{S}_e\} \to [0, 1]$ such that $\sum_{i=1}^{e} \mathcal{M}_t(\mathcal{S}_i) = 1$.

Users can independently declare mutational exposures for SBS and ID, specifying their validity in a temporal interval. So that the mutations of type $t$ emerging in a cell are distributed among the signatures of the exposure of $t$ that is valid when the cell is born. Section A.3.4 details how CLONES establishes the number of new mutations to be placed in a cell and the process that distributes them among the signatures.

##### *A.3.1. Germinal mutations*

CLONES supports SBS and ID germinal mutations and builds the germinal genome according to a specified VCF file. The mutation engine reads the VCF file and places the mutations in an object whose chromosomes and alleles are consistent with the reference genome. ProCESS provides a pair of pre-configured setups for the mutation engine containing a subset of the germlines published by the 1000 Genome Project [17] and allows users to select the germline of a specific individual.

##### *A.3.2. Passenger mutations*

**SID mutations and genome indices.** CLONES uses SBS and ID mutational signatures to simulate the passenger mutations. When an SBS is requested, the SBS and its context are randomly chosen according to the signature. Then, a locus consistent with the selected context must be found. Analogously, placing an ID requires CLONES to find a locus that satisfies the repeated sequence structure randomly selected according to the active ID signature. Because of the above reasons, the mutation engine builds two indices for the reference genome: one to quickly identify loci by their context bases (*context index*) and the other to retrieve loci by their repeated sequence type (*repeated sequence index*). These indices are built during the initialisation of the mutation engine and saved to disk. Every successive creation of an instance of the same mutation engine loads the saved indices, instead of building them. In order to reduce memory requirements, the two indices are pre-sampled to store a fraction of the original loci. ProCESS pre-samples 1 locus every 100 per context type in the context index and stores at most 500 000 loci per repeated sequence type in the repeated sequence index by default. Users can customise their values during mutation engine creation (see `MutationEngine` manual).

During the initialisation phase, the mutation engine is provided with a list of known driver mutations, and their loci are removed from the two indices. This step ensures that no passenger mutation is placed in a known driver mutation locus. ProCESS provides a set of driver INDEL and SNV mutations in the pre-configured mutation engine setups. This set consists in a curated list of driver genes for each tumour type was generated by integrating data from the COSMIC Cancer Gene Census (v102) and the IntOGen database (release 2024.09.20). The COSMIC Census Gene Mutations list was downloaded and filtered to retain only high-confidence somatic variants. Variants were excluded according to the following filters:

1. the `MUTATION SOMATIC STATUS` was annotated as *Confirmed somatic variant* or *Reported in another cancer sample as somatic*;
2. the `MUTATION DESCRIPTION` corresponded to *synonymous variant*, *3 prime UTR variant*, *5 prime UTR variant*, *intron variant*, or if the field was empty;
3. unknown amino acid changes `MUTATION AA`;
4. the `CDS` entries were labelled as *delins* or *inv*

Tumour type labels were assigned using the drivers provided by IntOGen. Genes were selected if the `CGC CANCER GENE` field was marked as True. The filtered IntOGen gene set was then joined with the curated COSMIC mutation list to associate each driver gene with its corresponding tumour type. This process yielded 138 876 driver mutations across 218 genes, mapped to 78 tumour types. ProCESS users can provide a custom set of known driver mutations during the initialisation of the mutation engine.

When a passenger SID is requested, the mutation engine randomly samples its reference and alternative sequences according to a mutational signature. Then, it identifies all the loci in the genome that are compatible with the chosen mutation using the appropriate index. The loci are uniformly sampled until the selected one is available for mutation. Finally, the mutation is placed and, if its locus is no longer available for mutation, it is removed from the indices.

**CNA mutations.** The passenger CNAs are, instead, uniformly sampled in a set of CNAs provided during the mutation engine initialisation. The set is partitioned by tumour type, and users can specify which tumour they are simulating to select only tumour-specific passenger CNAs. ProCESS packages a set of CNAs in the pre-configured mutation engine setups. A list of passenger copy number alterations (CNAs) was constructed using data from the Pan-Cancer Analysis of While Genome (PCAWG, 2778 samples) and Hartwig Medical Foundation (HMF, 3538 samples) cohorts. For PCAWG, we downloaded CNAs data from Zenodo [DOI:10.5281/zenodo.6410935], previously analysed with CNAqc [18] and categorised as either clonal or subclonal. For each patient, all segments with copy number states (CN) were retained, including the associated study and tumour type. For subclonal CNAs, the CN state of the clone with the highest cancer cell fraction (CCF) was selected. For the Hartwig cohort were segmentations files using the request forms that can be found at Hartwig Medical Foundation application form. We manually converted tumour-types in both PCAWG and HMF cohorts in order to match IntOGen database classification, the same used to identify driver mutations (section A.3.3). Copy number segments were then filtered according to the following principles:

1. Total copy number cannot exceed 15;
2. Segment length higher than 500 kbp.

This filtering process resulted in 375 244 CNA segments, with a median segment length of 3.7 Mb, an average total copy number of 4, and covering 63 tumour types.

##### *A.3.3. Driver mutations and genetic characterisation of mutants*

The tissue simulation abstracts genomics from the mutant specifications. Therefore, users must genetically characterise mutants and inform the mutation engine of the driver events – both mutations and WGD – and the SBS, ID, and CNA rates for each mutant. The mutation engine uses this data to generate the somatic mutations for each sampled cell, based on its mutant and those of its ancestors.

ProCESS allows to specify each driver mutation by reporting its locus, allele, reference and alternative sequences or, limited to the set of known driver mutations, by using its name.

##### *A.3.4. Simulating cells' DNA*

For each root in the sample forest, the mutation engine builds an object to model the genome mutations of the cell associated with that root. Then, it chooses a user-specified number of pre-neoplastic SBSs and IDs according to the age-related mutational signatures SBS1 and ID1, and places them. The resulting object serves as a template for the mutations in the genomes of the cells represented by the phylogenetic forest nodes.

The mutation engine recursively traverses the sample forest from the root to the leaves, processing each node in turn. As soon as a sample forest node is considered, the mutation engine builds an equivalent node in the phylogenetic forest and labels it with a copy of the parent node's genome mutations (the template genome mutations for forest roots). If the parent cell belongs to a different mutant, the mutation engine also applies the user-specified driver mutations.

The number $m$ of passenger mutations emerging during the cell duplication is sampled from a Poisson distribution

$$\mathcal{P}_{\mu_m}(X = m) \stackrel{\text{def}}{=} \frac{\mu_m^m}{m!} e^{-\mu_m}. \tag{16}$$

The distribution mean $\mu_m$ is the product of the amount of DNA in the cell genome and the mutant-specific mutation rate declared when mutants are genomically characterised (see Section A.3.3). It is important to stress that the amount of DNA, $D$, differs from the genome length $G$: the former is the mass of DNA in a cell and is affected by the number of alleles; the latter is allele-agnostic and is the sum of the longest alleles per chromosome. From the software perspective, CLONES measures the amount of DNA as the sum of the lengths of all allele fragments in the genome. For instance, the amount of DNA in a human wild-type cell is twice its length because all chromosomes, except chromosome X in men, have two alleles.

The mutation engine associates each of the passenger mutations with a mutational signature by sampling a multinomial distribution

$$\mathcal{P}_{p_1,\ldots,p_e}(X_1 = x_1, \ldots, X_e = x_e) \stackrel{\text{def}}{=} \frac{\left(\sum_{i=1}^{e} x_i\right)!}{\prod_{i=1}^{e}(x_i!)} \prod_{i=1}^{e} p_i^{x_i} \tag{17}$$

whose probabilities $p_1, \ldots, p_e$ are defined by the mutational exposure that is active at the birth of the affected cell. Then, the mutations are applied to the cell genome as described in Section A.3.2. Instead, the CNAs are uniformly sampled in the

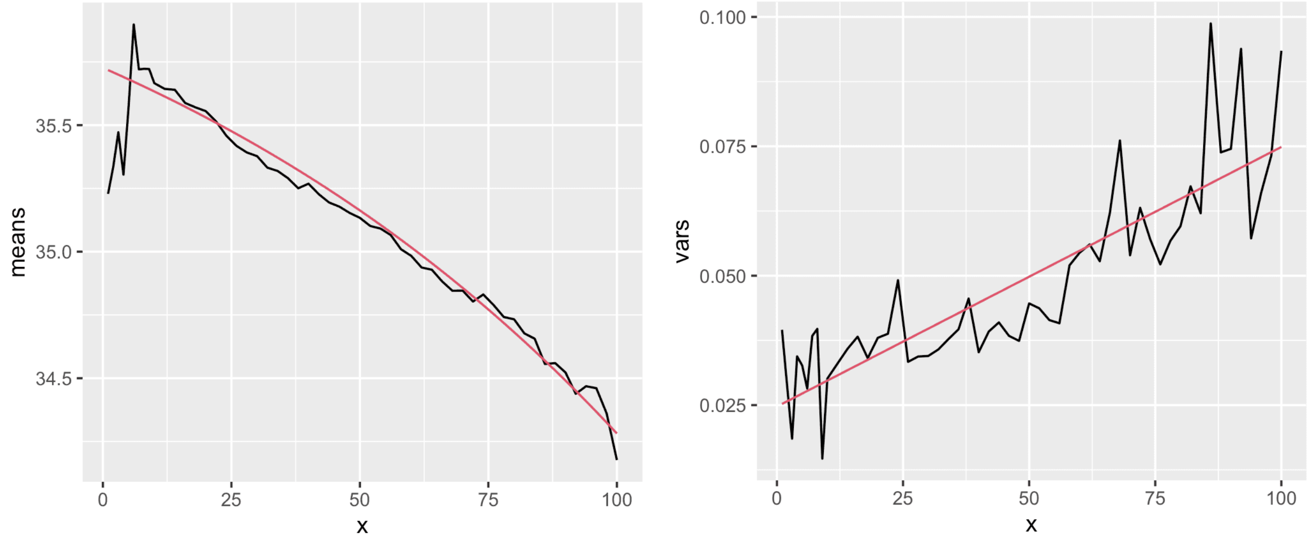

**(a)** The mean per read position of the Phred quality scores of the considered data set. The smooth line represents the best-fitting quadratic function for the means $\mu_q(x)$.

**(b)** The variance per read position of the Phred quality scores of the considered data set. The smooth line represents the best-fitting quadratic function for the standard deviations $\sigma^2_q(x)$.

**Supplementary Fig. S4.** In order to model the base quality score, we considered a real cohort of Illumina reads having size 100. The mean and the variance of the quality scores per base position $x$ were fitted into two quadratic functions, $\mu_q(x)$ and $\sigma^2_q(x)$, that model the relation between the parameters of the normal distribution ruling the quality score and the base position on the read. These functions can be scaled to mimic the quality scores of reads longer than 100 bases.

passenger CNA set maintained by the mutation engine and applied in agreement with Section A.3.2. After applying all the somatic mutations to the node, its children are considered. The mutation engine completes the generation of the forest on the leaves and labels them with the corresponding `GenomeMutations` objects.

#### A.4. Sequencing level

CLONES can simulate DNA sequencing of the sampled cells by sampling reads from the reference genome and reconstructing the corresponding sampled-cell reads based on the phylogenetic forest mutations that lie on them. The number of templates to simulate for each sample is computed as the product of the target coverage and the reference genome size, divided by the product of the number of reads per template and the read length. The total amount of DNA per sample is also evaluated as the sum of the sizes of all the allele fragments in the sample. The number of templates coming from each allele fragment is sampled from a binomial distribution $B(n'_t, p_t)$ in which $n'_t$ is the number of templates that haven't been simulated yet and $p_t$ is the hit probability evaluated as the ratio between the allele fragment size and the size of the sample DNA that hasn't been processed yet. If an allele fragment is shorter than the simulated template, no template is simulated, and the following allele fragment is considered. For each simulated template, the insert size is drawn from a Binomial distribution $B(n_i, p_i)$ whose parameters are computed from the corresponding normal distribution's mean and standard deviation. In particular, if $\mu_i$ and $\sigma_i$ are the mean and the standard deviation provided by the user for the insert size normal distribution, respectively, then $p_i = 1 - \sigma_i^2/\mu_i$ and $n_i = \lfloor \mu_i/p_i \rceil$. The template initial position is uniformly sampled in the interval from 1 to the fragment size minus the template size to ensure that the fragment allele fully contains the template. The final position of the template is univocally determined by the sampled initial position and the template's size. CLONES retrieves the SIDs placed in the considered allele fragment over the template position from the phylogenetic forest and applies them to the reference region from which the template is coming. The SBSs are simulated by simply replacing a base in the sequence. Instead, insertions increase the template size and require removing bases from the template ends. Analogously, deletions shrink the template size and force base feeding from the reference.

##### *A.4.1. Sequencing errors and quality score simulation*

Sequencing errors are simulated at a user-defined rate by adding a single base error whenever a sample from a uniform distribution on $[0, 1]$ does not exceed the rate. Instead, the Phred quality scores are sampled from a set of normal distributions depending on the base positions and whose parameters are fitted from an in-house cohort 21 high coverage whole genome sequencing samples. These samples comprises both normal and tumour, sequenced respectively at 30x and 100x with Illumina Novaseq600. The mean and the variance of the quality scores per base position $x$ were fitted into two quadratic functions, $\mu_q(x)$ and $\sigma^2_q(x)$, that model the relation between the parameters of the normal distribution ruling the quality score and the base position on the read (see Figure S4). Modelling the mean and variance of the quality score using two functions allows CLONES to scale them along the position axis, handling reads of different lengths. When the read size is $l$, ProCESS samples the quality scores of a base in position $x$ from the normal distribution with means $\mu_q\left(x\frac{l}{100}\right)$ and $\sigma^2_q\left(x\frac{l}{100}\right)$.

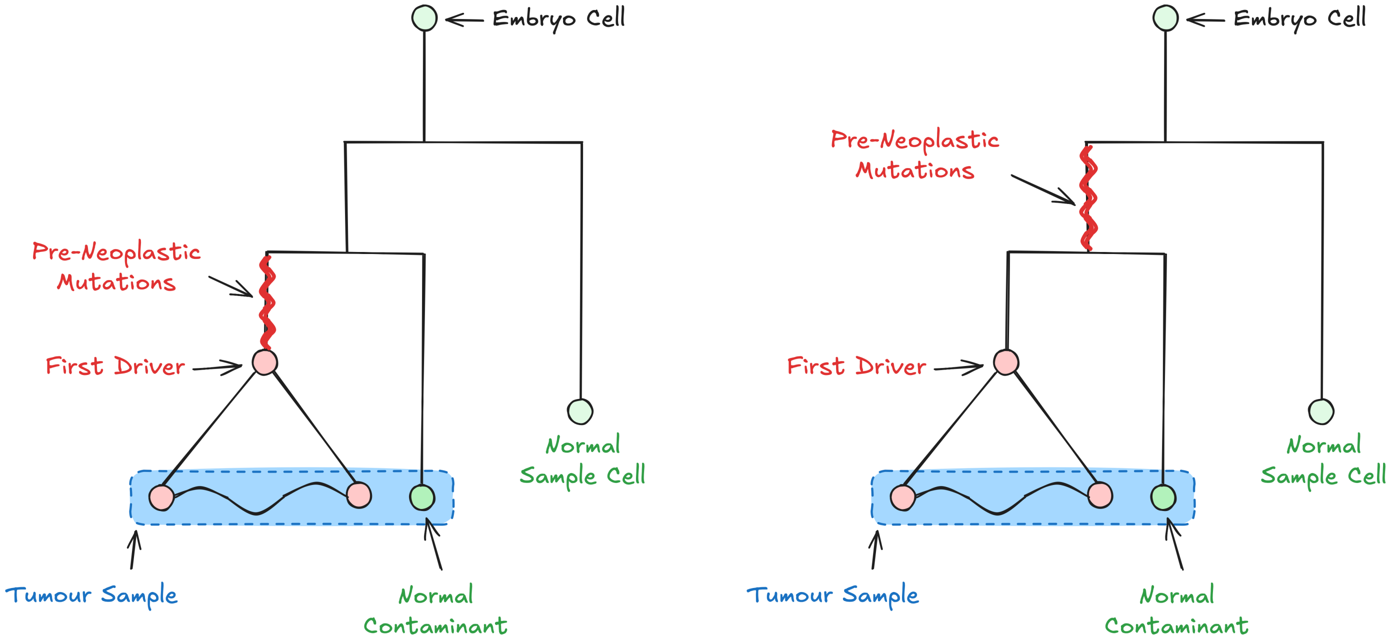

**(a)** The default simulated sequencing genotypes. Pre-neoplastic mutations exclusively appear in the tumour cells. Contaminants and normal cells have the same genotype and only contain germline mutations.

**(b)** The alternative sequencing genotypes supported by ProCESS. Pre-neoplastic mutations appear in tumour and contaminant cells. Instead, normal (non-contaminant) cells exclusively contain the germline mutations.

**Supplementary Fig. S5.** The default simulated sequencing genotypes. The tumour sample contains both tumour cells and normal contaminant cells. The number of contaminant cells depends on the specified sample purity. Tumour cells contain germline mutations, pre-neoplastic mutations, and somatic mutations, which are simulated within the tumour phylogenetic forest.

##### *A.4.2. Normal sample, contaminant, and purity*

The samples are exclusively composed of tumour cells by construction. However, CLONES can alter the sample's purity during simulated sequencing. This feature is achieved by temporarily adding sufficient normal contaminant cells to the sample to achieve the targeted purity. If $p$ is the aimed purity and the considered sample contains $n_s$ tumour cells, then CLONES adds $\left\lceil n_s \frac{1-p}{p} \right\rceil$ normal cells to the sample. All contaminant cells have the same genotype, and, by default, they exclusively contain germinal mutations. Whenever it is required, the default behaviour can be changed, and pre-neoplastic mutations can be inserted into the normal contaminant genome. CLONES can also simulate the sequencing of a normal sample by simulating a single-cell sample whose genome contains only germinal mutations. Although the reads produced by this kind of sample do not contain sufficient variability to effectively simulate wild-type cells, they can serve as the reference for genome sequencing analysis tools such as nf-core/sarek [19]. Figure S5 summarises the default simulated sequencing genotypes and their alternatives.

##### *A.4.3. Sequencing simulation results*

As a result of the simulated sequencing, ProCESS outputs an R object that contains the simulation parameters and a data frame reporting, for each SID in the sampled cells and for each of the samples, the coverage of the corresponding locus, the number of occurrences in the simulated reads, the variant allele frequency (VAF), and the SID class, i.e., driver, passenger, pre-neoplastic, or germinal. When the SID is a passenger mutation, the data frame also reports the mutation signature that caused the mutation; when the SID is a driver mutation, the mutant characterised by the mutation.

| chr | chr_pos | ref | alt | causes | classes | S1.occurrences | S1.coverage | S1.VAF | S2.occurrences | … | S3.occurrences | S3.coverage | S3.VAF |
|---|---|---|---|---|---|---|---|---|---|---|---|---|---|
| 22 | 16050185 | A | C | B | driver | 0 | 36 | 0 | 0 | … | 6 | 11 | 0.545 |
| 22 | 16052080 | G | A | | germinal | 7 | 23 | 0.304 | 22 | … | 6 | 11 | 0.545 |
| 22 | 16078613 | A | C | SBS1 | pre-neoplastic | 15 | 30 | 0.5 | 17 | … | 6 | 12 | 0.5 |
| 22 | 16147672 | GTTT | G | ID1 | pre-neoplastic | 24 | 39 | 0.615 | 17 | … | 2 | 6 | 0.333 |
| 22 | 16147892 | AC | A | | germinal | 12 | 26 | 0.462 | 12 | … | 6 | 10 | 0.6 |
| 22 | 16242273 | CTT | C | | germinal | 11 | 22 | 0.5 | 19 | … | 6 | 10 | 0.6 |
| 22 | 16250626 | A | G | SBS3 | passenger | 2 | 34 | 0.059 | 0 | … | 0 | 10 | 0 |
| 22 | 19276404 | A | AC | ID9 | passenger | 0 | 26 | 0 | 0 | … | 1 | 27 | 0.037 |
| 22 | 21689059 | CAAA | C | ID2 | passenger | 16 | 33 | 0.485 | 0 | … | 3 | 28 | 0.107 |

**Supplementary Table T2.** Some lines of a data frame returned by ProCESS sequencing a phylogenetic forest of the tumour model depicted in Figure S3. The pre-neoplastic and germinal mutations affect all the samples. Instead, most passenger mutations occur exclusively in one sample. However, some of them appear between the root and the MRCA of two samples in the phylogenetic forest and are sequenced in more than one sample, e.g., (22:21689059)[CAAA>C]. The first line in the data frame reports the driver mutation (22:16050185)[A>C] of the mutant B. The mutation does not occur in the reads of samples S1 and S2 because these samples exclusively contain cells of mutants A and C, which are not sub-mutants of B.

ProCESS also offers plotting functions to summarise and compare sequencing results. Moreover, it optionally saves the produced reads in a SAM file[20] version 1.6, reporting their position in the reference genome, their mutations (field CIGAR), and the originating sample (attribute read group). The SAM file can eventually be processed by SAMtools [20] to extract the corresponding FASTQ file or convert it to a BAM or CRAM file [21].

#### A.5. Algorithmic analysis and performances

This section provides insight into the algorithms and data structures implemented at each of the CLONES' levels.

##### *A.5.1. Tissue Level*

The overall time cost of a tissue-level simulation depends on the modelled evolution and cannot be represented by a general formula. In this section, we focus on the cost of a single simulation step, which consists of stochastically selecting an event and applying it.

As mentioned above, CLONES implements a main loop that chooses the next event to simulate, its timing, and the affected mutant using Gillespie's first-reaction method, selects a cell from the affected mutant that is available for the chosen event, then applies the event, updating the cell's availability status in the simulation. The Gillespie's first-reaction method is linear in the number $k$ of the simulated mutants [15]. The cancer cells in the simulation are organised by their availability status and mutants in *indexed RB-trees*. This data structure supports inserting and removing cells, accessing the $i$-th cell in the map, and searching for a cell by its identifier in time logarithmic with respect to the number of cells in the map. Hence, each random selection and availability status update takes logarithmic time with respect to the number of cancer cells in the simulation.

The costs of each event and the number of availability status updates per event depend on both event type and the adopted growth model.

**Cell deaths** In both homogeneous and border-driven growth models, every cell is available for death. Thus, the death of any cell forces CLONES to remove it from those available for duplication, mutation, and death. This task scales logarithmically with the number of cancer cells in the simulation. In the border-driven growth model, the death of a cell $c$ updates the tumour border and may also trigger changes in the availability status for mutation and duplication of cancer cells in the neighbourhood of $c$. However, any cell has at most 8 neighbours in a 2-dimensional grid. Hence, in general, the asymptotic time complexity of a cell death is $O(\log n)$ where $n$ is the number of cancer cells in the simulation.

**Cell duplications and mutations** Both duplicating a cell $c$ and mutating $c$ cause $c$ to die and be removed from the set of available cells for duplication, mutation, and death. This step takes time $O(\log n)$. Then, one of $c$'s children is placed in the former position of $c$, while the position of the other is selected with a probability that depends on the distance of the original cell $c$ from the border according to Section A.2. The distance from the border is evaluated on-the-fly, visiting the tumour mass from the original cell position in all the possible directions $x \in \mathcal{D}$. The asymptotic complexity of this process depends on the tumour shape. In any case, it is upper-bounded by $O(\sum_{x\in\mathcal{D}} d_{b,x}(c))$, where $d_{b,x}(c)$ is the distance of $c$ from the border towards direction $x$. In the worst-case scenario, the tumour evolves in a straight line of cells and $O(\sum_{x\in\mathcal{D}} d_{b,x}(c))$ corresponds to $O(n)$ where $n$ is the number of cancer cells in the simulation. However, this case is unlikely because the tumour mass should expand uniformly across the tissue according to Eq. 12, and, at the steady state, the expected asymptotic complexity to evaluate the border distance from $c$ on a 2-dimensional grid is $O(\sqrt{n})$. Once the border distance has been evaluated, CLONES chooses a position for the second child of the original cell and pushes cancer cells in that direction. This task has the same asymptotic cost as evaluating the border distance: $O(n)$ and expected $O(\sqrt{n})$.

Finally, depending on the selected growth model, the children of the original cell are inserted among those available for mutation, duplication, and death in time $O(\log n)$.

In the homogeneous growth model, every cell is available for duplication, mutation, and death, and no further action is required. Thus, the asymptotic complexity of a duplication or a mutation is $O(\log n + n + \log n) = O(n)$, while its expected complexity is $O(\log n + \sqrt{n} + \log n) = O(\sqrt{n})$.

In the border-driven growth model, only border cells are available for duplications and mutations. A push may alter the availability status of the pushed cells by moving some of them away from the border and others near it. The status of a pushed cell changes only if it was in the neighbourhood of a non-cancer cell before the push, but not after, and vice versa. Thus, the number of changes is upper-bounded by the length $\ell$ of the push. Once again, if the tumour evolves as an elongated mass, a cell push may prompt $O(n)$ changes in availability status for an overall cost of $O(n \log n)$. However, the push direction is randomly selected with a probability proportional to the border distance in that direction. As a consequence, no cells must be pushed with high probability in the border-driven growth model. Moreover, when some cells are pushed, these cells are a contiguous sequence of cancer cells ending in a non-cancer cell. Thus, if we assume a uniform distribution of cancer cells along the region containing the pushed cells and their neighbourhood, then the expected length of such sequences is

$$E[\ell] = \sum_{k=1}^{\infty} k p_c^{k-1}(1 - p_c) = \frac{1}{1 - p_c} \tag{18}$$

where $p_c$ is the potentially unknown probability for each cell in the tumour mass to be a cancer cell. It is easy to see that the neighbourhood of a straight vertical, horizontal, or diagonal segment having length $\ell$ contains at most $2\ell + 6$ cells in a

**Supplementary Table T3.** The strict and expected asymptotic time costs for event selection and simulation for both homogeneous and border-driven growth models. The values $k$ and $n$ are the number of mutants and the number of cells in the simulation, respectively.

| | Homogeneous | Border-driven |
|---|---|---|
| Event selection | $O(k)$ | $O(k)$ |
| Death | $O(\log n)$ | $O(\log n)$ |
| Duplication/Mutation | $O(n)$ (exp. $O(\sqrt{n})$) | $O(n \log n)$ (exp. $O(\sqrt{n})$) |

2-dimensional grid. Thus, the expected number of non-cancer cells in the neighbourhood of such a segment is $(2\ell + 6)(1 - p_c)$, and the number of expected non-cancer cells in the neighbourhood of a push $X_b$ is

$$E[X_b] = (2E[\ell] + 6)(1 - p_c) = \left(2\frac{1}{1 - p_c} + 6\right)(1 - p_c) = 2 + 6(1 - p_c). \tag{19}$$

Since $E[X_b]$ is upper bounded by 8, the expected number of availability status changes due to a single mutation or duplication is constant in the border-driven growth model. Hence, the asymptotic complexity of the same event is $O(\log n + n \log n + \log n) = O(n \log n)$, while its expected complexity is $O(\log n + \sqrt{n} + \log n) = O(\sqrt{n})$.

Supplementary Table T3 summarises the asymptotic complexity for selecting a single event and simulating it.

**Simulation expected complexity** This section provides a rough estimate of the expected asymptotic time complexity $E$ of simulating the expansion of a single-mutant tumour from a single cell to $N$ cells. Each duplication increases the number of cells in the simulation by one. Analogously, a death removes one cell from the simulation. Hence, if the number of deaths during a simulation is $d$, then the number of events required to reach $N$ cells from 1 cell is $N - 1 + 2d$, and $N - 1 + d$ of them are duplications. Let us focus on simulations that end when the number of simulated cells reaches $N$. The number of their cells is always in the interval $[1, N]$. According to Supplementary Table T3, the time costs of a single duplication and a single death in such simulations are upper-bounded by $O(\sqrt{N})$ and $O(\log N)$, respectively. Hence, the cost of each of these simulations is $O((N - 1 + d)\sqrt{N} + d \log N)$. By using the reflection principle, we can prove that the number of the considered simulations is

$$W_{N,d} = \sum_{k=-\infty}^{\infty} \left( \binom{N + 2d - 1}{d - k(N+1)} - \binom{N + 2d - 1}{d - k(N+1) - 1} \right). \tag{20}$$

In the homogeneous growth model, all cells are active for both duplication and death. Thus, the probabilities of these two events do not depend on the number of cells in the simulation, and $p_\lambda = \lambda/(\lambda + \delta)$ and $p_\delta = \delta/(\lambda + \delta)$ throughout the simulation. As a consequence, the expected complexity $E$ of simulating the expansion of a single-mutant tumour from a single cell to $N$ cells is

$$E = O(\sum_{d=0}^{\infty} \underbrace{W_{N,d}}_{\substack{\text{Num. of sim.}\\ \text{with } d \text{ deaths}}} \underbrace{p_\lambda^{N-1+d} p_\delta^d}_{\substack{\text{Prob. of sim.}\\ \text{with } d \text{ deaths}}} \underbrace{\left((N - 1 + d)\sqrt{N} + d \log N\right)}_{\substack{\text{Cost of sim.}\\ \text{with } d \text{ deaths}}}) = O\left(p_\lambda N^{3/2} + p_\delta N \log N\right). \tag{21}$$

When $\lambda > \delta$, the expected complexity is $O(N^{3/2})$.

Instead, the border-driven growth model allows only cell duplications at the tumour border, and the number of border cells depends on the tumour shape. In this case, we cannot express the complexity of reaching a mass of $N$ cells in closed form, and we may ask under which conditions the tumour can reach a mass of $N$ cells.

**Sampling** Sampling removes sample cells from the simulations and saves their identifiers to a file for later construction of the sample forest. Cell removal is handled as a cell-death operation, requiring asymptotic time $O(\log n)$ and $O(1)$ disk accesses per sampled cell. Moreover, each sample triggers a snapshot of the simulation to disk, requiring $O(n)$ disk writes. Hence, if $n_s$ is the number of sampled cells, the overall cost of collecting one sample is $O(n_s \log n + n)$ with $O(n_s + n)$ disk writes. When $n_s$ is smaller than $n/\log n$, the dominant term is $O(n)$.

**Building sample forests** Building the sample forest requires retrieving the ancestors of the sampled cells. This is done by using a binary log file containing the simulated cell's data. For each simulated cell, the log file records the cell identifier, the parent cell identifier, the cell's mutant, and the birth time. Entries are saved to the log file when the corresponding cells are created, and each cell requires the same amount of space. Moreover, cell identifiers are progressive. Thus, retrieving a cell's data using its identifier as the index takes a constant number of disk accesses.

Each node in the sample forest corresponds to a cell in the simulation and stores the cell's data, as maintained in the log file, along with the identifiers of the cell's children. The forest maintains an RB-Trees that associates the identifiers of the cells represented in the forest with the corresponding nodes. At the beginning of the construction, the map is empty, and the identifiers of the sampled cells are processed one at a time. The data corresponding to the considered identifier are retrieved from the log file by using a constant number of disk reads, and the cell node is created and added to the map in time $O(\log q_t)$ where $q_t$ is the number of cells in the map at the time $t$. If the considered identifier is that of one of the cells placed on the tissue at the beginning of the simulation, then the node is added among the forest roots, and the next sampled cell is considered. Otherwise, the algorithm checks whether the node corresponding to the parent cell is already in the tree in time $O(\log q_t)$. If this is the case, the list of the children in the parent node is updated in time $O(\log q_t)$ and the next sampled cell is considered. If, instead, the parent cell's identifier is not associated with any node, the cell's data is retrieved from the log file in time $O(1)$, and the corresponding node is constructed and added to the map in time $O(\log q_t)$. Then, the algorithm recursively traverses one of the forest roots, considering the parent's identifier of the new node. The data for each cell in the forest is retrieved from the log file at most once during the build process. Since every cell in the simulation has two children at most, every node in the forest is visited at most twice. Thus, building the sample forest is $O(q \log q)$, where $q$ is the number of nodes in the final forest, and requires $O(q)$ disk reads in total.

The number of nodes in the sample forest $q$ cannot be determined by the number of sampled cells $n_s$ alone. We know that $n_s \leq q$ because all sampled cells are represented as leaves in the sample forest, and the number of nodes at each level must be less than or equal to the number of leaves in the forest. However, the forest height and the average node height, together with $q$, depend on many factors. For example, the greater the simulated time at which the sample is collected, the more ancient the lineage of the sample cells, and the taller the sample forest. Analogously, the greater the ratio between the duplication and death rates of the simulated mutants, the more level the forest, and the larger the set of forest nodes. The growth model and the position of the collected sample also affect the number of nodes in the forest. In the border-driven growth model, only cells in the tumour border can duplicate. Hence, the cells at the tumour border tend to have more ancestors than those at the centre of the tumour mass.

##### A.5.2. Mutation Level

**Genome representation operations** Behind lazy-copy, which takes constant time, CLONES implements deep-copy, which fully duplicates the genome data structure with no reference to the original object in time $\Theta(n_f + M_G)$ where $n_f$ and $M_G$ are the number of allele fragments and SIDs in the genome, respectively. Moreover, it supports three operations for inserting and removing mutations into and from the data structure representing the cell genome: `insert_CNA`, `insert_SID`, and `remove_SID`. `insert_CNA` inserts a CNA, being either an amplification or a deletion, in the genome. Before the insertion, `insert_CNA` triggers a full duplication of the objects that will be changed by the insertion and that are referenced more than once. The complexity of `insert_CNA` depends on the kind of CNA: an amplification requires time $\Theta(n_a + n_{f,a} + M_a)$, where $n_a$ is the number of alleles in the affected chromosome, and $n_{f,a}$ and $M_a$ are the number of fragments and mutations in the affected allele, respectively; a region deletion requires time $\Theta(n_{f,a} + M_a)$. Instead, `insert_SID` inserts either an SBS or an ID in the genome, avoiding data structure duplications. This operation alters all objects referencing the allele fragment that contains the mutation, and its cost is $\Theta(\log M_f)$ where $M_f$ is the number of mutations in the affected fragment. Finally, `remove_SID` removes a mutation from a data structure, again, avoiding data structure duplications. Also in this case, the operation affects all objects referencing the allele fragment from which the mutation is removed, and the execution time is $\Theta(\log m_f)$.

The asymmetry in the sematic of `insert_CNA`, which produces a new genome referencing to the unchanged part of the original structure, and those of `insert_SID`, and `remove_SID`, which alter a genome and all those referencing to it, is due to efficiency pourposes and plays a crucial role in the asymptotic complexity mutation placing.

**Building phylogenetic forests** Phylogenetic forests are built from the corresponding sample forests by labelling each node with the mutations of the cell represented by the node. This task is implemented by a recursive procedure that visits the sample forests depth-first and enriches the node's genome with mutations arising in that node. Depending on whether the node is a forest's root or has a parent, the per-enrichment genome is either the wild-type genome with the pre-neoplastic mutations or the genome inherited from the node's parent.

The recursive procedure takes as input a node in the sample forest representing a cell $c$ and either the genome of $c$'s parent, when $c$ has a parent, or the wild-type genome added with the pre-neoplastic mutations, when $c$ is one of the cells placed on the simulated tissue. As the first step, the procedure makes a lazy copy of the second parameter to represent the $c$'s genome in time $\Theta(1)$. Then, the cell's mutant and the corresponding mutation rates and drivers are retrieved from two maps in time $O(\log(q) + \log(k))$, where $q$ is the number of nodes in the forest, and $k$ is the number of mutants in the simulation. Having the mutation rates for indels, SBSs, and CNA, the procedure samples the number of mutations to add for each of the three kinds and inserts them in $c$'s genome using `insert_CNA` and `insert_SID`. After that, if the node is a leaf, the procedure saves the genome in the list of the sampled cell genomes by performing a deep copy in time $O(n_{f,c} + M_c)$, where $n_{f,c}$ and $M_c$ are the number of allele fragments and somatic mutations in $c$'s genome, respectively. If the node has any children, the procedure handles them recursively. Finally, before returning the control to the calling function, the SIDs that were added to $c$'s genome are removed from the genome of $c$'s parent using `remove_SID`.

The number of mutations arising at cell birth depends on the amount of DNA in the genome, i.e., the number and length of the alleles. These quantities are affected by CNAs. When an amplification occurs, the amount of DNA increases by the length of the amplified fragment. Symmetrically, when a deletion takes place, the amount of DNA decreases by the length of the CNA. If the number of CNAs and their lengths are low, the change in DNA quantity is negligible, and we can ignore it. However, if the rate of the CNAs is high, the global effect on the genome cannot be predicted, and the asymptotic complexity cannot be evaluated by a closed formula.

For simplicity, we will focus on the asymptotic analysis of building a phylogenetic forest for samples coming from a single-mutant tumour without CNAs. The number of SIDs to place in a node is sampled at each cell birth by a Poisson distribution whose mean is $\mu_m = (\text{SBS} + \text{ID})D$, where SBS, ID, and $D$ are the SBS rate, the ID rate, and the amount of DNA in the cell's genome, respectively. Thus, the expected number of SIDs arising in each node is $\mu_m$. If there are $m_p$ pre-neoplastic SIDs, then each cell $c$ in the $j$-th level of the phylogenetic forest is expected to both inherit $j\mu_m + m_p$ somatic SIDs from its parent and be affected by $\mu_m$ new SIDS. Thus, the total number of somatic mutations on a sampled cell $c$ whose height in the forest is $h_c$ is expected to be $h_c\mu_m + m_p$. As far as time complexity may be concerned, each call to the recursive procedure retrieves the mutation rates in time $O(\log(q))$ and is expected to add $\mu_m$ new mutations to the genome by using $\mu_m$ successive `insert_SID` in time $\Theta(\sum_{i=1}^{\mu_m} \log(i + j\mu_m + m_p)) = \Theta(\mu_m \log((j+1)\mu_m + m_p))$. The functions `insert_SID` and `remove_SID` have the same complexity, and the cumulative time cost of adding and, subsequently, removing $\mu_m$ SIDs during a single recursive call is the same. It follows that, if no CNA occurs, a single call of the recursive procedure on any internal node of the $j$-th level is expected to require asymptotic time $O(\log(q) + \mu_m \log((j+1)\mu_m + m_p))$. Each recursive call to a forest leaf also requires a deep copy of the corresponding sampled cell genome. The expected overall cost of deep copying all sampled cell genomes is $\Theta(\sum_{c \in C_s}(h_c\mu_m + m_p)) = \Theta(n_s(\overline{h_s}\mu_m + m_p))$ where $C_s$ is the set of all the sampled cells and $\overline{h_s}$ is the average depth of the leaves in the forest. Thus, if no CNA occurred, the expected asymptotic time cost of building a phylogenetic forest whose $j$-th level contains $q(j)$ nodes is

$$\begin{aligned} &O\left(n_s(\overline{h_s}\mu_m + m_p) + \sum_{j=0}^{h} \left(q(j)\left(\log(q) + \mu_m \log\left((j+1)\mu_m + m_p\right)\right)\right)\right) \\ &\subseteq O\left(n_s(\overline{h_s}\mu_m + m_p) + q\log(q) + q\mu_m \log\left((h+1)\mu_m + m_p\right)\right). \end{aligned} \tag{22}$$

##### *A.5.3. Sequencing Level*

The sequencing task takes as parameters a phylogenetic forest, the aimed coverage $C$, the read length $\ell_r$, and both the mean and the standard deviation of the template length $\mu_{\ell_t}$ and $\sigma_{\ell_t}$, respectively. ProCESS generates $n_r = CG/\ell_r$ reads, where $G$ is the wild-type genome size. In the case of paired reads, $n_t = \lceil n_r/2 \rceil$ templates are needed because each sequencing template contains two paired reads. These templates come from the genomes of all the sampled cells. For each sampled cell genome and for each allele fragment $f$ in the genome, the number, $n_{t,f}$, of templates coming from $f$ is sampled over a binomial distribution $B(n'_t, \ell_f/D')$ where $\ell_f$, $n'_t$, and $D'$ are the length of the allele fragment, the number of templates that have not been produced, and the amount of DNA that has not been considerated yet. The cumulative asymptotic time to sample the number of templates to be extracted from each allele fragment is $\Theta(n_f)$ where $n_f$ is the number of allele fragments of all sampled cell genomes.

For each allele fragment $f$ and for each of the $n_{t,f}$ templates to be extracted from $f$, the length $\ell_t$ of the template and the initial position $b_t$ of the template in $f$ are sampled in time $O(1)$ from the normal distribution $\mathcal{N}(\mu_{\ell_t}, \sigma_{\ell_t})$ and the discrere uniform distribution over the interval $[1, \ell_t - \ell_t + 1]$, respectively. Then, the nearest somatic SID to $b_t$ that comes before it in $f$ is found in time $O(\log M_f)$ where $M_f$ is the number of somatic mutations in $f$. The allele to which $f$ belongs may be absent from the wild-type genome. However, CLONES tracks allele origins and maps any allele $a$ to the allele, $\text{wt}(a)$, which is $a$'s original allele in the wild-type genome in time $O(1)$. This helps in identifying the nearest germinal SID to $b_t$ that comes before it in $a$ in time $O(\log \overline{M_f})$ where $\overline{M_f}$ is the number of geminal mutations in $\text{wt}(a)$. Afterwards, the read sequences are built from 5' to 3' using the reference genome and the germinal and somatic mutations lying on it. Since the reads have length $\ell_r$, the number of somatic and germinal mutations on it is at most $\ell_r$. Scanning $\ell_r$ consecutive values from a map containing $M_f$ and $\overline{M_f}$ takes time $O(\ell_r + \log M_f)$ and $O(\ell_t + \log \overline{M_f})$, respectively, and building the read sequence has asymptotic time complexity $O(\ell_r + \log M_f + \log \overline{M_f})$. Then, CLONES simulates sequencing errors in time $\Theta(\ell_r)$. Hence, producing paired reads cumulatively takes asymptotic time $O(\ell_r + \log(M_f\overline{M_f}))$. In a coarse, but conservative estimation, generating the reads for all the sampled cells takes expected asymptotic time $O(n_s + n_f + n_t(\ell_r + \log(M_{\max,f}\overline{M_{\max,f}})))$ where $n_s$, $n_f$, $n_t$, $M_{\max,f}$, and $\overline{M_{\max,f}}$ are the number of sampled cells, the number of allele fragments in the sampled cells, the number of template to be produced, the maximum number of somatic mutations in a fragment, and the maximum number of germinal mutations in a fragment.

Sequencing simulation returns mutation frequencies in the simulated reads. To achieve this goal, CLONES uses a map that associates each mutation in the sampled cells with the number of occurrences of that mutation in reads generated for each sample. Initially, the number of occurrences for each mutation is set to 0 in time $O(M_{\text{D}} \log M_{\text{D}})$ where $M_{\text{D}}$ is the total number of distinct mutations in the sampled cells. Then, every time a simulated read containing a mutation is generated, the number of occurrences of that mutation is updated in time $O(\log M_{\text{D}})$. In total, the cost of tracking the mutation frequencies is $O((M_{\text{D}} + M_{\text{S}}) \log M_{\text{D}})$ where $M_{\text{S}}$ is the number of mutations in the simulated reads. The reader must be aware that $M_{\text{S}}$ is proportional to the coverage $C$ and it is not affected by the number of sampled cells, whereas $M_{\text{D}}$ is directly affected by the number of sampled cells and it is unrelated to the coverage.

Since $n_t \in \Theta(CG/\ell_r)$, the overall asymptotic cost of sequencing simulation is:

$$O\left(n_s + n_f + CG + \frac{CG}{\ell_r}\left(\log\left(M_{\max,f}\overline{M_{\max,f}}\right)\right) + (M_{\text{D}} + M_{\text{S}})\log M_{\text{D}}\right). \tag{23}$$

#### A.6. Performances

This section describes the tests used to validate the asymptotic analysis and measure the performance.

##### *A.6.1. Materials and methods*

Each test was repeated 5 times with different random seeds to account for stochastic variability. The test results were collected on a MacBook Pro M1 13" (2020) with 16GB of RAM, with macOS Tahoe 26.3.1, ProCESS version 1.1.5, and R version 4.5.2.

**Number of mutants** We considered a $7000 \times 7000$-cell simulated tissue, and we simulated tumour expansion from 240 cells to 1 million cells, with the initial cells belonging to 1, 10, 30, 60, 120, and 240 mutants, all with the same duplication and death rates (0.4 and 0.01, respectively). Each simulation used the border-driven growth model. The initial cells were placed along the circumference of a circle centred on the tissue's midpoint, with a radius of 1750. In particular, the $j$-th cell was placed in position $(3500 + 1750\cos(2\pi j/240), 3500 + 1750\sin(2\pi j/240))$ and assigned to the $(j \mod k)$-th mutant, where $k$ is the number of mutants in the simulation. Each tumour cell was equally distant from its neighbours, and the mutants were uniformly distributed on the circumference.

**Tumour size and duplication rates** We considered a $7000 \times 7000$-cell simulated tissue, and we simulated the expansion of a single-mutant tumour from a single cell placed at the centre of the simulated tissue using both the border-driven and the homogeneous growth models, and we measured the time required to grow the mass to $4 \times 10^5$, $8 \times 10^5$, $1.2 \times 10^6$, $1.6 \times 10^5$, and $2 \times 10^6$ simulated cells. The mutant's death rate was 0.01 in all the tests. Instead, the duplication rate was set to one of 0.1, 0.2, or 0.4, and, as with the death rate, it remained constant throughout each test.

In this set of tests, we used the *Corrected Akaike Information Criterion* (AICc) [22, 23] to identify the most likely models relating the number of cells to computation time. We held the duplication rate and growth models fixed, and we evaluated the AICcs of the $1/N$, $N$, $N\log N$, $N^{3/2}$, $N^2$, and $e^N$ models. The model with the smallest AICc is the one most likely to properly represent the relation.

**Sampling** We considered 2-million-cell single-mutant tumours initiated from a single cell placed at the centre of a $7000\times7000$-cell simulated tissue. The mass was grown using both border-driven and homogeneous growth models. The mutant's duplication and death rates were set to $\lambda \in \{0.1, 0.2, 0.3, 0.4, 0.5\}$ and $\delta \in \{0.01, \lambda/10\}$, respectively. For each setup, we collected $n_S \in [1, 10]$ samples, either at the edge or at the centre of the tumour. A sample consisted of a $71 \times 71$-cell tissue square containing at least 80% of tumour cells. When the samples were collected at the centre of the tumour, they formed a spiral. The first sample is centred at the tissue centre. The next eight samples are taken from the neighbourhood of the first sample, while the last sample is the farthest from the tumour centre. For each test, we measured the sampling time.

AICcs of the $1/n_s$, $n_s$, $n_s\log n_s$, $n_s^{3/2}$, $n_s^2$, and $e^{n_s}$ models were used to identify the model that better represents the relation between the number of samples and the sampling time when the growth model and position of the samples are fixed. The lowest AICc corresponds to the model that most likely fits the data.

**Building sample forests** We built the sample forests corresponding to the sampling tests and measured the number of sampled cells, forest heights, the number of nodes, and the time required to build them. To reduce the relative error, the latter quantity was evaluated as the average time required to build each sample forest 40 times.

We investigated the relationships between the number of collected samples and the height of the corresponding forests and between the number of sampled cells and the number of nodes in the corresponding forests, as the growth model and sampling position changed. We also studied how the ratio of the duplication rate over the death rate affects the number of nodes in the forests. Finally, we related the number $q$ of nodes in the forests to the time required to build them, and we used the AICcs of the $1/q$, $q$, $q\log q$, $q^{3/2}$, $q^2$, and $e^q$ models to identify, among them, the most suited ong to represent this relation.

**Building mutation engine** For this set of tests and for the following ones, we used the ProCESS setup `GRCh38`, which includes the homonimum human reference genome.

The phylogenetic forest generation requires the mutation engine to place mutations. The first initialisation of this component downloads the required files – e.g., the reference genome, mutational signatures, and germinal mutations – from the internet and uncompresses them. Then, two indices associating the SBS and ID types to their possible loci are built and saved to disk. Any successive mutation engine setup loads the two indexes and requires less time. To reduce memory and disk requirements, the two indices are presampled. The SBS index stores one locus per COSMIC SBS mutation type every `context_sampling`; the ID index maintains at most `max_repetition_storage` loci per COSMIC ID mutation type [16].

The download time is affected by network speed. Thus, we pre-downloaded and compressed all the required files to avoid measurement bias. We generated the engine varying `context_sampling` and `max_repetition_storage` in $\{20, 40, 60, 80, 100\}$ and $\{4\times10^5, 8\times10^5, 1.2\times10^6, 1.6\times10^6, 2\times10^6\}$, respectively. During each test, we measured the cumulative time spent building and saving the mutation engine. Once the indices were built, we repeated the mutation engine construction using the same `context_sampling` and `max_repetition_storage` values to also measure the mutation engine's loading times.

**Building phylogenetic forests** For each test in this section, we measured only the time required to build the phylogenetic forests and excluded the time to generate or load the mutation engine. We considered the sampled forests generated by the previous set of tests from single-mutant tumours with duplication and death rates $\lambda = 0.4$ and $\delta = 0.01$, respectively. The mutation engine was always built by using the ProCESS setup `GRCh38` and the ID and SBS mutational exposures were set to be $\{ID2 = 0.6, ID13 = 0.2, ID21 = 0.2\}$ and $\{SBS13 = 0.2, SBS1 = 0.8\}$, respectively.

In a first set of tests, we took as input the sample forests built after collecting 10 samples, and we generated the phylogenetic forests imposing 2000 pre-neoplastic mutations and setting the mutation rate to $5 \times 10^{-10}$, $7.5 \times 10^{-10}$, $1 \times 10^{-9}$, $1.25 \times 10^{-9}$, and $1.5 \times 10^{-9}$ for both indels and SBSs. Then, we fixed the mutation rate to be $1 \times 10^{-9}$, and the number of pre-neoplastic mutations was evenly distributed between indels and SBSs and set to 6000, 8000, 10 000, 12 000, and 14 000. In a third set of tests, the mutation rate was $1 \times 10^{-9}$, no preneoplastic mutations were added, and the only mutant had 0 to 10 amplifications of chromosome 1 as driver mutations. Since chromosome 1 is about one-tenth of the wild-type human genome and almost every chromosome has 2 alleles, every amplification of this chromosome increases the amount of DNA by about one-twentieth with respect to the wild-type, and 10 amplifications increased it by 50%. While these scenarios are biologically unlikely, they allowed us to systematically analyse the effect of CNAs on the amount of DNA in a cell and, consequently, on the average number of somatic mutations per generation. Finally, we evaluated the impact of the number of sampled cells on phylogenetic forest generation time by setting the mutation rate, the number of pre-neoplastic mutations, and the number of CNAs to be $1 \times 10^{-9}$, 2000, and 0, respectively, and by considering the sample forests built by collecting from 1 to 10 $70 \times 70$-cell samples. In this last set of tests, the building time depends not only on the number of sampled cells but also on the total number of somatic mutations, which increases with the number of sampled cells.

**Sequencing** We considered the phylogenetic forests built during the previous tests when 10 samples were collected, and 2000 pre-neoplastic mutations were placed. From these forests, we extracted the subforests associated with the last sample, and we simulated its sequencing.

As the first set of tests, we dealt with the forests built using mutation rates in $\{5 \times 10^{-10}, 7.5 \times 10^{-10}, 1 \times 10^{-9}, 1.25 \times 10^{-9}, 1.5 \times 10^{-9}\}$ for both SBSs and indels, and we simulated 0.1-coverage sequecing. Then, we focused on forests generated with a mutation rate of $1 \times 10^{-9}$ and simulated sequencing at coverages of 0.2, 0.4, 0.6, 0.8, and 1.0. The third set of tests used the forests from the second set of tests and measured the time required to simulate 0.1-coverage sequencing with read sizes of 50, 100, 150, 200, and 250.

Finally, we considered the sample forests generated when 10 samples were collected, and we built the corresponding phylogenetic forests with no pre-neoplastic mutations, assuming the only mutant in the simulation has 0, 200, 400, 600, 800 and 1000 driver CNAs amplifying $1 \times 10^4$ base pairs in chromosome 1. Each amplification increases the number of alleles and fragments per cell by 1, and the amount of DNA by about $1/6000\%$ of the baseline. While the amount of DNA in a cell does not appear to be significantly altered by 1000 amplifications, the change may increase the number of mutations in the sampled cells enough to affect the sequencing simulation time. Thus, we set SBS and ID based on the number of CNAs to maintain the number of somatic mutations per test. In particular, both SBS and ID were set to $1 \times 10^{-9} * D/(D + iA)$ where $D$ is the amount of DNA in the wild-type cell, $i$ is the number of driver CNAs, and $A = 1 \times 10^4$ is the size of the amplification. From the resulting phylogenetic forests, we again extracted the subforests associated with the last sample and simulated 0.1-coverage sequencing.

##### *A.6.2. Results and Discussion*

**Number of mutants** The initial setup maximised the likelihood of an equivalent expansion across all mutants and minimised the probability that the populations of some mutants would disappear. Since all mutants had the same duplication and death rates, the expected number of events to complete the simulation is fixed, and the difference in the simulation time is exclusively related to the number of mutants. Supplementary Fig. S6a shows the test results and supports the asymptotic analysis, suggesting that the computation time is linear in the number of mutants.

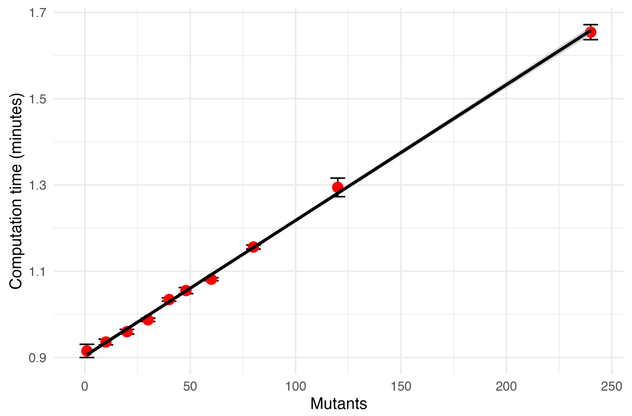

**(a)** Selecting an event to be simulated required a time that is linear in the number of mutants in the simulation. So, it is the overall computation.

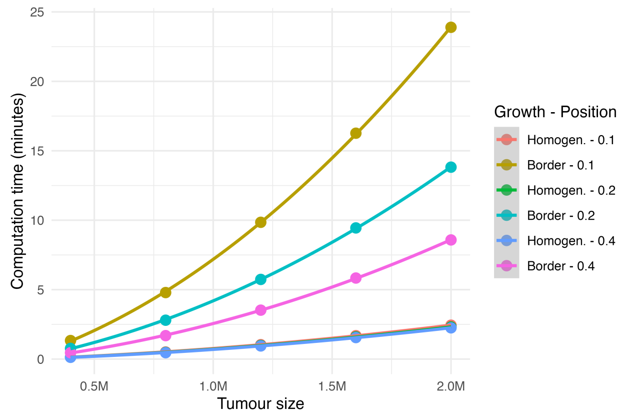

**(b)** Computation time to grow a single-mutant tumour from a cancer cell as the duplication rate and the growth model change.

**Supplementary Fig. S6.** Single mutant tissue-level simulation computation time. This figure represents the computation time required to grow a single mutant mass in both the border-driver and homogeneous growth models. The duplication rate is one among $\{0.1, 0.2, 0.4\}$, while the death rate is set to 0.1. Each measurement is repeated 5 times to account for stochastic variability.

**Tumour size and duplication rates** Supplementary Fig. S6b summarises the results and shows the computation time difference between the border-driven and the homogeneous growth models as the rate varies.

The $N^{3/2}$ model has the smallest AICc among $1/N$, $N$, $N \log N$, $N^{3/2}$, $N^2$, and $e^N$ models for any of the considered rates and for both the border-driven and homogeneous growth models (see Table T4). This result is consistent with the expected asymptotic complexity for the homogeneous growth model reported in Section A.5.1.

**Supplementary Table T4.** The AICc of the $1/N$, $N$, $N \log N$, $N^{3/2}$, $N^2$, and $e^N$ models representing the relation between the aimed number of tumour cells in a simulated tumour and the computation time when the death rate is 0.01. The lowest value corresponds to the most likely model that fits the data. Each row reports the AICs for a particular combination of growth model and duplication rate. The minimum AICcs in each row are underlined. The values whose difference from the row minimum is less than 1% are reported in bold font. The $N^{3/2}$ model has the smallest AICc for both the homogeneous and the border-driven growth models.

| Growth model | Dup. rate | $1/N$ | $N$ | $N \log N$ | $N^{3/2}$ | $N^2$ | $e^N$ |
|---|---|---|---|---|---|---|---|
| Border-driven | 0.1 | 357.7 | 290.57 | 285.32 | **230.46** | 241.17 | 292.91 |
| Homogeneous | 0.1 | 242.56 | 171.6 | 165.75 | **94.89** | 135.6 | 180.55 |
| Border-driven | 0.2 | 330.09 | 262.47 | 257.21 | **205.49** | 220.6 | 266.89 |
| Homogeneous | 0.2 | 240.07 | 169.53 | 163.74 | **94.64** | 131.96 | 178.03 |
| Border-driven | 0.4 | 306.71 | 239.5 | 234.23 | **178.19** | 189.18 | 242.44 |
| Homogeneous | 0.4 | 238.9 | 168.72 | 162.98 | **95.64** | 129.92 | 176.78 |

**Sampling** Considering the first batch of tests, in all four combinations of growth model and sampling position, the AICc of the linear model relating the sampling time to $n_s$ is smaller, than those of $1/n_s$, $n_s \log n_s$, $n_s^{3/2}$, $n_s^2$, and exponential model by more than 1% (Supplementary Table T5). The dominant term in the time cost appears to be disk writing, as conjectured in Section A.5.1.

**Supplementary Table T5.** The AICc of the $1/n_s$, $n_s$, $n_s \log n_s$, $n_s^{3/2}$, $n_s^2$, and $2^{n_s}$ models representing the relation between the number of sampled cells and the sampling time when the duplication and death rates are 0.4 and 0.01, respectively. The lowest value corresponds to the most likely model that fits the data. Each row reports the AICs for a particular combination of growth model and sampling position. The minimum AICcs in each row are underlined. The values whose difference from the row minimum is less than 1% are reported in bold font.

| Growth model | Sampling Pos. | $1/n_s$ | $n_s$ | $n_s \log n_s$ | $n_s^{3/2}$ | $n_s^2$ | $e^{n_s}$ |
|---|---|---|---|---|---|---|---|
| Border-driven | Edge | 328.44 | **59.08** | 70.88 | 173.73 | 231.13 | 228.86 |
| Homogeneous | Edge | 336.33 | **124.4** | 125.78 | 187.03 | 238.98 | 237.76 |
| Border-driven | Center | 246.98 | **-21.25** | -3.26 | 97.53 | 152.93 | 149.57 |
| Homogeneous | Center | 248.69 | **75.86** | 79.12 | 120.39 | 162.16 | 159.19 |

Figure S7 shows how the sampling time scales with the number of sampled cells.

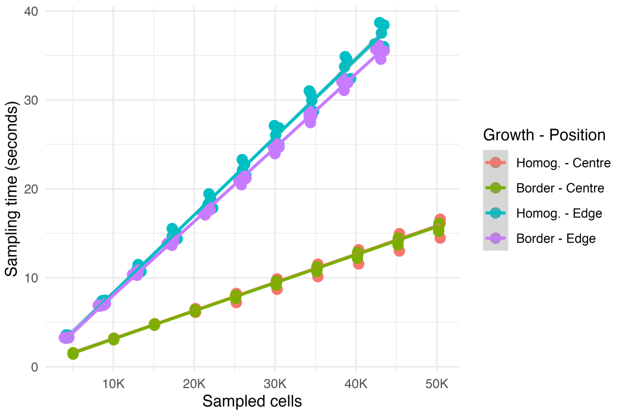

**Supplementary Fig. S7.** Single-mutant sampling time scales linearly in the number of sampled cells. This figure shows the sampling time for a $2 \times 10^6$-cell tumour with duplication and death rates of 0.4 and 0.01, respectively. Each measurement is repeated 5 times to account for stochastic variability.

**Building sample forests** Supplementary Fig. S8 shows the relation between the growth model, the sampling position, and the height of the sample forest. In the border-driven growth model, cells at the tumour border tend to be younger than those at the centre, and the expected lifespan of a cell is inversely proportional to its distance from the tumour centre. The more distant a cell is from the centre, the more ancestors it has. The tumour genealogy trees are strongly unbalanced, with the shortest branches going to cells at the tumour centre and the longest branches ending in cells on the external tumour border. Thus, when the border-driven growth model is adopted, the sample forest height tends to be proportional to the distance between the sample and the tumour centre. The farther apart the samples from the tumour centre, the taller the forest. When the samples are collected at the centre of the tumour, the difference in height between forests built solely using the nearest sample to the tumour centre and the other forests is clearly visible in the figure. By contrast, the heights of the forests built using from 2 to 9 samples are not significantly different because the distances from the tumour centre of the samples between the second and the ninth do not change (Supplementary Fig. S8A "*Border - Centre*"). In any case, the forests built over samples on the tumour edge are taller than those on the other samples (Supplementary Fig. S8A "*Border - Edge*"). Instead, when the homogeneous growth model is adopted, the youngest cells are distributed throughout tumours, and the tumour genealogy trees are balanced. Consequently, at each time instant, the forests in the homogeneous growth model tend to have the same heights throughout a mutant mass. Moreover, forests generated by the border-driven growth model are generally taller than those of the homogeneous growth model (Supplementary Fig. S8A "*Homog. - Centre*" and "*Homog. - Edge*").

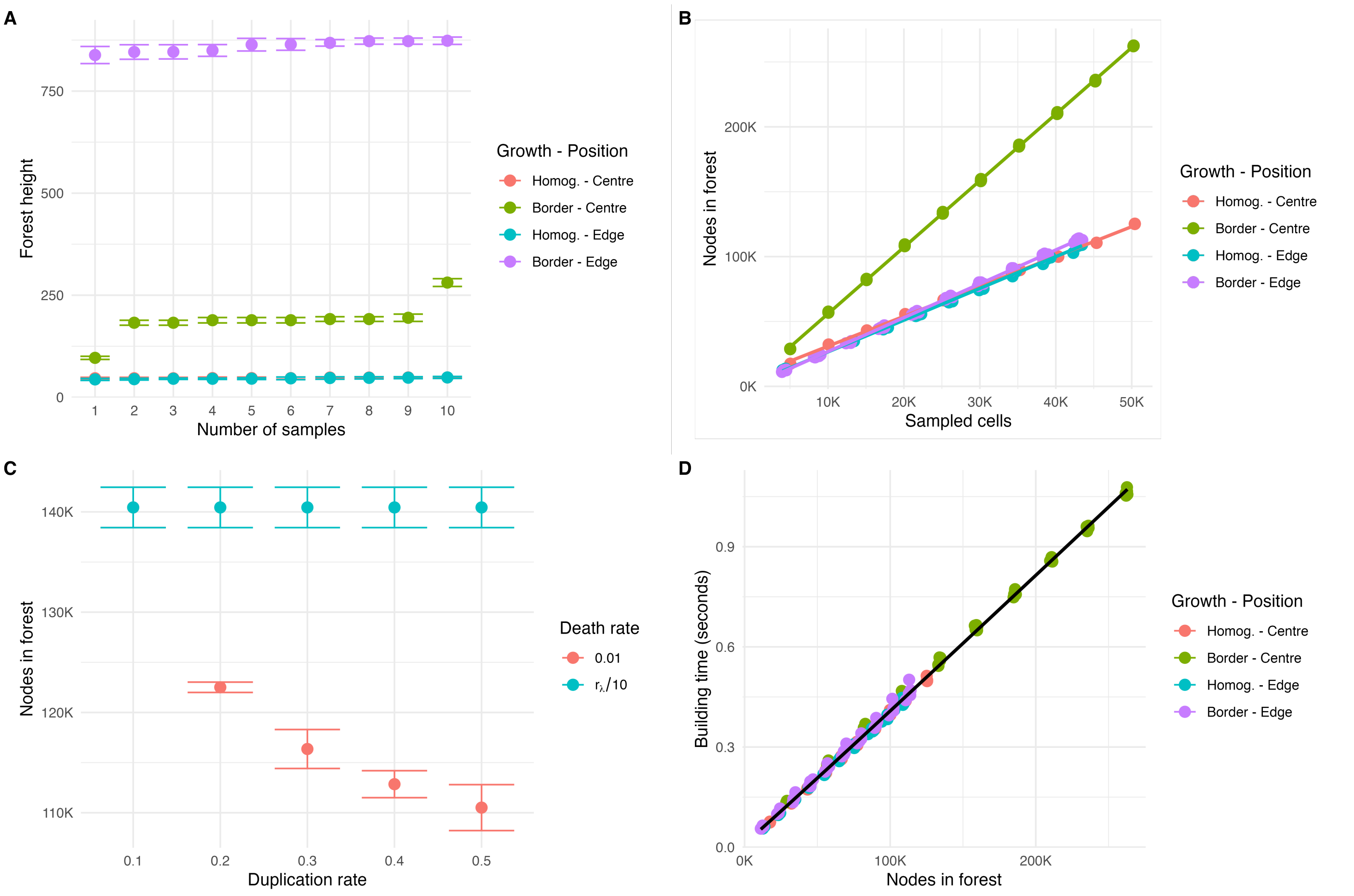

**Supplementary Fig. S8.** Single mutant tissue-level simulation computation time. This figure represents the computation time required to grow a single mutant mass in both the border-driver and homogeneous growth models. Each measurement is repeated 5 times to account for stochastic variability. **(A)** Sample position, growth model, and forest height. When the homogeneous growth model is adopted, the position of the samples does not affect forest height. Instead, when the border-driven model is used, the forest height directly depends on the sample distance from the tumour centre, i.e., the tumour origin. When the samples come from the tumour centre, the forest height did not change significantly as the number of samples increased from 2 to 9 because all samples are equally distant from the centre. As soon as we added the tenth sample, which is the farthest from the centre, the height increased. The tallest forests are those due to samples at the tumour edge. **(B)** The relation between the number of sampled cells and the number of nodes in the forest as the growth model changes among border-driven and homogeneous growth models, and the position of the samples varies between the centre and the edge of the tumour. The data represented in this figure were collected by setting the duplication and death rates to 0.4 and 0.01, respectively. **(C)** The ratio of the duplication and death rates affects the number of nodes in the forest. This plot shows the relationship between the ratio of the two rates and the number of nodes when the border-driven model is adopted, with samples collected at the tumour edge and $\delta$ set to either 0.01 or $\lambda/10$. **(D)** The relation between the number of nodes in the forest $q$ and the sample forest building time when duplication and death rates were set to 0.4 and 0.01, respectively. Even though the number of nodes depends on the sampling position, the relationship between building time and the number of nodes appears to be independent of the growth model or the sampling position.

The data suggest a linear relation between the number of sampled cells and the number of nodes in the resulting forest. As for forest height, there's no apparent dependence on sample position when the homogeneous growth model is used. Instead, in the border-driven model, the closer a sample is collected to the tumour centre, the greater the first-order coefficient that maps the number of sampled cells to the number of forest nodes. Supplementary Fig. S8B relates the number of sampled cells, the growth model, the sampling position, and the number of nodes in the resulting forest.

The ratio $r = \lambda/\delta$ of duplication to death rates affects the number of nodes in the forest. When the border-driven model is adopted, the samples are collected at the tumour edge, and $\delta = 0.01$, the number of nodes in the sample forests grows as $1/r$. When, instead, $r = 10$, the number of nodes in the forest is constant without any dependency on $\lambda$. Supplementary Fig. S8C depicts the relation between $r$ and the number of nodes as $\lambda$ changes.

Although the time complexity for building a sample forest is $O(q \log q)$, Supplemenatry Table T6 highlights a linear behaviour in most cases. This phenomenon may be due to the $O(q)$ disk accesses. When the number of nodes in the forest grows, for instance by decreasing the ratio between the duplication rate and the death rate (Supplementary Fig. S8C), the $O(q \log q)$ trend tends to arise. The growth model and sampling positions apparently do not affect the relation between the number of nodes and the building time (Supplementary Fig. S8D).

**Supplementary Table T6.** The AICc of the $1/q$, $q$, $q \log q$, $q^{3/2}$, $q^2$, and $e^q$ models representing the relation between the number of nodes in a sample forest and its building time when the sampling occurred on a single-mutant simulation having 0.01 as the death rate. The lowest value corresponds to the most likely model that fits the data. Each row reports the AICs for a particular combination of growth model and sampling position. The minimum AICcs in each row are underlined. The values whose difference from the row minimum is less than 0.5% are reported in bold font.

| Growth model | Sampling Pos. | Dup. rate | $1/q$ | $q$ | $q \log q$ | $q^{3/2}$ | $q^2$ | $e^q$ |
|---|---|---|---|---|---|---|---|---|
| Border-driven | Edge | 0.1 | -58.74 | **-202.83** | **-203.71** | -182.34 | -147.48 | -148.1 |
| Homogeneous | Edge | 0.1 | -105.17 | **-240.25** | **-242.08** | -227.73 | -195.83 | -199.05 |
| Border-driven | Center | 0.1 | 23.48 | **-17.22** | -17 | -13.68 | -7.2 | -12.28 |
| Homogeneous | Center | 0.1 | -117.55 | **-347.63** | **-346.52** | -268.21 | -211.12 | -215.21 |
| Border-driven | Edge | 0.2 | -82.01 | **-250.04** | **-249.03** | -212.46 | -170.89 | -174.45 |
| Homogeneous | Edge | 0.2 | -133.85 | **-401.58** | -393.73 | -287.26 | -228.18 | -234.3 |
| Border-driven | Center | 0.2 | -2.99 | -128.51 | **-130.08** | -119.53 | -92.52 | -99.07 |
| Homogeneous | Center | 0.2 | -123.19 | **-343.45** | **-344.04** | -274.53 | -218.23 | -223.29 |
| Border-driven | Edge | 0.3 | -94.27 | **-265.7** | **-268.14** | -237.8 | -193.87 | -198.27 |
| Homogeneous | Edge | 0.3 | -134.24 | **-389.95** | -383.1 | -286 | -228.19 | -233.99 |
| Border-driven | Center | 0.3 | -31.48 | **-298.96** | -286.16 | -181.15 | -124.92 | -135.07 |
| Homogeneous | Center | 0.3 | -124.87 | **-394.71** | **-393.84** | -282.5 | -220.82 | -225.32 |
| Border-driven | Edge | 0.4 | -110.74 | **-311.52** | **-310.58** | -255.93 | -205.8 | -212.34 |
| Homogeneous | Edge | 0.4 | -123.58 | **-392.94** | -381.87 | -275.88 | -218.07 | -224.6 |
| Border-driven | Center | 0.4 | -30.75 | **-343.57** | -304.58 | -177.96 | -121.92 | -129.63 |
| Homogeneous | Center | 0.4 | -115.98 | **-395.54** | -390.4 | -272.56 | -211.31 | -216.07 |
| Border-driven | Edge | 0.5 | -101.76 | **-310.8** | -307.11 | -244.8 | -194.37 | -199.06 |
| Homogeneous | Edge | 0.5 | -132.46 | **-410.19** | -403.4 | -288.54 | -228.34 | -233.55 |
| Border-driven | Center | 0.5 | -46.39 | **-299.91** | -285.23 | -190.53 | -136.93 | -145.12 |
| Homogeneous | Center | 0.5 | -126.25 | **-384.68** | -379.41 | -278.85 | -219.61 | -224.5 |

**Building mutation engine** Supplementary Fig. S9 shows the mutation engine building and loading time as the parameters `context_sampling` and `max_repetition_storages` vary. Supplementary Fig. S9A shows that the parameter `max_repetition_storag` positively impacts both the index building and loading times, even though, by increasing it by 5 time, the building and loading times increase by less than 10%. By contrast, the building and loading times are not significantly affected by the parameter `context_sampling` (Supplementary Fig. S9B).

**Building phylogenetic forests** Supplementary Fig. S10 depicts the tests about phylogenetic forest building time.

When the homogeneous growth model was adopted, or the samples were collected at the tumour edge, the building time is linear with respect to the mutation rate (Supplementary Fig. S10A). When, instead, the border-driven growth model was used, and the samples were collected at the tumour centre, a non-linear behaviour, possibly due to the large difference in the number of nodes of the considered forests (Supplementary Fig. S8B), emerged. In the second batch of tests, the mutation rate was set to $1 \times 10^{-9}$, and the pre-neoplastic mutations varied. The differences in building time due to the sampling position became less relevant as the number of pre-neoplastic mutations increased (Supplementary Fig. S10B). In the third batch of tests, the CNAs altered the amount of DNA in cells, increasing the number of somatic mutations in the sampled cells, even though the mutation rate was fixed to $1 \times 10^{-9}$. This change produced a linear dependency between the number of CNAs and the phylogenetic forest generation time (Supplementary Fig. S10C).

Supplementary Fig. S10D depicts the relation between the number of sampled cells and the number of somatic mutations when the mutation rate and the number of pre-neoplastic mutations were set to $1 \times 10^{-9}$ and 2000. The two quantities appear to be linearly related. The tests that adopted the border-driven growth model and collected samples from the tumour border yielded the highest number of somatic mutations per sampled cell, due to the highest forest heights (Supplementary Fig. S8A).

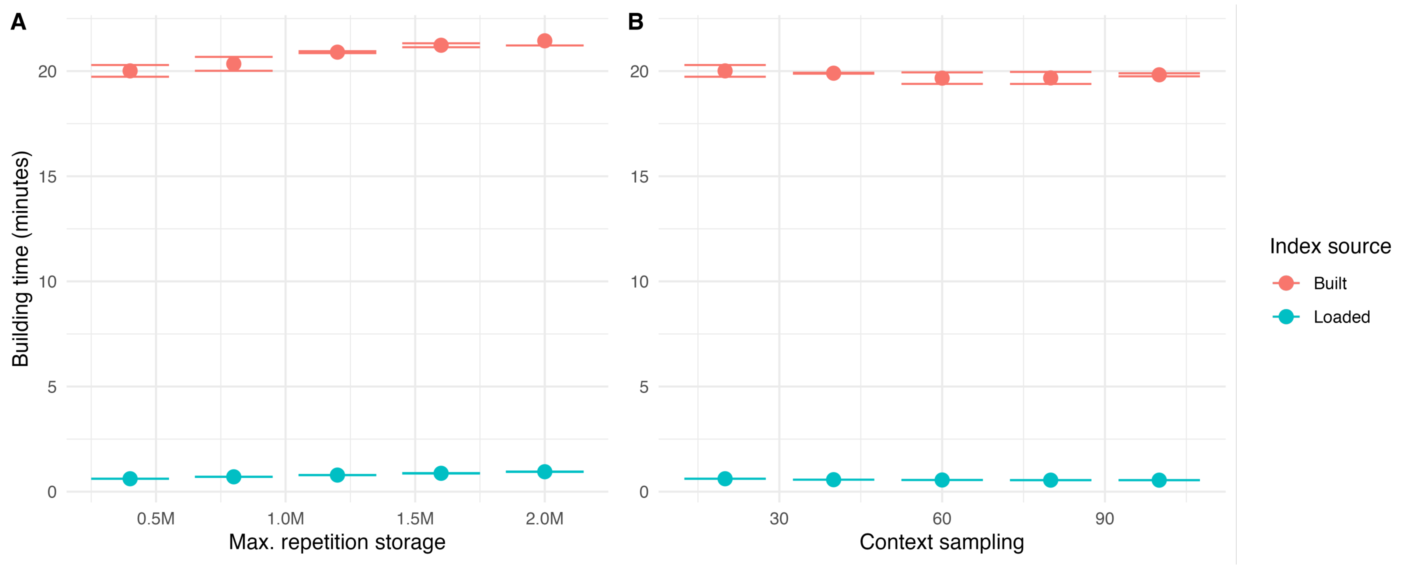

**Supplementary Fig. S9.** Building and loading the mutation engine as the pre-sampling parameters, `context_sampling` and `max_repetition_storage`, change. The building times reported in these figures do not account for the download and unpack time. **(A)** The mutation engine building and loading time when `context_sampling` is 20. **(B)** The mutation engine building and loading time when `max_repetition_storage` is $4 \times 10^5$.

This peak in the number of somatic mutations highlighted the logarithmic term of the complexity expression when the relation between the number of sampled cells and the phylogenetic forest generation time is drawn (Supplementary Fig. S10E).

**Sequencing** The sequencing simulation time increases with the mutation rate (Supplementary Fig. S11A) and the coverage (Supplementary Fig. S11B). As forecasted by Eq. S11, the sequencing simulation time decreases as $1/\ell_r$ (Supplementary Fig. S11C). Finally, the number of amplifications affects the sequencing simulation time: as expected, the more amplifications, the longer the simulation time (Supplementary Fig. S11D).

#### B. The SCOUT cohort

The Simulated Cohort of Universal Tumours (SCOUT) is a carefully curated collection of prototypical tumours generated with the ProCESS framework, each accompanied by synthetic whole-genome sequencing (WGS) data. Every tumour reflects biologically realistic profiles reported in the literature, including canonical driver events, copy-number alterations, and characteristic mutational signatures. At the same time, specific features are deliberately engineered to challenge analytical methods and enable systematic evaluation of key computational tasks, including:

- Accurate calling of somatic variants, encompassing single nucleotide variants (SNVs), insertions/deletions (INDELs), and copy-number alterations (CNAs);
- Mutational signature deconvolution, enabling the identification and quantification of underlying mutagenic processes shaping tumour genomes;
- Subclonal deconvolution of complex VAF mixtures, allowing reconstruction of intratumour heterogeneity by resolving the composition and prevalence of distinct tumour clones.

Beyond serving as a benchmark for analytical methods in cancer genomics, SCOUT functions as a ”manual” for interpreting bulk WGS data in the context of tumour evolution. The cohort spans a wide range of evolutionary and clinical scenarios, highlighting the main interpretational challenges that link genomic observations to underlying tumour dynamics. These challenges include:

- Reconstructing latent clonal dynamics, lineage replacement, and the emergence of late subclones;
- Associating mutational processes with specific subclones and temporal intervals;
- Distinguishing plastic, non-genetic relapse from genetically driven resistance;
- Integrating the effects of therapy-induced mutagenesis with pre-existing clonal architecture to elucidate the impact of clinical interventions.

Each sample within the SCOUT cohort is generated through a fully reproducible computational pipeline (Supplementary Fig. S12), structured into four main steps:

1. Simulating tumour growth and the accumulation of mutations using the ProCESS framework;
2. Simulating sequencing reads that mimic whole-genome sequencing data derived from the simulated tumours;

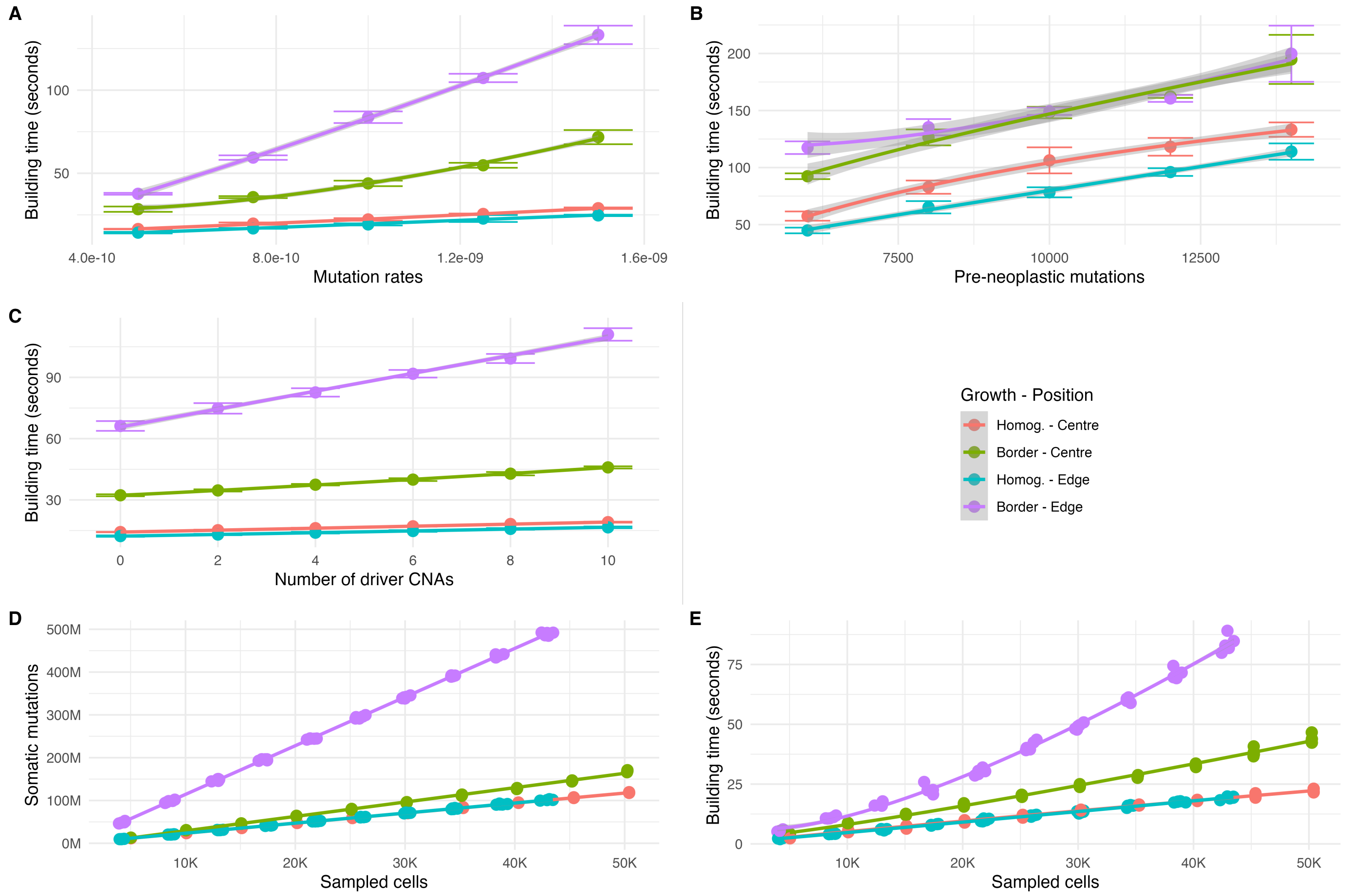

**Supplementary Fig. S10.** The time required to build a phylogenetic forest. We considered the sample forest generated during the previous tests and used the ProCESS setup `GRCh38`, which includes the homonimum human reference genome, to build the phylogenetic forests corresponding to the sampled forests generated by the previous set of tests when $\lambda = 0.4$ and $\delta = 0.01$. Moreover, we set $\{ID2 = 0.6, ID13 = 0.2, ID21 = 0.2\}$ and $\{SBS13 = 0.2, SBS1 = 0.8\}$ as ID and SBS mutational exposures, respectively. **(A)** The phylogenetic forests generation time when 2000 pre-neoplastic mutations were evenly spread between indels and SBSs and the mutation rate ranged between $5 \times 10^{-10}$, $7.5 \times 10^{-10}$, $1 \times 10^{-9}$, $1.25 \times 10^{-9}$, and $1.5 \times 10^{-9}$ for both indels and SBSs. **(B)** The building time as the number of pre-neoplastic mutations varied in the set $\{6000, 8000, 10\,000, 12\,000, 14\,000\}$ when the mutation rate is set to $1 \times 10^{-9}$. **(C)** The mutation rate was set to $1 \times 10^{-9}$, no preneoplastic mutations were added, and the only mutant had between 0 and 10 amplifications of chromosome 1 as driver mutations. **(D)** The relation between the number of sampled cells and the number of overall somatic mutations when the mutation rate, the number of pre-neoplastic mutations, and the number of CNAs were $1 \times 10^{-9}$, 2000, and 0, respectively. **(E)** The relation between the number of sampled cells and the phylogenetic forest building time when the mutation rate, the number of pre-neoplastic mutations, and the number of CNAs were $1 \times 10^{-9}$, 2000, and 0, respectively.

3. Applying the nf-core/sarek pipeline for standardized variant calling and quality control from the simulated sequencing data;
4. Executing the nf-core/tumourevo pipeline to perform downstream modelling of tumour evolution, including clonal deconvolution and mutational signature analysis.

Each of these steps is described in detail in the sections that follow.

#### B.1. ProCESS simulation of tumour growth and mutational data

In this step, we simulate tumor growth and sampling, carefully tailoring the process to reflect the unique characteristics of each sample. The number of cells in each clone is recorded at every simulation step, enabling fine-grained reconstruction of evolutionary lineages.

At any key moment in the evolutionary process such as before the introduction of a new clone, before and after simulating a treatment, or at the final stopping time, we collect tissue information by generating a spatial plot of the tumour. This provides a snapshot of the cancer tissue at specific time points, offering spatial and clonal context for downstream analysis.

The sampling is performed according to the specific experimental design of each sample, ensuring that the simulation mimics realistic biological and technical constraints.

Once a sample forest representing the phylogenetic relationships among the collected cells is obtained, we construct the mutation engine. This engine introduces somatic alterations to the nodes of the phylogenetic tree, thus capturing the evolutionary history and genetic diversity of the tumour.

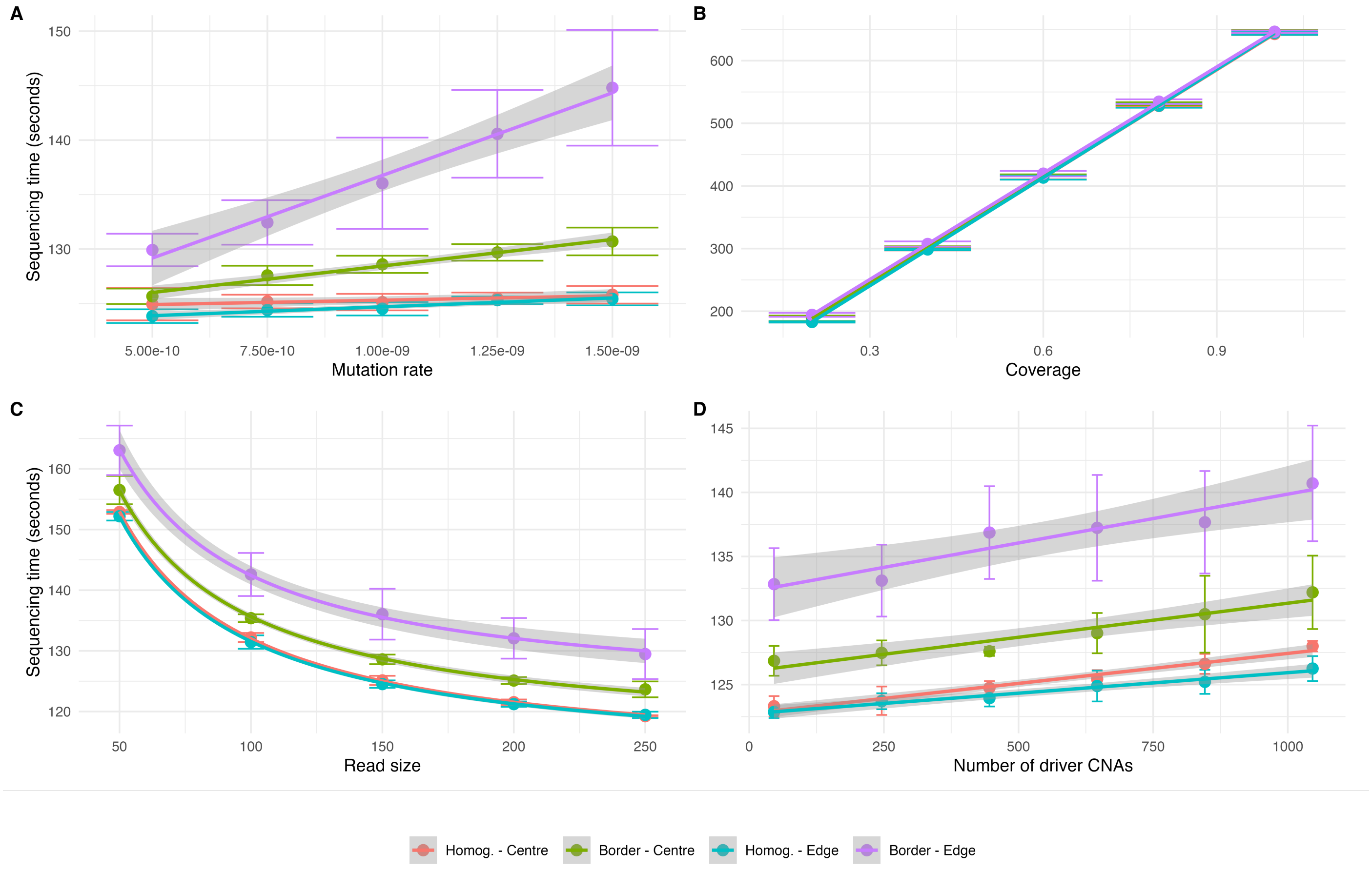

**Supplementary Fig. S11.** The time required to simulate a sequencing. We considered the phylogenetic forests built during the previous sets of tests when 10 samples were collected, and 2000 pre-neoplastic mutations were placed. From these forests, we extracted the subforests associated with the last sample, and we simulated its sequencing. **(A)** The time required to produce a 0.1-coverage output is linear in the mutation rates. **(B)** When the mutation rate is set to $1 \times 10^{-9}$ and the number of pre-neoplastic mutations varies in $\{6000, 8000, 10\,000, 12\,000, 14\,000\}$, the sequencing simulation time is linear in the coverage. **(C)** Instead, it decreases as the reciprocal of the read size when the mutation rate is set to $1 \times 10^{-9}$ and the number of pre-neoplastic mutations is 2000. **(D)** Finally, the simulation time grows as the number of CNAs in each sampled cells.

Specifically, the engine allows the user to set the reference genome of sequencing data, configure mutation rates for each clone in the simulation, including rates for single nucleotide variants (SNVs), insertions/deletions (INDELs), and copy number aberrations (CNAs). In addition, it supports the nesting of specific mutations and/or CNAs within all cells of a given clone to define its genotype.

Finally, through the mutation engine, we select the catalogue of mutational signatures to include in the simulation, specifying their periods of activity and relative exposures according to the specific characteristics of each sample.

#### B.2. ProCESS simulation of sequencing

We simulate DNA WGS sequencing with ProCESS for 12 combinations of coverage and purity. We considered tumour sequencing with coverages 50×, 100×, 150×, 200× and purities 0.3, 0.6, 0.9. We also obtain a further set of reads corresponding to a 30X simulated sequencing of normal tissue. The cells in the normal sample share with cancer cells the so-called germline mutations. The cohort is made available as tables of simulated mutation and copy number data and as raw FASTQ files output of ProCESS sequencing. Raw data can also be regenerated using a singularity image of ProCESS, which is the same we used to generate this version of the SCOUT cohort. On a standard HPC system, generating a single SPN (tumour and matched normal samples) takes about 36 hours thanks to the data generation algorithm implemented in ProCESS, which runs in parallel for each chromosome. To further parallelise, we divided the maximum target sequencing depth of 200× into 40 chunks of 5× coverage each. Sequencing reads were simulated for all combinations of tumour purity and coverage using the `simulate_seq` and `simulate_normal_seq` functions from the ProCESS package, and outputs were saved in SAM format. To ensure compatibility with standard bioinformatics workflows, a custom post-processing step was developed to merge chromosome-specific SAM files into sample-specific paired-end FASTQ files. The workflow starts with the creation of chromosome-level SAM files directly from ProCESS. These individual files are then combined into a single BAM file using `samtools merge`. After merging, the BAM file is divided into separate files corresponding to each simulated sample. Each of these sample-specific BAM files is then converted into paired-end FASTQ files using `samtools fastq` command. As a

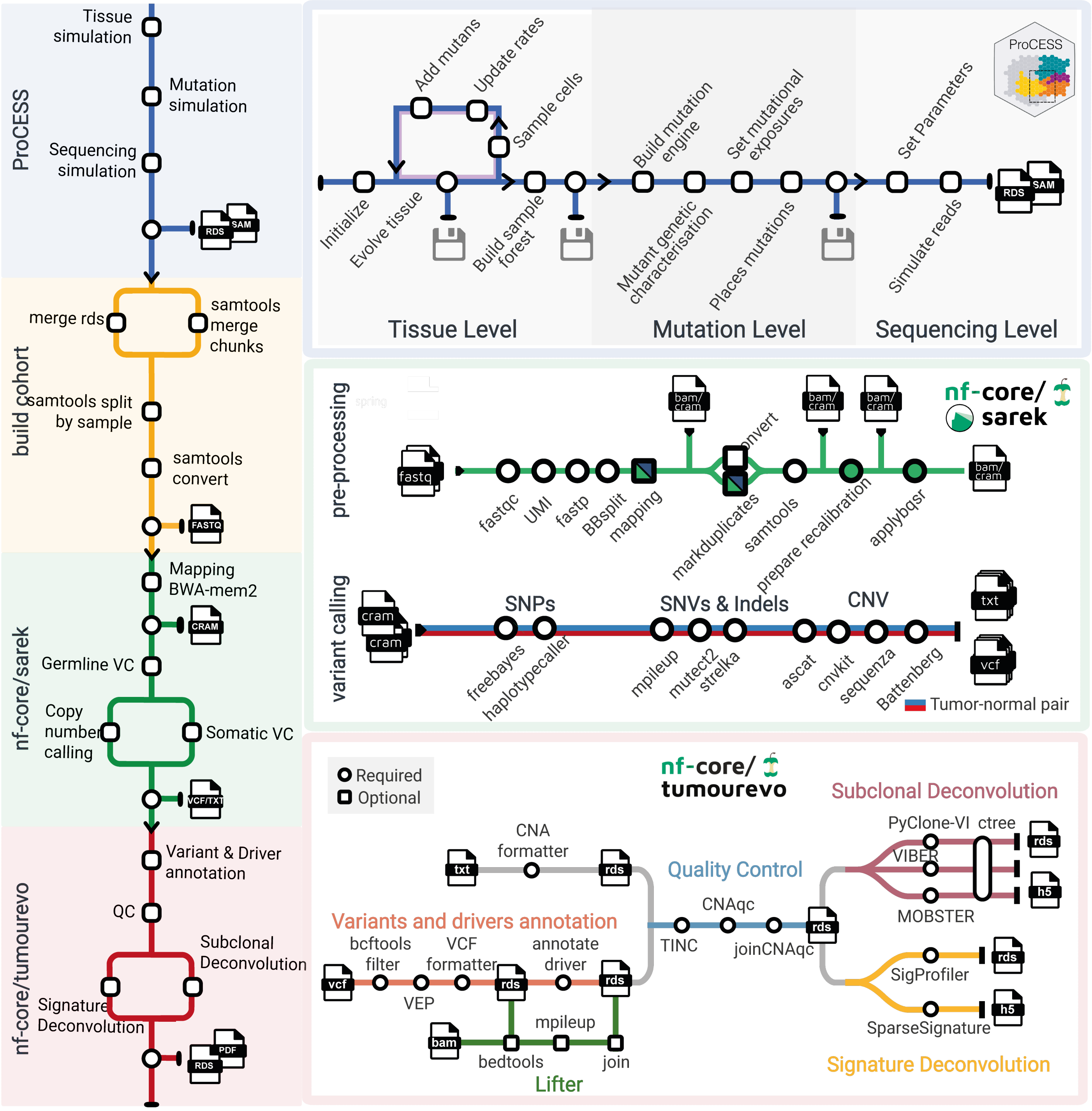

**Supplementary Fig. S12. SCOUT cohort generation pipeline.** The four main steps of data generation and analysis: (i) Tissue simulation and mutation engine set up with ProCESS; (ii) Sequencing of raw reads for different coverages by parallelizing into low-coverage chunks: ; (iii) Alignment, somatic and germline variant calling with nf-core/sarek; (iv) Variant annotation, quality control, subclonal and signature deconvolution with nf-core/tumourevo.

result, 40 paired-end FASTQ files were generated for each sample at each purity level. To reconstruct the target sequencing depths required for downstream alignment and preprocessing, subsets of these FASTQ files were merged. Specifically, effective coverages of 50×, 100×, 150×, and 200× were obtained by combining 10, 20, 30, and 40 paired-end FASTQ files, respectively. Publicly available sequencing data are typically distributed as a single paired-end FASTQ file corresponding to the maximum coverage (200×). Therefore, all downstream analyses were performed on the reconstructed 50×, 100×, and 150× datasets, with the original 200× dataset serving as the full-coverage reference.

#### B.3. Cohort description

##### *B.3.1. Overview*

The simulated cohort spans a wide range of realistic tumour types and experimental settings. It includes adenocarcinomas (colorectal and lung), haematological malignancies (chronic lymphocytic leukaemia and acute myeloid leukaemia), breast cancer, and glioblastoma. All tumours undergo multiple subclonal expansions during their evolutionary history; in some cases, these occur early and the tumour appears largely monoclonal at the time of sampling, whereas in others they arise later, resulting in a polyclonal structure at the final observation time.

For selected tumours, we simulated the effects of therapy following clinically established treatment protocols, incorporating resistance mechanisms driven either by the expansion of pre-existing clones or by de novo resistant populations emerging under treatment. Tumour evolution is further shaped by a variety of mutagenic processes acting throughout disease progression in a tumour-type- and treatment-dependent manner. These include endogenous processes, such as mismatch repair deficiency marking specific evolutionary stages, as well as exogenous, therapy-associated processes that are active within defined temporal windows.

For each simulated tumour, we generated multi-region and/or longitudinal sequencing data under different configurations of tumour purity and sequencing coverage, as described in the previous section. Analysis of the resulting genomic data, including subclonal reconstruction and mutational signature deconvolution, enables recovery of the latent evolutionary dynamics and, when treatment is present, allows these dynamics to be aligned with the clinical timeline.

In the following sections, we provide a detailed description of the simulated evolutionary processes, sampling strategies, and genomic analyses for each tumour in the cohort. For each case, we also highlight the specific analytical challenge posed by reconstructing tumour evolution from bulk WGS data, illustrating scenarios in which polyclonality, temporal ordering of driver events, therapy-induced mutational signatures, or subtle subclonal expansions contribute to interpretational complexity.

##### *B.3.2. General aspects of the simulation*

We distinguish between general and sample-specific features in our simulation framework. General features define baseline properties of the system that apply across all tumours, independently of their specific evolutionary trajectories, whereas sample-specific features govern the clonal dynamics and experimental configurations of individual simulations. We first describe the general aspects of the simulation setup, followed by the specific configurations adopted in this study.

We selected simulation parameters with a particular focus on supporting the practical analysis of clonal structure and evolutionary dynamics from bulk WGS data:

- Border-growth model: Tumor growth is modeled using a boundary-driven ("border") growth model, which preserves a correlation between spatial location and clonal ancestry. This enables genetically distinct regions to arise even in tumours that are monoclonal at the time of sampling, making the model well suited for studying spatial heterogeneity and sampling effects.
- `sim$death_activation_level = 50`: This parameter prevents newly emerging subclonal populations from going extinct due to stochastic effects before they can establish. By reducing early cell death in low-density regions, it allows nascent clones to expand before density-dependent turnover becomes dominant.
- Treatment simulation: Therapeutic interventions are simulated by modifying the birth and death rates of existing clones according to their sensitivity to the administered treatment. In selected cases, therapies also induce specific mutational processes, which are activated in the mutation engine during predefined treatment windows.
- Reference genome: Synthetic bulk whole-genome sequencing (WGS) data are generated using the human reference genome GRCh38.
- `num_of_preneoplastic_SNVs = 800`: We introduce 800 single nucleotide variants (SNVs) during a pre-neoplastic phase to model background mutational accumulation prior to malignant transformation.
- `num_of_preneoplastic_indels = 200`: Additionally, 200 insertions and deletions (INDELs) are introduced to capture early genomic variability.
- Aging-related mutational signatures: All tumours are exposed to clock-like SNV signatures SBS1 and SBS5, as well as the INDEL signature ID1, with exposure levels varying across evolutionary stages.

Sample-specific features are summarised across five tables. Supplementary Table T7 recapitulates the main characteristics of each tumour's evolutionary history, including simulated treatments and experimental design (multi-region and/or longitudinal sampling).

For computational reasons, clonal growth rates are not kept constant throughout the simulations. Following the emergence of a new subclone, the growth rates of pre-existing clones are progressively reduced. This adjustment mitigates the computational burden associated with exponential expansion of dominant populations: without it, newly introduced subclones would require prohibitively long simulation times to reach detectable sizes. By dynamically scaling down the growth rates of older clones, we ensure that emerging subclones can expand within a computationally feasible timeframe. Implementation details are provided in the simulation code associated with each sample. Supplementary Table ST8A reports the mean birth and death rates of each clone over the course of the simulation, summarising overall population dynamics.

Supplementary Table ST8B provides information on the collected samples for each tumour, including the number of sampled cells, sampling time points, and clonal composition. Driver mutation profiles, mutation rates, and copy-number alteration rates for each clone are reported in Supplementary Table ST8C-E-F-G-H. Finally, Supplementary Table ST8D-I summarises the mutational signatures simulated in each tumour, together with their relative exposures and the evolutionary time windows during which they are active.

| Patient ID | Type | Samples | Description | Tumour Type |
|---|---|---|---|---|
| SPN01 | P (n=4) | b=3 (t=1) | Micro-satellite stable (MSS) colorectal cancer with a single clonal population, sampled at a single time-point with two multi-region biopsies. | CRC |
| SPN02 | P (n=3) | b=2 (t=1) | Micro-satellite instable (MSI) colorectal cancer composed of a main clonal population, sampled at a single time-point with two multi-region biopsies. | CRC |
| SPN03 | P (n=3) | b=4 (t=4) | Chronic lympocitic laeukemia in a watch-and-wait (i.e., untreated) scenario, with multiple subclones sampled longitudinally over four time-points. | CLL |
| SPN04 | P/R (n=3) | b=2 (t=2) | Acute myeloyd laeukemia with a dominant clone, relapsing platinum-based chemotherapy with, sampled longitudinally before treatment and at relapse. | AML |
| SPN05 | P (n=5) | b=3 (t=1) | Mismatch deficient breast cancer, with many subclones, sampled at one time-point with with three multi-region biopsies. | BRCA |
| SPN06 | P/R (n=6) | b=5 (t=3) | Smoking-associated lung adenocardinoma with many subclones, treated with two lines of chemotherapy, both time relapsing with a de-novo clone, sampled at three time-points with five multi-region biopsies. | LUAD |
| SPN07 | P/R (n=6) | b=5 (t=2) | Glioblastoma with many subclones, treated with a line of chemotherapy, relapsing with a hypermutant clone, sampled at three time-points with five multi-region biopsies. | GBM |

**Supplementary Table T7.** The SCOUT cohort statistics: Primary (P), Relapse (R), Number of simulated clones (n), Number of sampled time-points (t), Total number of biopsies (b).

##### *B.3.3. SPN01*

SPN01 represents a prototypical case of microsatellite-stable (MSS) colorectal cancer, reflecting a common evolutionary trajectory reported in the literature [24, 25] (Supplementary Fig. S13). Tumour expansion begins with a truncating mutation in *APC* (R1450*), followed by complete inactivation of the gene through loss of heterozygosity (LOH) of the wild-type allele. The competition between the founding clone C1 and the homozygous *APC*-inactivated clone C2 results in a clonal sweep, establishing C2 as the dominant population. Subsequent partial inactivation of the tumour suppressor *TP53* (R175H) drives the emergence of a new subclone, C3, originating from C2.

Later in tumour evolution, oncogenic activation of *PIK3CA* (R88Q) occurs within C3, coinciding with a whole-genome doubling (WGD) event. This gives rise to clone C4, which outcompetes other populations and becomes the predominant clone. At the time of sample collection, the tumour is largely composed of C4 cells carrying all accumulated driver events.

In addition to clock-like mutational processes, three further mutational signatures are active from early tumour development: SBS88, associated with *E. coli* exposure; SBS18, linked to reactive oxygen species–induced damage; and SBS17b, a signature of unknown aetiology frequently observed in colorectal cancer [26]. These signatures produce characteristic mutations that accumulate throughout the tumour's lifetime.

Mutation rates for single nucleotide variants (SNVs) and insertions/deletions (INDELs) are constant across all clones, ensuring that observed heterogeneity arises from clonal dynamics rather than differences in mutational processes.

SPN01 is profiled using a multi-region sampling strategy, with three spatially distinct samples collected. Specifically, SPN01_1.2 is dominated by clone C3, SPN01_1.3 by the WGD subclone C4, and SPN01_1.1 exhibits polyclonality, with a larger contribution of C3 alongside C4. The copy-number profiles of SPN01_1.3 and SPN01_1.2, are tetraploid and diploid, respectively, revealing a late-emerging WGD-driven subclonal expansion.

The principal challenges are twofold. First, accurately calling the subclonal WGD event is non-trivial: standard copy-number callers optimised for diploid or uniformly polyploid tumours may miscall the ploidy state of the mixed sample SPN01_1.1 or fail to resolve the coexistence of diploid and tetraploid clones. Second, discriminating mutation clusters associated with genuine clonal expansions from clusters arising purely from spatial sampling variability across the three regions requires careful multivariate VAF analysis. Tracking the frequency of both passenger and driver mutation clusters across all sample combinations resolves the temporal sequence of key evolutionary events, including early TP53 loss-of-heterozygosity and the subsequent emergence of the WGD subclone, while also delineating the polyclonal structure at the time of sample collection.

###### B.3.4. SPN02

SPN02 represents a prototypical case of microsatellite-instable (MSI) colorectal cancer, reflecting the evolutionary dynamics of POLE-driven hypermutation [24, 25] (Supplementary Fig. S14). The tumour harbors a somatic BRAF V600E mutation present in the ancestral clone C1, which is characterised by clock-like mutational processes and a stable mutation rate [24, 25].

Progression occurs through sequential acquisition of driver events, resulting in a nested clonal architecture. The emergence of a PIK3CA E545K mutation gives rise to clone C2, which confers a proliferative advantage and leads to a clonal sweep, without substantially altering either the active mutational processes or the overall mutation rate. Subsequently, acquisition of a POLE V411L mutation generates clone C3, which induces a microsatellite instable, hypermutant phenotype. This transition is associated with a marked increase in both SNV and indel burdens and the activation of mismatch repair–related mutational signatures, including SBS10a and SBS10b for SNVs and ID7 for indels [24, 25]. Clock-like mutational processes remain detectable in C3 but at reduced relative contribution. The increased fitness conferred by the hypermutated state leads to a final clonal sweep, such that at the time of disease detection the tumour is entirely dominated by clone C3.

SPN02 is profiled using a multi-region sequencing design, with two samples collected simultaneously. Despite the monoclonal structure of the final tumor, the spatial organization of somatic mutations in solid tumours induces a non-trivial coupling between genotype and sampling location. As a result, both samples SPN02_1.1,SPN0_1.2 exhibit a subset of mutations that are private to each region, even in the absence of additional evolutionary bottlenecks or true subclonal diversification. This pattern is also reflected in the structure of the latent phylogenetic forest, which reveals two distinct branches onto which these private mutations map. Importantly, these branches do not correspond to independent evolutionary lineages, but rather to spatial segregation of mutations within a single dominant clone. This introduces a key interpretative challenge: distinguishing clusters arising from spatial segregation effects from those reflecting genuine evolutionary events. The difficulty is further exacerbated by the POLE-driven hypermutator phenotype, which generates an extremely high burden of SNVs and indels and leads to an inflated rate of candidate driver calls. In this context, the abundance of putative driver mutations can obscure the distinction between spatially structured but evolutionarily neutral clusters and true subclonal expansions.

###### B.3.5. SPN03

SPN03 represents a prototypical case of chronic lymphocytic leukaemia (CLL), characterized by stepwise clonal evolution and sequential acquisition of canonical driver events (Supplementary Fig. S15). The simulated tumour comprises three major clones that emerge through successive selective sweeps during the evolutionary process.

The founding clone C1 originates from a somatic NOTCH1 p.P2514Rfs*4 mutation accompanied by a loss of heterozygosity (LOH) on chromosome 13q14.2, both well-established lesions promoting aberrant B-cell proliferation [27–29]. A descendant subclone C2 subsequently arises from within C1 after the acquisition of a TP53 p.R175H mutation, conferring increased genomic instability and a proliferative advantage. In the later stages of evolution, a third clone C3 emerges through a MAP2K1 p.P124L mutation [30, 31]. This clone outcompetes its parental lineage and start to become dominant toward the end of the simulation, illustrating a typical pattern of subclonal replacement.

Throughout tumour evolution, clock-like mutational processes, SBS1 and SBS5, are active, modelling the slow accumulation of somatic mutations typical of indolent CLL, and SBS9, typically associated with CLL that possess immunoglobulin gene hypermutation (IGHV-mutated) [32, 33]. No treatment-associated or hypermutant processes are simulated, and none of the driver events induce secondary mutagenic effects.

The tumour is simulated under a watch-and-wait clinical scenario, mimicking active surveillance prior to therapeutic intervention. Four longitudinal samples are collected at distinct evolutionary time points to capture the temporal dynamics of clonal competition and replacement. Specifically, the first sample (SPN03_1.1) is monoclonal and composed exclusively of clone C1. In the second sample (SPN03_2.1), both C1 and C2 are represented, marking the onset of clonal competition. A subsequent selective sweep leads to fixation of C2 in the third sample (SPN03_3.1). Finally, the last sample (SPN03_4.1) exhibits renewed polyclonality, comprising C2 together with the newly emerged and rapidly expanding C3 population.

The copy-number profile of chromosome 13q in sample SPN03_1.1, inferred from bulk WGS data, reveals the ancestral clonal deletion of the q arm, as evidenced by the allele-specific B-allele frequency (BAF) and depth-ratio profiles and confirmed by the corresponding VAF distribution. Multivariate analysis of VAFs across samples resolves clusters of mutations shared across time points, corresponding to ancestral clones, alongside private clusters marking newly emerging subclones. Annotated driver mutations co-segregate with these clusters, anchoring the inferred evolutionary structure. By comparing how both passenger and driver mutation clusters shift in frequency across the multivariate VAF distributions of all longitudinal sample combinations, it is possible to reconstruct the sequence of clonal competition, selective sweeps, and re-emergence of polyclonality over time.

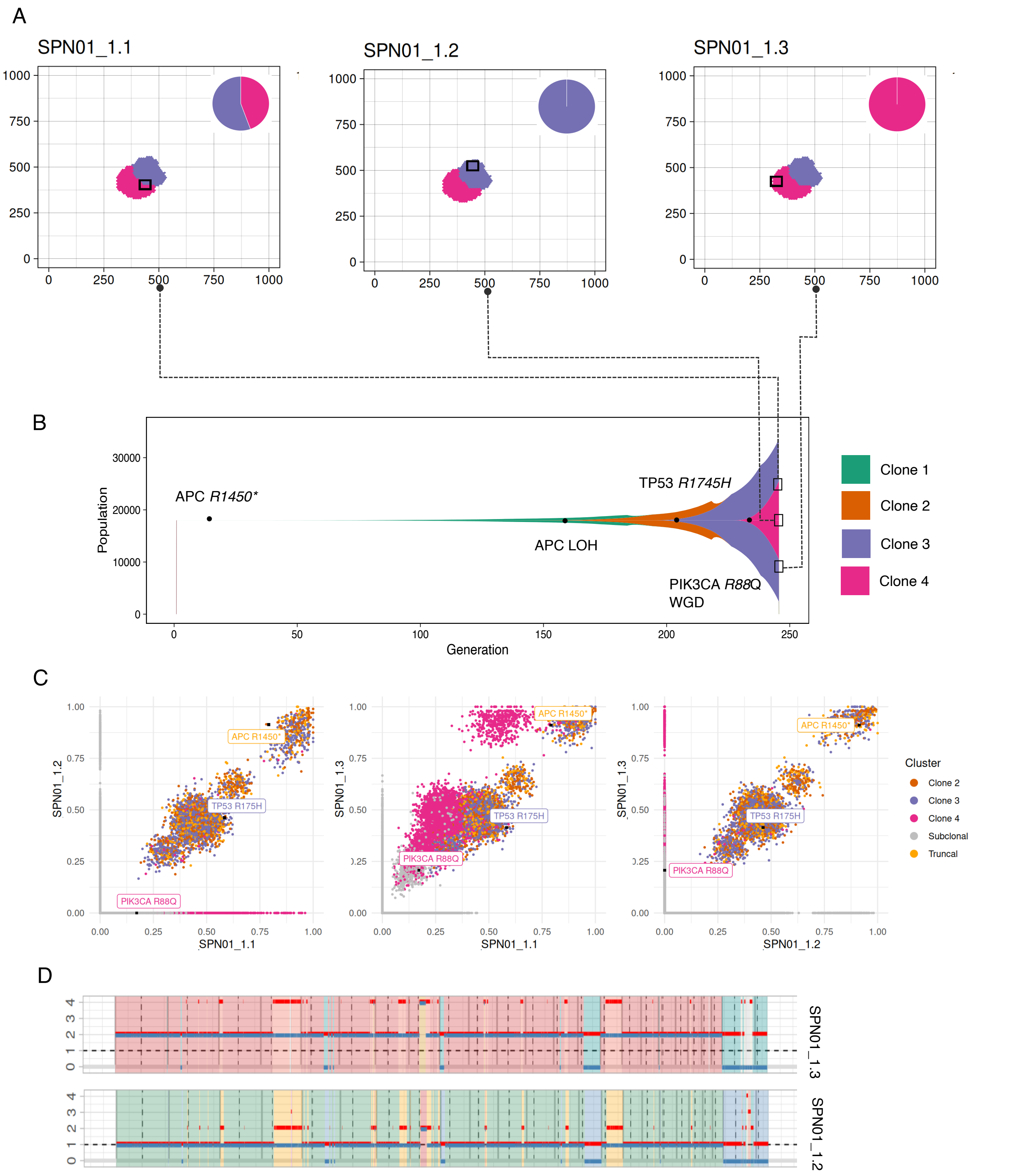

**Supplementary Fig. S13. SPN01 simulation.** (A) Sampling scheme showing the three spatially distinct tumour regions profiled at a single time point. (B) Muller plot of the simulated evolutionary dynamics, with sampling times and driver mutations annotated. (C) Multivariate VAF analysis across all samples, resolving clonal and subclonal structure with key driver mutations highlighted. (D) Allele-specific copy-number profiles of SPN01_1.3 and SPN01_1.2, illustrating the tetraploid and diploid ploidy states respectively.

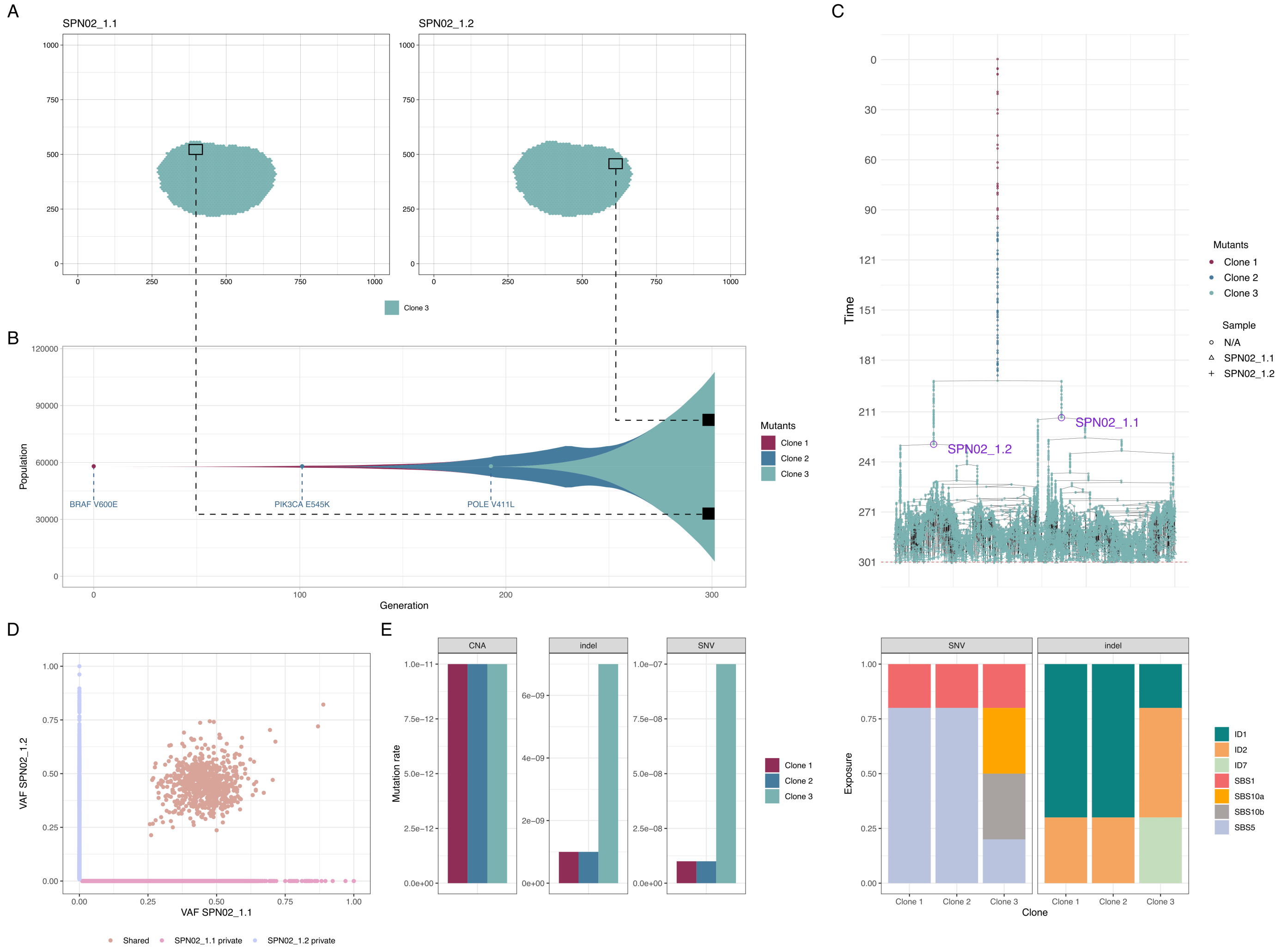

**Supplementary Fig. S14. SPN02 simulation.** (A) Sampling scheme showing the two spatially distinct tumour regions profiled at a single time point. (B) Muller plot of the simulated evolutionary dynamics, with sampling times and driver mutations annotated. (C) Phylogeny of the two samples with most recent common ancestors (MRCAs) annotated. (D) Multivariate VAF analysis across all samples, resolving clonal and spatially private mutations. (E) Mutation rates for CNAs, INDELs, and SNVs across the three clones. (F) SBS and ID mutational signatures associated with each clone.

This analysis recapitulates the underlying evolutionary history and provides a direct link between bulk sequencing data and the latent clonal dynamics.

The main analytical challenge in performing the above logitudinal analysis lies in the interpretation of the multivariate VAF structure across longitudinal samples, where the primary difficulty consists in distinguishing genuine clonal expansions from spurious clusters induced by sampling variability. In particular, such artefactual clusters may be incorrectly interpreted as evidence of new subclonal lineages, complicating the reconstruction of the true sequence of clonal sweeps. This is especially relevant in a longitudinal setting, where temporal sampling is intended to resolve the order of selective events but may also introduce apparent structure unrelated to underlying evolutionary dynamics.

An additional layer of complexity arises from the relatively low mutational burden of this tumour, which limits the resolution of mutational signature inference at the level of individual samples. As a consequence, accurate identification of the active mutational signatures and reliable estimation of their exposures become challenging, as low mutation counts lead to increased stochastic variability in the assignment of mutations to signatures.

##### B.3.6. SPN04

SPN04 represents a prototypical case of acute myeloid leukemia (AML), characterised by a largely linear evolutionary trajectory and relapse driven by non-genetic plasticity following platinum-based chemotherapy (Supplementary Fig. S16). The simulated tumour evolves through a sequence of successive driver events that establish a dominant clonal population prior to treatment.

The founding clone C1 arises from an IDH1 R132C mutation, a canonical early event in AML [34]. A subsequent amplification of MYC gives rise to clone C2, conferring a proliferative advantage and promoting clonal expansion. Tumour progression culminates in the acquisition of an NRAS Q61K mutation within C2, generating clone C3, which becomes the dominant population at diagnosis. Throughout this pre-treatment phase, mutagenesis is driven exclusively by clock-like processes, and both mutation and copy-number alteration (CNA) rates remain stable across clones.

Platinum-based chemotherapy is introduced after clonal dominance of C3 has been established. Treatment activates the chemotherapy-associated mutational signature SBS25, which remains active during the exposure window but does not alter baseline mutation or CNA rates. Although therapy substantially reduces the overall tumour burden, a subset of C3 cells survives treatment and retains proliferative capacity. Upon therapy cessation, these surviving cells re-expand, giving rise to relapse.

Relapse is characterised by a monoclonal architecture derived entirely from clone C3, with no evidence of newly acquired driver events or clonal replacement. This evolutionary pattern indicates that resistance is mediated by non-genetic plasticity rather than by positive selection of a genetically distinct resistant subclone. Longitudinal genomic profiling confirms the absence of an evolutionary bottleneck during therapy, as well as the stability of mutation and CNA rates throughout tumour evolution.

The tumour is profiled using paired pre-treatment and post-treatment samples (SPN04_1.1 and SPN04_2.1). In the pre-treatment sample, the allele-specific B-allele frequency (BAF) and depth-ratio profiles of chromosome 8 reveal the clonal MYC amplification, consistent with early clonal selection. Multivariate analysis of variant allele frequency (VAF) distributions across the pre-treatment and relapse samples shows no private mutation clusters emerging at relapse, indicating the absence of a genetically driven evolutionary bottleneck. From a mutational process perspective, signature inference correctly identifies the dominant constitutive processes; however, the chemotherapy-associated signature SBS25 is not detected in the relapse sample. This is expected, as the relapse is non-genetic, and therefore SBS25-associated mutations remain confined to low-frequency variants in the tail of the VAF distribution, making them difficult to detect with standard signature inference methods.

##### B.3.7. SPN05

SPN05 is a prototype mismatch repair-deficient breast cancer [35–37] composed of five distinct clones (Supplementary Fig. S17). The ancestral clone, C1, harbors a TP53 R240QWL mutation, which initiates tumour growth. In the early stages of tumour evolution, the cancer acquires biallelic inactivation of BRCA2 through the sequential acquisition of a missense mutation (E2028*, clone C2) and a loss of heterozygosity (LOH) event affecting the wild-type allele (clone C3).

This alteration induces mismatch repair (MMR) deficiency, leading to an elevated mutation rate and the activation of specific mutational signatures, including SBS3 and ID7. These signatures emerge following BRCA2 inactivation and persist throughout tumour evolution, co-occurring with the background clock-like mutational processes. The homozygous BRCA2-deficient clone (C3) eventually becomes dominant.

In the later stages of tumour development, two additional driver events give rise to independent subclonal expansions: an LOH of the TP53 wild-type allele (clone C4) and an LOH of CDKN2A (clone C5). These clones expand in parallel with C3 and coexist at the time of sampling, reflecting a highly structured subclonal architecture.

A multi-region sampling strategy is employed for SPN05, involving three biopsies collected at a single time point that capture the spatial heterogeneity of the tumour at the time of profiling. One biopsy is monoclonal and consists exclusively of cells from clone C4. The other two biopsies are polyclonal, containing mixtures of C3 and C4 in sample SPN05_1.2, and C4 and C5 in samples SPN05_1.1 and SPN05_1.3.

The mismatch repair-deficient and hypermutated state leads to a very high mutational burden, which in turn results in a large number of candidate driver calls, many of which are likely false positives. This complicates downstream interpretation, as driver assignment becomes less reliable in distinguishing biologically meaningful events from spurious signals. As a consequence, the main analytical challenge in SPN05 lies in the correct classification of mutation clusters, in order to separate true driver-associated subclonal expansions from clusters that arise purely as a consequence of spatial sampling effects across biopsies.

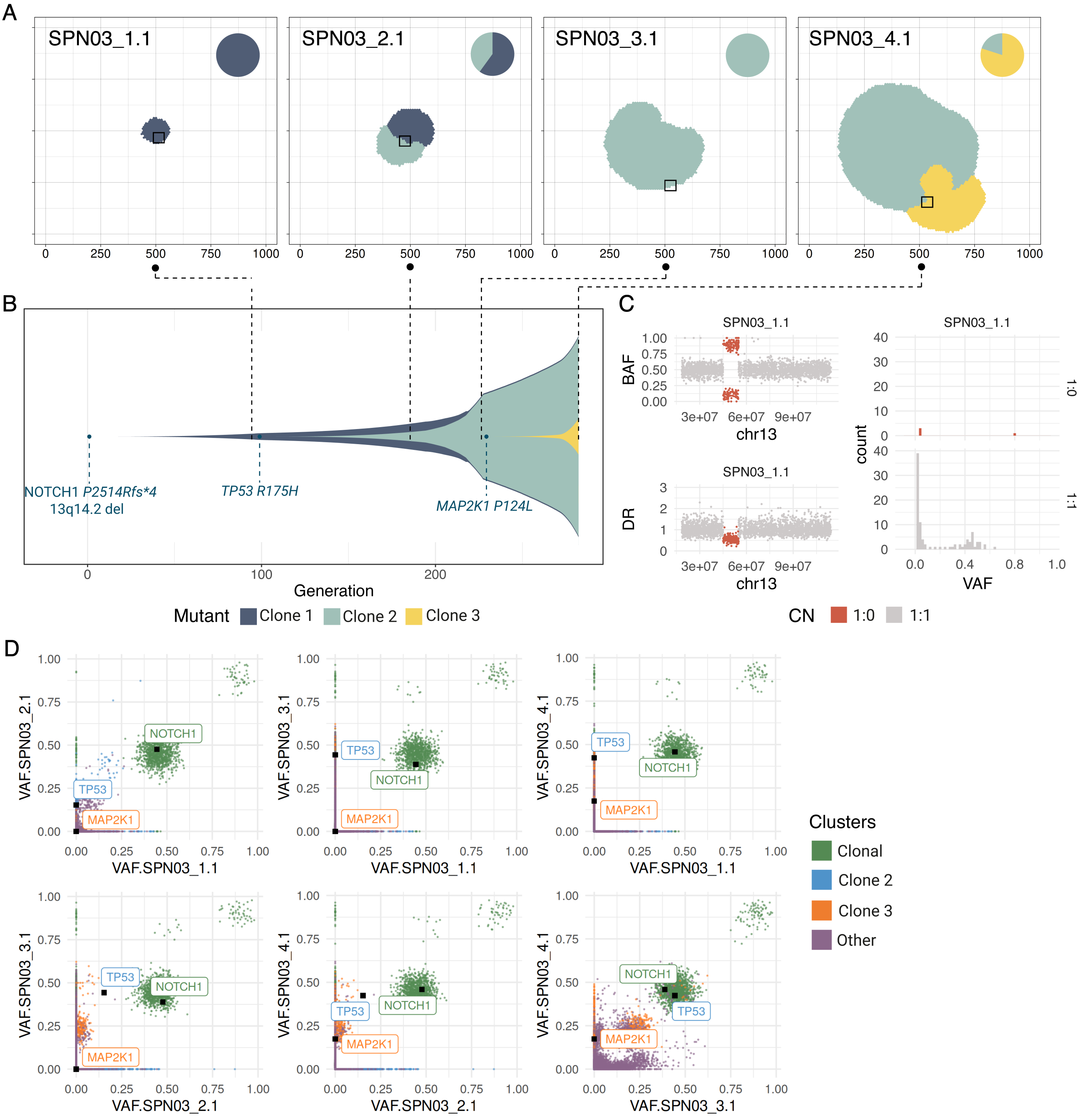

**Supplementary Fig. S15. SPN03 simulation.** (A) Sampling scheme showing the four longitudinal samples collected at distinct evolutionary time points. (B) Muller plot of the simulated evolutionary dynamics, with sampling times and driver mutations annotated. (C) Allele-specific copy-number profile of chromosome 13q in SPN03_1.1, showing the ancestral 1:0 deletion pattern in BAF, depth ratio, and VAF. (D) Multivariate VAF analysis across all samples, resolving clonal and subclonal structure with key driver mutations highlighted.

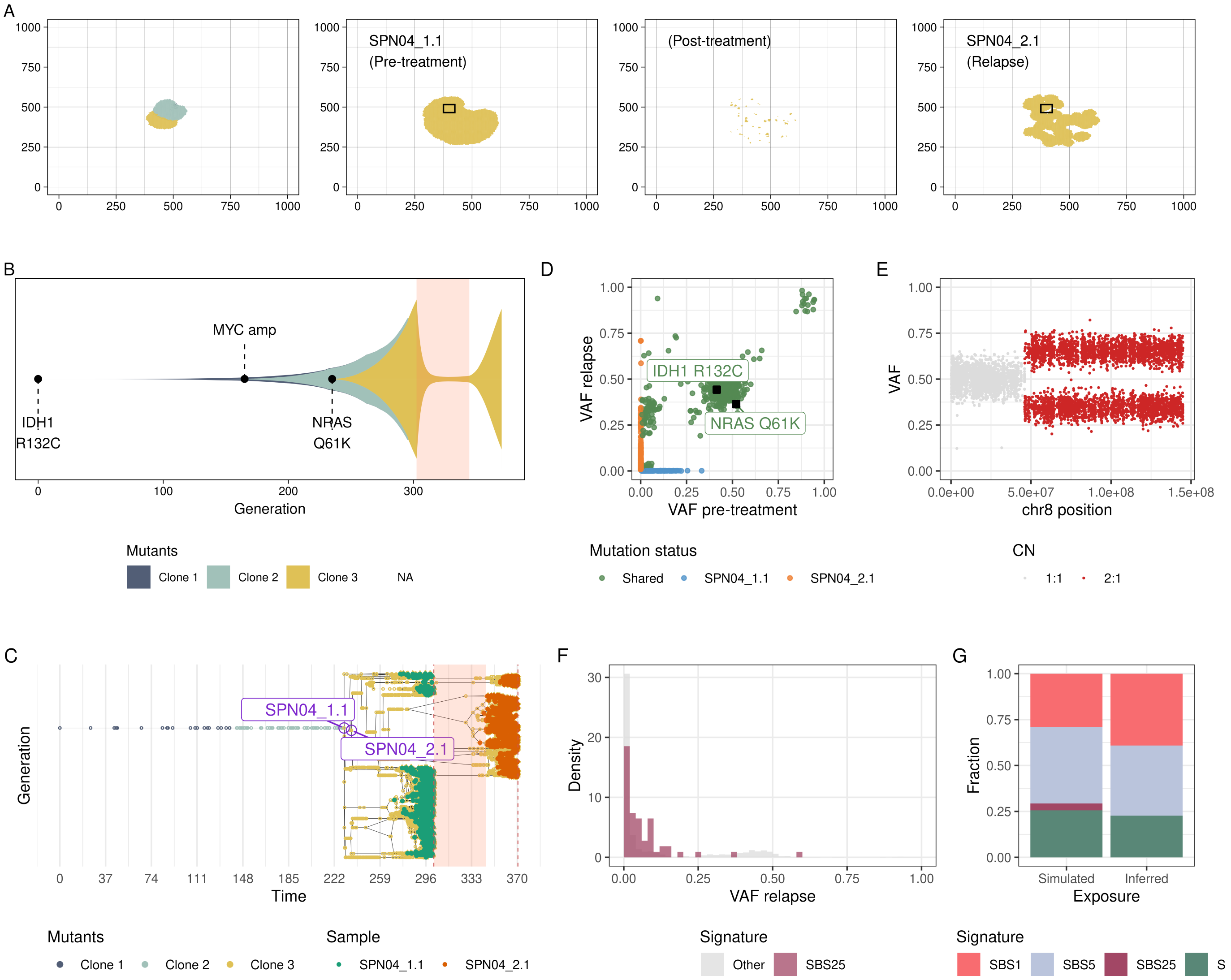

**Supplementary Fig. S16. SPN04 simulation.** (A) Sampling scheme showing the two longitudinal samples collected at pre-treatment and relapse time points, with tissue structure illustrated before and after treatment. (B) Muller plot of the simulated evolutionary dynamics, with the treatment period indicated by a shaded region. (C) Sample forest with the most recent common ancestor (MRCA) of the sampled cells annotated. (D) Multivariate VAF analysis across all samples, resolving shared and private mutations with driver mutations highlighted. (E) Multivariate VAF highlighting SBS25-associated (treatment-induced) mutations, present at low frequency in the relapse sample. (F) Allele-specific copy-number profile along chromosome 8, illustrating the clonal MYC amplification. (G) Univariate VAF distribution of the relapse sample with SBS25-associated mutations highlighted.

In parallel, analysis of mutational contexts reveals a highly stable mutational spectrum across samples, consistent with a persistent mismatch repair-deficient hypermutator state. In particular, SBS3 and ID7 profiles remain remarkably consistent across all biopsies, indicating that the underlying mutational processes are stable throughout tumour evolution despite substantial variation in clonal composition.

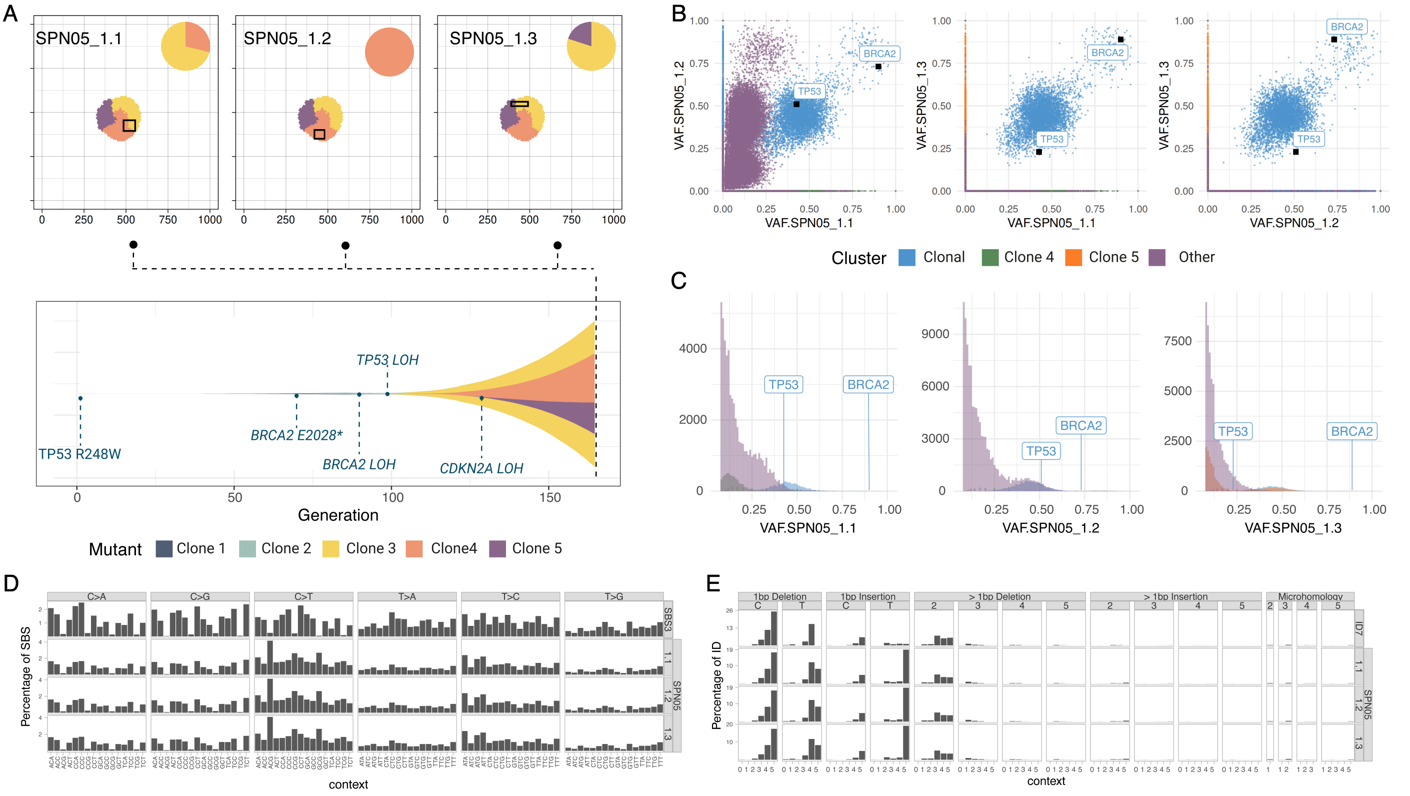

**Supplementary Fig. S17. SPN05 simulation.** (A) Sampling scheme showing the three spatially distinct tumour regions profiled at a single time point, with driver mutations annotated. (B) Muller plot of the simulated evolutionary dynamics, with sampling time and driver mutations annotated. (C) Multivariate VAF analysis across all samples, resolving clonal and subclonal structure with key driver mutations highlighted. (D) Univariate VAF distributions for all three samples with driver mutations annotated. (E) SBS mutational profile for SBS3, associated with homologous recombination deficiency, across the three samples. (F) ID mutational profile for ID7, associated with mismatch repair deficiency, across the three samples.

##### *B.3.8. SPN06*

SPN06 is a prototype smoking-associated lung adenocarcinoma [38] comprising seven clones that undergo dynamic competition during two lines of treatment and subsequent relapses driven by de novo resistance mechanisms (Supplementary Fig. S18). Tumor evolution begins with the acquisition of a TP53 R248W mutation (clone C1), followed by a loss-of-heterozygosity event affecting STK11 (clone C2).

At the time of the first treatment, the tumour exhibits a polyclonal architecture generated by two independent subclonal expansions arising from clone C2. These expansions are driven by distinct driver alterations: an EGFR amplification (clone C3) and a KEAP1 R413H mutation (clone C4). The first line of therapy consists of platinum-based chemo-immunotherapy, which reduces tumour burden. During this treatment window, clones C2, C3, and C4 are eradicated, whereas a KRAS G12D-driven subclone (clone C5), also derived from C2, survives therapy and drives the first relapse.

Chemotherapy induces the mutational signature SBS25, which is active only during its administration. Additional mutational processes, including the clock-like signatures and the smoking-associated SBS4, remain active throughout the evolutionary history of the tumour. Importantly, both the mutation rate and the copy-number alteration (CNA) rate remain constant over time and are not modified by therapeutic interventions.

Clone C5 continues to expand until the second line of chemotherapy, which does not induce any therapy-specific mutational signature. During this phase, an amplification of the KRAS mutant allele emerges (clone C6), enabling resistance and leading to a second relapse. A final subclonal expansion (clone C7) arises following a whole-genome doubling (WGD) event, which occurs late in tumour evolution and remains subclonal at the final sampling time.

SPN06 is profiled using a longitudinal, multi-region sampling strategy designed to assess the impact of therapy on tumour evolution. Two samples collected before the first treatment display polyclonal compositions: SPN06_1.1 contains clones C2 and C3, SPN06_1.2 includes C2, C3, and C4, while SPN06_1.3 is also polyclonal with varying proportions of these lineages. A third sample, SPN06_2.1, obtained prior to the second treatment, reveals a monoclonal population composed exclusively of C5. At the time of the second relapse, two additional spatially distinct samples, SPN06_3.1 and SPN06_3.2, show that clone C6 has survived therapy and that the late-arising WGD-driven clone C7 is present as a subclonal population.

Clonal inference based on multivariate VAF analysis shows substantial variability across samples. A major analytical limitation arises from spatial confounders combined with a high degree of similarity in mutational processes across subclonal populations, which reduces the separability of their genomic profiles and complicates robust reconstruction of the underlying clonal structure from bulk data. The presence of whole-genome doubling further exacerbates these difficulties by distorting copy-number profiles and clonal proportions, making subclonal WGD events especially difficult to reliably resolve.

Within an agnostic genomic analysis, both the KRAS G12D mutation and the subsequent amplification of the KRAS mutant allele would emerge as plausible candidate resistance drivers. Indeed, each alteration becomes clonal following a major evolutionary bottleneck, with the KRAS mutation defining the dominant population at the first relapse and the KRAS amplification defining the dominant population at the second relapse. From an observational perspective, such patterns would naturally nominate both events as candidate mediators of genetic resistance. However, the two cases differ in the information provided by mutational signatures. In the first relapse, the co-occurrence of the therapy-associated signature SBS25 would support the interpretation that the resistant clone emerged de-novo during treatment. In contrast, the second relapse lacks a corresponding treatment-associated signature. As a result, although KRAS amplification would still represent a strong candidate resistance event, standard mutational signature analysis alone would not allow one to determine whether the amplified clone emerged during therapy or originated from a pre-existing resistant population. Discriminating between these alternatives would require dedicated timing-aware evolutionary analyses beyond conventional mutational signature deconvolution.

##### *B.3.9. SPN07*

SPN07 is a prototype of glioblastoma comprising six clones that compete over time during a single line of treatment and a subsequent relapse driven by a de novo genetic resistance mechanism [39–42]. Tumour expansion is initially driven by PTEN inactivation, beginning with a PTEN R130G mutation (clone C1) followed by loss of heterozygosity of the remaining wild-type allele (clone C2). At the time of treatment, the tumour exhibits a polyclonal structure, with two subclonal lineages independently derived from C2 through an NF1 R192* mutation (clone C3) and an ATRX R907* mutation (clone C4).

Temozolomide chemotherapy selects for a hypermutant subclone (clone C5) initiated by an MSH6 R361H mutation acquired during treatment. This mutation in glioblastoma is strongly associated with acquired resistance to temozolomide chemotherapy, as it disrupts the mismatch repair pathway and enables survival under treatment-induced DNA damage. This alteration substantially increases the single-nucleotide variant mutation rate in C5. Prior to therapy, only clock-like mutational processes are active. Temozolomide exposure induces the characteristic therapy-associated signature SBS11, which becomes clonally established within the resistant population. Following acquisition of the MSH6 mutation, additional hypermutation-related signatures SBS25 and SBS26 become detectable within subclonal fractions. Within the relapse population, a further subclone emerges through acquisition of a TP53 R248W mutation (clone C6).

SPN07 is profiled using a longitudinal, multi-region sampling strategy designed to capture clonal dynamics and treatment impact. Three samples are collected prior to temozolomide administration and two additional samples at relapse. Sample composition across SPN07_1.1, SPN07_1.2, and SPN07_1.3 reflects the coexistence and shifting proportions of multiple subclonal lineages, while relapse samples SPN07_2.1 and SPN07_2.2 are each dominated by single resistant clones, together demonstrating a structured relapse driven by a genetically distinct resistant lineage.

In this setting, the multivariate VAF distribution reveals the polyclonal architecture and clonal competition throughout tumour history. The complexity of SPN07 poses several challenges for downstream analysis. First, driver mutation calling is strongly affected by the hypermutated state. The elevated mutation rate increases the probability of identifying recurrent mutations in known cancer genes that do not correspond to true selective events in the simulated evolutionary history, effectively generating false drivers. Second, subclonal deconvolution is complicated by spatial sampling effects within each time point, which induce apparent cluster structure that does not reflect true evolutionary branching and can lead to misclassification of mutation clusters. Third, mutational signature deconvolution is affected by both hypermutation and subclonal structure. Signatures such as SBS25 and SBS26 are often confined to subclonal populations and are therefore difficult to resolve, especially at lower purity or coverage, while detection of the treatment-associated SBS11 signature can be challenging at low variant allele frequency despite its clonal nature in the resistant population.

Further information regarding the relapse mechanism is obtained from mutational signature analysis at the cluster level of the marginal VAF distributions of samples SPN07_1.3 and SPN07_2.2. Sample SPN07_1.3 shows clock-like mutations in both clonal and tail clusters, whereas the clonal cluster in SPN07_2.2 is dominated by SBS11 mutations. This pattern is consistent with an MSH6-driven relapse, where exposure to SBS11 is triggered by acquisition of the alteration. Moreover, MSH6 deficiency is known to promote a hypermutant phenotype characterized by an increased mutation rate. The comparison of mutational burden between SPN07_1.3 and SPN07_2.2 is consistent with this mechanism, particularly in tail mutation clusters, where the number of mutations reflects the underlying mutation rate.

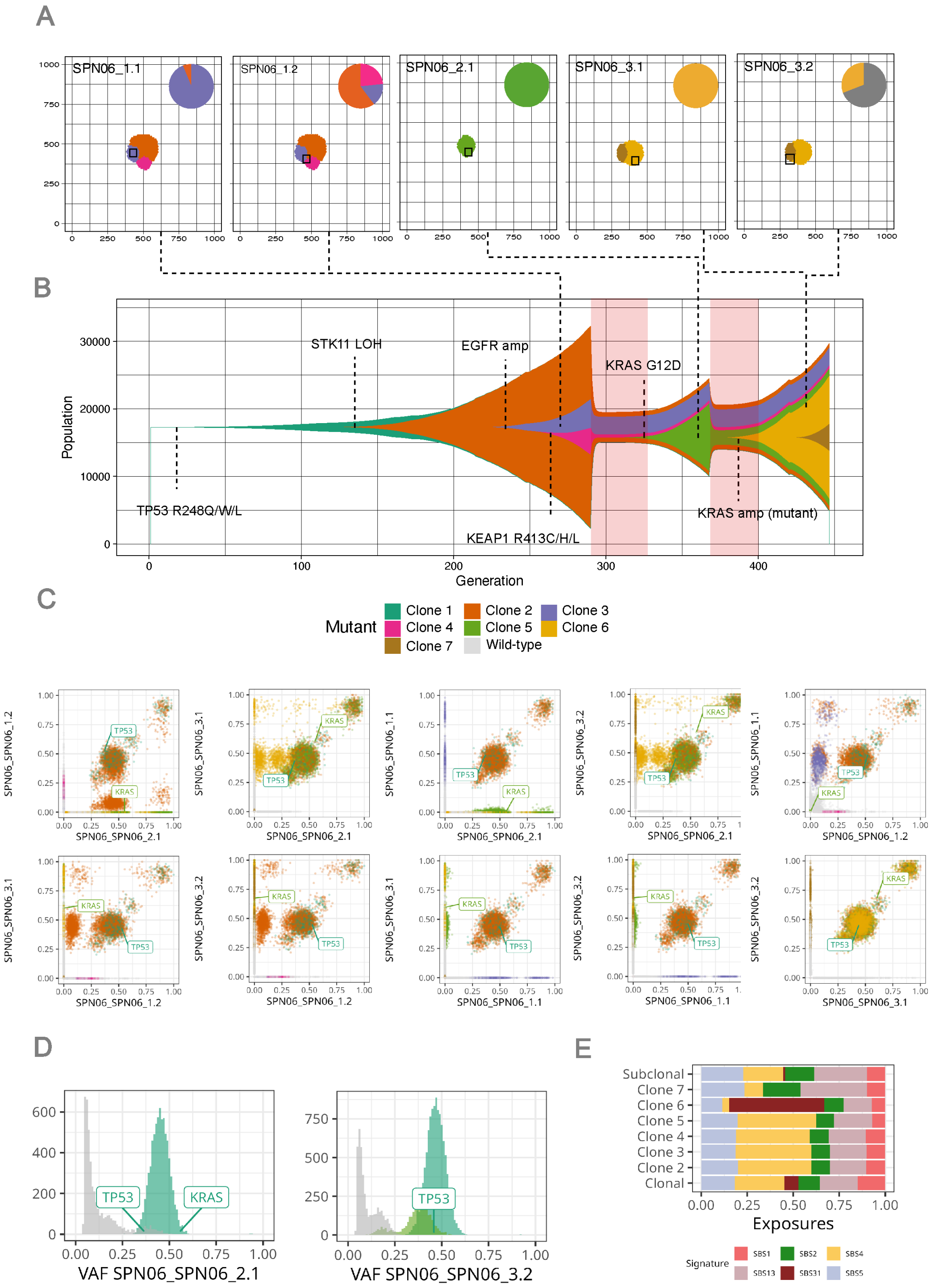

**Supplementary Fig. S18. SPN06 simulation.** (A) Sampling scheme showing the five samples collected at three time points: two pre-treatment, one at first relapse, and two at second relapse. (B) Muller plot of the simulated evolutionary dynamics, with treatment intervals highlighted by shaded regions and driver mutations annotated. (C) Multivariate VAF analysis across all samples, resolving shared and private mutations with driver mutations highlighted. (D) Univariate VAF distributions for the two relapse samples SPN06_2.1 (after first-line chemo-immunotherapy) and SPN06_3.2 (after second-line chemotherapy). (E) Simulated mutational signature exposures across clones, illustrating the emergence of the treatment-associated SBS25 signature during first-line therapy.

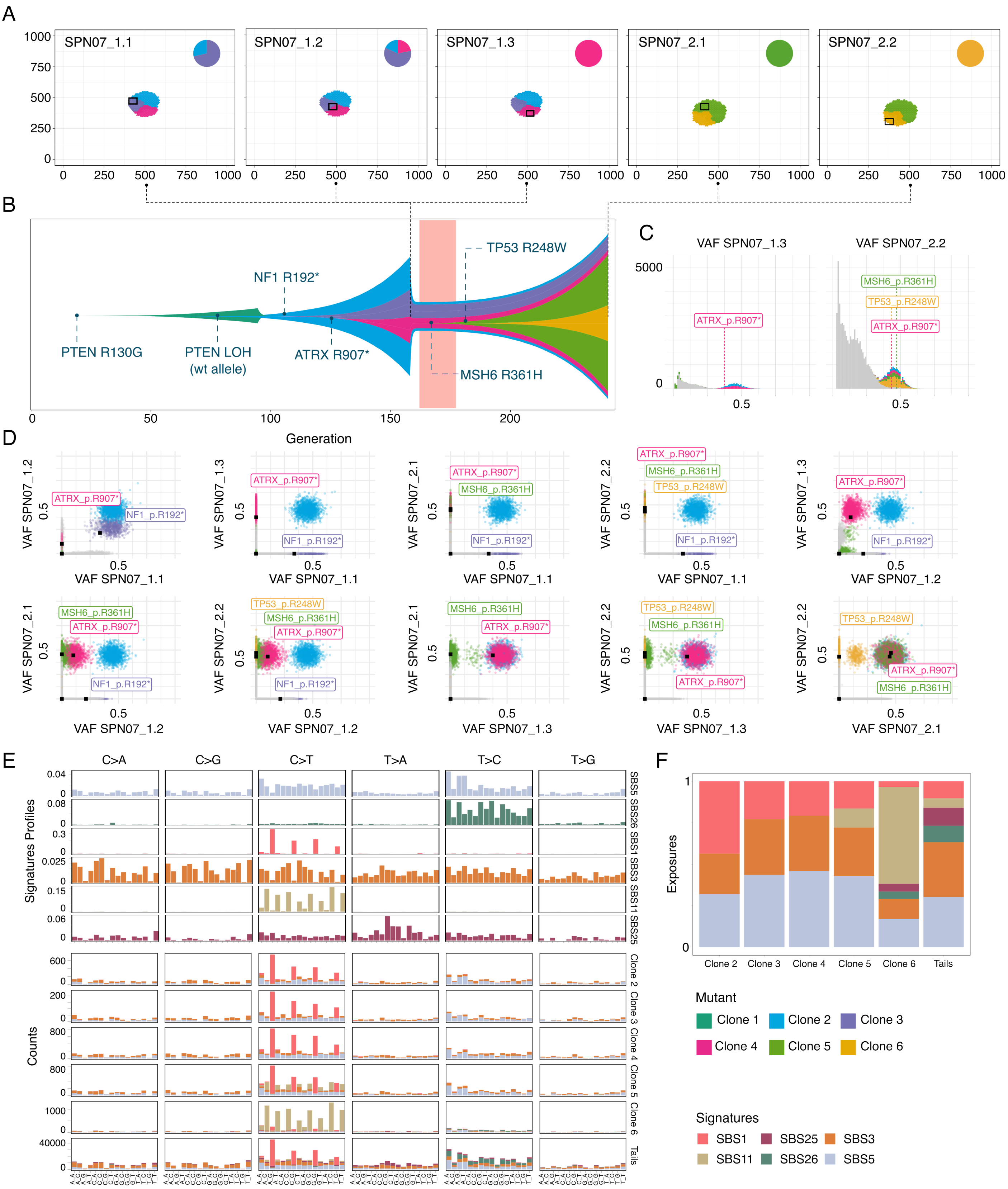

**Supplementary Fig. S19. SPN07 simulation.** (A) Sampling scheme showing the five samples collected at two time points: three pre-treatment and two at relapse, with tissue structure illustrated before and after temozolomide (TMZ) treatment. (B) Muller plot of the simulated evolutionary dynamics, with the TMZ treatment period indicated by a shaded region and driver mutations annotated. (C) Multivariate VAF analysis across all samples, resolving shared and private mutations with driver mutations highlighted. (D) Univariate VAF distributions for a pre-treatment sample (SPN07_1.3) and a relapse sample (SPN07_2.2), illustrating the markedly higher mutation burden in the hypermutant relapse clone. (E) SBS mutational profiles across clones, showing clock-like signatures (SBS1, SBS5), hypermutation-related signatures (SBS25, SBS26), and the TMZ-induced signature (SBS11), with mutation counts by context coloured according to simulated aetiology. (F) Simulated mutational signature exposures across all samples.

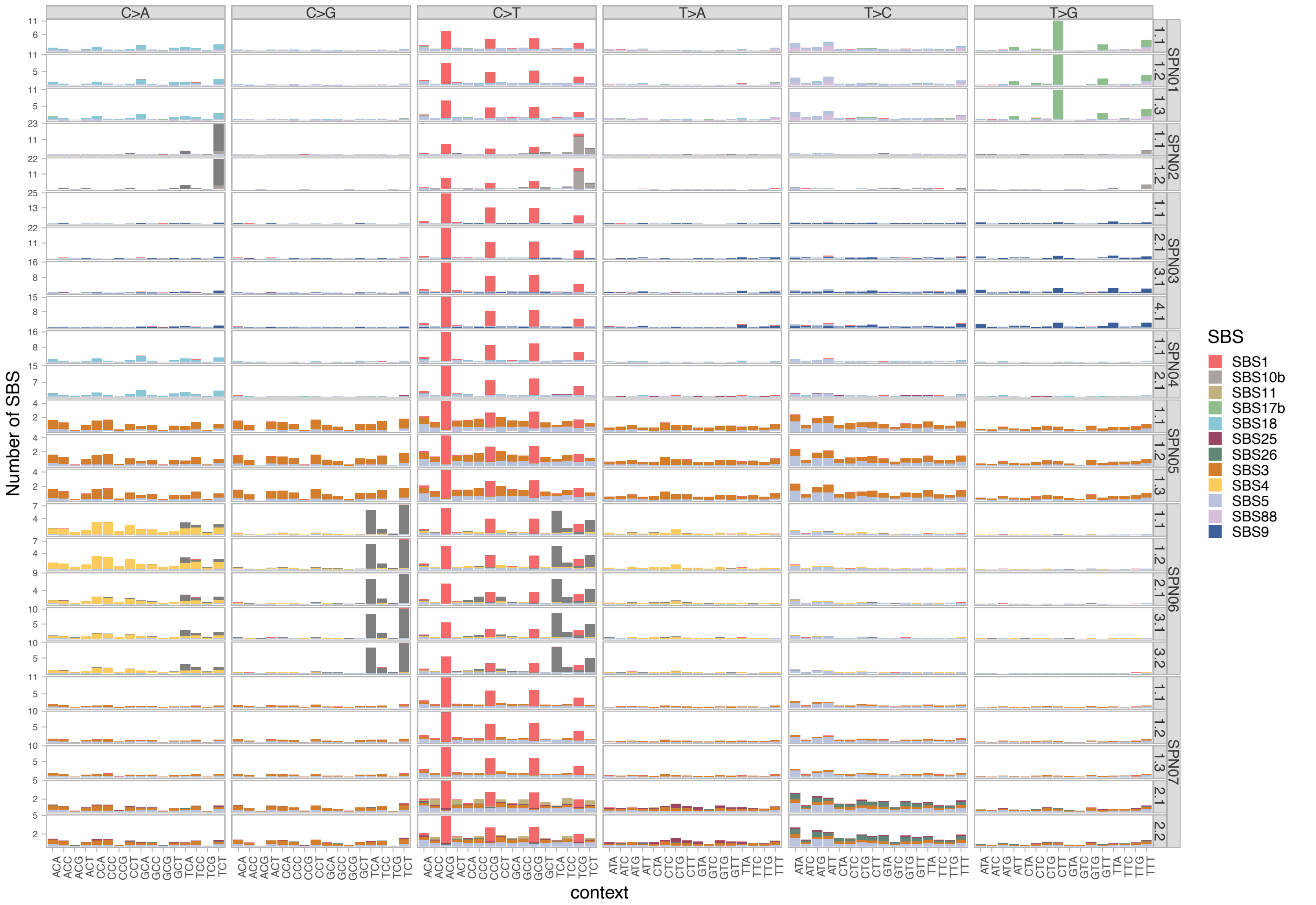

**Supplementary Fig. S20. SBS profile for SCOUT cohort.** SBS signature profile for each of the sample composing the SCOUT cohort.

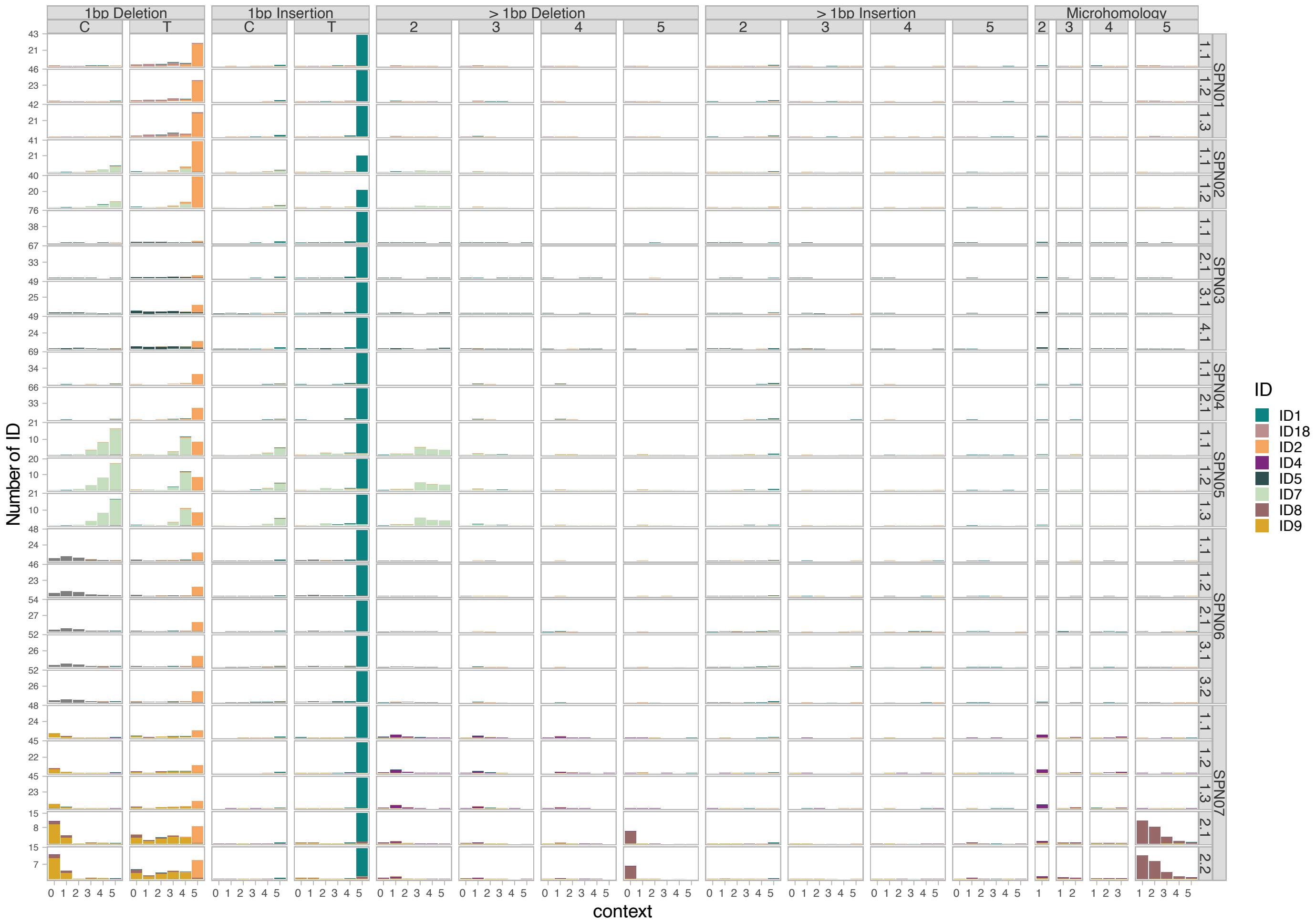

**Supplementary Fig. S21. ID profile for SCOUT cohort.** ID signature profile for each of the sample composing the SCOUT cohort.

#### C. Benchmarking of the SCOUT cohort with nf-core/sarek

We benchmarked the nf-core/sarek pipeline by comparing somatic and germline variant and copy-number calls against the molecular ground truth provided by the ProCESS simulations. Results are organised in two parts: a cohort-level analysis summarising performance trends across all SPNs, purity conditions, and coverage levels, followed by per-SPN results.

##### C.1. Cohort results

###### *C.1.1. Somatic variant callers*

We evaluated the performance of three somatic variant callers, Strelka2 [43], FreeBayes [44], and Mutect2 [45], across simulated tumours generated at varying purity and coverage levels, using the matched tumour-normal whole-genome sequencing data produced by ProCESS.

A variant was considered truly somatic in the simulated BAMs when its tumour VAF was ≥ 0.02. For Strelka2 and Mutect2, somatic calls corresponded to variants marked `PASS` in their somatic VCFs. For FreeBayes, which lacks explicit somatic calling, we classified variants as somatic if they met the following empirical thresholds: quality ≥ 20, tumour VAF ≥ 0.01, normal VAF ≤ 0.02, and depth ≥ 10. Based on these definitions, we derived precision, recall, accuracy and False Discovery Rate (FDR).

To assess the impact of tumour subclonality, we computed these metrics across cancer cell fraction (CCF) bins spanning 0–5%, 5–10%, 10–25%, 25–50%, 50–75%, 75–95%, and clonal (CCF ≥ 95%), enabling detailed comparison of caller performance across the full spectrum of allelic representation. For visualisation, bins were further aggregated into a subclonal low-frequency group (CCF < 10%), a subclonal high-frequency group (CCF 10–95%), and a clonal group. All metrics were computed separately for SNVs and INDELs across all combinations of tumour purity and sequencing coverage.

Across the cohort, clonal SNV recall was high and similar for Strelka2 (0.922) and Mutect2 (0.921), with FreeBayes slightly lower (0.910). All three callers showed a strong dependence on tumour purity: at purity 0.3, recall dropped to 0.890, 0.892, and 0.862 for Strelka2, Mutect2, and FreeBayes respectively, recovering to 0.943, 0.942, and 0.939 at purity 0.9. FDR was lowest for Strelka2 (cohort mean 0.044), while Mutect2 (0.109) and FreeBayes (0.107) showed higher false-discovery rates, driven primarily by hypermutant samples where the elevated mutation burden increases the probability of passenger mutations passing somatic filters. Recall of subclonal variants at high frequency (CCF 10–95%) was substantially lower than recall of clonal variants for all three tools: Strelka2 0.711, Mutect2 0.668, FreeBayes 0.455. The drop was most severe in the low-CCF bin (CCF < 10%), where recall fell to 0.171, 0.160, and 0.091 for Strelka2, Mutect2, and FreeBayes respectively, reflecting the fundamental difficulty of distinguishing rare variants from sequencing noise. Among the callers, Mutect2 showed the highest sensitivity for INDELs while Strelka2 performed best for clonal SNVs, and FreeBayes displayed the lowest subrecall of clonal variants across all CCF bins. A full summary of recall and FDR per SPN, CCF bin, purity, and coverage is reported in Supplementary Fig. S22 and Supplementary Table ST9A.

###### *C.1.2. Germline variant callers*

We evaluated the performance of normal germline variant detection by benchmarking three callers, Strelka [43], FreeBayes [44], and HaplotypeCaller [46], against the known germline mutations included as ground truth in the simulated data. For Strelka and HaplotypeCaller, we considered variants as true calls if their FILTER field was marked as `PASS`. Since FreeBayes does not use a FILTER field, we treated variants with a quality score ≥ 20 as passing. Precision, recall, and accuracy were computed. To assess the accuracy of B-allele frequency (BAF) estimation, we additionally computed the Pearson correlation coefficient and root mean squared error (RMSE) between predicted and true BAFs, considering only true positive variants (i.e., those detected by the caller and present in the ground truth). Unlike the somatic analysis, CCF binning was not applied, as germline variants are by definition present in all cells of the normal at a diploid allele frequency.

All three callers achieved high F1 scores for germline SNV detection across all SPNs and conditions. Performance differences were most evident in BAF estimation accuracy: Strelka achieved the highest mean Pearson correlation between inferred and true BAF (0.906, RMSE 0.089), followed closely by HaplotypeCaller (0.897, RMSE 0.090) and FreeBayes (0.895, RMSE 0.092). The small absolute differences reflect the fact that all three tools accurately recover heterozygous germline variants; the ranking is driven by the tails of the BAF distribution where FreeBayes estimates are more variable. Full per-SPN results are reported in Supplementary Fig. S23 and Supplementary Table ST9B.

###### *C.1.3. Copy number callers*

We evaluated the accuracy of copy number (CN) inference using three tools: ASCAT [47], Sequenza [48], and Battenberg [49]. ASCAT and Sequenza support allele-specific CN estimation, while Battenberg additionally resolves subclonal CN states. To account for the heterogeneity in copy-number complexity across simulated tumours, samples were stratified by the fraction of genome altered (FGA), defined as the proportion of the autosomal genome deviating from a diploid state:

$$\mathrm{FGA} = \frac{L_{\mathrm{altered}}}{L_{\mathrm{total}}} \times 100 \tag{24}$$

where $L_{\mathrm{altered}}$ is the total length of segments with copy number changes (CN ≠ 1 : 1 or ratio < 1), and $L_{\mathrm{total}}$ is the total length of autosomal segments. Samples with FGA > 15% were classified as High FGA; all remaining samples were designated Low FGA. All evaluations were performed separately within each category.

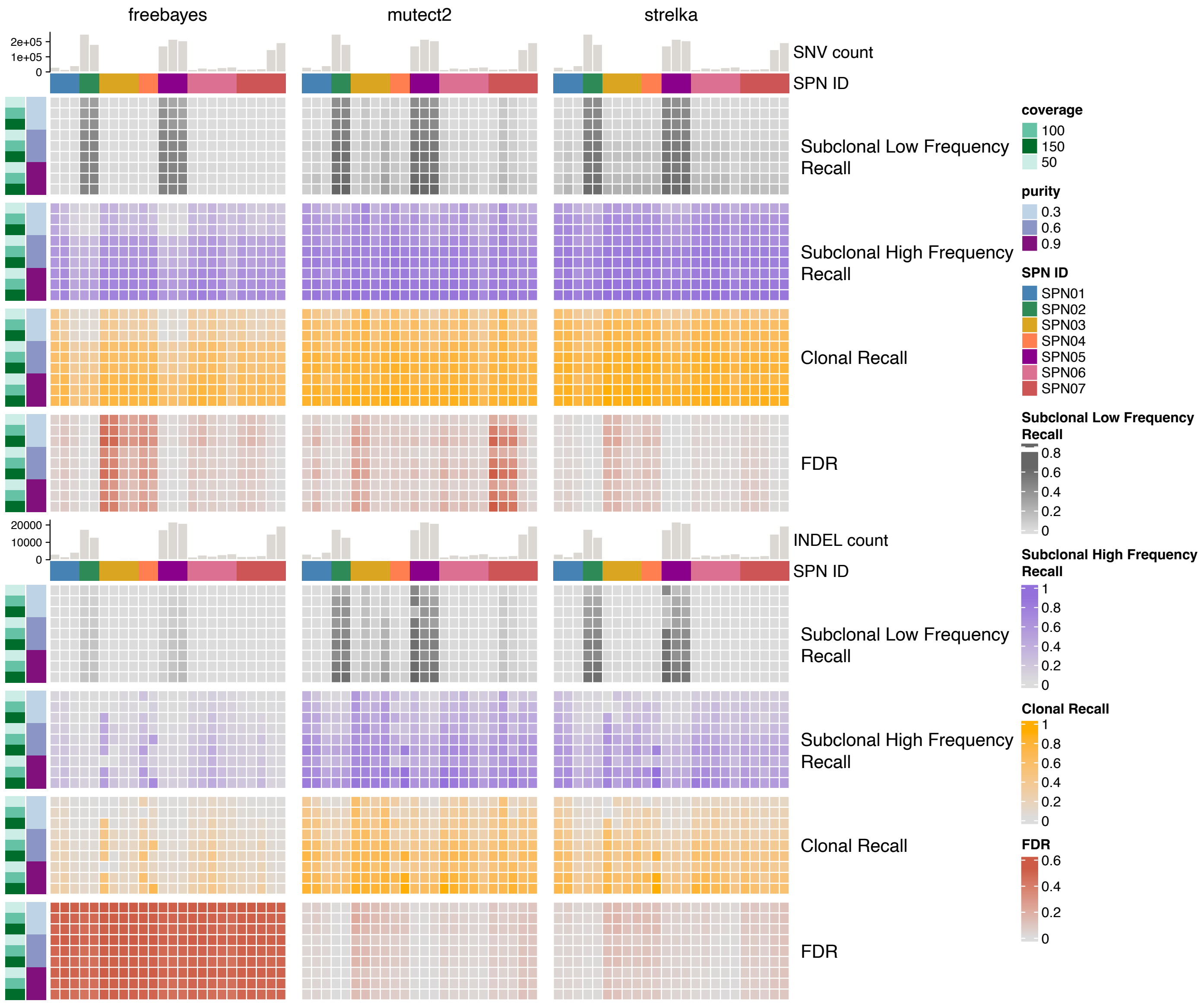

**Supplementary Fig. S22. Overview of SCOUT somatic benchmarking**. Heatmaps showing FreeBayes, Mutect2, and Strelka performance, recall and FDR, for each SPN sample across simulated tumour purity (0.3, 0.6, 0.9) and coverage (50×, 100×, 150×). SNV count per samples is also reported.

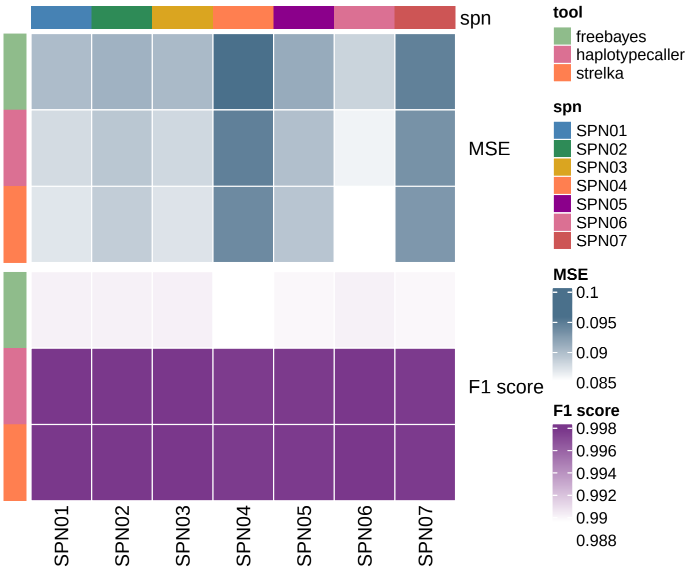

**Supplementary Fig. S23. Overview of SCOUT germline benchmarking**. Heatmaps showing FreeBayes, Strelka, and HaplotypeCaller performance for each SPN in term of Root Mean Squared Error (RMSE) and F1 score.

We first assessed purity and ploidy estimates from each caller by computing the absolute error between the predicted and simulated ground-truth values ($\delta_\pi$ for purity and $\delta\sigma$ for ploidy). We then evaluated breakpoint detection accuracy and CN state inference. Because ProCESS simulations contain both clonal and subclonal CN segments, the latter defined by a cancer cell fraction CCF < 1, segments were categorised accordingly prior to evaluation. Breakpoint detection was assessed using a tolerance-based matching procedure with a window of $\delta = 10$ kb: predicted breakpoints matching a true breakpoint within $\delta$ were counted as true positives (TP), unmatched predictions as false positives (FP), and unmatched true breakpoints as false negatives (FN), from which precision and recall were derived. For CN state accuracy, we computed the proportion of the clonal genome correctly inferred ($\rho_{\text{clonal}}$) at base-pair resolution:

$$\rho_{\text{clonal}} = 1 - \frac{\sum_{s \in S_{\text{clonal, incorrect}}} \text{length}(s)}{\sum_{s \in S_{\text{clonal}}} \text{length}(s)} \tag{25}$$

An analogous evaluation was performed for subclonal segments. Samples were further stratified by the fraction of genome subclonal (FGS), defined as:

$$\text{FGS} = \frac{L_{\text{subclonal}}}{L_{\text{total}}} \times 100 \tag{26}$$

where $L_{\text{subclonal}}$ is the total length of subclonal segments.

Purity estimation accuracy differed markedly by tool and FGA class. ASCAT showed the most consistent purity errors across FGA classes (Low FGA: 0.043, High FGA: 0.059) and improved substantially at higher purity (errors of 0.055, 0.070, and 0.026 at purity 0.3, 0.6, and 0.9 respectively), reflecting its optimisation for the clonal peak. Battenberg and Sequenza showed larger purity errors in Low FGA samples (0.170 and 0.161 respectively) but performed better on High FGA samples (0.024 and 0.039), where the richer allelic imbalance pattern provides stronger signal for their more flexible models. $\rho_{\text{clonal}}$ showed a pronounced FGA dependence: ASCAT achieved near-perfect $\rho_{\text{clonal}}$ in Low FGA samples (0.998) but dropped to 0.619 in High FGA samples; conversely, Battenberg reached $\rho_{\text{clonal}} = 0.878$ in High FGA samples, its best regime, compared to 0.762 in Low FGA. Sequenza maintained high $\rho_{\text{clonal}}$ in Low FGA samples (0.962) with a moderate drop in High FGA (0.809). Breakpoint precision was generally high across tools and FGA classes (ASCAT 0.815–0.849, Battenberg 0.738–0.756, Sequenza 0.494–0.676), while breakpoint recall varied more substantially and declined in High FGA samples for Battenberg (0.623) and FreeBayes in general, as expected given the greater segmentation complexity. Full results, stratified by FGA class, purity, coverage, and SPN, are reported in Supplementary Fig. S24 and Supplementary Table ST9C.

##### C.2. Per-SPN results

The following subsections report detailed benchmarking results for each SPN in the SCOUT cohort, integrating somatic variant calling, germline variant calling, and copy-number inference.

###### *C.2.1. SPN01*

SPN01 comprises three multi-region samples collected at a single time-point, with distinct copy-number profiles: sample 1.2 is mostly diploid heterozygous (FGA 20%, ploidy 2.1), while 1.1 and 1.3 are almost completely tetraploid (FGA 100%, ploidy 3.0 and 4.1 respectively), the latter arising from a whole-genome doubling (WGD) event. The resulting heterogeneity in ploidy was the primary driver of variable performance in copy number calling, across samples. In particular, 1.2 was well handled by all callers: ASCAT provided the most accurate purity estimates and very small ploidy errors, with Battenberg and Sequenza achieving similar accuracy in purity (mean absolute errors $\delta\pi = 0.005$ and 0.041) and a lower proportion of correctly inferred clonal CNAs (mean absolute errors $\rho_{clonal} = 0.999$ and 0.953). The two tetraploid samples were substantially harder: in 1.1 all callers showed average purity errors in range $\delta\pi \in 0.058$–$0.061$ and lower proportions of correctly inferred clonal CNAs; in 1.3 these errors further increased ($\delta\pi \in 0.102$–$0.125$) and the proportion of correctly inferred clonal CNAs collapsed to $\rho_{clonal} = 0$ for Battenberg and Sequenza, making SPN01 overall the most demanding case in the cohort for CNA inference. In general, all CNA callers struggled to correctly infer ploidy in highly aneuploid samples, and tended to underestimate it, e.g. 1.3 instead of $\approx 4.12$ and 1.1 instead of 3.0. This bias arose most likely because these methods are not designed to detect WGD.

On the other hand, somatic variant calling was uniformly high for all three samples and across callers in terms of recall of clonal variants (Strelka2: 0.921–0.928, Mutect2: 0.918–0.926, FreeBayes: 0.907–0.918); False discovery rates FDR were also low, with the largest values observed in the diploid sample (1.2, Mutect2: 0.117), where the mixed subclonal architecture generated more ambiguous calls. Recall of subclonal variants with relatively high-frequency was lower for the tetraploid sample 1.3 across all callers (Strelka2: 0.630, Mutect2: 0.568, FreeBayes: 0.294), reflecting the difficulty of resolving subclonal variants in the contex of a complex copy number profile. Germline VAF correlation in this sample was also slightly lower than in simpler SPNs (Strelka: 0.902, HaplotypeCaller: 0.866, FreeBayes: 0.863).

###### *C.2.2. SPN02*

SPN02 comprises two spatially distinct biopsies collected at a single time-point (1.1, 1.2), both almost completely diploid with low-FGA (FGA 2% and 7%, ploidy 2.0) and harbouring a POLE-driven hypermutator phenotype that generates approximately

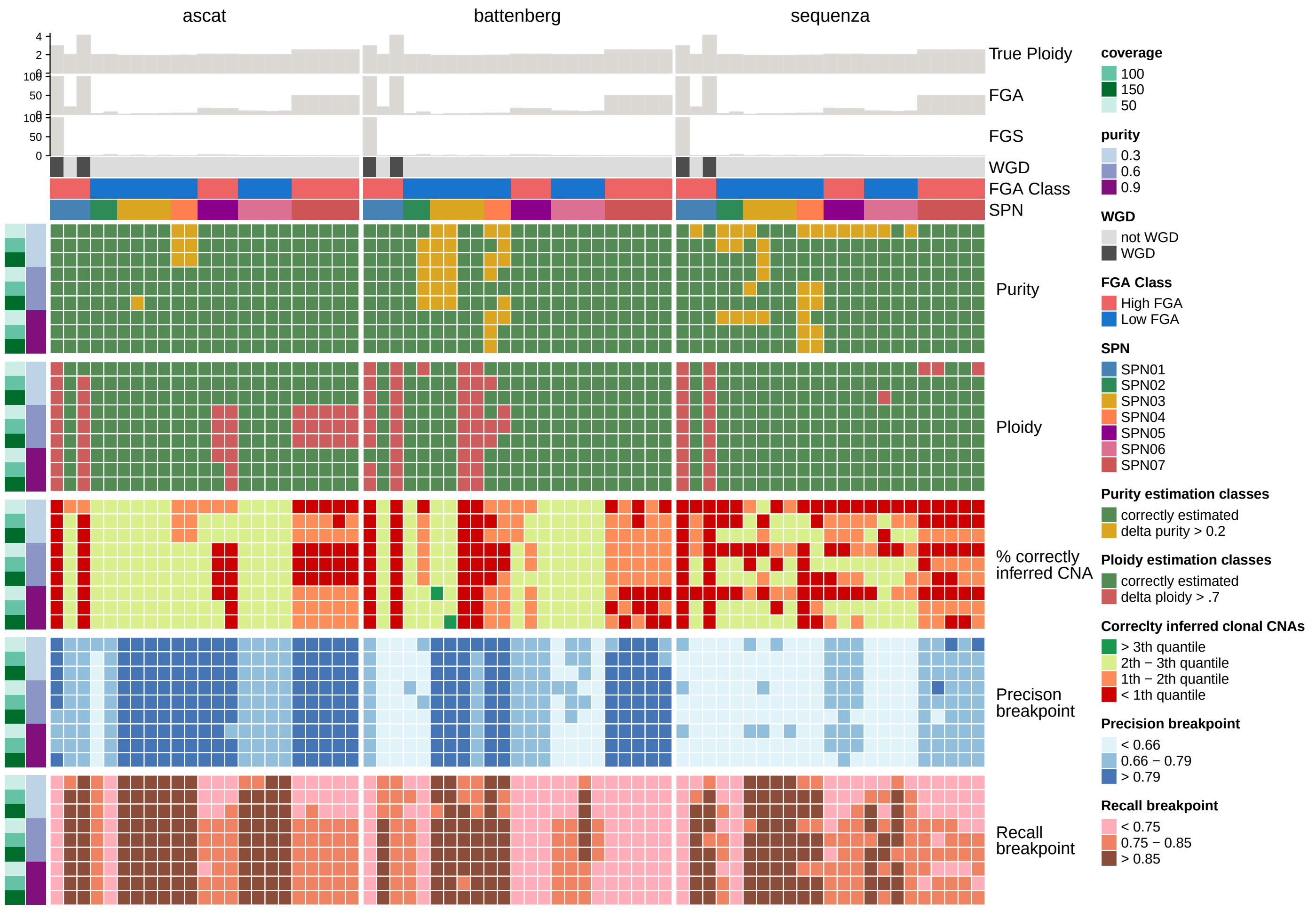

**Supplementary Fig. S24. Overview of SCOUT CNA benchmarking**. Heatmap showing ASCAT, Sequenza, and Battenberg performance for each SPN sample across simulated tumour purity (0.3, 0.6, 0.9) and coverage (50×, 100×, 150×). True ploidy, FGA and FGA for each samples are reported on top of the heatmap.

514 000 and 413 000 SNVs for each sample, respectively. The two samples are structurally equivalent in terms of copy-number simplicity and clonal composition, and their somatic variant calling performance is correspondingly similar: recall of clonal variants was relatively large (Strelka2: 0.922–0.924, Mutect2: 0.918–0.920, FreeBayes: 0.907–0.910), with very low FDR for Strelka2 (≤ 0.003) and FreeBayes (≤ 0.010) across both samples, and moderate FDR for Mutect2 (0.038–0.040). Recall of subclonal, high-frequency mutations was notably lower for FreeBayes (0.301 in both samples) compared to Strelka2 (0.617–0.641) and Mutect2 (0.649–0.663). Germline VAF correlation was uniformly high (Strelka: 0.941, HaplotypeCaller: 0.941, FreeBayes: 0.939). Concerning CNA inference, ASCAT performed well on both samples, with purity errors of 0.016 (1.1) and 0.010 (1.2), and correctly inferring all clonal CNAs ($\rho_{clonal} = 1$). Battenberg was more accurate on 1.1 ($\delta\pi = 0.006$, $\rho_{clonal} = 0.998$), but showed a larger error on 1.2 ($\delta\pi = 0.306$, $\rho_{clonal} = 0.866$), suggesting a substantial sensitivity to spatial mutation patterning in the second biopsy. Sequenza showed consistent purity errors around $\delta\pi \in 0.211$–$0.221$ and large proportions of correctly inferred clonal CNAs ($\rho_{clonal} \in 0.915$–$0.934$) in both samples.

##### C.2.3. SPN03

SPN03 comprises four longitudinal samples (1.1 to 4.1), all with low-FGA and almost completely diploid (FGA 0.3–2.7%, ploidy 1.9) genomes. Low absolute SNV counts (4 000–11 000) constituted the defining challenge of this SPN. Clonal SNV recall was comparable across all four time points and callers (Strelka2: 0.920–0.923, Mutect2: 0.917–0.920, FreeBayes: 0.906–0.910), but the FDR was large relative to other non-hypermutant SPNs with higher mutation counts, because false positives contribute disproportionately at low burden. The FDR was higher in the early time points (1.1 and 2.1, FreeBayes: 0.376–0.410, Strelka2: 0.155–0.168) and lower in the later samples (3.1 and 4.1, FreeBayes: 0.189–0.203, Strelka2: 0.070–0.071), tracking the increase in clonal SNV count accompanying the dominant clonal expansion. Recall of subclonal variants at high frequency was consistently high for Strelka2 (0.755–0.790) and Mutect2 (0.644–0.762) across all time points, and lower for FreeBayes (0.526–0.586). Germline VAF correlation was uniform across all callers (∼ 0.864–0.867). For CNA inference, ASCAT performed consistently well across all four time points with near-zero purity errors ($\delta\pi \in 0.001$–$0.057$) and full proportion of correctly inferred clonal CNAs ($\rho_{clonal} = 1.000$). It also achieved the best breakpoint performance in the cohort (precision 0.942–0.988, recall 0.988). Battenberg showed a variable behaviour: a good proportion of correctly inferred clonal CNAs in 1.1 and 2.1 ($\rho_{clonal} = 0.997$) but complete failure on 3.1 and 4.1 ($\rho_{clonal} = 0.000$), with consistently large purity errors ($\delta\pi \in 0.119$–$0.369$). Sequenza maintained good proportion of correctly inferred clonal CNAs ($\rho_{clonal} \in 0.967$–$0.989$) with moderate purity errors ($\delta\pi \in 0.022$–$0.262$).

##### C.2.4. SPN04

SPN04 comprises two longitudinal samples (1.1 pre-treatment, 2.1 at relapse), both low-FGA and near-diploid (FGA 3.5%, ploidy 2.0). Relapse is driven by non-genetic plasticity rather than new somatic mutations, so the two samples share the same clonal architecture. The low SNV count (5 000–6 500 per sample) is here the defining challenge. The two samples showed very similar somatic variant calling performance: recall of clonal variants relatively high (Strelka2: 0.909–0.910, Mutect2: 0.915 (both samples), FreeBayes 0.909 (both samples). The FDR was high for FreeBayes (0.234–0.275) and lower for Strelka2 (0.085–0.102) and Mutect2 (0.060–0.073), consistent with the low absolute mutation count. Recall of subclonal variants at high frequency was relatively lower (Strelka2: 0.727–0.763, Mutect2: 0.677–0.754, FreeBayes: 0.541–0.544). Germline VAF correlation was uniformly high (∼ 0.859–0.861). For CNA inference, performance was identical between the two samples, as expected from their shared copy-number profile: ASCAT achieved near-perfect proportion of correctly inferred clonal CNAs ($\rho_{clonal} = 0.988$) and breakpoint performance (precision 0.983, recall 1.000) on both samples, with purity errors of $\delta\pi \in 0.203$–$0.210$ that were inflated by failures at low purity/coverage conditions rather than reflecting systematic bias. Battenberg showed large purity errors ($\delta\pi \in 0.354$–$0.371$) and intermediate proportions of correctly inferred clonal CNAs ($\rho_{clonal} \in 0.536$–$0.750$). Sequenza achieved high proportion of correctly inferred clonal CNAs ($\rho_{clonal} \in 0.956$–$0.957$) with large purity errors (0.350–0.414). This was a common behavior of CNA callers, which tended to overestimate purity in nearly diploid samples due to the limited copy-number signal available for accurate model fitting.

##### C.2.5. SPN05

SPN05 comprises three spatially distinct biopsies collected at a single time point. All three samples share the MMR-deficient hypermutator background driven by BRCA2 inactivation, resulting in very high SNV counts (519 000–622 000 per sample) and a uniformly high FGA (∼ 15–16%, ploidy 2.1). Somatic variant calling performance was consistent across all three samples: recall of clonal variants was relatively high (Strelka2: 0.928–0.932, Mutect2: 0.927–0.930, FreeBayes: 0.916–0.919), with FDR very low for Strelka2 (0.003–0.005) and FreeBayes (0.011–0.025), and moderate for Mutect2 (0.042–0.084). Recall of subclonal variants at high frequency was similar across Strelka2 (0.640–0.659) and Mutect2 (0.621–0.697), with FreeBayes substantially lower (0.286–0.322). Germline VAF correlation was uniformly high (Strelka: 0.941, HaplotypeCaller: 0.941, FreeBayes: 0.939). For CNA inference, sample 1.1 was the easiest, with ASCAT and Battenberg both achieving low purity errors ($\delta\pi = 0.004$ and 0.008) and high proportion of correctly inferred clonal CNAs ($\delta\pi = 0.996$ and 0.994); 1.2 and 1.3 were more challenging for ASCAT ($\delta\pi = 0.070$ and 0.083, $\rho_{clonal} = 0.553$ and 0.330), while Battenberg remained accurate in these samples ($\delta\pi = 0.006$ for both samples, $\rho_{clonal} \in 0.976$–$0.994$). Sequenza showed consistent moderate purity errors ($\delta\pi \in 0.044$–$0.050$) and good proportion of correctly inferred clonal CNAs ($\rho_{clonal} \in 0.947$–$0.956$) across all three samples.

###### *C.2.6. SPN06*

SPN06 comprises five samples across three time points: two pre-treatment biopsies (1.1, 1.2), a sample at first relapse (2.1) and two samples at second relapse (3.1, 3.2) with a WGD-driven treatment-resistant subclonal population. All samples, except the last are low-FGA and near-diploid (FGA 8–9%, ploidy 2.0). Somatic variant calling performance was consistent across all samples: recall of clonal variants was relativley high (Strelka2: 0.916–0.933, Mutect2: 0.918–0.929, FreeBayes: 0.905–0.928), with low FDR for all callers. The late-relapse samples 3.1 and 3.2 showed slightly better recall, likely reflecting the higher clonal SNV counts at that stage. Recall of subclonal variants at high frequency ranged in 0.639–0.755 for Strelka2, 0.497–0.701 for Mutect2, and 0.350–0.555 for FreeBayes, with the best performance achieved consistently in the pre-treatment and first-relapse samples. Germline VAF correlation was relatively large (Strelka: 0.903, HaplotypeCaller: 0.870, FreeBayes: 0.868). For CNA inference, the low-FGA profile produced excellent results across all samples and callers: ASCAT and Battenberg both achieved purity errors of $\delta\pi \in 0.002–0.005$ and near-perfect proportion of correctly inferred clonal CNAs ($\rho_{clonal} = 0.999$) on all four available samples; ASCAT showed higher breakpoint recall (0.875–0.959) while Battenberg had slightly lower recall (0.739–0.834); Sequenza showed somewhat higher purity errors ($\delta\pi \in 0.036–0.103$) but good proportion of correctly inferred clonal CNAs ($\rho_{clonal} \in 0.962–0.976$), making SPN06 the best-fitted SPN for purity estimation in the cohort.

###### *C.2.7. SPN07*

SPN07 comprises five samples across two time points: three pre-treatment samples (1.1, 1.2, 1.3) with 44 000–48 000 SNVs each, and two relapse samples (2.1, 2.2) dominated by the MSH6-driven hypermutant resistant clone with 449 000 SNVs each. All samples share the same high-FGA copy-number profile (FGA 50%, ploidy 2.5). The difference between pre-treatment and relapse samples was clearly reflected in somatic variant calling performance. The recall of clonal variants was consistently high across both time points and all callers (Strelka2: 0.917–0.921, Mutect2: 0.912–0.923, FreeBayes: 0.900–0.906), but the FDR diverged sharply: in the pre-treatment samples, the FDR was 0.295–0.388 with Mutect2, compared to Strelka2 ( 0.044–0.059) and FreeBayes ( 0.071–0.099); in the relapse samples Mutect2 FDR dropped to 0.025–0.084, comparably to the other callers, due to the dominant hypermutant clone providing a strong clonal signal. Recall of subclonal variants at high frequency was lower (Strelka2: 0.701–0.732, Mutect2: 0.631–0.753, FreeBayes: 0.428–0.514) across all samples. Germline VAF correlation was uniformly high (Strelka: 0.938, HaplotypeCaller: 0.937, FreeBayes: 0.936). For CNA inference, performance was remarkably consistent across all five samples given the uniform copy-number profile: ASCAT showed purity errors of $\delta\pi \in 0.064–0.066$ and proportion of correctly inferred clonal CNAs $\rho_{clonal} \in 0.639–0.644$ on all samples; Battenberg achieved near-zero purity errors ($\delta\pi \in 0.008–0.012$) and high proportion of correctly inferred clonal CNAs ($\rho_{clonal} \in 0.962–0.964$) consistently; Sequenza also performed well with purity errors $\delta\pi \in 0.013–0.018$ and proportion of correctly inferred clonal CNAs $\rho_{clonal} \in 0.841–0.908$, with the relapse sample 2.1 showing the highest proportion of correctly inferred clonal CNAs.

#### D. The nf-core/tumourevo analysis pipeline

nf-core/tumourevo is a novel bioinformatics pipeline to model tumour evolution from whole-genome sequencing (WGS) data. The pipeline performs state-of-the-art analyses of somatic mutations from tumour-normal matched sequencing assays, downstream of standard variant calling. The output of the workflow enables the reconstruction of the evolutionary processes leading to the observed tumour genome. All the analyses can be done at the level of single samples, multiple samples from the same patient (multi-region/longitudinal assays), and of multiple patients from distinct cohorts.

##### D.1. Workflow overview and inputs

A schematic overview of nf-core/tumourevo is shown in Supplementary Fig. S25. The workflow accepts somatic variant calls generated by widely used variant callers such as Mutect2[45], Strelka2[43], Platypus [50]. Copy number segmentation files produced by tools such as ASCAT[51], Sequenza[48], Battenberg[49] are incorporated to estimate the underlying tumour karyotype. The workflow automatically adapts to single-sample or multi-sample sequencing settings, enabling analysis of spatially distinct tumour regions or longitudinal samples collected over the course of disease progression or treatment. Many tumour evolution studies involve multi-region or longitudinal sequencing data, but variant calling is often performed independently for each sample. This can lead to incomplete mutation matrices. To address this, nf-core/tumourevo offers an optional "lifter" subworkflow that harmonises mutation information across samples from the same patient by performing targeted pile-ups of sample-specific mutations across the other samples. This recovers sequencing depth at loci where mutations are present in one sample but absent in others, providing all the necessary information for multivariate subclonal deconvolution models, and enabling more accurate inference of shared and sample-specific tumour subpopulations.

##### D.2. Driver annotation

As workflows for de novo driver annotation are already available as independent pipelines[52–54], nf-core/tumourevo focuses on providing a simple and effective mechanism for annotating driver mutations using a precompiled catalogue, which can be supplied by the user. Alterations that have a disruptive or effectiveness-altering effect on proteins, such as those assigned a 'HIGH' or 'MODERATE' impact by the Ensembl Variant Effect Predictor[55] (VEP), and affect a cancer gene from the chosen catalogue (IntOGen[52] by default), are identified as potential tumour driver alterations.

##### D.3. Quality control and filtering of genomic data

Reliable evolutionary inference depends on a thorough assessment of data quality and potential technical confounders. To tackle this, nf-core/tumourevo incorporates two quality control steps that collectively evaluate tumour purity estimates, copy number segmentation, and somatic mutation calls. Copy number segments are examined using CNAqc [56], which checks the consistency of observed variant allele frequencies (VAF) with the reported tumour purity and ploidy. Segments that fail these consistency checks are marked as low confidence and can optionally be excluded from further analysis. The pipeline also uses TINC [57] to estimate the proportion of tumour-derived reads in the normal sample (tumour-in-normal contamination). Significantly, this filtering step reduces the likelihood of false subclonal signals caused by inaccurate copy number estimates and high contamination levels, which may lead to false-negative mutation calls and thus act as a major confounder for subsequent evolutionary studies. By integrating these quality metrics early in the workflow, nf-core/tumourevo enables users to identify problematic samples, filter out unreliable genomic regions, and ensure that subsequent analyses are performed on high-confidence data.

##### D.4. Subclonal deconvolution and phylogenetic reconstruction

To determine the genetic composition of tumours, nf-core/tumourevo performs probabilistic clustering of somatic mutations based on their VAF across samples. The pipeline supports various clustering methods, including MOBSTER [58], PyClone-VI [59], and VIBER [58]. All these methods use beta-binomial mixture models to model read counts and estimate the cancer cell fraction (CCF) of mutations that distinguish distinct tumour cell populations. For datasets with multiple samples, multivariate clustering models exploit information across samples to identify shared subclonal populations. The resulting mutation clusters are then used to reconstruct clone trees with ctree [60] that align with the observed mutation accumulation patterns, revealing evolutionary trajectories.

##### D.5. Identification of mutational processes

Somatic mutation patterns reflect the activity of mutational processes such as exposure to carcinogens, defects in DNA repair pathways, or endogenous enzymatic activity. nf-core/tumourevo quantifies these processes through mutational signature analysis. Mutation counts are computed for single-nucleotide substitutions (SBS), dinucleotide substitutions (DBS), and small insertions or deletions (IDs) in their trinucleotide sequence contexts, and successively modelled with SparseSignatures [61] and SigProfiler [62], supporting both de novo signature extraction and assignment to reference signatures in the COSMIC catalogue.

##### D.6. Identification of mutational processes at clonal levels

To disentangle the mutational processes operating within distinct tumour cell populations, nf-core/tumourevo performs mutational signature analysis at the level of individual clones identified during subclonal deconvolution. Mutation counts are computed for each clone detected by VIBER or PyClone-VI, and signature assignment is carried out using the SigProfiler [62] assignment module to map dataset-level signatures to clone-specific mutations. This approach enables the characterisation of how mutational processes vary across the evolutionary history of a tumour, linking signature activity to specific subclonal populations and their inferred phylogenetic relationships.

##### D.7. Automated genome interpretation

The Genome Interpreter module integrates three complementary lines of evidence to compute a composite expansion score $\phi_C \in [0, 1]$ for each mutation cluster $C$ inferred in the subclonal deconvolution step.

The composite score $\Phi_C \in [0, 1]$ is defined as a weighted average of three complementary component scores, each capturing a distinct dimension of the evidence for a clonal expansion. The first component, $\phi_{\text{tail}} \in [0, 1]$, is defined as the proportion of mutations in cluster $C$ that are never labelled as `TAIL` across any of the univariate subclonal deconvolution runs. A high value of $\phi_{\text{tail}}$ therefore indicates that the mutations comprising the cluster are robustly supported as a genuine subclonal signal across samples, rather than being attributable to neutral evolutionary dynamics or sequencing noise. The second component, $\phi_{\text{signature}} \in [0, 1]$, captures the degree to which the mutational signature exposure profile of cluster $C$ diverges from both the clonal cluster and the whole-sample background. This divergence is quantified by combining cosine dissimilarity, computed as one minus the cosine similarity between exposure vectors, with the proportion of signatures that differ between the clonal cluster and $C$. Since SBS and ID profiles provide complementary and partially independent views of the underlying mutational processes, the final value of $\phi_{\text{signature}}$ is obtained by retaining the minimum cosine similarity and the maximum proportion of differing signatures across the two mutation types, adopting a conservative criterion that favours sensitivity in detecting any signature divergence. A high value of $\phi_{\text{signature}}$ therefore indicates that the mutational processes active during the expansion differ substantially from those that shaped the clonal background, which may reflect the acquisition of a novel mutagenic process associated with the expansion. The third component, $\phi_{\text{driver}} \in \{0, 1\}$, is a binary indicator equal to 1 if at least one driver mutation is annotated in cluster $C$, and 0 otherwise, providing a direct link between the inferred expansion and a functionally relevant selective event. The three components are then combined as:

$$\Phi_C = \frac{w_1 \cdot \phi_{\text{tail}} + w_2 \cdot \phi_{\text{signature}} + w_3 \cdot \phi_{\text{driver}}}{w_1 + w_2 + w_3} \tag{27}$$

where the weights $w_1, w_2, w_3 \in \{0, 1\}$ act as binary switches that allow the user to include or exclude each component based on the biological context of the analysis and the availability of the required data. This modular design ensures that $\Phi_C$ can be tailored to different analytical scenarios, ranging from a single-component heuristic, for instance relying solely on driver annotation when signature data are unavailable, to a fully integrated three-component score. Based on the continuous index $\Phi_C$, each cluster is assigned to one of four ordered expansion tiers:

$$\text{Tier}(C) = \begin{cases} \text{Tier 1} & \text{if } (C \text{ is the clonal cluster}) \vee (\Phi_C \geq 0.9), \\ \text{Tier 2} & \text{if } 0.55 \leq \Phi_C < 0.9, \\ \text{Tier 3} & \text{if } 0.20 \leq \Phi_C < 0.55, \\ \text{Tier 4} & \text{if } \Phi_C < 0.20. \end{cases} \quad (28)$$

ranging from Tier 1, which encompasses the founding clonal cluster as well as subclonal expansions supported by the strongest evidence ($\Phi_C \geq 0.9$), to Tier 4, which groups clusters with no appreciable evidence of a biologically meaningful expansion ($\Phi_C < 0.20$).

##### D.8. Portability and reproducibility

The pipeline is built using Nextflow[63, 64], a workflow tool to run tasks across multiple compute infrastructures in a very portable manner. It comes with docker containers, singularity images and biocontainers that make the installation process trivial and ensure highly reproducible results. The Nextflow DSL2 implementation of this pipeline, that uses one container per process, makes maintenance and software updating easy. All the relevant processes have been implemented as independent nf-core/modules, making them available to any other nf-core pipeline, and to everyone within the Nextflow community.

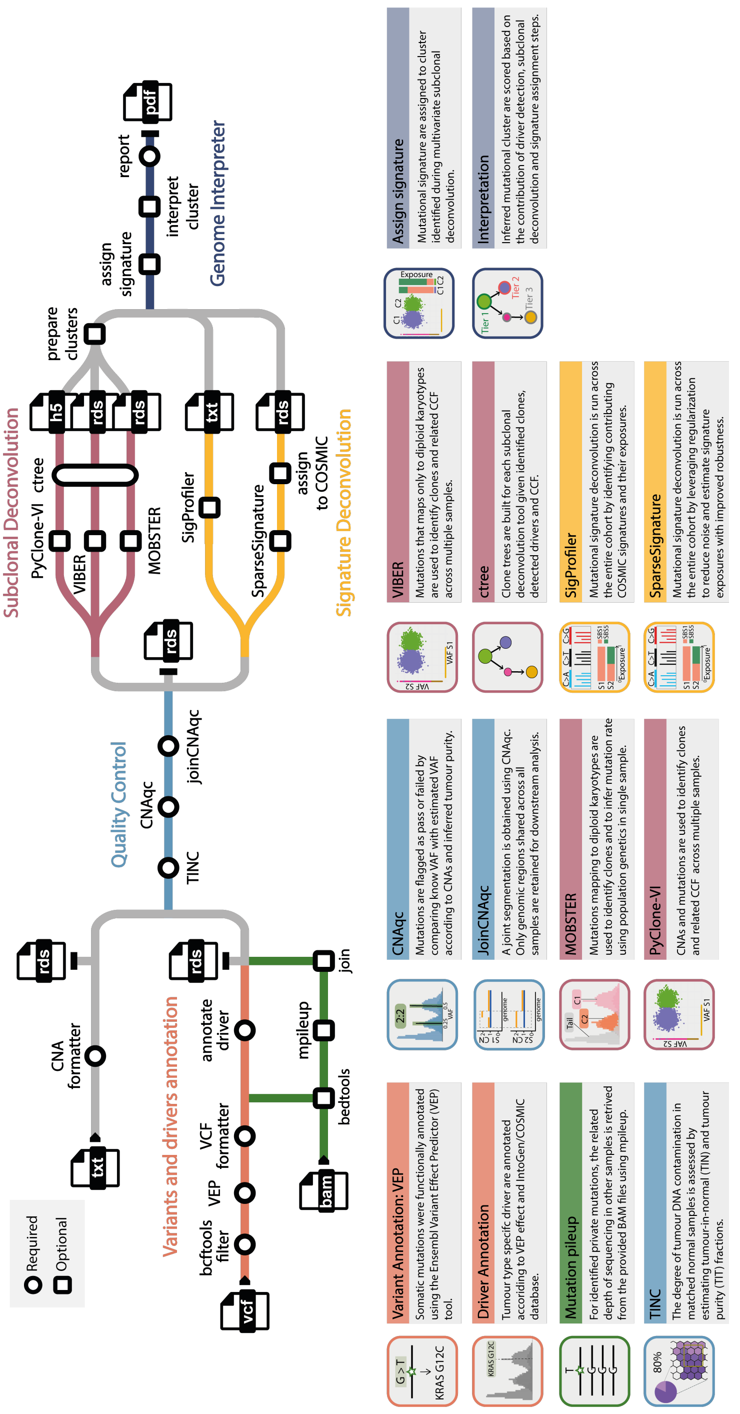

**Supplementary Fig. S25.** nf-core/tumourevo **workflow.** A schematic overview including some of the main analysis software implemented in the nf-core/tumourevo workflow

.

### E. Benchmarking of the SCOUT cohort with nf-core/tumourevo

We benchmarked the nf-core/tumourevo pipeline by evaluating its ability to accurately reconstruct driver mutations, quality-control variant calls, deconvolve mutational signatures, infer subclonal architectures, and interpret clonal expansions, all against the molecular ground truth provided by the ProCESS simulations. Results are organised in two parts: a cohort-level analysis summarising performance trends across all SPNs, purity conditions, and coverage levels, followed by per-SPN results that integrate all modules into a unified description for each simulated patient.

#### E.1. Cohort results

##### *E.1.1. Quality control*

Quality control of variant and copy-number calls is an essential step before any downstream evolutionary analysis. Within nf-core/tumourevo, two dedicated tools are applied at this stage: CNAqc [65] and TINC [57].

**CNAqc** CNAqc assesses the internal consistency between somatic variant allele frequencies (VAFs) and the inferred allele-specific copy-number profile by testing whether the observed VAF peaks match the theoretical positions expected under the estimated tumour purity and ploidy. For each segment, the expected VAF of a heterozygous mutation is a deterministic function of the major and minor allele copy numbers and tumour purity; CNAqc uses a peak-fitting procedure to quantify the deviation between observed and expected VAF distributions, assigning a quality label to each sample. We reported CNAqc final state, PASS or FAIL, for each sample, across all purities and coverage combinations, as well as copy number (CN) and somatic variant callers combinations. The accuracy of CNAqc is strictly influenced by CN caller performance, as previously reported in Supplementary Fig. S24. Results are summarised in Supplementary Fig. S26 and Supplementary Table ST9E.

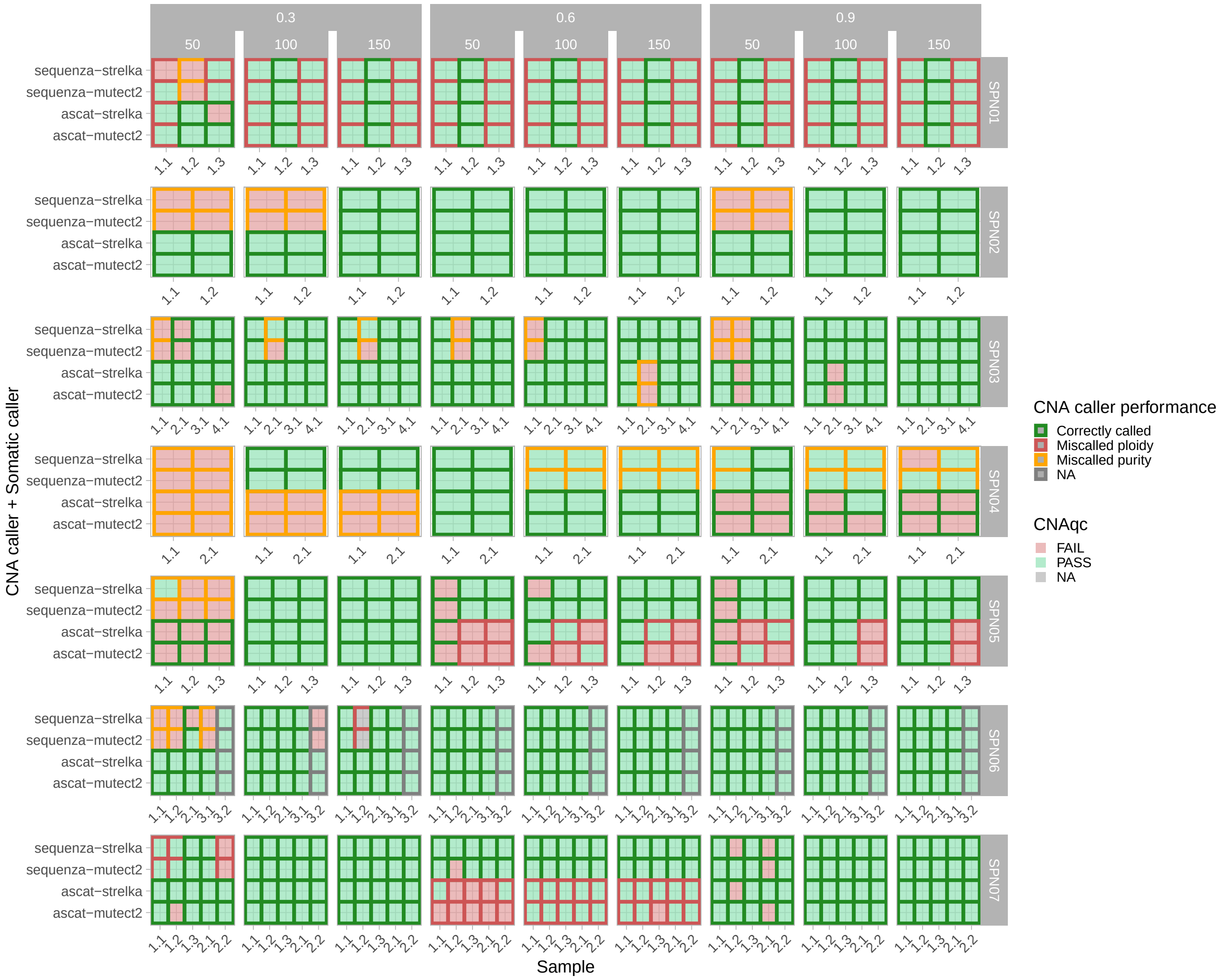

**Supplementary Fig. S26. Overview of SCOUT CNAqc benchmarking.** Boxes are colored according to CNAqc performance, while the contour color indicates the performance of the CNA caller (rows), evaluated in terms of purity and total ploidy.

**TINC** TINC (Tumour-in-Normal Contamination) detects and quantifies the presence of tumour-derived DNA in the matched normal sample, a common source of artefactual somatic calls that can bias downstream analyses. For each sample, TINC estimates a contamination score by modelling the allele frequency distribution of heterozygous germline variants in the normal sample: contamination shifts these frequencies away from the expected 0.5, and TINC uses a mixture model to infer the contamination fraction. To evaluate TINC, we compared its TIT (tumour-in-tumour) and TIN (tumour-in-normal) contamination estimates against the ground truth encoded in the ProCESS simulations. For TIT, we computed the absolute error between the TINC estimate and the simulated purity values across all purity, coverage, and caller configurations. For TIN, since ProCESS simulates no normal contamination, estimates are expected to be $\approx 0$. Results are summarised in Supplementary Fig. S27 and Supplementary Table ST9F.

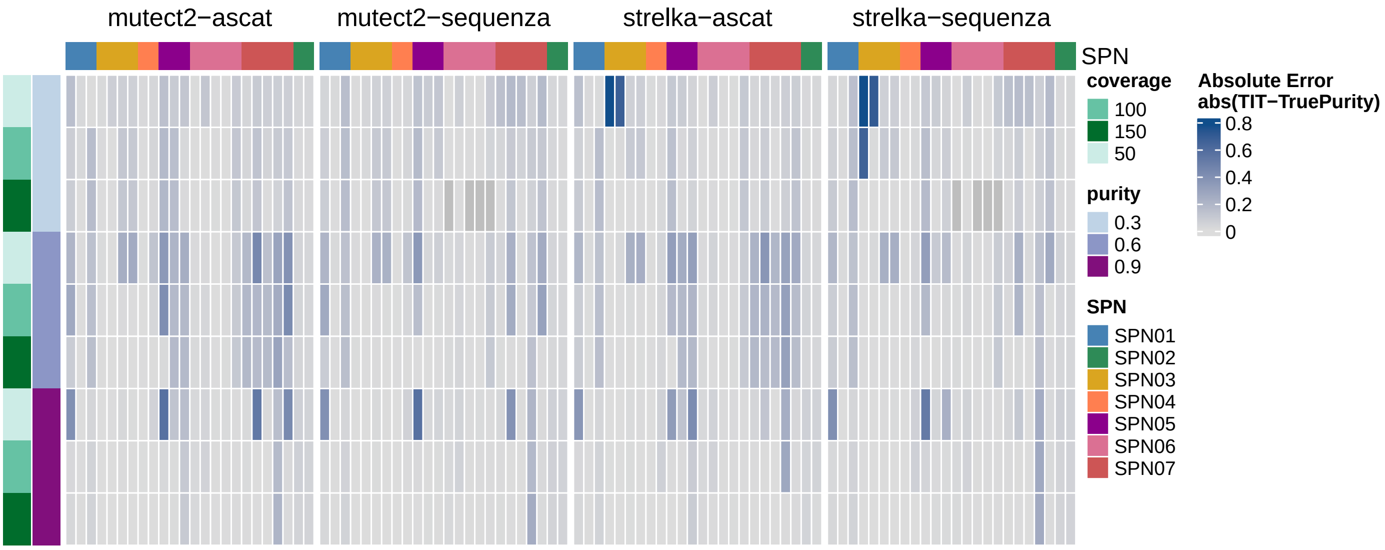

**Supplementary Fig. S27. Overview of SCOUT TINC benchmarking.** Heatmap showing the absolute error between TIT TINC estimate and simulated true purity for each combination of mutation and copy number caller (mutetct2-ascat, mutetct2-sequenza, strelka-ascat, strelka-sequenza), purity (0.3, 0.6, 0.9) and coverage (50×, 100×, 150×) for each SPN.

##### *E.1.2. Driver detection*

Driver identification performance was evaluated using standard binary classification metrics, Precision, Recall, and F1, comparing the set of driver mutations inferred by the nf-core/tumourevo annotation module against the ground-truth driver events encoded in the ProCESS simulations. True positives (TP) are simulated driver mutations correctly identified by the pipeline; false negatives (FN) are simulated drivers that were not recovered; and false positives (FP) are variants annotated as drivers in genomic regions where no driver was simulated. All metrics were computed in the multisample setting, yielding a single performance value per SPN rather than per individual sample. To assess whether performance varies across mutation rate regimes, SPNs were stratified into hypermutant ($\mu \geq 1 \times 10^{-7}$; SPN02, SPN05, SPN07) and non-hypermutant ($\mu < 1 \times 10^{-7}$; SPN01, SPN03, SPN04, SPN06) categories. The evaluation was repeated across all combinations of somatic variant caller and CNA caller, as well as across all purity and coverage conditions, to characterise how upstream calling choices propagate to driver annotation performance.

Non-hypermutant SPNs (SPN01, SPN03, SPN04, SPN06) achieved a cohort mean F1 of 0.805, with recall of 0.891 and precision of 0.771, indicating reliable recovery of true drivers with a modest rate of false positives. Hypermutant SPNs (SPN02, SPN05, SPN07) showed similarly high recall (0.822), as the true driver mutations are still detectable, but precision collapsed to 0.242 and F1 to 0.356. This degradation reflects the elevated mutation burden increasing the probability that passenger mutations fall within known driver genes, inflating false-positive driver calls regardless of caller combination, purity, or coverage. The finding confirms that driver annotation in hypermutant contexts is a fundamental biological challenge rather than an artefact of any specific upstream calling strategy. Full results are reported in Supplementary Fig. S28 and Supplementary Table ST9D.

##### *E.1.3. Mutational signature deconvolution*

For each signature deconvolution tool, SigProfiler [66] and BASCULE [67], we benchmarked its ability to accurately identify the simulated mutational signatures active in each sample, both in the SBS96 and ID83 contexts, and to correctly estimate their corresponding exposures. The evaluation was structured around two complementary tasks: signature detection, which assesses whether the correct set of signatures is identified, and exposure estimation, which assesses whether the inferred contribution of each signature to the overall mutation catalogue is accurate.

For signature detection, the set of signatures inferred by each tool was compared against the known ground-truth signatures from the ProCESS simulation for each sample. True positives (TP) were defined as signatures present in both the ground truth

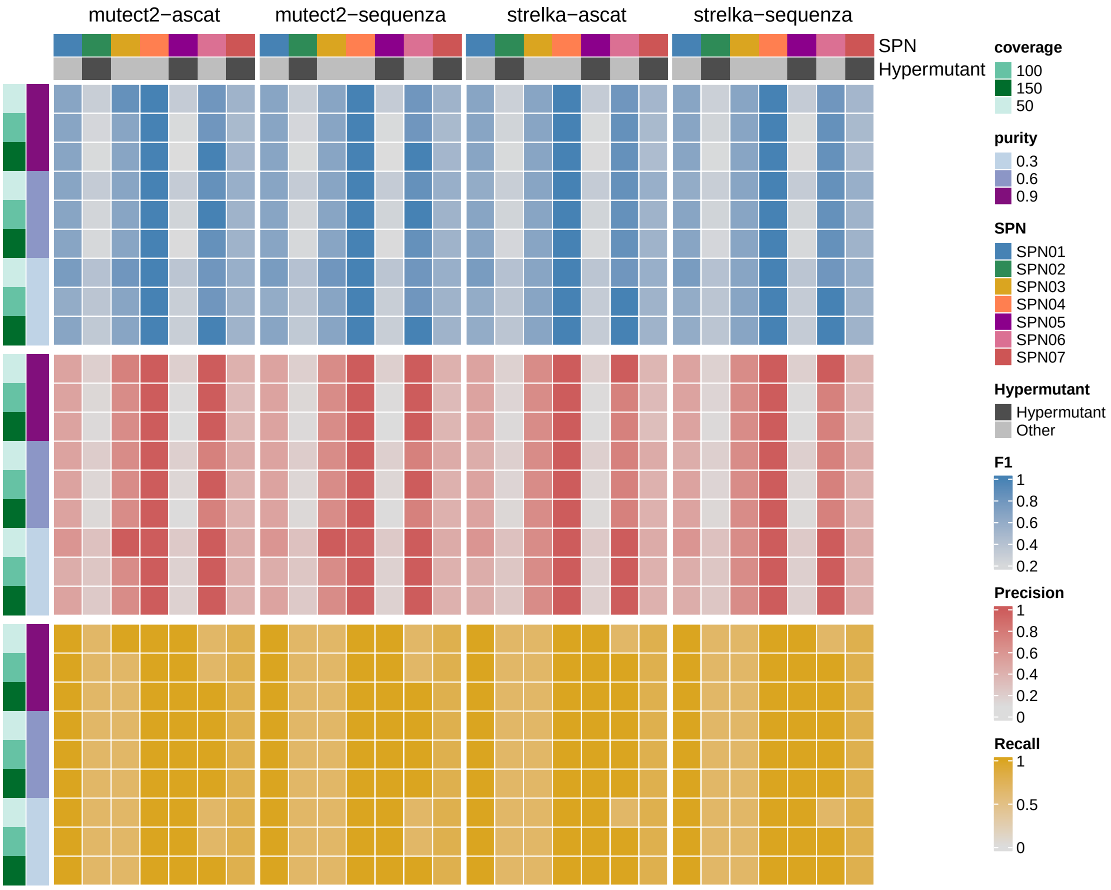

**Supplementary Fig. S28. Overview of SCOUT Driver Annotation benchmarking**. Heatmap showing the F1 score, Precision and Recall for driver detection for each combination of mutation and copy number caller (mutetct2-ascat, mutetct2-sequenza, strelka-ascat, strelka-sequenza), purity (0.3, 0.6, 0.9) and coverage (50×, 100×, 150×).

and the inferred solution; false positives (FP) as signatures inferred by the tool but absent from the ground truth; and false negatives (FN) as ground-truth signatures not recovered by the tool. These counts were used to compute recall and precision, allowing us to assess separately the sensitivity of each tool in recovering all active signatures and its specificity in avoiding the inference of spurious ones.

For exposure estimation, we computed two complementary metrics between the ground-truth exposure vector $\mathbf{x}$ and the inferred exposure vector $\mathbf{y}$. The first is cosine similarity, which captures the directional agreement between the two vectors regardless of their magnitude:

$$\text{cosine}(\mathbf{x}, \mathbf{y}) = \frac{\mathbf{x} \cdot \mathbf{y}}{\|\mathbf{x}\|_2 \, \|\mathbf{y}\|_2} = \frac{\sum_{i=1}^{n} x_i y_i}{\sqrt{\sum_{i=1}^{n} x_i^2} \sqrt{\sum_{i=1}^{n} y_i^2}} \tag{29}$$

A cosine similarity of 1 indicates perfect agreement in the relative composition of the exposure profile, while lower values reflect increasing divergence in the inferred signature contributions. The second metric is the mean squared error (MSE) between the ground-truth and inferred exposures, which penalises absolute deviations and is therefore sensitive to both the direction and magnitude of errors. The MSE was computed for each inferred exposure and summarised as the average MSE per sample.

To evaluate the potential confounding effect of subclonal mutational signatures on whole-sample deconvolution, samples were stratified into high- and low-complexity groups based on the degree of similarity between the signature compositions of clonal and subclonal mutations. For each sample, the set of mutational signatures assigned to clonal clusters and the set assigned to subclonal clusters were compared using the Jaccard similarity index. Values of $J$ approaching 1 indicate that the same mutational processes are active throughout tumour evolution; such samples were classified as low-complexity. Samples with $J \leq 0.9$ were classified as high-complexity, reflecting the emergence of distinct mutational processes associated with subclonal expansion. The Jaccard index was averaged across all coverage–purity combinations and computed separately for SBS and ID mutation types using the ProCESS ground truth data.

Across the cohort, both tools achieved high cosine similarity in low-complexity samples, where the same signatures are active in clonal and subclonal mutations, and lower similarity in high-complexity samples where specific signature are present only in subclonal mutations. For SigProfiler, mean SBS96 cosine similarity was 0.987 in low-complexity and 0.931 in high-complexity samples; ID83 followed a similar pattern (0.982 vs. 0.903). BASCULE performed comparably in SBS96 (0.985 low, 0.933 high) but showed a larger drop in ID83 (0.960 low, 0.873 high), indicating greater sensitivity to subclonal indel signature complexity. The degradation in high-complexity samples is attributable to signatures confined to subclonal populations whose contribution is diluted in the whole-sample mutation catalogue, making them difficult to separate. Full results, stratified by tool, mutation type, complexity class, purity, and coverage, are reported in Supplementary Fig. S29 and Supplementary Table ST9G.

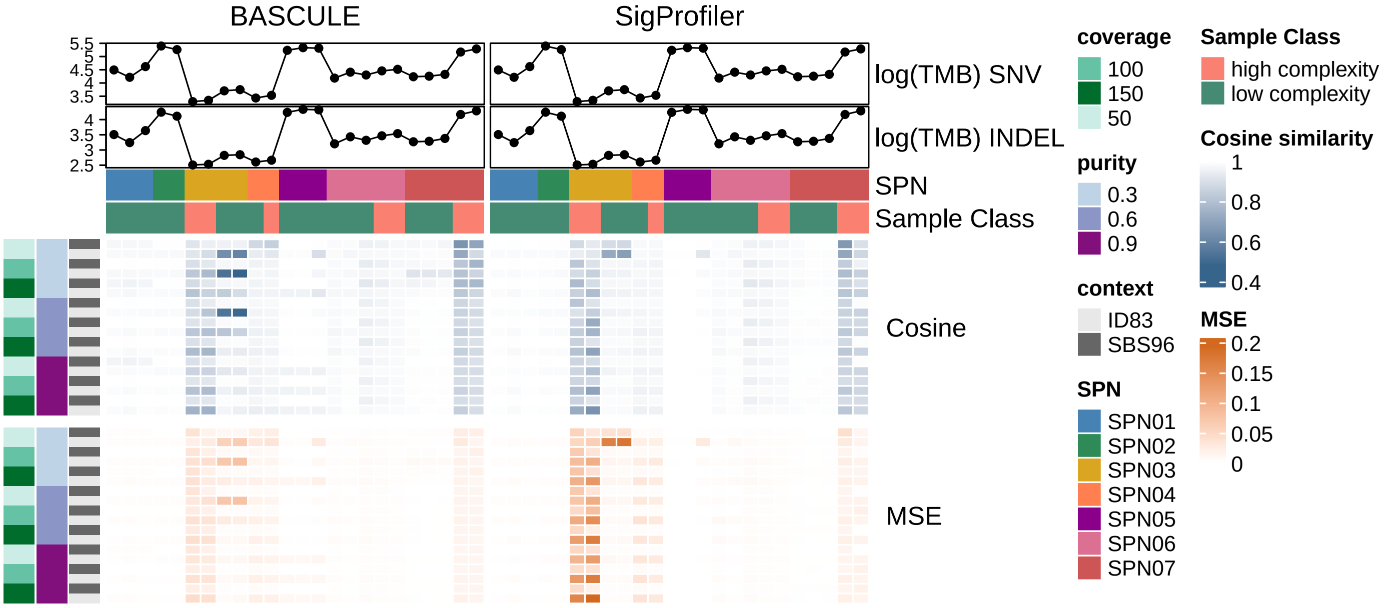

**Supplementary Fig. S29. Overview of SCOUT signature deconvolution benchmarking**. Heatmap of cosine similarity and MSE between inferred and simulated mutational signature exposures across all combinations of mutation and copy number caller (Mutect2–ASCAT, Mutect2–Sequenza, Strelka–ASCAT, Strelka–Sequenza), purity (0.3, 0.6, 0.9), and coverage (50×, 100×, 150×) for each SPN. Results are stratified by SBS and ID signatures, with samples annotated by complexity class (high or low). Log-transformed tumour mutational burden (TMB) for SNVs and INDELs is displayed above the heatmap.

##### *E.1.4. Subclonal deconvolution*

**Univariate methods** Univariate subclonal deconvolution methods were evaluated on their ability to detect subclonal structure at the single-sample level using MOBSTER [68]. Each sample was annotated according to the presence or absence of at least one subclone in the ProCESS ground truth, and accordingly classified as monoclonal (no subclonal expansion present) or polyclonal (at least one subclonal expansion present). Within each of these two broad categories, samples were further stratified into four classes reflecting distinct biological and technical conditions: monoclonal without sampling bias (monoclonal samples without any spatial confounder), with sampling bias (monoclonal samples in which spatial sampling bias may introduce artefactual VAF heterogeneity, mimicking spurious subclonal signals) and polyclonal strong evidence (polyclonal samples with clear subclonal structure), and weak evidence (polyclonal samples in which subclonal populations produce only weak VAF signals, for instance due to low cancer cell fraction, making them harder to distinguish from sequencing noise). The evaluation was deliberately restricted to this binary detection task, whether any subclonal structure is present at all, without assessing the accuracy of the inferred number of subclonal clusters or their specific cellular prevalences. This design choice reflects the practical importance of reliable subclone detection as a prerequisite for any downstream inference.

To quantify detection performance within each category, we defined true positives (TP) and false negatives (FN) as follows. For monoclonal samples, a TP is a sample correctly classified as monoclonal (i.e., no spurious subclones are inferred), while a FN is a monoclonal sample incorrectly classified as polyclonal (i.e., at least one subclone is erroneously detected). Conversely, for polyclonal samples, a TP is a sample correctly classified as polyclonal (i.e., at least one subclone is successfully identified), while a FN is a polyclonal sample incorrectly classified as monoclonal (i.e., existing subclonal structure goes undetected). In both cases, recall was used as the primary performance metric, as it directly captures the method's sensitivity to the relevant signal in each category: absence of subclones in the monoclonal case, and presence of subclones in the polyclonal case. Recall was computed as a function of increasing tumour purity and stratified across the four sample classes.

Across the cohort, recall for strong evidence polyclonal samples (CCF well separated from noise) was 0.629, and for the weaker-evidence polyclonal class 0.611, confirming that MOBSTER can detect most subclonal structure when the signal is present. The main source of error was in the opposite direction: monoclonal samples were systematically over-called as polyclonal. Monoclonal samples without sampling bias were correctly classified (as monoclonal) only 58% of the time (specificity 0.582), and samples with spatial sampling bias showed similar over-calling (specificity 0.577), indicating that spatial mutation patterning within a single clone generates VAF heterogeneity that mimics subclonal signals and confounds univariate deconvolution regardless of whether sampling bias is formally present. These patterns were consistent across all caller combinations and coverage conditions. Full per-class recall stratified by purity, coverage, and caller is reported in Supplementary Fig. S30 and Supplementary Table ST9H.

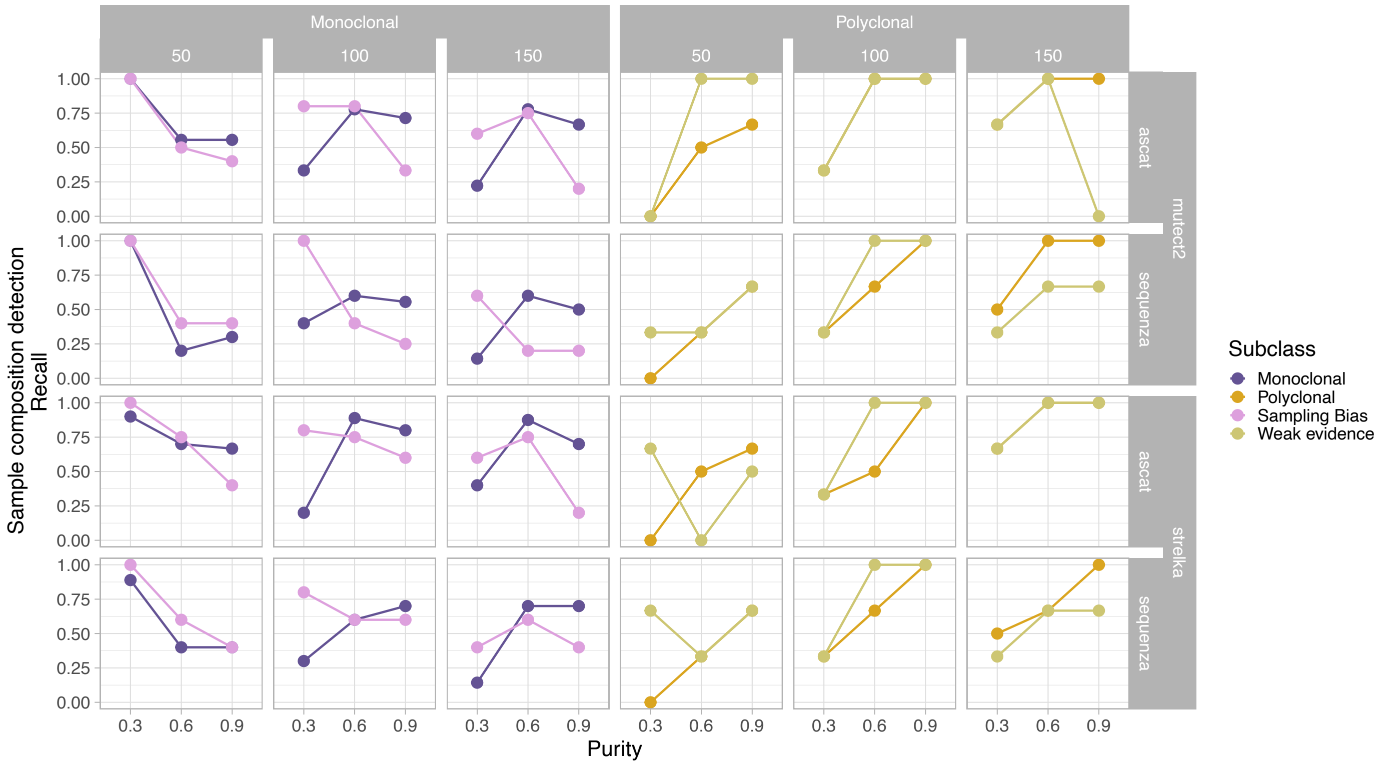

**Supplementary Fig. S30. Overview of SCOUT univariate subclonal deconvolution benchmarking**. Recall for the detection of monoclonal and polyclonal samples inferred by MOBSTER, stratified by tumour purity, sequencing coverage, somatic mutation calling tool, and copy number calling tool. For each stratification variable, recall is reported separately for monoclonal and polyclonal samples, and samples are also stratified by the bias for which the ground truth simulation is affected (Monoclonal-sampling bias or Polyclonal-weak evidence).

**Multivariate Methods** We further evaluated the multivariate subclonal deconvolution methods VIBER [68] and PyClone-VI [59], which jointly analyse mutation data across multiple samples from the same tumour. For these methods, no ground-truth comparison was performed; instead, we characterised the complexity of the inferred clonal architectures by computing the ratio of the number of inferred clusters to the number of samples, a normalisation that accounts for the fact that the expected complexity of the clonal architecture scales with the number of available samples. This ratio serves as a proxy for how much subclonal structure each tool infers relative to the amount of available samples, and enables comparison across SPNs of varying size. This ratio was compared across SPNs with spatial sampling bias (SPN02, SPN03, SPN04) and without (SPN01, SPN05, SPN06, SPN07). Spatial sampling bias arises when the physical locations of biopsies do not represent the full clonal diversity of the tumour, for example when spatially proximate samples are over-represented relative to more distant regions harbouring distinct subclones. By stratifying results according to the presence or absence of such bias, we aimed to assess the extent to which spatial distribution of samples affects the ability of multivariate methods to recover the true clonal architecture, and the additional effect of tumour purity on this task.

SPNs without sampling bias showed cluster-to-sample ratios of 1.41–2.40× for VIBER and 2.49–3.58× for PyClone-VI. SPNs with spatial sampling bias showed higher ratios of 1.58–3.03× for VIBER and 2.69–4.14× for PyClone-VI, with the most complex solutions inferred for SPN02 (3.03× and 4.14×), where two biopsies capture spatially segregated mutations within a single dominant clone. PyClone-VI consistently inferred more clusters than VIBER across all SPNs. Both tools inferred more complex solutions at lower purity, where wider VAF distributions promote spurious clusters inference. Full results stratified by SPN, purity, coverage, and caller combination are reported in Supplementary Fig. S31 and Supplementary Table ST9I.

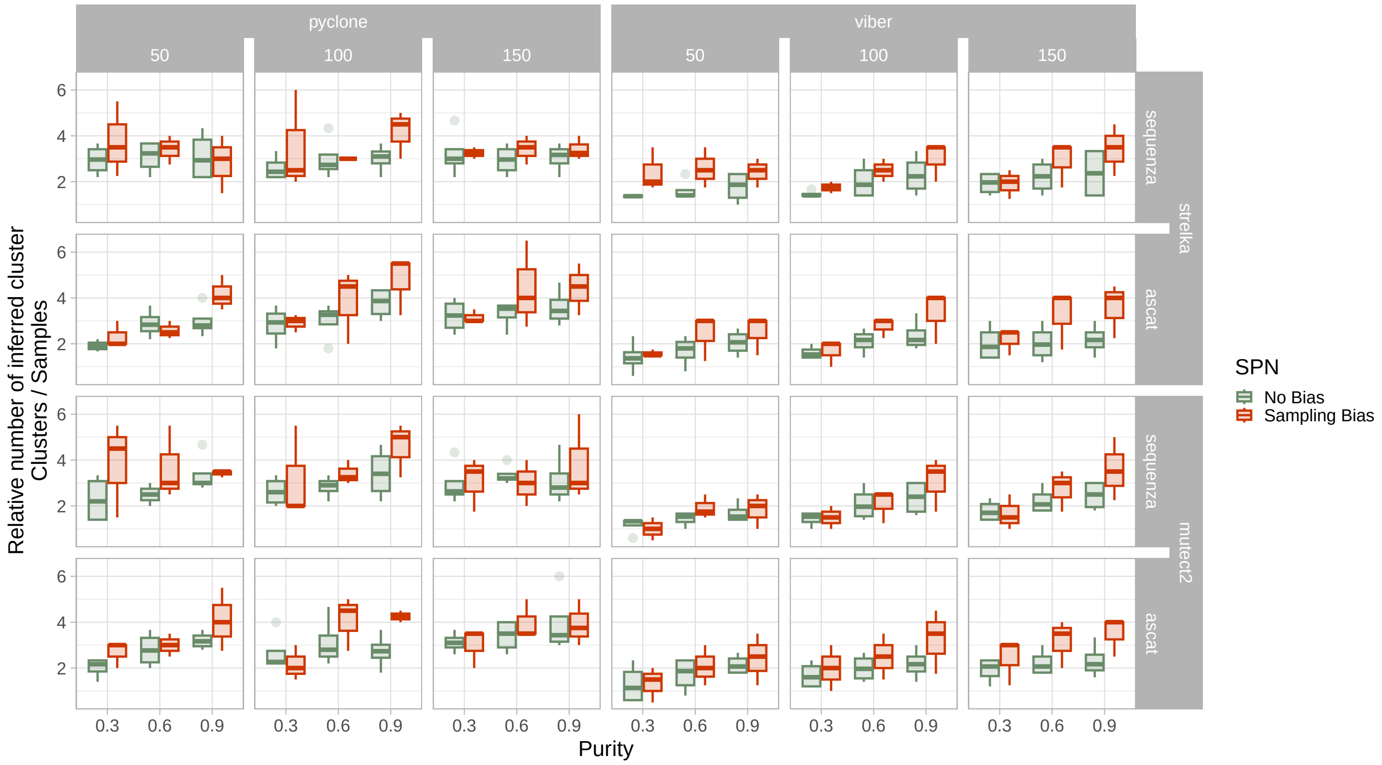

**Supplementary Fig. S31. Overview of SCOUT multivariate subclonal deconvolution benchmarking**. Ratio of the number of inferred clonal clusters to the number of samples per simulated patient (SPN), reported for PyClone and VIBER, stratified by tumour purity, sequencing coverage, somatic mutation calling tool, copy number calling tool, and presence or absence of spatial sampling bias.

##### *E.1.5. Genome interpreter*

The Genome Interpreter module of nf-core/tumourevo assigns a composite score $\Phi_C \in [0, 1]$ to each inferred mutation cluster $C$, integrating three sources of evidence: the robustness of the cluster's mutational content with respect to univariate subclonal deconvolution ($\phi_{\text{tail}}$), the divergence of its mutational signature exposure profile relative to the clonal cluster and the whole-sample background ($\phi_{\text{signature}}$), and the presence of at least one annotated driver mutation ($\phi_{\text{driver}}$). Based on $\Phi_C$, each cluster is assigned to one of four ordered expansion tiers, ranging from Tier 1 (clonal cluster or $\Phi_C \geq 0.9$) to Tier 4 ($\Phi_C < 0.20$), reflecting decreasing evidence of a biologically meaningful subclonal expansion.

To benchmark the Genome Interpreter, each mutation was annotated with two independent Tier labels: the Tier assigned by the nf-core/tumourevo pipeline to the cluster in which the mutation was detected, and the corresponding Tier derived from the ProCESS ground-truth simulation. Tier membership was then treated as a binary classification problem at the mutation level: for a given Tier, mutations belonging to that Tier in the ground truth were considered positive instances and all remaining mutations negative instances. Recall, precision and accuracy were computed for each Tier, quantifying how accurately individual mutations are assigned to their correct expansion evidence class. An analogous evaluation was performed at the cluster level: ground-truth clusters were matched to inferred clusters on the basis of shared driver mutations, and Tier-level recall and precision were computed over matched cluster pairs, assessing the schema's ability to correctly characterise entire groups of co-occurring mutations.

At the mutation level, Tier 1 (clonal) assignment was highly accurate across all purity conditions: recall was 0.887 at purity 0.3, rising to 0.973 at purity 0.9, with precision consistently above 0.88. Tier 2 recall was more variable and did not scale monotonically with purity (0.729 at purity 0.3, dropping to 0.597 at purity 0.6 before recovering to 0.761 at purity 0.9), reflecting the sensitivity of intermediate-tier assignment to the precise cluster boundaries inferred by upstream deconvolution. Tier 3 recall was substantially lower and strongly purity-dependent: 0.071 at purity 0.3, improving to 0.531 at purity 0.9, consistent with the difficulty of resolving weak subclonal expansions at low tumour content. Tier 4 recall remained very low across all purity conditions (0.022–0.063), as mutations with minimal expansion evidence are the hardest to assign correctly. At the cluster level, Tier 1 precision and recall were 0.875 and 0.918 respectively; Tier 2 showed lower precision (0.365) due to merging of inferred clusters; Tier 3 achieved high precision (0.909) but lower recall (0.552), indicating that when a Tier 3 cluster is identified it is usually correct but many true Tier 3 clusters are missed. Cluster-level and mutation-level evaluations were broadly concordant. Full accuracy per Tier, stratified by purity, coverage, and caller combination, are reported in Supplementary Fig. S32 and Supplementary Table ST9J-K.

#### E.2. Per-SPN results

The following subsections summarise per-SPN benchmarking results across all nf-core/tumourevo modules.

##### *E.2.1. SPN01*

Driver detection achieved perfect recall (1.000) with moderate precision (0.495), resulting in an F1 score of 0.661. The three true drivers were consistently recovered but approximately three false positives were detected across all conditions. CNAqc achieved a PASS rate of 0.963, with only four cases classified as FAIL, all occurring at relatively low simulated purity. Despite the overall strong performance, CNAqc was unable to detect miscalled ploidy in samples with WGD (1.1 and 1.3). Signature deconvolution was consistently accurate across tools: the average cosine similarity achieved for SBS96 and ID83 signatures, with respect to ground-truth profiles, was respectively 0.9986 and 0.9893 for SigProfiler, 0.9889 and 0.9852 for BASCULE. For univariate subclonal deconvolution, only sample 1.1 was considered, since 1.2 and 1.3 were WGD. Although 1.2 was monoclonal, a subclone was incorrectly inferred in 68% of the possible combinations of tools. Multivariate deconvolution inferred a mean of 7.2 clusters (VIBER) and 10.1 (PyClone-VI) against 3 samples (2.40 and 3.35 per sample, respectively), confirming a systematic inflation of cluster number estimates in the absence of true polyclonality.

##### *E.2.2. SPN02*

Driver detection in this SPN was impacted by the hypermutant background (POLE): 2/3 of the true drivers were recovered on average (recall 0.667), but a mean of approximately ten false positives resulted from inflation of passenger mutations in driver genes, producing low precision (0.182), and thus low F1 (0.280). POLE was not detected since it was missing from the list of known drivers used for annotation. CNAqc PASS rate was 0.833, while all CNAqc FAIL samples were associated with miscalled purity by Sequenza: in particular at low simulated purity, Sequenza overestimates it to 1, giving rise to unexpected VAF peak. Cosine similarity of inferred versus true signatures was excellent for both SigProfiler (SBS96: 0.9983, ID83: 0.9874) and BASCULE (SBS96 0.9986, ID83 0.9865), most likely due to the dominant activity of the well-characterised POLE signature and the low complexity of the overall genome architecture. Univariate deconvolution correctly classified all samples as monoclonal, except 1.2, most likely because of the strong simulated sampling bias. Multivariate deconvolution inferred a mean of 6.1 (VIBER) and 8.3 (PyClone-VI) clusters across 2 samples (3.03 and 4.14 per sample), resulting in the highest inflation ratio in the cohort, consistent with the confounding effect of high mutation rates in driver annotation and the designed sampling bias.

##### *E.2.3. SPN03*

Driver detection achieved a mean F1 of 0.679 (precision 0.688, recall 0.676), with approximately one false positive per condition on average. The primary source of missed detections was MAP2K1, which was not annotated as a driver due to its absence from the list of CLL driver genes in the catalogue. CNAqc PASS rate was 0.854 across all samples, with lower values for the first two time point samples, characterized by low mutation counts. Miscalled purity samples were corrected flagged as FAIL by CNAqc, despite few cases. CNAqc flagged as FAIL seven samples despite correct CN caller inference especially for sample SPN03_2.1. Signature deconvolution performance was lower than in other SPNs, reflecting the high complexity of the distribution of active mutational processes in this patient, in particular due to the presence of SBS5, SBS9, ID5 and ID2 only at very low frequencies in the first two time points. The average cosine similarity for SBS96 and ID83 was 0.9381 and 0.8989, respectively, with SigProfiler, 0.9610 and 0.8002 with BASCULE. The latter was the lowest achieved cosine similarity for ID83 signatures of the whole analysis. This degradation was primarily driven by two factors: (i) the low overall TMB, which drastically reduced the sample size down to less than required for statistical significance, and (ii) the accumulation of SBS5, SBS9, ID5 and ID2 mutations at low variant allele frequencies only in the early time points, making the corresponding mutational signals hard to distinguish from noise. For univariate deconvolution, both Polyclonal (recall 0.667) and Weak Evidence (recall 0.556) classes were only partially recovered; Monoclonal and Sampling Bias samples produced no true-positives, indicating consistent over-calling of subclonal clusters. Multivariate deconvolution inferred a mean of 6.3 clusters (VIBER) and 10.8 (PyClone-VI) across 4 samples (1.58 and 2.69 per sample), the lowest ratios in the cohort despite the sampling bias presence.

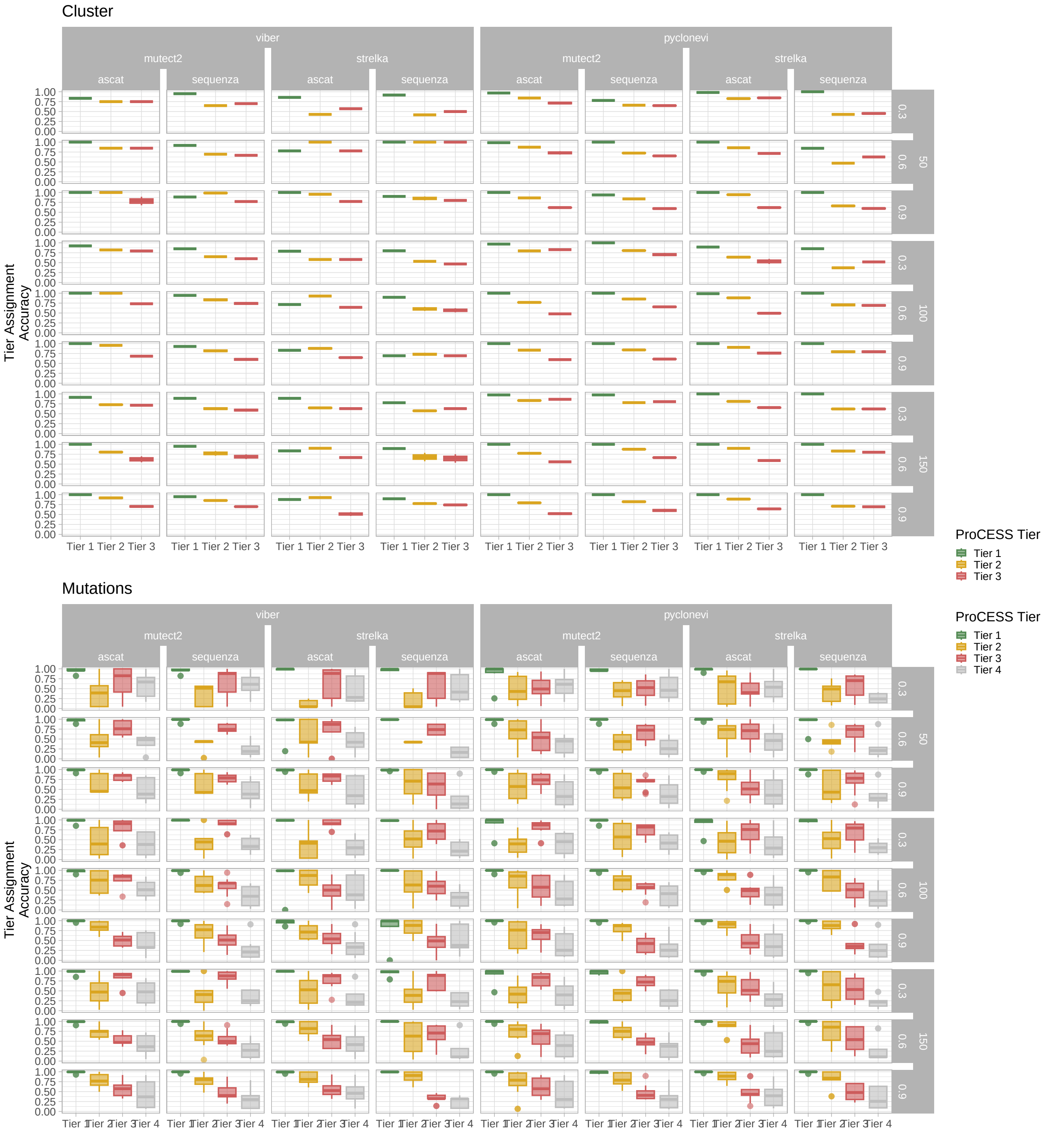

**Supplementary Fig. S32. Overview of SCOUT genome interpreter benchmarking**. Mutation- and cluster-level accuracy for assignment to the correct simulated expansion Tier, stratified by tumour purity, sequencing coverage, somatic mutation caller, and copy number caller. Mutation-level evaluation assessed the concordance between inferred and ground-truth Tier labels for individual mutations, while cluster-level evaluation compared matched inferred and simulated clusters based on shared driver mutations.

###### E.2.4. SPN04

Driver detection in this SPN was perfect: all two simulated drivers were recovered with no false positives across all conditions (F1, precision, recall 1.000). This was the best achievement concerning driver annotation, across the whole analysis. CNAqc achieved an overall PASS rate of 0.611, with performance strongly influenced by the choice of the CNA caller. ASCAT contributed most of the FAIL calls due to systematic purity misestimation across both high (0.9) and low (0.3) simulated purity, thereby reducing overall CNAqc performance. ASCAT tended to overestimate low purities, which CNAqc correctly flagged as FAIL. Conversely, CNAqc was more permissive on Sequenza inputs, returning PASS more frequently despite underestimated purity. This was likely due to imperfect modelling of tail mutations introducing spurious low-VAF peaks, and to the presence of the sampling bias further distorting the VAF distribution, all of which affected peak detection. Performance in signature deconvolution was intermediate: for SBS96 and ID83 signatures, the cosine similarity was respectively 0.9783 and 0.9669 with SigProfiler, 0.9621 and 0.9577 with BASCULE. The modest decrease relative to other SPNs was consistent with the low absolute mutation count in this SPN. Univariate deconvolution was affected by the spatial sampling bias in sample 2.1, where 31% of the tool combinations yielded spurious subclonal clusters, particularly at high purity. Noticeably, in the monoclonal sample 1.1, multiple subclones were inferred despite the absence of this bias, likely because of unassigned tail mutations incorrectly grouped into an additional subclone. This effect may have been driven by the low mutation rate, which limited the resolution of tail mutations. Multivariate deconvolution inferred a mean of 5.5 clusters (VIBER) and 7.4 (PyClone-VI) against 2 samples (2.75 and 3.71 per sample).

###### E.2.5. SPN05

Driver detection here was severely impaired by the hypermutant background (MMR deficiency): perfect recall (1.000) for the two true drivers was accompanied by a mean of approximately 12.5 false positives (precision 0.151, F1 0.260), the worst value of precision achieved in the whole analysis. CNAqc achieved a PASS rate of 0.667, with FAIL calls correctly tracking cases with poor purity and ploidy estimates. However, a consistent number of cases were incorrectly flagged as FAIL despite correct CN caller estimates ($\sim$ 0.44% of CNAqc FAIL samples); these were predominantly due to low simulated purity, particularly in 1.1, a polyclonal sample with weak evidence systematically assigned to subclonal clusters. Signature deconvolution performance in this SPN was the most accurate of the whole analysis: SigProfiler achieved 0.9997 cosine similarity for SBS96 and 0.9950 for ID83 signatures; BASCULE achieved 0.9997 and 0.9744 cosine similarity for SBS96 and ID83 signatures, respectively. The extremely good performance was primarily driven by the dominant and easily separable MMR-deficiency signature. Univariate deconvolution successfully recovered true subclones despite weak evidence in polyclonal samples (1.1 and 1.3) in nearly all cases at high purity. In contrast, the monoclonal sample 1.2 was fitted with multiple spurious subclonal clusters in 78% of tool combinations. The effect was most pronounced at higher purities. Univariate deconvolution recovered Weak Evidence polyclonal samples with recall 0.648; Monoclonal samples produced no TP calls. Multivariate deconvolution inferred a mean of 7.0 clusters (VIBER) and 10.8 (PyClone-VI) against 3 samples (2.32 and 3.58 per sample).

###### E.2.6. SPN06

Driver detection here achieved mean F1 of 0.878, precision 0.903, recall 0.889, with a mean of approximately 0.4 false positives per condition. CNAqc PASS rate was 0.939, the second highest in the analysis, consistent with the near-perfect CNA caller performance for this SPN. All the CNAqc FAIL cases were consistent with miscalled purity estimated of Sequenza copy number caller at low purity and coverage. Signature deconvolution performance was intermediate: cosine similarity of SBS96 and ID83 signatures was, respectively, 0.9713 and 0.9849 with SigProfiler, 0.9730 and 0.9812 with BASCULE. Performance in ID83 deconvolution was higher than for SBS96 for both tools, due to the prominence of strong signals from indel signatures related to APOBEC activity. Perfomance in univariate deconvolution was consistently robust across both monoclonal and polyclonal samples, with errors generally lower than average (16.1%) and driven by a common failure mode related to unresolved tail mutations. Deviations occurred only for low or intermediate purity across samples 1.1 (monoclonal; 9.7% miscalls), 1.2 (polyclonal; 16.1% missed subclones), 2.1 (monoclonal; 9.7% miscalls), and 3.1 (monoclonal; 16.1% miscalls). In monoclonal contexts, errors manifested as spurious subclonal clusters primarily driven by unassigned tail mutations, while in polyclonal samples they appeared mostly as missed subclones at low purity. Multivariate deconvolution inferred a mean of 7.5 clusters (VIBER) and 12.5 (PyClone-VI) against 5 samples (1.50× and 2.49×), the lowest absolute inflation ratio in the cohort for a No Bias SPN.

###### E.2.7. SPN07

Driver detection was partially impaired by the hypermutant relapse samples: the mean F1 was 0.526 (precision 0.393, recall 0.800), with all four true drivers recovered on average but a mean of approximately six false positives arising predominantly from the MSH6-driven relapse clone. Notably, MSH6 itself was not annotated as a driver due to its absence from the driver catalogue used by the pipeline. CNAqc PASS rate was 0.9, with most PASS calls corresponding to correctly called CNA profiles from both ASCAT and Sequenza (0.32 and 0.44, respectively). FAIL CNAqc calls were relatively infrequent in cases with correct CNA calls, while they were mainly correctly associated with miscalled ploidy. A small fraction of correctly called but low-confidence configurations also resulted in QC FAIL. Notably, CNAqc classified multiple miscalled ploidy cases as PASS ($\sim$ 15% of the CNAqc PASS samples). This suggested that spurious VAF peaks induced wither by sampling bias or by subclonal CNAs not resolved by the CNA caller allow certain ploidy errors to evade detection by the QC framework. Signature deconvolution yielded the lowest cosine similarities in the cohort, reflecting the high complexity of the mutational signature landscape driven primarily by the two relapse samples. The coexistence of treatment-associated signatures (SBS11, SBS25,

SBS26) with the hypermutant phenotype and a complex ID distribution creates a challenging deconvolution setting where pre-treatment and post-treatment mutational processes are active simultaneously across samples. SigProfiler achieved cosine similarity of 0.9524 for SBS96 signatures and 0.9322 for ID83 signatures; BASCULE reached cosine similarities of 0.9323 and 0.9342 for SBS96 and ID83 signatures, respectively. The performance was further degraded by the subclonal nature of the post-treatment signatures, whose contributions were diluted in the mutation count matrix and were therefore harder to resolve. Univariate deconvolution recovered Polyclonal samples with recall 0.296, the lowest in the whole analysis, while Monoclonal and Sampling Bias samples produced no TP calls; the bimodal mutation burden between pre-treatment and relapse samples likely confounded the monoclonal/polyclonal classifier. Multivariate deconvolution inferred a mean of 7.0 clusters (VIBER) and 12.8 (PyClone-VI) against 5 samples (1.41× and 2.56×), with PyClone-VI showing the highest absolute cluster count in the cohort.
